# BOTANIC-1: a series of long-context plant genomic foundation models in the agentic era

**DOI:** 10.64898/2026.09.04.749355

**Authors:** Amélie Barozet, Vincent Cabeli, Jean Ogier du Terrail, Alexey Rukhovich, Thomas Janssoone, Gary Klajer, Zeinab Sheikhitarghi, Gregory Andrews, Cyril Veran, Léonard Strouk

## Abstract

The development of climate-resilient crops would be greatly accelerated by models able to reason directly over plant genomic sequences and to pinpoint trait-associated regions or loci. Anticipating the impact of DNA base changes (variants) remains challenging, and understanding regulatory mechanisms is still an active area of research. Through self-supervised training on unannotated genomic data, genomic language models (gLMs) can learn DNA syntax and grammar that go beyond current annotations, thus complementing standard bioinformatics analyses that rely on rules established by decades of genomics research. Here we present our agent-powered *Model Factory* and its first outputs: the Botanic1 family of gLMs designed for plant research, which operates reliably on sequences from hundreds of base pairs up to 128 kbp. These models outperform all generalist and plant-specific gLMs (as well as specialised baselines) on one of the largest sets of plant genomics evaluation tasks reported to date, at a much smaller budget than concurrent models. Mechanistic interpretability analysis identifies features associated with biologically meaningful sequence properties including coding region boundaries and splice site motifs, demonstrating that these models are a source of biological insight beyond their benchmark performance. Finally, because a gLM only becomes practically useful when embedded in a broader workflow, we integrate Botanic1 as a specialised genomic layer callable by a generalist large language model (LLM) agent, illustrating how such hybrid systems could accelerate plant biology research. To support the plant genomics research community, we release the four Botanic1 models, their pre-training corpus and the trained sparse autoencoder for research use at https://huggingface.co/spaces/living-models/botanic1-report.

## 1 Introduction

Accelerating climate change is intensifying the search for crops that can adapt to increasingly adverse growing environments and withstand extreme events such as hail, drought, and heat stress. Crop pests and pathogens are expanding into higher latitudes due to the warming climate [1, 2], deployed resistance genes are frequently overcome within a few growing seasons [3], and anticipated yield gains remain insufficient to meet projected mid-century demand [4, 5]. All of these trends call for rapid action, and yet the breeding cycle remains slow, with the timeline from trait discovery to varietal release spanning up to eight years [6]. Plant genomics is complex, and translating trait-associated loci into mechanistic, causal understanding remains a bottleneck. Establishing which variants are causal and how they shape gene regulation and phenotype therefore still requires extensive experimentation in the field [7, 8].

Computational methods and analyses are natural and well-studied avenues to save time on crop research. Bioinformatics pipelines are thus employed to identify causal variants from genome-wide association studies (GWAS) and bulk segregant analysis (BSA) data [9, 10], but they face two limitations: first, they rely on extensive experimental data, which is expensive and slow to generate [11]; second, due to population biases such as linkage disequilibrium (LD), reliably separating causal variants from statistically correlated ones is a difficult problem, known as fine-mapping, whose resolution currently depends on fine genomic annotations and well-understood genomic regulation mechanisms [12].This problem is particularly acute in plants where genomic annotation data is mostly limited to reference genomes [13, 14], and genomic regulation mechanisms have been less studied than in humans and little mapped [15, 16, 17]. In other areas of biology, machine-learning (ML) models have overcome similar bottlenecks. AlphaFold-like models predict three-dimensional protein structures directly from a sequence in a few minutes or hours, where this would have required years of work from crystallographers less than a decade ago [18]. Protein language models pre-trained on sequence alone learn representations that transfer to functional annotation and variant effect prediction tasks [19, 20]. Finally, generative models based loosely on the same techniques and data produce binders that engage their intended targets in wet-lab experiments [21]. Considering these successful examples, it is plausible that strong plant genomic language models (gLMs) could address the limitations described above. Pre-trained on genomes spanning many species, they would learn general features of DNA sequence, from coding structure to regulatory grammar, that could transfer to untested genotypes or under-annotated species. They could draw on their internal representations of sequence constraints to help rank variants even in the absence of curated annotations, thus bypassing the indistinguishability caused by LD blocks. This raises the question of how to effectively use ML methods to construct such strong models.

The rigorous data-centric discipline behind the success of modern large language model (LLM) pre-training [22, 23] has seldom been applied to gLMs, although recent works such as Evo 2 [24] and NTv3 [25] have begun to address this gap. Similarly, a concurrent effort, MarinDNA [26] (released by Open Athena through the Marin collaboration [27]), showed on a small scale that careful data curation and hyperparameter tuning, rather than model size alone, are critical to performance on variant effect prediction (VEP) tasks. Its scope, however, is limited to mammalian genomes, 255 bp context windows, and single-variant scoring. Developed independently, our work extends similar conclusions at larger scale in the realm of plant genomics, through our first *agentic* model factory [28, 29].

Agents play a substantial role in the development of Botanic1 models: they help reliably operate the experimental infrastructure required to build new models. Built on agents, our *Model Factory* streamlines and industrialises a process that is typically carried out manually, and at smaller scale, within individual model development efforts: curating and versioning data, training control proxy models, evaluating competing design choices, and iterating on the resulting pre-training pipeline. We use this *Model Factory* to systematically ablate each component of the data pipeline under matched token, step, and epoch budgets, replicating experiments across seeds and data contexts, while coding agents launch experiments, terminate unpromising runs, monitor infrastructure health, and repair failures. This allows researchers to focus on experimental design and ideas while agents help with everything else. Thanks to this change of paradigm, gLM development turns from a sequence of bespoke experiments into a scalable search where, given enough compute, the limiting factor is human creativity. The first results of this *Model Factory* research process validate several heuristics emerging across the gLM literature while identifying additional avenues for predictive performance and training efficiency. Our *Model Factory* enables us to scale pre-training methodically on all axes while maintaining downstream performance. One of these scaling axes is species coverage, which to this day has been under-exploited in plants: earlier studies all trained on fewer than 100 species (42 plant genomes for PlantBiMoE [30], 48 crop genomes [31, 32], 65 for PlantCAD2 [33]), whereas the new *Model Factory* employs a 320-genome corpus spanning the land plants, thus yielding, to our knowledge, the broadest taxonomic coverage of any plant gLM to date.

We present the first models to emerge from this factory: Botanic1, a family of plant gLMs designed to make genomic sequence directly accessible to agentic systems. We evaluate Botanic1 across a broad set of regimes, from frozen representations and zero-shot log-likelihood to adaptation and long context, including cross-species transfer assessment. We rely as much as we can on established external plant genomics benchmarks, in particular the Plant Genomic Benchmark (PGB) [31] and the PlantCaduceus datasets suite [34], and we also introduce a new and harder zero-shot causal-variant discovery benchmark based on experimentally validated variants, totalling 545 loci across 14 species. On 22 frozen-model evaluation tasks, our four Botanic1 models occupy the top four positions in aggregate, ahead of every other model. Botanic1-S achieves this performance with fewer than half the trainable parameters of PlantCAD2-L [33] (318M versus 694M) and roughly 13× fewer pre-training tokens (314.6B versus ~ 4T), without hard-wiring reverse-complement equivariance into its architecture. Hard-wired reverse-complement equivariance is computationally heavy*^‡^* and less well understood than standard alternatives; in addition, it can be limiting for strand-specific prediction tasks, making the model blind to the biological signal. Beyond frozen-model evaluation, these gains persist when Botanic1 is adapted to new biological problems, either via full fine-tuning or via parameter-efficient adaptation with low-rank adaptation (LoRA) [35] or IA^3^ [36]. Adapted Botanic1-S is competitive or superior across diverse regulatory tasks and outperforms task-specific models trained from scratch where comparable baselines are available. In particular, adapted Botanic1-S surpasses a widely-used specialised ChromBPNet [37] baseline on a base-resolution chromatin accessibility prediction task, whereas the same Botanic1 architecture trained from scratch does not, thus emphasising the contribution of pre-training to Botanic1-S’s performance. To make Botanic1 able to understand long-range interactions, we further extend its accessible context from 8 kbp to 128 kbp. We first validate that this does not degrade short-context performance. Using synthetic data, we then perform nested-context and retrieval experiments to confirm that the extended models increasingly exploit information distributed across longer genomic intervals as context grows. We further assess these long-context capabilities on a real biological task: a chromatin accessibility task borrowed from PlantCAD2 [33], tested at several input context lengths. While both adapted PlantCAD2-S and Botanic1-S saturate before reaching 8 kbp context, we show that this saturation can be overcome by employing smart pooling approaches that leverage the full context while still focusing on a central window, improving the performance of Botanic1-S up to 32 kbp context.

The representations learned by Botanic1 models encode biologically meaningful structure. In silico mutagenesis experiments and sparse autoencoders (SAEs) [38, 39] recover interpretable features associated with identifiable genomic elements including regulatory motifs, gene coding structure, and genomic boundaries such as splice sites. These features provide insight into the signals underlying Botanic1’s predictions. While the above results on academic benchmarks are further validated by those new interpretability signals, turning those results into biological answers requires integrating gLMs with the analytical workflows biologists already use.

Although plant gLMs (and our Botanic1 models in particular) can solve problems where standard bioinformatics pipelines stumble, these more classical approaches can still be applied successfully to other problems. Even among gLMs themselves, no single model dominates across all tasks or meets all user constraints, which is one of the reasons why we choose to develop a family of models. The most efficient approach thus appears to be combining gLMs and bioinformatics tools, using each where it performs best. This could be made possible through the use of generalist agentic systems, which are rapidly permeating the scientific community and are already producing new science [40, 41]. Specifically in the domain of life sciences, initiatives like Anthropic’s Claude for Science and OpenAI’s GPT-Rosalind [42, 43, 44] recently showed that such systems can execute standard bioinformatic analyses more rapidly by combining their well-validated code-generation capabilities with dedicated bioinformatics harnesses. There is already substantial evidence that faithfully exposing non-text modalities to a model benefits from modality-specific representations: vision-language models bring images and video within reach of LLMs by projecting learned visual embeddings into the language model’s token space [45, 46], and audio-language and speech-text models integrate sound and real-time speech similarly through audio encoders [47, 48, 49, 50]. Another way to connect the specialised capabilities to generalist agent is at tool level: allowing it to reason over inputs and outputs of a dedicated model such as AlphaFold [18] for protein structure prediction. The plausibility of this approach was already shown in a recent preliminary research report [51], which used a generalist agent to orchestrate published structure-design, sequence-design, and co-folding AI models, thus producing candidate binders *in silico* whose validity was later confirmed by wet-lab experiments. The same architectural principles apply to plant genomics: an agent could call plant gLMs alongside conventional bioinformatics pipelines and use each where it is most appropriate. We illustrate this generalist-agent-based approach via a retrospective analysis of a published melon (*Cucumis melo*) sex-determination locus [52], where bulk-segregant analysis localises the causal mutation to a chromosome 2 region but leaves thousands of candidate variants unresolved, the causal base being later validated as a substitution in the ethylene-signalling gene <u>CmEIN3</u>. Given those candidates and no access to the publication, a generalist language-model agent (Gemma 4 [53]) ranks the causal variant no better than chance from sequence alone. With conventional bioinformatics tools, it reaches the limit of what current standard methods can do and yields a shortlist of variants that still require further experiment work. With Botanic1 callable as a tool, the causal variant is ranked first. The effect extends beyond this example: whereas standard annotations can only assign variants to discrete consequence classes, Botanic1 provides a complementary continuous measure of sequence constraint that enriches experimentally validated causal variants near the top of the ranking.

## 2 Results

### 2.1 Introduction

Rather than optimising a single model architecture at a fixed point in time, we develop the Botanic1 series within a *model factory* framework [28, 29] designed to support the continuous improvement of plant gLMs. This framework, inspired from LLM research [54, 55, 22, 56, 23, 57, 58], combines increasingly refined training data, additional training and evaluation signals, and systematic experimentation across model scales. All results reported here correspond to non-causal, encoder-only architectures trained on plant genomes using masked language modelling (MLM) (Figure 1a), although our codebase supports any data source and any model architecture (Sections 4.1 and 4.1.6). We optimise our evaluation pipeline so that it is lightweight enough to be used throughout training as an additional supervisory signal, more informative than train and test language log-likelihood. We define a proxy aggregate metric called *S*_bal_, which averages species’ scores within each of nine task families and then averages the families’ scores, and use its training variant 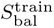 as the optimisation target (Section 4.3). We extensively test our models’ capabilities across diverse tasks and evaluation methods (Figure 1b) and at different context sizes (Figure 1c).

**Figure 1.**
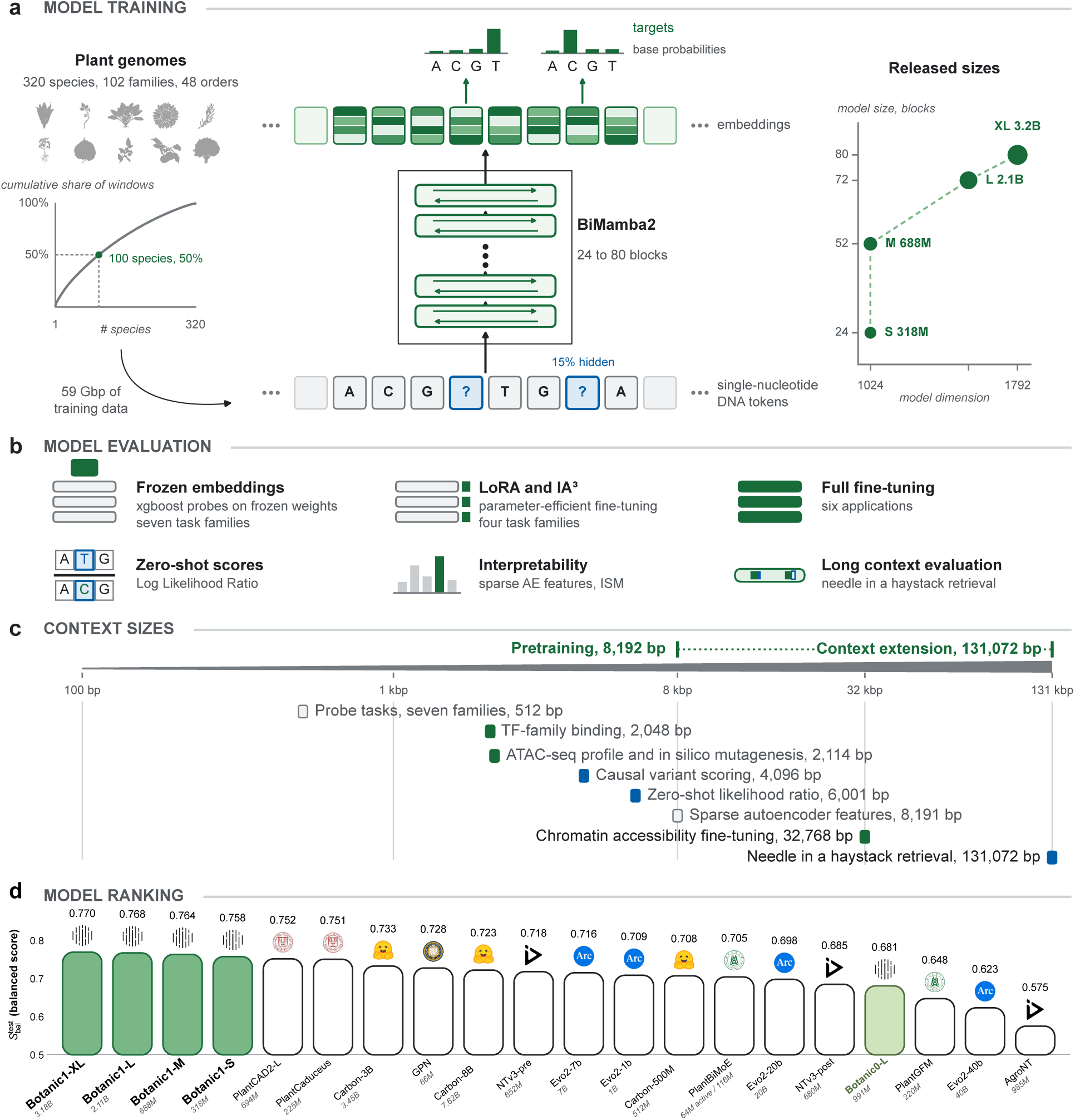
The Botanic1 model factory, from pre-training to the 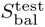 leaderboard. **a**, <u>Model training.</u> Corpus is plant genomes only. Training is masked language modelling on single nucleotide positions: 15% of eligible positions are hidden and the model is tasked to predict them. The backbone is a stack of bidirectional Mamba2 blocks, released at four sizes from 318M to 3.2B parameters. **b**, <u>Model evaluation.</u> We test our model under the six evaluation regimes detailed in the panel, from probes on frozen representations through zero-shot likelihood, adaptation and mechanistic interpretability to needle-in-a-haystack retrieval. **c**, <u>Context sizes.</u> Together they span context sizes from 100 bp probe tasks through the 8,192 bp base pre-training context out to the 131,072 bp reached by context extension and retrieval. **d**, <u>Model ranking.</u> 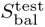 for 20 models, each bar topped by its organisation’s logo and labelled with its parameter count. The axis starts at 0.5, so bar length is not proportional to the score. Filled bars are ours, Botanic0-L in light green; all four Botanic1 sizes rank above PlantCAD2-L (0.752).

This evaluation-driven process allows us to assess heuristics from the literature quantitatively, distinguish robust design principles from spurious findings, and incorporate the validated improvements into successive generations of the Botanic family (Figure 1). We present the results of the latest Botanic generation below.

We further increase the speed of our experimentation thanks to LLM agents (Supplementary Figure S1): agents are responsible for infrastructure management and experiment throughput while research is driven by humans, who formulate hypotheses, design architectures, propose losses, and add or retire data sources and preprocessing steps. Each two-week-long research programme yields a small set of candidate models of different sizes, which become either major or minor versions of the Botanic1 series and can then be tested extensively on a larger evaluation suite and undergo fine-tuning, mechanistic interpretability and agentic discovery tests.

### 2.2 Training Botanic1

#### 2.2.1 Botanic1: careful data-mixture curation is the engine of the model factory

Our data pipeline acts at three levels, *inter-species*, *intra-species* and *inter-window* (Figure 2a). We systematically evaluate different genomic pre-training data design decisions (Section 4.2) one at a time, holding the parameter and training budget fixed, and measure their impact on downstream task performance using the scalar metrics defined in Section 4.3.

**Figure 2.**
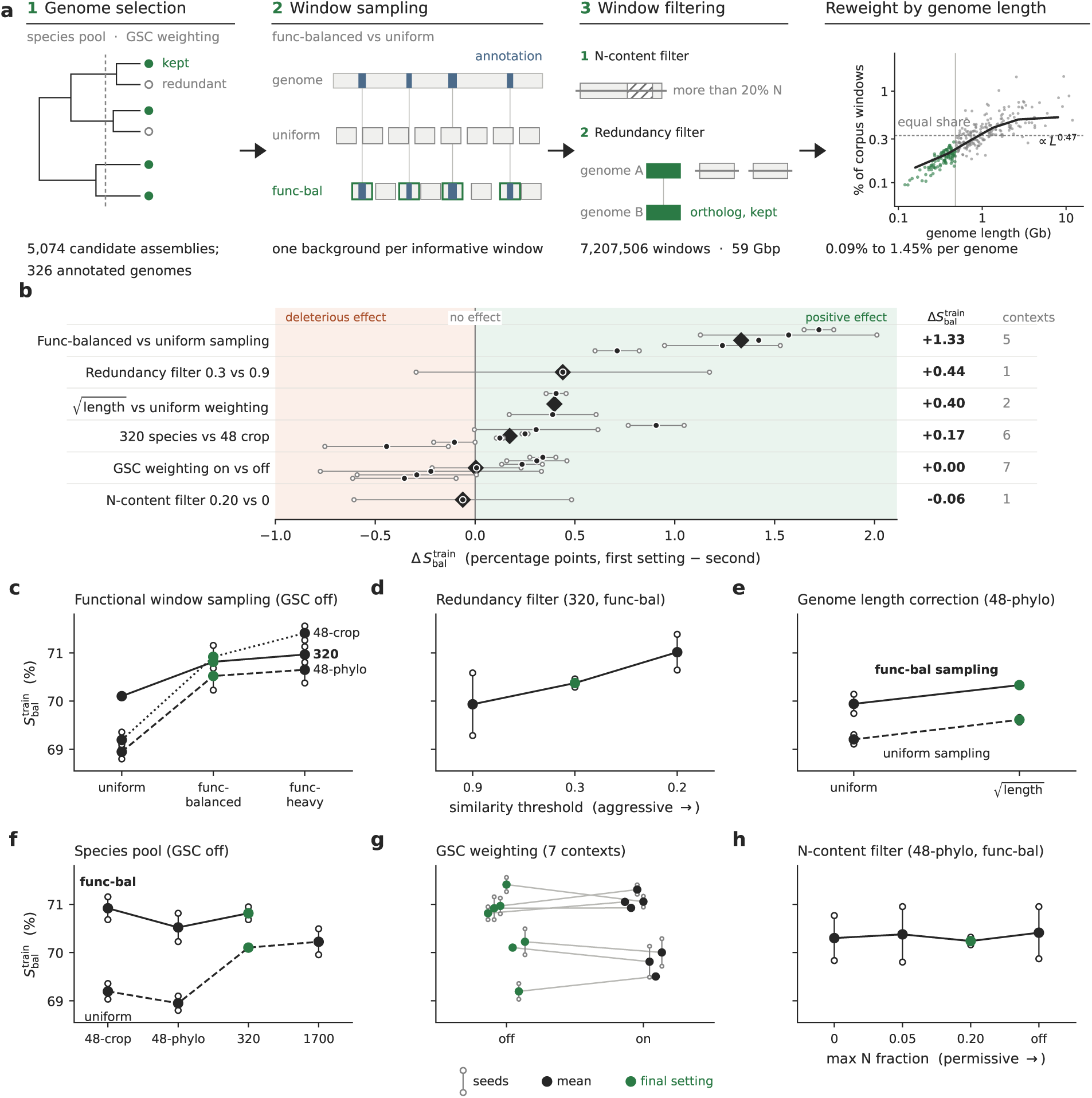
Pre-training-data preparation and ablations. **a**, (1) *Genome selection* (inter-species): input species catalogues such as Ensembl or NCBI are merged, filtered on assembly quality, and sourmash distances (*d* = 1 *−* ANI) clustered by complete linkage, keeping one representative genome per cluster. (2) *Window sampling* (intra-species): within a species, windows are sampled according to precise sets of rules relying on genomic region annotations or repeats. (3) *Window filtering* (inter-window): after sampling, some windows are filtered out due to redundancy or quality criteria (presence of N), or downsampled for large species in order to equalise the sampling across species irrespective of their sizes. The fourth cell, *Reweight by genome length*, visualises how far that equalisation goes: one point per genome (*n* = 306), green where the genome-length weight exceeds its floor, dashed line an equal split, solid line the binned median. Thresholds and parameters: Section 4.2. **b** to **h**, Each ablation experiment trains a 300M-parameter model at iso-step, iso-token and iso-effective epoch, downsampling the unique token pool across different settings if necessary. **b**, Each row reports the downstream performance difference of the first-named setting minus the second, the rest of the configuration being otherwise fixed, ordered by average impact across comparison contexts (sets of experiments where only the specified setting differed). Filled circles mark each context’s two-seed mean, the whiskers and open circles the spread of its two seeds, and the filled diamond the mean over contexts; GSC is measured on minus off, while the final mix uses GSC “off”. **c** to **h**, Absolute downstream performance when varying each setting individually. Panels follow the row order of **b** and share the same y-axis. Open circles are two seed replications, the filled marker their mean, and a green marker the chosen setting. Where a panel draws several series those are the different comparison contexts each with its own line style; the seven comparison contexts in **g** are unnamed. Species pools: the 48 crop-centric dataset of Botanic0 and AgroNT, a 48 phylogenetically diverse set, all 326 non-redundant annotated species, and 1700 species, for which only uniform sampling is available due to lack of annotations. **f** to **h** show inconclusive results.

##### Intra-species control

We find that the intra-species control (how windows are sampled from the whole genome) has the largest impact on performance (Figure 2b, with per-setting absolute scores in the remaining panels of Figure 2). Biasing window sampling to include more functional regions consistently outperforms uniform sampling of the genomes across all species pools (Figure 2c). Different ratios of functional to non-functional windows (1:1 for functional-balanced or 4:1 for functional-heavy, Section 4.2.7) do not separate cleanly in our experiments, so we choose to favour the highest scaling potential: functional-balanced can produce more unique tokens by drawing from a larger pool of non-functional windows, and includes more intergenic and repetitive regions which may contain important regulatory elements, supporting our choice for the 1:1 ratio. Similarly, whenever a data-ablation result is inconclusive, we usually favour the most scalable option that preserves the most unique tokens.

The second most impactful parameter is a new preprocessing step for gLM training: inter-window redundancy reduction within species using a MinHash [59] filter. We compare three settings based on the similarity threshold: starting from 8M sequences (65 Gbp), 0.9 removes barely any windows (0.8%), 0.3 removes around 20% and 0.2 is an extreme setting leaving only 4.3% of windows or 2.81 Gbp. All experiments train for the same number of optimiser steps on the same number of tokens (iso-step and iso-token), and their dataset is trimmed down where necessary so that the pool of unique tokens matches as well, which equalises the number of passes over that pool (iso-effective-epoch). The most aggressive threshold (0.2) scores higher on average but does not rank first across all seeds, and the gain does not warrant the steep cost in unique windows retained, so we adopt the intermediate 0.3 threshold (Figure 2d).

##### Inter-species control

For inter-species sampling, we find that correcting for genome length is also beneficial (Figure 2e). The weighting scheme and its effect on the composition of the corpus are detailed in Section 4.2.3.1.

##### Other controls, negative results

Neither N-content filtering (Section 4.2.7) nor manual curation of the species pool carries significant improvement over the other choices (Figure 2f and Figure 2h). Broadening the species pool from 48 crop genomes originally in [31, 32] to 326 Embryophytes has no detrimental effect on S^train^, built with various functional public data and mainly focused on *Arabidopsis thaliana* and crop species. Sampling species with phylogenetic-distances-derived Gerstein-Sonnhammer-Chothia (GSC) weights [60] instead of using a uniform sampling does not have a consistent effect on downstream tasks (Figure 2g).

##### Reporting the correct metrics is crucial when comparing different pre-training schemes

Some of the tested settings have a large effect on the proportion of repeated regions included in the pre-training dataset, functional window sampling being a prime example. We measure the correlation between the final training loss and the downstream performance across all data ablation experiments and find a strong positive correlation (r ≈ 0.75), indicating that downstream evaluation improves when loss worsens, and highlighting the importance of reporting the right metric when comparing training data preparation strategies, including different repeat-token fractions (see Supplementary Figure S2).

We also test to which extent data specialisation influences downstream task performance on any given species. We train models on a Brassicales dataset (the clade to which Arabidopsis belongs) and on a more phylogenetically diverse species pool, including or leaving out Arabidopsis in each case. We measure the performance of each of the four resulting models on the Arabidopsis downstream tasks and on all the other tasks on other species, and we find that training on Brassicales, even without Arabidopsis itself, improves performance on Arabidopsis tasks more than including the test species itself in the pre-training (Supplementary Figure S3, protocol in Section 4.2.7).

#### 2.2.2 Scaling Botanic1 improves pre-training and downstream performance

To characterise how the Botanic architecture scales, we pre-train our bidirectional-Mamba2 (BiMamba2) backbone on the data mixture from Section 2.2.1 at four capacities: Botanic1-S (318M), Botanic1-M (688M), Botanic1-L (2.1B) and Botanic1-XL (3.2B) (see Section 4.1). All four models share the same optimisation recipe and the same 314.6B-token horizon (200k steps for Botanic1-S, Botanic1-M and Botanic1-L and 100k steps for Botanic1-XL), with the learning rate fully decayed at the reported horizons (Section 4.1). All models are pre-trained on sequences of 8,192 tokens, a <cls> token followed by 8,191 nucleotides (Section 4.1).

The backbone choice is supported by a controlled comparison: a 352M encoder-only Transformer pre-trained with the same data, tokeniser, window length and learning-rate schedule needs about six times more tokens than Botanic1-S to reach its best 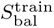. What is more, training the Transformer further, to 500B tokens, still leaves it well below the value Botanic1-S attains at 314.6B tokens, trailing in eight of the nine task families (Section 4.1, Supplementary Figure S9).

We find that, for each run, training loss decreases approximately linearly with the logarithm of the number of tokens seen: a *b* log_10_ D, with D the number of tokens. Fitting over the constant-learning-rate phase (3.9B to 283B tokens) gives *b* = 0.074, 0.084, 0.095 and 0.099 for Botanic1-S, Botanic1-M, Botanic1-L and Botanic1-XL respectively (R^2^ 0.99). Our 314.6B-token horizon also sits well above what a Chinchilla-like scaling law would recommend [61], whose compute-optimal allocation of roughly twenty tokens per parameter would stop at 6B tokens for Botanic1-S and 64B for Botanic1-XL. However, extending the token horizon at fixed capacity does not improve the downstream score much and does not generally close the model size performance gap (Supplementary Figure S4).

Comparing models of different sizes at matched token counts, we see that both training loss and 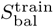 improve with scale (Section 4.3), the latter rising from 0.735 for Botanic1-S through 0.743 for Botanic1-M and 0.746 for Botanic1-L to 0.748 for Botanic1-XL. Under 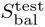 (Figure 3), the same four checkpoints occupy the top leaderboard positions from 0.758 to 0.770 with the same ordering.

**Figure 3.**
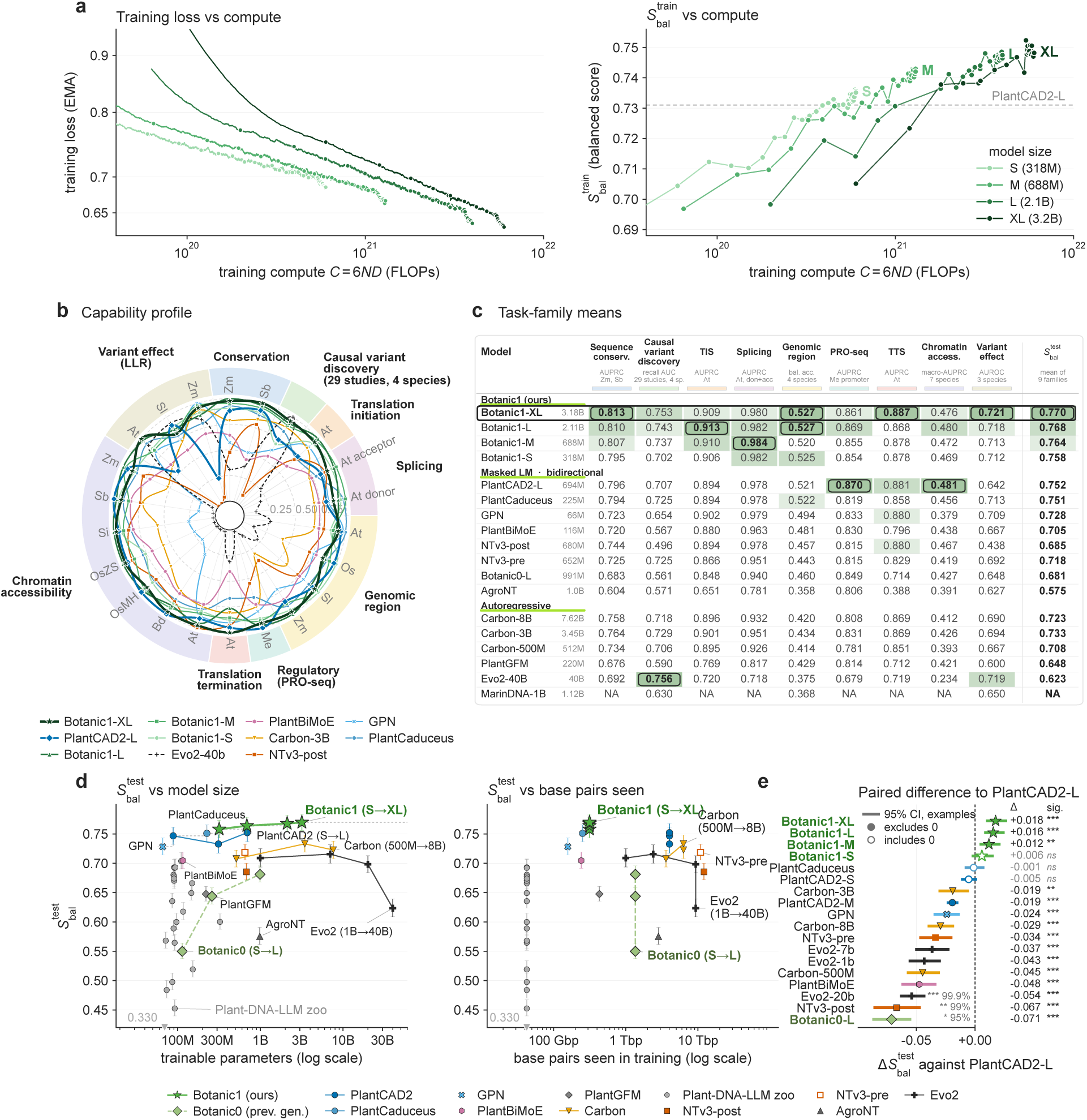
Botanic pre-training scaling, capability profile, and training efficiency. *S*_bal_ is the mean over nine task families of their per-family species means (22 metrics; higher is better), every model scored under the same test protocol, 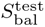, with variations in pooling type for auto-regressive models. **a**, Training loss and 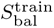 against training compute *C* = 6*ND*; markers are evaluation checkpoints, the dashed line is PlantCAD2-L, and both are scored with the training-specific faster evaluation protocol. **b**, One number per task and species, grouped into the nine task families; the radius is the model’s min-max normalised rank within the 43-model cohort, (*r −* 1)*/*(*n −* 1). Species codes are genus and species initials (At, *Arabidopsis thaliana*; OsMH and OsZS, the MH63 and ZS97 accessions of *Oryza sativa*). Causal variant discovery (subset) computes scores over a fixed subset of 29 curated studies from four species (13 *A. thaliana*, 11 *O. sativa*, 4 *Triticum aestivum*, 1 *Sorghum bicolor* ), each causal SNP ranked by signed LLR against 100 background SNPs of its locus (Section 4.10); the full 545-study benchmark results are reported in Section 2.5.1.2. **c**, The same breakdown in absolute value, each column shaded on its own scale; TIS and TTS are the translation initiation and termination sites. MarinDNA-1B is scored only on the families that fit its 255 bp context. **d**, 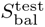 against model size and against base pairs seen, reconstructed from each publication at 7 nt per BPE token. Evo 2-20B is pruned from Evo 2-40B and the two Carbon models are iso budget. Vertical bars are 95% confidence intervals from the per-sample bootstrap (20,000 replicates, Section 4.3.1). **e**, Difference in 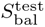 to PlantCAD2-L for every model within 0.10 of it, with its paired 95% interval from the per-sample bootstrap; a filled marker means the interval excludes zero. The right-hand column gives the difference and its significance level (*^∗^*95%, *^∗∗^*99%,*∗∗∗*99.9%). Models more than 0.10 below PlantCAD2-L and the Plant-DNA-LLM suite are omitted. Supplementary Figure S18 gives every pair of the 11 leading models, Supplementary Table S7 the family-level test.

#### 2.2.3 Context-extending Botanic1-S to 131 kbp windows

Regulatory interactions in plant genomes often span tens to hundreds of kilobases, far beyond the 2^13^ = 8,192 bp window seen by Botanic1 models during their pre-training. To make Botanic1 able to capture such interactions, we continue pre-training a 318M BiMamba2 backbone (equivalent to Botanic1-S but pre-trained for 500 billion tokens) for an extra 300B tokens on longer windows. To ease its way to long context, we proceed with a four-stage curriculum that gradually introduces longer contexts, doubling the longest window seen at each stage: 2^14^ = 16,384, 2^15^ = 32,768, 2^16^ = 65,536 and finally 2^17^ = 131,072 bp. The learning rate is warm-restarted at the start of each stage (Sections 4.1.5 and 4.2.8, Supplementary Figure S4). At each stage we keep some shorter windows in the training mix to avoid catastrophic forgetting: without it, a model would improve on the recently seen long windows but regress on short ones. The stability of *S*_bal_ on our short-context downstream tasks (scored on windows of 100 to 1,000 b, Section 4.3) across the stages of the curriculum is already reassuring: it remains near its 2^13^ bp value (≈ 0.73) up to a 2^17^ bp context (Supplementary Figure S4). Masked-language-modelling loss drops sharply at every context transition and decreases monotonically. We evaluate the capabilities that the extended context unlocks in Section 2.3.3. This series of context-extended models is used only for long-context tasks; elsewhere we use the regular Botanic1 family models.

### 2.3 Botanic1 is a state-of-the-art encoder of plant genomes

#### 2.3.1 Botanic1 has the best pareto performance profile across all competitors

##### 2.3.1.1 Botanic1 outperforms competing models on frozen model benchmarks

###### Best overall performance and efficient training

Figure 3 shows the ranking of a large panel of both generalist gLMs (gLMs trained on diverse organisms’ genomes) and plant specialist gLMs trained exclusively on plants (Section 4.3.6), with the per-family values reported in Figure 3c. All four Botanic models rank above the strongest competitor, PlantCAD2-L (aggregate score 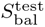 = 0.752; the two aggregate scores used in this figure are defined in Section 4.3), and their performance scales monotonically with model size, from Botanic1-S (0.758) through Botanic1-M (0.764), Botanic1-L (0.768) to Botanic1-XL (0.770), moving the frontier on plant genomic frozen-model benchmarks. Plotting the aggregated score 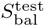 against model size (Figure 3d) highlights the efficiency of this frontier: the Botanic family leads at every scale, and even Botanic1-S (318M parameters) outperforms baselines with more than an order of magnitude larger, including Carbon-8B (7.6B, 0.723) and Evo 2 (7B, 0.716), both margins significant under both tests (95% CIs [+0.025, +0.046] example-level and [+0.015, +0.058] family-level against Carbon-8B, [+0.029, +0.056] and [+0.019, +0.068] against Evo 2); the most parameter-efficient baseline, GPN (66M, 0.728), still trails on performance.

The right panel of Figure 3d replots the same models against the total number of genomic base pairs seen during training: the tokens seen, reconstructed for every model from its publication or model card (Section 4.3.7). We convert tokens to base pairs using approximations for 6-mer models which tokenise the ambiguous nucleotide N as a single token. The Botanic scores are attained after only 314.6B base pairs, where the strongest competitor, PlantCAD2-L, had to process 4.03T base-pairs and NTv3 12.1T; among the models with their 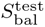 larger than 0.7, only PlantCaduceus (251.7B), PlantBiMoE (241.3B) and GPN (157.3B) saw fewer base pairs, and all three trail every Botanic in this benchmark. Interestingly, the Evo 2 family does not scale well with capacity: it peaks at 7B (0.716, from 0.709 at 1B) and declines through 20B (0.698) to 40B (0.623), so even at 40B it is not competitive on plant benchmarks, specifically on probing tasks, indicating that scale alone is not sufficient to achieve strong performance on plant-specific benchmarks.

As for plant specialists, the 22-model Plant-DNA-LLM suite [62] (70–325M parameters) covers a wide range of scores from 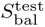 = 0.330–0.704. Its strongest score is obtained with ModernBERT-single-base but still trails every Botanic model as well as the leading plant-specific competitors. PlantBiMoE [30], a sparse mixture-of-experts with BiMamba masked encoders trained on 42 plant genomes (116M trainable parameters, 64M active per token), reaches 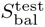 = 0.705, on par with that strongest Plant-DNA-LLM checkpoint and below GPN, a smaller backbone trained on even fewer species. Splitting the size axis by pre-training domain (Supplementary Figure S17) shows that training on plants alone does not necessarily lead to a better performance on plant-specific benchmarks: the majority of plant-only models score below the best generalist. Botanic1-S beats MarinDNA-1B on all 8 tasks compatible with MarinDNA’s 255 bp context, with a mean of 0.617 against 0.507, despite being about 3.5 times smaller. Note that while the gap between genomic region classification performance is large (0.525 vs 0.368), the above-average zero-shot log-likelihood ratio (LLR) performance of MarinDNA shows that mammalian pre-training transfers to plant variant ranking to some extent. However, specialising to plants is more efficient in terms of model size: GPN, trained on 8 Brassicales species, reaches an 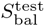 of 0.728 with 66M parameters, matching the best model trained beyond plants, Carbon-3B (0.733; difference −0.005, 95% CI [ 0.020, +0.009] under per-sample bootstrap), at a fiftieth of its size, and well ahead of Evo 2 at six hundred times its size (0.623).

###### Plant gLM benchmarks measure different aspects of sequence representations

Zooming in per-task (Supplementary Figure S5) shows that Botanic1 leads overall without dominating every capability: the flagship Botanic1-XL tops the benchmark and three task families, LLR (0.721), conservation and translation termination, Botanic1-L leads two more (genomic-region classification, 0.527 versus 0.521 for PlantCAD2-L, and translation initiation) and Botanic1-M leads splicing (donor and acceptor mean 0.984 versus 0.978 for NTv3-650M-post, which uses test annotations during post-training; see Section 3). The autoregressive Evo 2 [24] is close behind on variant-effect prediction, where its 7B model reaches an LLR area under the receiver operating characteristic curve (AUROC) of 0.710 and its 40B model 0.719 against 0.721 for Botanic1-XL, and its 40B model leads causal-variant discovery (0.756 against 0.753); PlantCAD2-L [33] leads on chromatin accessibility (0.481 versus 0.480 for Botanic1-L) and PRO-seq (0.870 versus 0.869). Botanic1-XL is the only model that stays competitive across all nine families. The two NTv3 checkpoints also present widely different performance profiles: the pretrain-only NTv3-pre reaches 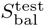 = 0.718 against 0.685 for the released post checkpoint (a gap of 0.033, 95% CI [+0.015, +0.050] over examples), as post-training degrades evolutionary-based tasks (Section 3). In order to gauge the validity of our choice of aggregation method (averaged per-task type and across species, see Section 4.3), we resample the nine families of metrics (paired bootstrap, 20,000 replicates, see Section 4.3.1) and show that the rankings hold for most pairs of models: the Botanic1-L and Botanic1-XL 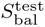 margins over PlantCAD2-L are significant: +0.016 (95% confidence interval (CI) [+0.002, +0.034]) and +0.018 ([+0.003, +0.036]), leading on respectively six and seven of the nine families. Botanic1-M stands at +0.012 ([ −0.002, +0.030]) with five families ahead of PlantCAD2-L, but the difference is not statistically significant. The Botanic1-S margin over PlantCAD2-L is not statistically significant either: +0.006 with a 95% interval of [ −0.006, +0.024]. Regarding statistical significance, when assuming the choice of aggregation, we do bootstrap draws at the sample-level within each of the 22 tasks and species, with the same draws for both models (Section 4.3.1). The Botanic1-L and Botanic1-XL margins over PlantCAD2-L are significant under both tests: +0.016 ([+0.002, +0.034] over families, [+0.008, +0.025] over examples) and +0.018 ([+0.003, +0.036] and [+0.009, +0.027]), leading on respectively six and seven of the nine families. Botanic1-M stands at +0.012 with five families ahead: its familylevel interval [ 0.002, +0.030] includes zero and its example-level interval [+0.004, +0.022] is significant. The Botanic1-S margin, +0.006, is not significant under either test ([ −0.006, +0.024] and [ −0.003, +0.016]). The family-level intervals are longer than the example-level ones for these pairs, and the two tests agree on most pairs of Supplementary Table S7 except Botanic1-M against PlantCAD2-L. Figure 3e draws the example-level test against PlantCAD2-L for every competitor model, and Supplementary Figure S18 extends it to all pairs of the leading models: it separates significantly every Botanic size from PlantCaduceus, PlantCAD2-S and PlantCAD2-M, at the 99.9% level for Botanic1-M, Botanic1-L and Botanic1-XL and at the 95% level for Botanic1-S against PlantCaduceus (+0.007) and PlantCAD2-S (+0.011), whereas PlantCAD2-L and PlantCaduceus do not separate significantly (+0.001, ns) and neither do PlantCAD2-M, GPN and Carbon-3B from one another. The ranking is also robust to the aggregation rule: replacing 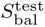 with the unweighted mean of the 22 metrics or with a subset of the 22 tasks or with the first principal component of the score matrix does not change the fact that Botanic1 models occupy the top of the leaderboard, with Spearman ρ ≥ 0.89 against 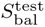 over 24 models (Section 4.3, Supplementary Figure S43).

In spite of the lack of statistical significance using this test, Botanic1-S wins nine of the 22 metrics, and PlantCAD2-L’s edge is mainly in one task: chromatin accessibility, where it leads all seven species by a tiny margin 0.001 to 0.029. Because *S*_bal_ gives that family a ninth of the score irrespective of its number of species (7) (Equation (1)), the consistent advantage of PlantCAD2-L there offsets Botanic1-S’s wins elsewhere, led by LLR in maize (+0.163, Supplementary Figure S12), LLR in *A. thaliana* (+0.055), genomic region in maize (+0.026) and translation initiation (+0.011).

Outside of tasks included in *S*_bal_, Botanic1-S outperforms PlantCAD2-L. On adaptation, Botanic1-S beats PlantCAD2-L on all four tasks under each of the three fine-tuning regimes of Table 1, scored on Pearson *r* for the two regression tasks and mean AUROC for the two classification tasks, with margins up to 0.32 AUROC on long non-coding RNA (lncRNA) classification. On out-of-species splice transfer, Botanic1-S leads 0.826 to 0.818 on the six-cell mean and the larger sizes lead by far more (Section 2.3.1.2). This reflects the inherent instability of PlantCAD2-L fine-tuning.

**Table 1:** Fine-tuning across four plant-regulatory tasks. Held-out test metric per model fine-tuning adaptation regime; best in each row across all four models in **bold**. The two regression tasks report Pearson *r* <u>and</u> R^2^. Classification tasks report mean AUROC over six species. The CNN column is the best public taskspecific model trained from scratch, and is placed under “full fine-tuning”; its metrics are recomputed from the authors’ released test-set predictions (Section 4.3.8). Classification cells report global per-species: <u>global</u> tunes hyperparameters once across all species, while <u>per species</u> selects the best hyperparameter set for each species on held-out validation (top-two runoff, Section 4.3.8); *^↑^* marks a per-species lift of 0.05. For Terminator strength we train a single head for both species, so it has no per-species run; Promoter per-species tuning moves the two-tissue mean Pearson *r* by at most 0.009 in either direction and is not shown. Notes: *^a^* full fine-tuning recovers under an fp32 classification head at a low learning rate (1–3 10*^−^*^6^): five of six species reach 0.91–0.97 and only *Chlamydomonas reinhardtii* remains near random (0.50), with a mean of 0.870 (in bf16 all six metrics collapse to 0.50); *^b^* all six species near random (mean AUROC 0.53); *^c^ Arabidopsis thaliana* collapses to constant output and is excluded; mean over the remaining five species; *^d^ Manihot esculenta* collapses to constant output and is excluded; mean over the remaining five species; *^e^* one species (maize) is non-finite and excluded, so the mean is over the remaining five; *^f^* terminator from-scratch model: Gorjifard DenseNet; *^g^* promoter from-scratch model: Jores CNN; *^h^* values published by the AgroNT authors [31] for their own IA^3^ fine-tuning (their Figures 2b, 2f, 3c and 3d), listed against the IA^3^ row and excluded from the bold comparison.

| Method | Botanic1-S | AgroNT | AgroNT<br>(published)<sup>h</sup> | PlantCAD2-L | CNN<br>(from scratch) |
| --- | --- | --- | --- | --- | --- |
| *Terminator strength, merged (tobacco / maize)* |  |  |  |  |  |
| Pearson $r$ | | | | | |
| Full FT | **0.861** (0.882 / 0.839) | 0.834 (0.865 / 0.803) | — | 0.790 (0.820 / 0.760) | 0.85 (0.87 / 0.82)<sup>f</sup> |
| LoRA | **0.856** (0.889 / 0.822) | 0.809 (0.848 / 0.769) | — | 0.780 (0.828 / 0.731) | — |
| IA<sup>3</sup> | **0.839** (0.883 / 0.794) | 0.808 (0.846 / 0.770) | — | 0.812 (0.845 / 0.779) | — |
| $R^2$ | | | | | |
| Full FT | **0.736** (0.773 / 0.698) | 0.693 (0.742 / 0.643) | — | 0.624 (0.671 / 0.576) | 0.72 (0.76 / 0.67)<sup>f</sup> |
| LoRA | **0.717** (0.783 / 0.650) | 0.626 (0.710 / 0.542) | — | 0.590 (0.679 / 0.501) | — |
| IA<sup>3</sup> | 0.588 (0.702 / 0.474) | 0.583 (0.585 / 0.580) | — (0.77 / 0.67) | **0.624** (0.648 / 0.600) | — |
| *Promoter strength (tobacco / maize)* |  |  |  |  |  |
| Pearson $r$ | | | | | |
| Full FT | **0.896 / 0.874** | 0.864 / 0.831 | — | 0.853 / 0.831 | 0.85 / 0.82<sup>g</sup> |
| LoRA | **0.900 / 0.872** | 0.844 / 0.825 | — | 0.864 / 0.839 | — |
| IA<sup>3</sup> | **0.882 / 0.862** | 0.843 / 0.816 | — | 0.837 / 0.833 | — |
| $R^2$ | | | | | |
| Full FT | **0.800 / 0.760** | 0.744 / 0.648 | — | 0.717 / 0.684 | 0.71 / 0.67<sup>g</sup> |
| LoRA | **0.783** / 0.689 | 0.654 / 0.668 | — | 0.740 / **0.691** | — |
| IA<sup>3</sup> | 0.663 / **0.720** | **0.680** / 0.573 | 0.73 / 0.70 | 0.651 / 0.687 | — |
| *poly(A) site: mean AUROC (6 species), global → per-species* |  |  |  |  |  |
| Full FT | **0.956** → 0.962 | 0.941 → 0.941 | — | 0.870<sup>a</sup> → 0.943<sup>†</sup> | — |
| LoRA | **0.956** → 0.961 | 0.926 → 0.946 | — | 0.878 → 0.952<sup>†</sup> | — |
| IA<sup>3</sup> | **0.958** → 0.959 | 0.856<sup>c</sup> → 0.935<sup>†</sup> | 0.937 | 0.914 → 0.878 | — |
| *lncRNA: mean AUROC (6 species), global → per-species* |  |  |  |  |  |
| Full FT | **0.848** → 0.842 | 0.819 → 0.819 | — | 0.529<sup>b</sup> → 0.704<sup>†</sup> | — |
| LoRA | 0.685 → 0.783<sup>†</sup> | **0.809** → 0.800 | — | 0.572<sup>e</sup> → 0.769<sup>†</sup> | — |
| IA<sup>3</sup> | **0.828** → 0.827 | 0.785<sup>d</sup> → 0.783 | 0.832 | 0.667 → 0.795<sup>†</sup> | — |

Additionally, Supplementary Figures S5 and S6 decompose the size-vs-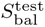 frontier of Figure 3d to the level of individual capabilities and metrics. Each panel plots performance against the trainable-parameter count (log axis) for every model family: one panel per capability sub-category (nine) in Supplementary Figure S5, and one panel per individual metric (34) in Supplementary Figure S6 including 12 that are excluded from *S*_bal_ (Section 4.3.4). We present another view of the same result in Supplementary Figure S17 which splits the size axis of Figure 3d by pre-training domain. Plant-specific pre-training is not a guarantee of a better performance on plant-specific benchmarks. The median 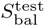 of the plant-only group (0.670) sits below the generalist models’ median (0.709), pulled down by the Plant-DNA-LLM zoo, and 30 of the 37 plant-only models do not outperform the best generalist, Carbon-3B (0.733). GPN scores 0.728 with 66M parameters, matching Carbon-3B at fifty times its size and well ahead of Evo 2 at six hundred times its size (0.623). Seven plant-specific models beat the best generalist, while Evo 2 loses 0.085 going from 1B to 40B.

##### 2.3.1.2 Botanic1 models lead out-of-species splice-site recognition

The *S*_bal_ splicing tasks are restricted to a single species: donor and acceptor recognition in Arabidopsis. These scores measure whether a representation captures local splicing grammar, but not whether the learned grammar transfers to other species. We therefore report a cross-species splicing benchmark built from the PlantCaduceus datasets suite [34]: models are trained on Arabidopsis donor and acceptor sites and evaluated on Arabidopsis, rice, sorghum and maize. Because class imbalance is strong in these datasets (near 10*^−^*^4^), the only informative metric that can be computed on a stratified sample is a prevalence-corrected area under the precision-recall curve (AUPRC). This estimator is itself biased and has high variability (Section 4.3.5); we therefore exclude the corresponding metrics from *S*_bal_, whose 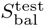 variant is used as the final ranking metric for all models. We perform eight measurements per model under the frozen-embedding protocol of Section 4.3.2. Every model is scored using the centre-pooling protocol of Section 4.3.6. Supplementary Figure S8 reports every transfer experiments for all models tested: Figure S8a shows the mean of all transfer metrics for the four Botanic sizes, the three PlantCAD2 sizes and the strongest member of each remaining family, and Figure S8b shows Botanic1-L against PlantCAD2-L at two sampling depths showing the impact of sampling more data points on the ranking between PlantCAD2-L and Botanic1-L: except for rice Botanic1-L keeps leading each task it was leading, by a wider margin, and leads most tasks where it did not lead before; Supplementary Figure S6 zooms in on each task giving performance as a function of model sizes including cross-species metrics. We notice that:

###### In-distribution scores are saturated and uninformative

Most models land between 0.95 and 0.98 AUPRC on Arabidopsis donor and acceptor; the best single Arabidopsis donor score is GPN (0.981), a 66M-parameter model trained only on Brassicales. Botanic1-XL, Botanic1-L and Botanic1-M take the top three positions (0.900, 0.878 and 0.873), ahead of every PlantCaduceus (0.830–0.855) and PlantCAD2 models at all sizes (0.818– 0.840) but the best models saturate the task. Therefore the margin between models is concentrated on hardest tasks: on maize donor Botanic1-XL reaches 0.958 and Botanic1-L 0.951 against 0.830 for PlantCAD2-L, and on sorghum acceptor Botanic1-L performance is 0.931 wrt Botanic1-XL at 0.915 and 0.910 for PlantCAD2-L. Under the per-example paired test (Section 4.3.1) Botanic1-L is significantly ahead of PlantCAD2-L on donor in maize (+0.121, 95% CI [+0.001, +0.221]); the other five splice cells do not separate the pair, acceptor in maize least of all (+0.266, [ −0.105, +0.511]). On each splice cell between 10 and 40 of the other 45 models sit significantly below PlantCAD2-L. PlantCAD2-L is the strongest model on both tasks on rice (0.930 donor, 0.916 acceptor) yet finishes thirteenth overall, because it drops to 0.695 on sorghum donor and 0.629 on maize acceptor. The performance of Botanic0, our previous generation of models, rises with capacity on this benchmark (0.142 0.605 0.808 for Botanic0-S, Botanic0-M and Botanic0-L), yet even Botanic0-L at 991M is outperformed by Botanic1-S (0.826) with only a third of its parameters.

###### Training objective dominates parameter count

Every model above 0.80 is an encoder pre-trained with a masked objective on plant genomes, and the strongest autoregressive model, the Plant-DNA-LLM “DNAMamba” with 4-mer tokenisation (91M, plant-trained), reaches 0.781. The four Evo 2 from 1B to 40B parameters reach only between 0.417 and 0.636, the three Carbon models from 0.5B to 7.6B between 0.497 and 0.647, while PlantCaduceus-l20, a 21M-parameter masked encoder, reaches as high as 0.842. The causal models are scored under the two-strand protocol of Section 4.3.6, which should give them an advantage over encoder-only models, but this advantage is not sufficient to close the gap. On the in-domain split, Arabidopsis, Carbon-3B (0.966) matches Botanic1-XL (0.966) and PlantCaduceus-l20 (0.968) on this saturated task, but on held-out species it loses 0.32 of AUPRC where the two encoders lose 0.07 and 0.13. Masked objective pre-training on plants by itself is not sufficient: the Plant-DNA-LLM encoders with BPE tokenisers reach only between 0.08 and 0.56.

###### Translation initiation and termination sites transfer the same way

The suite scores the same three held-out species on translation initiation sites (TIS) and translation termination sites (TTS) with the same estimator (Figure S8a), and the results are similar: Botanic and PlantCAD2 separate well from the rest of the models across all six tasks, models trained beyond plant genomes stay well below the plant-trained encoders, and performance is largely flat in parameter count above 10^8^. For these six tasks and species the per-sample paired bootstrap (Section 4.3.1) shows that between 30 and 45 of the other 45 models sit significantly below PlantCAD2-L depending on the task, whereas the leading models are rarely significantly different from one another. Botanic1-L is significantly ahead of PlantCAD2-L on TIS in rice (+0.178, 95% CI [+0.021, +0.333]); PlantCAD2-L is significantly ahead on TTS in sorghum ( 0.134, [ 0.195, 0.033]). No significant difference between the two models is measured on the other four tasks and species. The lower score of Botanic1-XL on rice TTS (0.717 against 0.829 for Botanic1-L) is not a measurable size effect: the 95% interval of that single score has a half-width of 0.19 and covers the scores of the three smaller sizes.

These cross-species scores are estimated from stratified subsamples because the full test sets are too large to evaluate exhaustively (Section 4.3.5). As a result, they are substantially noisier than the scores used in *S*_bal_. For the Botanic1 models and PlantCAD2-L, the median 95% bootstrap half-width is about 0.065 for the held-out cross-species tasks, compared with 0.034 for the *S*_bal_ tasks. Thus, a typical held-out score is about twice as uncertain as a typical *S*_bal_ score. In the noisiest held-out task, the half-width reaches 0.28, about eight times the *S*_bal_ median. This uncertainty also depends on which examples are sampled: drawing a fresh stratified sample of the same size changes the Botanic1-L minus PlantCAD2-L difference by as much as 0.092 in a single task. Therefore this benchmark provides only a rough ordering of the models. Increasing the number of sampled rows strengthens the lead of the Botanic1 models (Figure S8b).

#### 2.3.2 Botanic1-S adapts to hard plant-biology tasks by fine-tuning

We fine-tune the smallest of our models Botanic1-S in two settings: parameter-efficient adaptation to four plant-regulatory tasks, benchmarked against two other gLMs, and base-resolution assay for transposase-accessible chromatin using sequencing (ATAC-seq) profile prediction, benchmarked against a specialised task model. Botanic1-S leads both tasks.

##### 2.3.2.1 Parameter-efficient adaptation of Botanic1-S to four regulatory tasks

We first fine-tune Botanic1-S on four tasks including both regression and classification: terminator and promoter strength (STARR-seq transcriptional activity assayed in tobacco leaf and maize protoplast [63, 64]), poly(A)-site identification, and lncRNA classification, all from the PGB. Each task is learned under three adaptation regimes, from the most costly to the fastest (full fine-tuning, LoRA and IA^3^), and benchmarked against two competitors: AgroNT [31] (1B parameters) and PlantCAD2-L [33]. Hyperparameters are selected by grid search on a held-out validation split, independently for each task backbone fine-tuning regime. Since neither Botanic1-S nor PlantCAD2-L has attention layers, the LoRA and IA^3^ adapters are used on the two linear projections of every Mamba2 module (Section 4.3.8). Using the selected hyperparameters, the model is retrained once on the full training set and subsequently evaluated on the test set. For the hardest datasets, we also tune models per species and report the differences. In regression, Botanic1-S outperforms both competitors. For terminator-strength prediction, its Pearson *r* is 0.861, compared with 0.834 for AgroNT and 0.790 for PlantCAD2-L. For the two promoter-strength systems, it reaches 0.896 and 0.874, compared with 0.864 and 0.831 for AgroNT. Pearson *r* measures association and, as reported in [32], is sufficiently stable to guide training as an early-stopping criterion. By contrast, R^2^ measures prediction magnitude but is noisy during training and can take large negative values. Under this pointwise metric, Botanic1-S is the only model to outperform the previously unbeaten task-specific baselines trained from scratch. We report calibration of all models and fine-tuning regimes in Supplementary Figure S24, and the predicted-versus-observed densities in Supplementary Figure S23. For the poly(A) and lncRNA classification tasks, Botanic1-S also leads in every setting except LoRA on lncRNA, which performs poorly even when tuned globally across species; the per-species breakdown of both tasks is given in Supplementary Figure S25 and Supplementary Figure S26. Fine-tuning PlantCAD2-L with the authors’ code is too unstable to achieve strong performance on poly(A) and lncRNA, and requires several model modifications to resolve numerical instabilities (Section 4.3.8). Neither Botanic1-S nor AgroNT requires these modifications.

The AgroNT authors claim to have fine-tuned their model with IA^3^ only, and report R^2^ for the regression tasks and per-species AUROC for the classification tasks [31]. We add the published values next to our reproduction in Table 1. While our full fine-tuning of AgroNT reproduces the published poly(A) and promoter results to some extents: mean AUROC 0.941 against 0.937 published, every species within 0.01, and promoter R^2^ of 0.744 / 0.648 against 0.73 / 0.70 (tobacco / maize), we do not manage to reproduce their scores on terminator strength, and on lncRNA both by a wide margin. The authors do not publish their IA^3^ hyperparameters making reproduction harder.

##### 2.3.2.2 Pre-training drives Botanic1’s base-resolution ATAC-seq advantage over ChromBPNet

In addition to the binary peak-classification chromatin task from the short-context evaluation suite (Section 4.3.2), we assess whether Botanic1 models can predict the ATAC-seq accessibility profile at single-base resolution. We fine-tune the pre-trained Botanic1-S backbone to regress base-resolution ATAC-seq signal for Arabidopsis and maize, and benchmark it against ChromBPNet [37], a strong base-resolution model for DNase-seq and ATAC-seq profile prediction. The signal is derived from BigWig coverage tracks processed through our own plant-specific pipeline (Section 4.5.1). We evaluate two prediction designs built on the same backbone: a <u>dual-head</u> design, in which separate heads predict the accessibility profile and total read count, and a <u>single-head</u> design, in which one head predicts the per-base signal and the total count is obtained by summation. The single-head design therefore does not require the loss-balancing weight used in the dual-head design. All models are scored on held-out chromosomes over the called ATAC peak set. Botanic1 and ChromBPNet use the same genomes, ATAC-seq tracks, peak sets, and chromosome splits, and both models’ predictions are scored by the same metric implementation, which reproduces ChromBPNet’s published values.

###### Botanic1 outperforms ChromBPNet thanks to pre-training

Botanic1 outperforms ChromBPNet on both accessibility magnitude (count Pearson and Spearman) and base-resolution profile shape (Jensen-Shannon distance, JSD) in both species (Table 2). With the pre-trained backbone, peak count correlation increases from 0.666 to 0.750–0.765 in Arabidopsis and from 0.809 to 0.847–0.857 in maize (dual and single head, Table 2), with consistent gains in rank correlation and profile shape (Figure 4b). a paired bootstrap over the shared test peaks confirms that the Botanic1-S advantage is statistically significant. Specifically, we repeatedly resample the test peaks shared by the two models and compute the difference in their performance, allowing us to estimate a 95% CI for this difference. For every metric in both species, the entire CI lies above zero. This remains true even for the profile-shape metrics, where the differences between the models are smaller.

**Figure 4.**
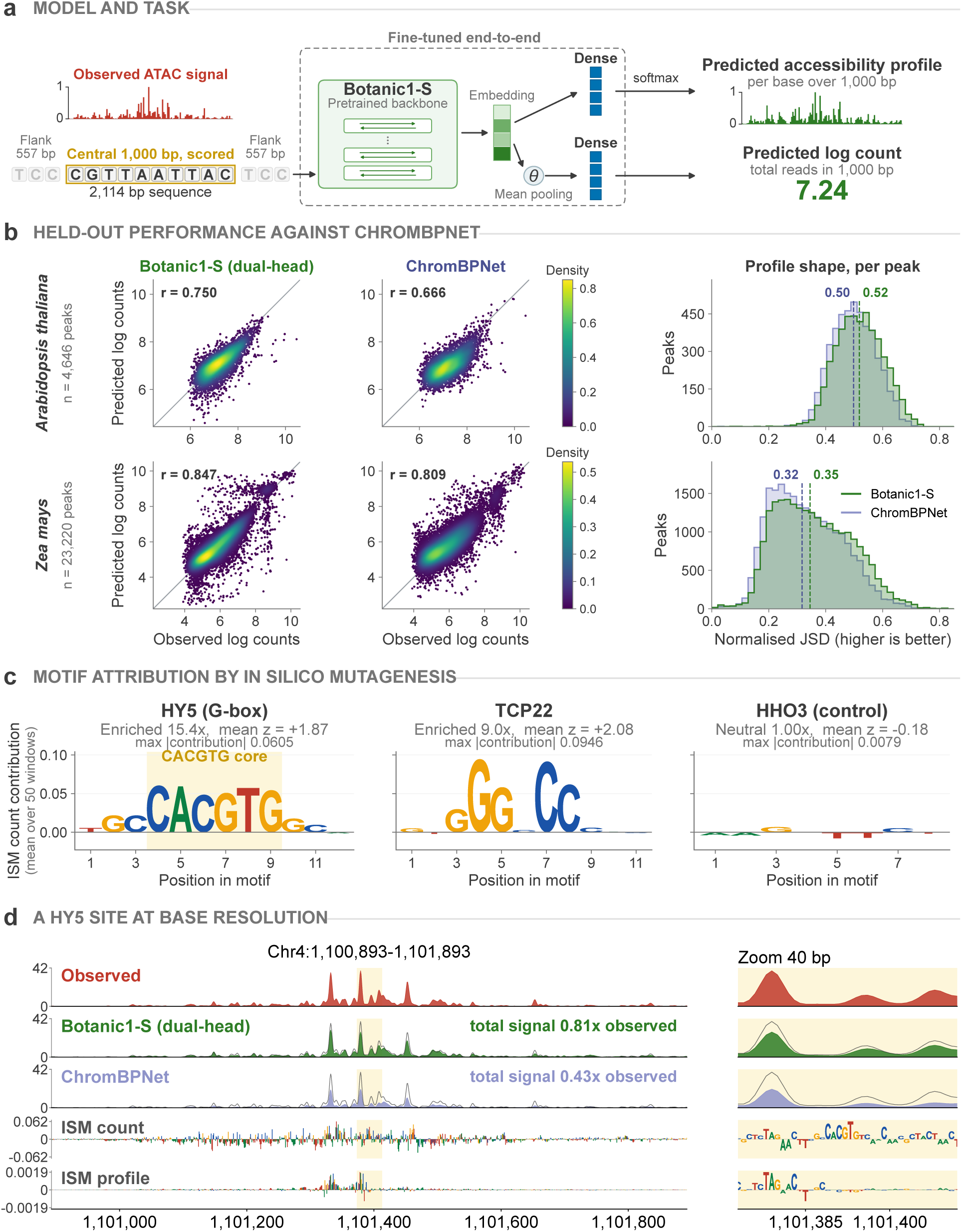
Base-resolution ATAC-seq prediction with Botanic1-S. **a**, a 2,114 bp window is read by the pre-trained Botanic1-S backbone; its last-layer embedding feeds a profile head (softmax, base-resolution accessibility over the central 1,000 bp) and a count head (mean-pool, log total count), fine-tuned end to end. Only the central 1 kbp is scored; the 557 bp flanks provide context. **b**, Botanic1-S versus ChromBPNet on held-out peaks (*Arabidopsis thaliana*, top, *n* = 4,646; *Zea mays*, bottom, *n* = 23,220): predicted versus observed per-peak log counts (density-coloured) for dual-head Botanic1-S and ChromBPNet with peak Pearson *r* (left, centre), and per-peak normalised JSD (higher is better) for both models with dashed lines at the medians (right). **c**, Per-base in silico mutagenesis (ISM) count contribution in three Arabidopsis motifs, averaged over 50 peak windows per motif: two functional motifs (HY5, a G-box; TCP22) and a control (HHO3), on one shared *y*-axis; each motif is a sequence logo (letter height is the contribution) and HY5’s CACGTG core is shaded. **d**, ISM at a representative HY5 motif’s site (Arabidopsis chromosome 4): lanes from top to bottom: observed ATAC signal, Botanic1-S (dual-head) predictions, ChromBPNet predictions, per-base ISM count contribution and per-base ISM profile contribution; the right panel zooms to 40 bp as a sequence logo, where the G-box (GCCACGTG) is the tallest feature. The three signal lanes share one *y*-axis, the observed profile is repeated as a dark outline on both model lanes, and each model lane states its total predicted signal relative to observed: Botanic1-S reaches 0.81 of the observed total at this locus, ChromBPNet 0.43.

**Table 2:**
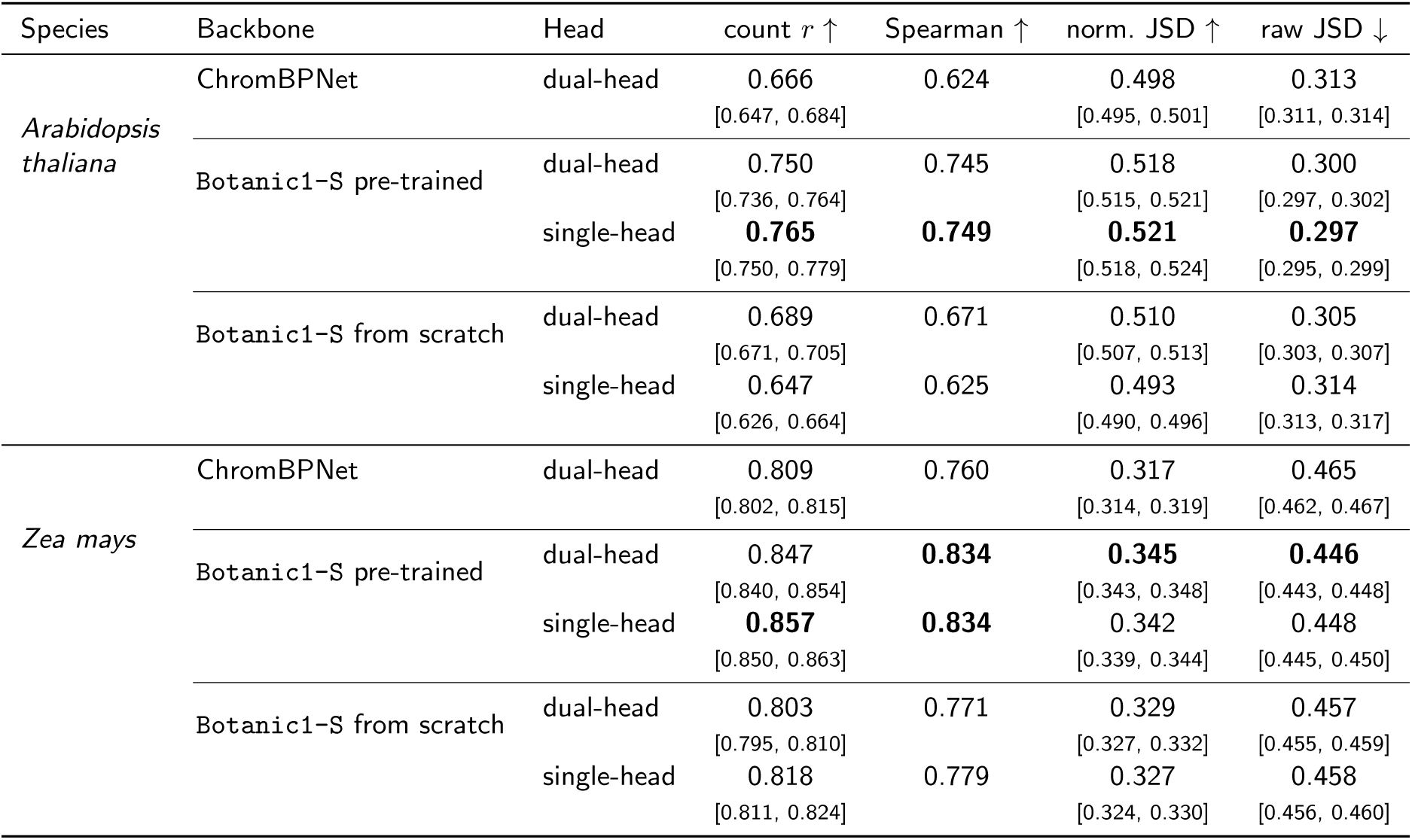
Base-resolution ATAC-seq prediction versus ChromBPNet. Held-out test metrics over the called ATAC peak set for the pre-trained and from-scratch Botanic1-S backbone under both head designs, against the specialised ChromBPNet baseline [37]. Test sets are held-out chromosomes: Arabidopsis chromosome 4 (4,646 peaks) and maize chromosomes 1 and 9 (23,220 peaks). Count *r* and Spearman measure the correlation between predicted and observed read totals across peaks. Profile shape is evaluated using normalised JSD (higher is better) and raw JSD (lower is better). Best per column within each species in **bold**. Values in brackets are 95% bootstrap confidence intervals (1,000 resamples over the test peaks). ChromBPNet is scored by its own pipeline, which does not provide confidence intervals.

To test whether this gain comes from pre-training rather than from the architectural differences between Botanic1-S and ChromBPNet, we fine-tune the same backbone from random weights. We see (Table 2) that architecture alone does not explain the strong performance of Botanic1-S. The advantage is thus attributable to the pre-training.

###### Motif attribution by in silico mutagenesis

We perform base-level attribution on the fine-tuned Botanic1-S to test whether its ATAC-seq predictions depend on real regulatory sequence features rather than on broader properties such as GC content or local nucleotide composition. For each sequence, we mutate each base individually, re-run the model, and measure the change in its prediction (Figures 4c and 4d). For each of three known Arabidopsis transcription factor (TF) motifs, we analyse 50 peak windows containing exactly one instance of that motif and compare the contribution scores at motif positions with those of the flanking bases within the same window, using paired comparisons across windows. We see a clear effect for two motifs from unrelated transcription-factor families. HY5, a bZIP G-box, has a mean *z* of +1.87, with the effect observed in 96% of windows, while TCP22, a TCP element, has a mean *z* of +2.08, with the effect observed in 78% of windows. In contrast, HHO3 occurs at a similar rate in peaks and background and is used as a negative control; it shows no such effect (z = 0.18, 20%; Supplementary Table S18). For the HY5 motif the model places roughly 16 more contribution on the CACGTG core than on the motif’s two outermost positions, and TCP22 concentrates on its GGG and CC blocks roughly 22× above its own outermost positions; both ratios are read off the contribution profile averaged over the 50 windows (Section 4.5.6), with the HY5 case shown at a representative locus in Figure 4d. This indicates that the model’s predictions depend on biologically meaningful motif sequence rather than only coarse local sequence composition (Figure 4c).

##### 2.3.2.3 Long-context Botanic1-S predicts genome-wide TF-family binding

We test whether our pre-trained models can be used to predict genome-wide transcription factor binding. Following O’Malley et al. [65], we group 568 *Arabidopsis thaliana* DAP-seq and ampDAP-seq assays into 46 transcription factor families and construct a multi-label task on non-overlapping 250 bp windows. Chromosomes 1 to 3 are used for training, chromosome 4 is used for hyperparameter and checkpoint selection, and chromosome 5 for testing.

We use our context-extended Botanic1-S model, taken at the end of the final 131 kbp stage of the curriculum of Section 2.3.3 (Section 4.6.2), in order to test its performance on a shorter-context task. We perform full finetuning, using 2,048 bp sequences centred on each labelled 250 bp window as inputs. To ensure a fair comparison between all gLMs, we test three learning rate settings for each model and use chromosome 4 to choose the best parameters (see Section 4.6). Our finetuned Botanic1-S model reaches a mean chromosome 5 macro average precision of 0.72 across three seeds, beating NTv3-652M (0.69), NTv3-106M (0.68), AgroNT (0.63) at a lower compute budget than all except NTv3-106M (Figure 5a,b), and also beating a bespoke architecture trained from scratch, DeepCistrome [66], which reaches 0.62 in our reproduction. In order to measure the added value of pre-training, we also finetune a Botanic1-S-like model with random weights instead of the pre-trained checkpoint on the same data. The same Botanic1-S architecture trained from random initialisation reaches 0.69 on chromosome 4, compared with 0.72 for the pre-trained checkpoint (Figure 5b). As an illustration of the task, Figure 5c shows the predicted tracks at a 40 kbp chromosome 5 locus, with model scores aligned to observed MYB-related binding windows.

**Figure 5.**
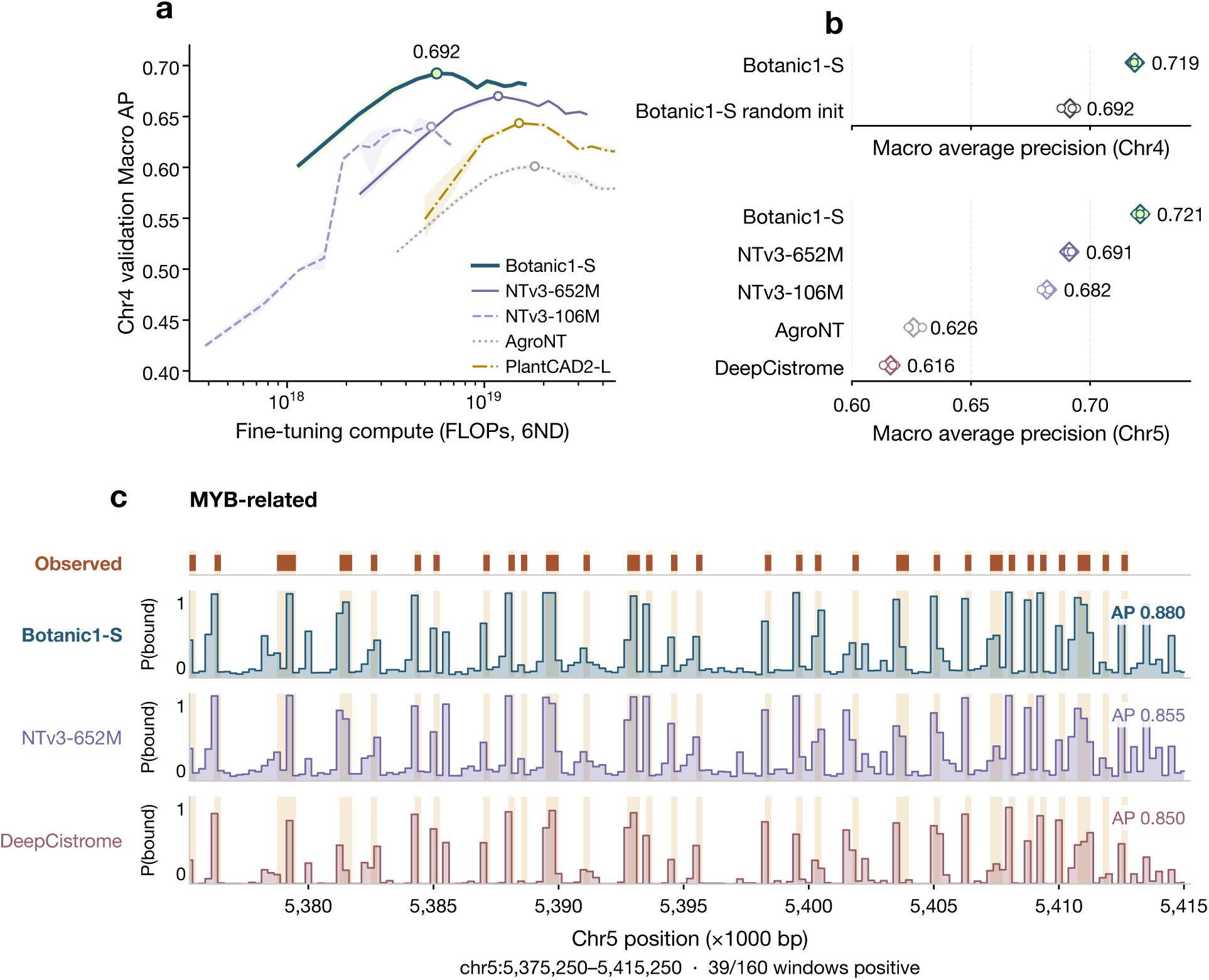
Genome-wide transcription factor family binding in Arabidopsis thaliana. **a**, Chromosome 4 validation macro average precision against estimated fine-tuning compute, 6ND. Lines show the mean across three seeds with shaded areas covering the minimum and maximum values, circles mark the peak of each mean curve. These values are not reverse-complement averages but single-orientation, unlike panel **b**. **b**, Macro average precision across three seeds; the top panel shows chromosome 4 validation for Botanic1-S only, including the random-weights initialised model, to illustrate the impact of pre-training; the bottom panel shows the test results on chromosome 5. **c**, Seed-averaged prediction tracks for the MYB-related family across one 40 kbp chromosome 5 region. In the top row, darker bars mark positive windows. AP values to the right report the average precision across the 160 windows.

Botanic1-S is also first across different trainings using subsampled data (Supplementary Figure S21). Finally, we find that fine-tuning of the larger Botanic1-M model, here the released 8,192 bp checkpoint, does not improve performance on this task, yielding a chromosome 5 macro average precision of 0.72.

To understand why Botanic1-S performance surpasses the bespoke architecture trained from scratch, we compare the prevalence of known 6-mer motifs for different TFs in the highest scoring vs the lowest scoring positive-labelled windows (Supplementary Figure S22). DeepCistrome shows stronger enrichment for the most common motif in all five families, and more generally in 25 of all 30 motifs. This suggests that the performance gain comes from other less subtle factors in the wider context of the target window, potentially complex genomic grammar across hundreds or thousands of base pairs, which is learned during pre-training.

#### 2.3.3 Botanic1 learns new capabilities through large context training without catastrophic forgetting of short context

We evaluate the benefits and downsides of longer context for gLMs. We compare the Botanic1-S model trained on 8 kbp windows with its versions obtained by progressive extension of the context on windows of length up to 131 kbp (2^17^ bp), every stage at the same token budget, writing the five stages Botanic1-S0 for the 8 kbp model through Botanic1-S4 for the longest (see Section 4.1.5, Supplementary Figure S4).

##### 2.3.3.1 Botanic1-S benefits from context-extension continued pre-training, and more context at evaluation

We test whether the further context-extended models can exploit the additional context they are given, and whether context-extension through continued pre-training (CPT) hurts short-context performance and improves it at longer evaluation lengths, by measuring pseudo-perplexity (PPPL) [67] on held-out sequences of various lengths. PPPL is the MLM analogue of perplexity (Section 4.4.1); lower is better.

The PPPL metric we report downweighs repetitive positions like our MLM training loss: each position is weighted by 1 (1 w)f, where *f* is the repeat binary flag from the soft-mask annotation and *w* = 0.1 (see Section 4.1.6). All comparisons are reported as paired deltas rather than absolute PPPL. To isolate the effect of giving a model more context, we fix the scored positions to a central 2^13^ bp subwindow and add surrounding context around it; we call this *nested-context* evaluation. Every pairing of training and evaluation length in the PPPL landscape of Figure 6a scores the same 30 positions in the same 1,251 sequences, so any two are directly comparable. The two margins read that landscape along its axes: the top margin takes each model against its own 2^13^ bp evaluation, the left margin takes each stage against Botanic1-S0 at the same evaluation window, both as a percentage of the baseline the delta is taken against.

**Figure 6.**
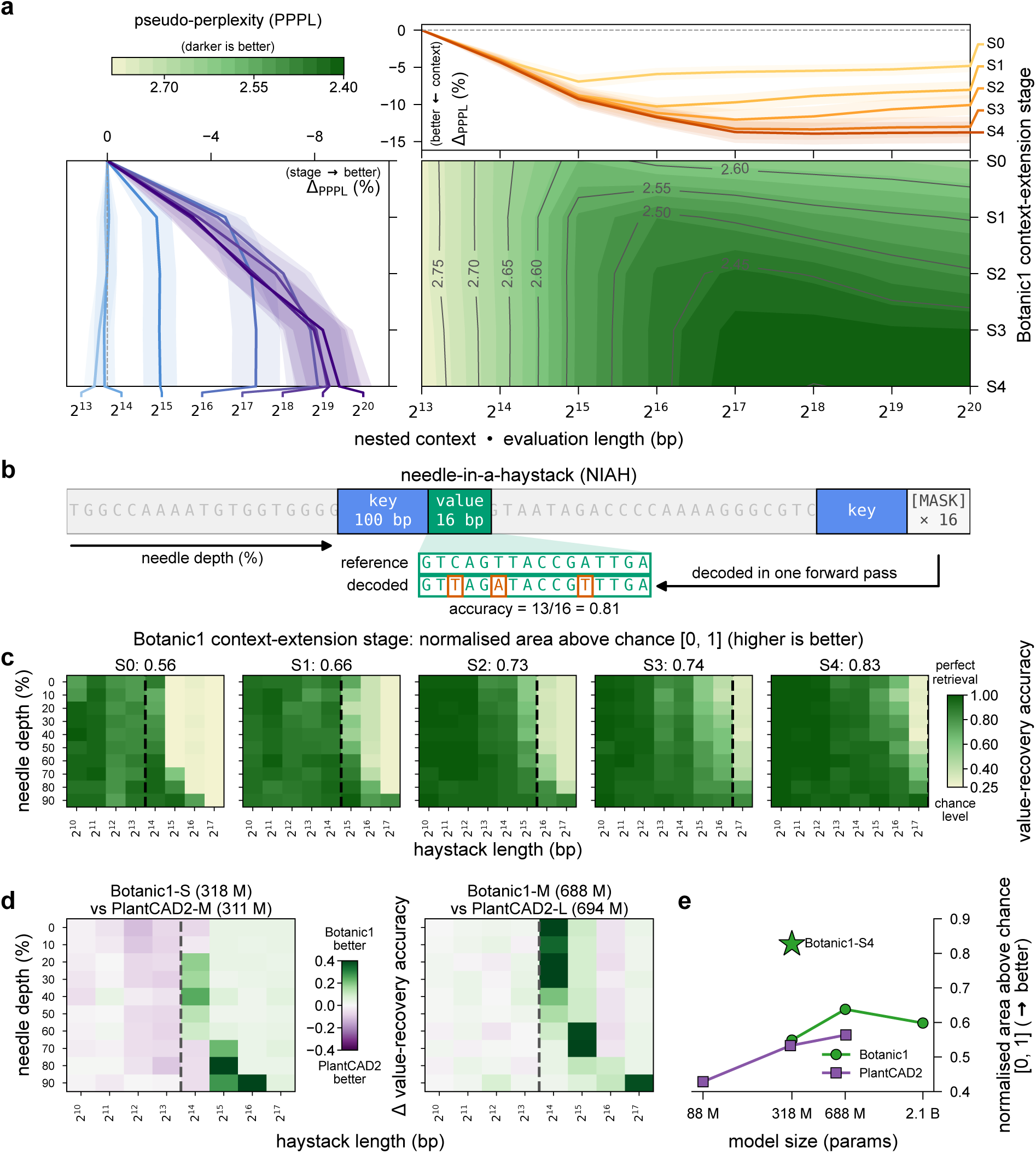
Long-context behaviour of the iso-token context-extended Botanic1-S models. **a**, Nested-context pseudo-perplexity, repeat-weighted (Section 4.4.1); lower is better. The 30 scored positions per sequence stay in the same central 2^13^ bp subwindow while surrounding context grows. Filled contour, corpus PPPL against evaluation length (*x*) and context-extension stage (*y*), Botanic1-S0 to Botanic1-S4 being the 2^13^ to 2^17^ bp stages. Top margin, each stage against its own 2^13^ bp evaluation; left margin, each stage against Botanic1-S0 at the same evaluation length. Both report a percentage of the baseline they are taken against and start from the zero that baseline defines; ribbons are simultaneous 95% intervals from a sequence-level paired bootstrap (10,000 resamples, common draws across models), so family-wise error is controlled at *α*_FWER_ = 5%. Held-out generalisation at each length separately is in Supplementary Figure S27. **b**, Task construction: a 100 bp key and a 16 bp value are planted at a controlled depth in a random haystack; the key is re-inserted at the end of the context with the value masked and all 16 value positions are decoded in one forward pass. **c**, Accuracy against haystack length and needle depth after each context-extension stage, Botanic1-S0 to Botanic1-S4; the dashed line marks that stage’s maximum training length. **d**, Botanic minus PlantCAD2 at matched parameter counts, both pretrained at 8,192 bp; green favours Botanic. **e**, Each grid summarised as the normalised area above chance against parameter count; the star is Botanic1-S4.

The top margin shows that every model is significantly better with surrounding context than on the central subwindow alone, and that the curves are U-shaped: PPPL reaches an optimum as surrounding context is added, then plateaus and eventually worsens. The further a model is context-extended the better that optimum and the longer the window at which it is reached. The landscape puts those optima at 2.58, 2.49, 2.44, 2.41 and 2.40, the top margin at a significant 6.9% to 13.9% reduction on the central subwindow, reached at 2^15^, 2^16^, 2^17^, 2^18^ and 2^18^ bp. Models extrapolate, reaching their optimum at a window four times the one they are trained on, except the Botanic1-S4, whose PPPL peaks at 2^18^ bp.

Taking the evaluation axis from its short end, at 2^13^ and 2^14^ bp the PPPL iso-contours are parallel to the context-extension axis, so PPPL depends on evaluation length alone and not on which stage we choose: PPPL ≈ 2.78 at 2^13^ bp and PPPL ≈ 2.67 at 2^14^ bp. We cannot detect any additional significant difference from Botanic1-S0 at these evaluation lengths apart from a mild 0.50% regression for Botanic1-S4. Context-extension CPT therefore costs little on short context.

At longer evaluation lengths from 2^15^ bp onwards, the iso-PPPL contours tilt as further context-extended models perform better, every stage now beating Botanic1-S0 significantly. The left margin gives the size of that effect at each evaluation window: at 2^15^ and 2^16^ bp the four stages gain much the same, 1.9% to 2.0% and 4.6% to 5.7%. From 2^17^ bp onwards the gain grows with the stage, 4.3%, 6.8%, 7.8% and 8.1% at 2^17^ bp, and the iso-contours turn almost parallel to the evaluation length axis: the stage now sets most of the PPPL. So at longer evaluation lengths PPPL improves monotonically, or stays on par, with further context-extension CPT.

The BiMamba2 backbone of the Botanic models costs time linear with respect to sequence length, so each doubling of the evaluation window roughly doubles the inference cost, making window length a test-time scaling axis: more compute buys lower PPPL. That axis stops paying beyond 2^17^ bp, where widening the window brings our context-extended models little improvement or even slight degradation, as the U-shaped curves above already show. Continued pre-training on longer context makes the axis more effective and extends it further: in our case Botanic1-S4 is best or on par across the useful 2^14^ to 2^17^ bp range of evaluation lengths, so additional context keeps improving its PPPL, and it keeps the best PPPL of the four stages out to 2^20^ bp if longer sequences have to be processed at inference anyway.

Scoring sequences of each source length instead leads to the same conclusions (Supplementary Figure S27): further context-extended models reach monotonically better or on-par PPPL, the one exception being the barely significant S3 → S4 step at 2^15^ bp, which raises it by 0.28% though it stays better than Botanic1-S0. That evaluation cannot compare PPPL across source lengths, which the nested-context one can.

##### 2.3.3.2 Context-extension improves Botanic1-S retrieval ability

Needle-in-a-haystack (NIAH) evaluation probes long-range modelling from a different angle: it tests associative recall, the ability to retrieve a specific needle from an ever larger haystack given a cue. We adopt here a key-value variant analogous to LLM NIAH benchmarks [68]: a key-value pair is inserted at a controlled depth within an arbitrary input; the 100 bp key is then repeated at the end of the context with its 16 bp value masked, and performance is measured by how much of the original value the model recovers, resolved by insertion depth and context length in a heatmap (Section 4.4.2). Throughout, we summarise one such heatmap by a single normalised score in [0, 1], where 0 is chance and 1 is perfect retrieval; all NIAH numbers below are on this scale. We evaluate the five context-extension stages and benchmark Botanic against PlantCAD2 [33], both pre-trained on 8,192 bp windows. Each context-extended model retrieves reliably up to roughly its own training context and degrades towards chance beyond it, and the score grows monotonically with the maximum context seen during pre-training, from 0.56 at Botanic1-S0 to 0.83 at Botanic1-S4 (Figure 6c). At matched parameter count and without context extension, Botanic1 retrieves better than the PlantCAD2 family (Figure 6d,e): 0.55 against 0.53 near 300M, and 0.64 against 0.56 near 650M. PlantCAD2 scores rise monotonically with capacity (0.43, 0.53 and 0.56 at 88M, 311M and 694M), whereas Botanic does improve from Botanic1-S to Botanic1-M but falls back slightly at Botanic1-L (0.60). The strongest model overall on this task is not the largest but the most context-extended: Botanic1-S4 scores 0.83 (the star in Figure 6e), above every point on either size curve and well clear of the best PlantCAD2 model (0.56 at 694M) at under half the parameters.

##### 2.3.3.3 Chromatin accessibility prediction benefits from longer context

Chromatin accessibility is a key determinant of gene expression regulation and is shaped in part by long-range interactions along the DNA [69, 70]. This makes chromatin accessibility prediction a natural task for assessing the benefit of extending the input context of genomic language models.

Zhai et al. [33] showed that longer context windows improve PlantCAD2-S’s performance on accessible chromatin region (ACR) prediction. However, this benefit was only demonstrated up to a 4,600 bp context, leaving open whether it persists at longer ranges. To test whether Botanic1-S can exploit its extended context, we evaluate its performance on the same task under several training strategies (see Section 4.7), using PlantCAD2-S and PlantCAD2-M as baselines.

We start with a training protocol employing LoRA [35] fine-tuning and a classifier head on hidden states mean-pooled over the full window width (Supplementary Figure S29). We apply this training strategy to PlantCAD2-S, PlantCAD2-M, and the two Botanic1-S variants trained at 8k and 128k context (denoted Botanic1-S-8k and Botanic1-S-128k, respectively). The range of tested context lengths is extended to include 8,000 bp. Zhai et al. [33] report an AUPRC of 0.665 at 600 bp rising to 0.707 at 4,600 bp for PlantCAD2-S on this task (in Figure 5G). Under our slightly different training protocol (different held-out chromosomes, classification head and effective learning rate (Section 4.7), the mean per-cell-type AUPRC of PlantCAD2-S rises from 0.664 to 0.691 over the same range (median 0.685 to 0.718). We thus reproduce the reported increase up to 4,600 bp context for PlantCAD2-S, and observe a decrease at 8,000 bp. a similar non-monotonic trend is observed for the other models, with a performance peak at 2,600 bp for PlantCAD2-M and at 1,600 bp for both Botanic1-S variants. We hypothesise that the performance decrease at longer contexts may be due to the relevant information being diluted by the mean pooling over the full input window. To check this hypothesis, we adjust the training strategy to focus the pooling on a 101 bp window centred on the ACR peak, and retrain Botanic1-S-128k with this strategy. Central-window pooling monotonically improves Botanic1-S-128k’s median AUPRC across the tested contexts, reaching 0.77 at 32,000 bp, against a peak of 0.74 for both full-window PlantCAD2-M and Botanic1-S-128k (Figure 7a). It thus seems that chromatin accessibility prediction does benefit from longer context, provided adjustments are made to the training strategy.

**Figure 7.**
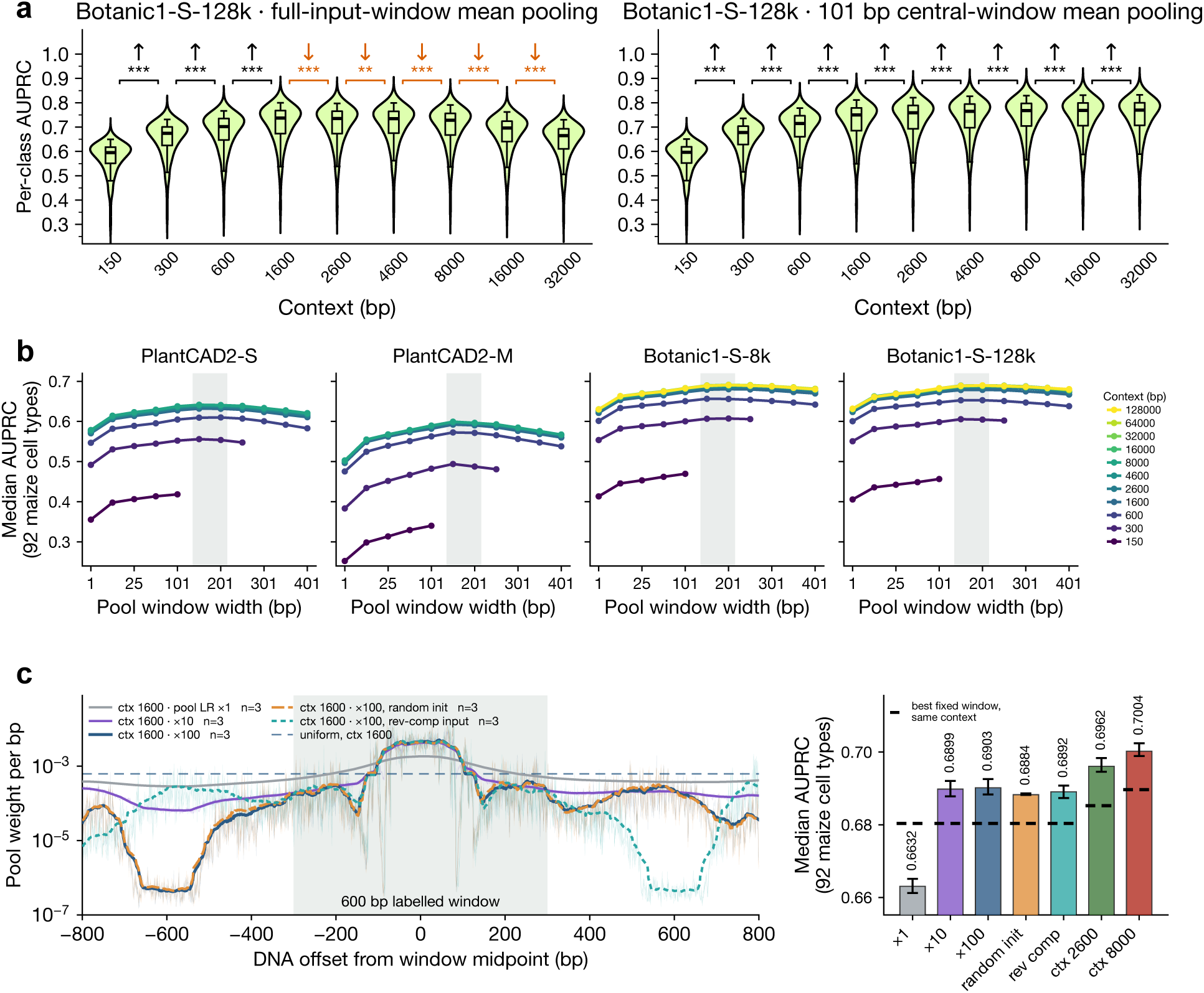
Chromatin accessibility prediction with Botanic1 models using window pooling. **a**, Performance of Botanic1-S-128k on cell-type-specific accessible chromatin regions prediction, using LoRA fine-tuning on fullinput-window mean-pooled embeddings (left) and on 101 bp central-window mean-pooled embeddings (right). For each context size, AUPRC is assessed for each of the 92 cell types and the resulting distribution is shown as a violin plot. Brackets between consecutive contexts show the direction of the shift in median AUPRC when it is statistically significant (an upward arrow shows a better performance at the longer context), and the significance marker after Holm-Bonferroni correction within the panel (class-paired Wilcoxon signed-rank test; * *p* < 0.05, ** *p* < 0.01, *** *p* < 0.001; ns otherwise). Central pooling avoids the long-context degradation seen with full-window averaging. **b**, Frozen-embedding probe on the same task, with each backbone kept frozen and only a single ESM-like head trained on hidden states mean-pooled over a peak-centred window. Each panel is one backbone, each line is the median per-class AUPRC against pooling window width, one line per input context from 150 bp (darkest) to 128,000 bp (lightest). Full results, with every cell value and its significance marker, are in Supplementary Figure S30; the same sweep under a linear head is in Supplementary Figure S31. **c**, Learned positional pooling weights on Botanic1-S-8k at 1,600 bp, three seeds per setting. The curves and values are shown for a uniform initialisation of the weights unless otherwise specified. Left: converged pool weight per base, log scale. Thicker lines represent the 51 bp rolling mean of each learned profile, while thinner lines are the raw weights without averaging. “x1”, “x10” and “x100” indicate the positional weights’ learning rate relative to the classifier head’s, the dashed line is the initialisation value 1/L. Right: median per-class AUPRC of each learned pool experiment against the best performance at the same context and fixed width pooling from panel **b**. 2,600 and 8,000 bp learned profiles are drawn in Supplementary Figure S32.

To characterise this further, we assess the performance of Botanic1-S over different context and pooling window sizes. For efficiency, we conduct this analysis without fine-tuning, using frozen embeddings pooled at various central window widths and a small ESM-style classification head trained on top (Figure 7b, Supplementary Figure S30). Median AUPRC rises steeply between 150 and ~ 1,600 bp of context for all backbones and keeps increasing at longer context, although to a lower extent, up to 16,000 bp context where it starts to plateau. PlantCAD2-M, whose performance is comparable to that of Botanic1-S-8k and Botanic1-S-128k with LoRA fine-tuning strategy, underperforms with a frozen backbone, with median AUPRCs even lower than those of the smaller PlantCAD2-S model across all cells. We also observe that Botanic1-S-8k and Botanic1-S-128k have similar performance up to 16,000 bp context, even on context lengths far greater than the window length Botanic1-S-8k is pre-trained on. The best performance on this task is observed for pooling window sizes between 101 and 201 bp, confirming that focusing on the embeddings at the peak region yields the best results.

We choose PlantCAD2-M as a reference to approximately match Botanic1-S’s model size, but it performs unexpectedly poorly. Analysing the pooled features of PlantCAD2-M, we find a possible explanation. PlantCAD2-M’s 1,024-dimensional representations can be effectively compressed into a handful of directions, with the first ten carrying 97.6% of the variance. The leading one alone carries 63%, against 12.9% for PlantCAD2-S and 40.7% for Botanic1-S-128k, an ordering that matches the AUPRC in this task, and more globally their 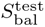 ranking (see Figure 3d). This theory would explain why a two-layer head recovers +0.165 AUPRC on PlantCAD2-M against +0.039 on Botanic1-S-128k: a collapsed representation is more readily recovered by a nonlinear classification than a linear head.

Finally, we set out to discover the best pooling strategy by replacing the fixed central window with positional weights learned during training. These weights are optimised with a separate learning-rate multiplier relative to the head: 1 , 10 or 100 . All learned masks converge towards a central peak higher than the uniform initialisation and progressively smaller weights moving away from the central position (Figure 7c). With at least 10× learning rate, learned positional pooling beats any fixed-width pooling across all context lengths: 0.690 against 0.680 with 1,600 bp context length, 0.696 against 0.685 at 2,600 bp, and 0.700 against 0.690 at 8,000 bp, each a mean over three seeds.

The learned profile is stable across context lengths, batch ordering, and random seeds (Supplementary Figure S32). One reverse complementation experiment shows that the shape is strand-dependent, although the task itself should not be. To understand this asymmetry and the unexpected ~ 600 bp pseudo-periodicity in the learned weights profiles upstream of the central peak, we investigate further the distribution of neighbouring ACR peaks at positions away from the central peak along the sequences. Looking at the fraction of rows belonging to an ACR at each position reveals a similar asymmetry in the negative dataset, with a rather flat and continuous profile downstream of the central position, but a broken profile on the upstream side, with what appears to be pseudo-periodic gaps every ~ 600 bp (Supplementary Figure S33). Examining how the negative dataset is built, we realise that while positive windows are built around 501 bp-wide ACR peaks, 600 bp-wide negative windows are cut consecutively from regions sitting in between ACR peaks, starting exactly 50 bp after the end of the ACR peak starting the interval. By construction, the negative windows thus all have an ACR peak sitting at positions [ 851 600k, 350 600k], *k* ∈ Z from their midpoint (ACR peaks are 501 bp wide, plus the 50 bp offset and the 300 bp half window length). Aligning all the negative sequences thus shows a heightened plateau at positions [851; 350] (corresponding to the left-most windows cut out from the inter-peak regions), another heightened plateau at [1451; 950] (corresponding to the second left-most windows), etc. This pattern is not observed downstream of the central window because the negative sequences are cut from left to right within each inter-peak gap. Therefore, the right-most negative window in the gap is at a variable distance from the next ACR (depending on the total gap length modulo 600) and we observe no periodicity. This observation is consistent with the learned pooling weights profiles we obtain: lower weights seem to be assigned to regions corresponding to positions enriched in ACR peaks in the negative dataset, probably thus contributing to decreasing the number of false positives.

Taken together, these experiments show that Botanic1-S models can exploit long sequence context for chromatin accessibility prediction, even without further fine-tuning of the backbone to this specific task. They also show that careful pooling is necessary to make the best use of the sequence embeddings when extending the input to longer contexts.

### 2.4 Sparse features of Botanic1 are readable genomic concepts

Sparse autoencoders [71, 38, 72] provide a modern approach for examining a model’s internal representations and identifying concepts it uses while processing input sequences. An autoencoder decomposes a token representation from the residual stream of one model layer into activations of discrete features. This decomposition is derived only from the model and activations collected from forward passes over a small dataset; it uses no biological annotations of those inputs. Some of these features carry meaningful biological information, which we validate by comparing their activation sets with known biological annotations. All results in this section describe features of one SAE trained on layer 35 of Botanic1-M. The training and methodology are described in Section 4.8 and summarised in Figure 8.

**Figure 8.**
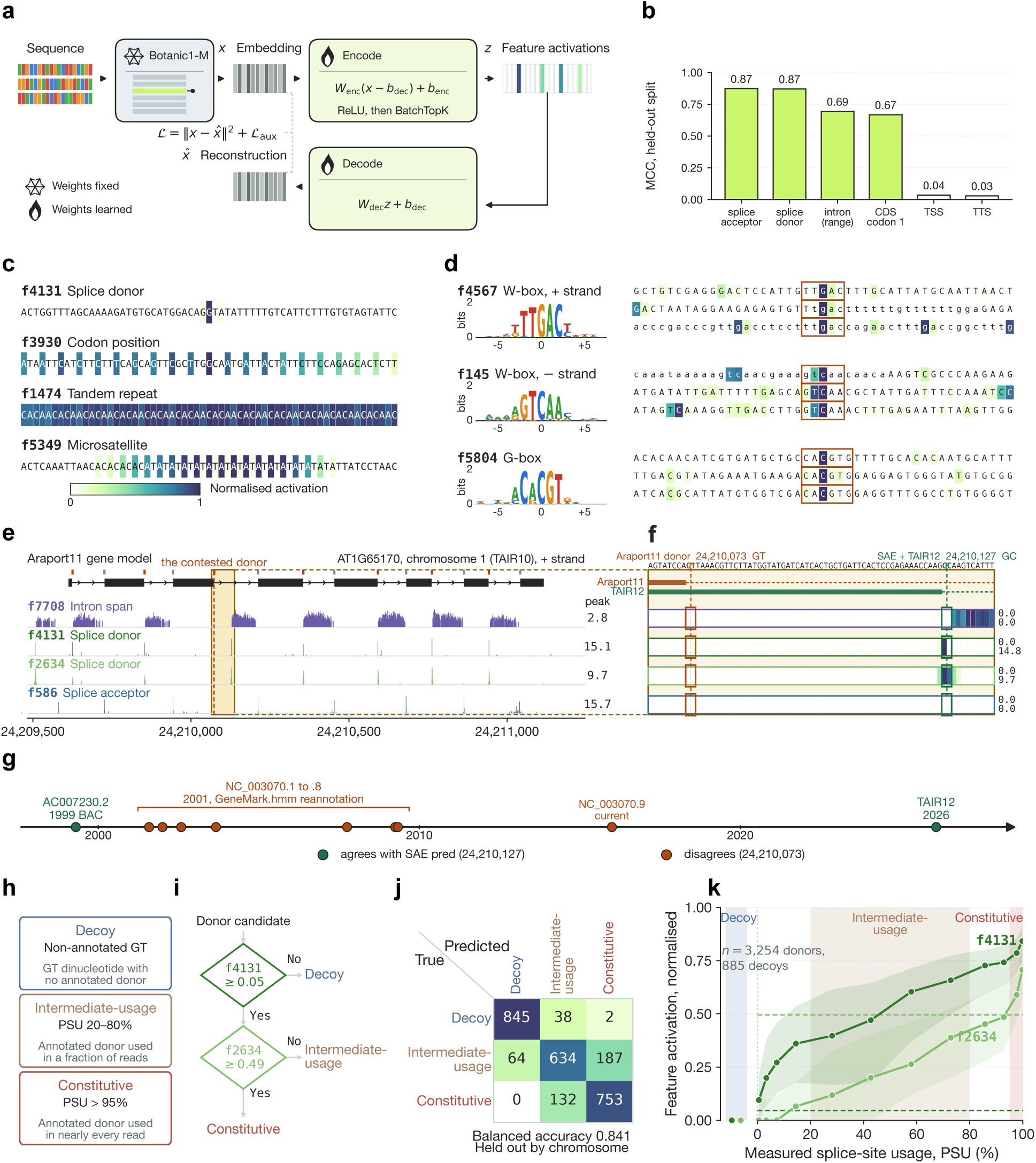
Features of SAE fitted to a frozen Botanic1 model detect splice boundaries, intron regions and codon positions, named cis-elements, and find errors in TAIR10/Araport11 annotations. **a**, Scheme of SAE training for interpretability analysis. The Botanic1 model is frozen, and no genomic annotations are used. **b**, The best single thresholded feature per target as a baseresolution classifier, MCC is the Matthews correlation coefficient. CDS codon 1 is the first base of each codon inside coding sequence. **c**, Per-base activation examples for top firings of four features show different patterns: isolated peak, 3-periodic peaks, region-based, alternating activations. **d**, Example of features found to correlate to named cis-elements. Left: motif built from feature’s 32 top firings (no motif database used). Right: three of that feature’s top windows, aligned on the top-firing (in the orange box). **e**, Four features over the <u>AT1G65170</u> gene compared to its Araport11 model, donor ticks are indicated in orange and acceptor ticks in grey; the feature track height is proportional to the activation. The dashed line indicates the annotated donor site at which both donor features are silent. The shaded rectangle marks the 73 bp window that panel **f** zooms into. **f**, Zoomed-in region near contested donor in AT1G65170. The annotated donor (orange, GT) and the one the SAE features predict and TAIR12 places 54 bp away (green, GC) are boxed in each feature row; numbers on the right are the activations at the two sites. The Araport11 and TAIR12 exon/intron bars at the top align with the Araport11 gene model in **e**. **g**, Published history of that donor drawn as circles on a time arrow: 1999 BAC submission, 2001 GeneMark.hmm reannotation and seven follow-ups gathered under a bracket (NC 003070.1 to .8), the standing RefSeq NC 003070.9, and the 2026 TAIR12 release. Green circles agree with the SAE prediction, orange circles do not. The SAE prediction matches the 1999 and the 2026 annotation and not the twenty-five years between them. **h**, The three donor splice site classes used in **i** to **k**: decoys are non-annotated GT dinucleotides, intermediate-usage donors are used in 20% to 80% of reads, and constitutive donors are used in more than 95%. **i**, Predicting the class of a donor splice site candidate follows a two-layer decision tree over two dictionary features. The thresholds shown are the median value chosen over the five cross-validation folds, expressed on the normalised scale used in **k** (each feature’s raw activation divided by its own 99th percentile). **j**, Confusion matrix of the predictions, with counts pooled over the five held-out chromosomes. Rows are the true class, columns the predicted class; the diagonal glyph in the top-left corner labels the two axes. **k**, The two selected features’ activation against PSU, over 3,254 annotated donors. The line marks the median within each PSU bin and the shaded band the 20th to 80th percentile. Activations for each feature are normalised on its own 99th percentile over these donors. Decoys have no PSU value and are drawn as a separate coloured slab to the left of the axis. The two horizontal dashed lines are the fitted thresholds from **i** on that same normalised scale (0.05 for f4131, 0.49 for f2634). Coloured spans mark the three classes used in the classification task, in the same palette as **h**, **i** and **j**.

#### 2.4.1 Base and region-level features

We identify both base-level and region-level features. For example, base-level feature f4131 recognises “splicing donor” genomic annotations (Matthews correlation coefficient (MCC) 0.871 at its validation-selected threshold) and f586 recognises “splicing acceptor” annotations (MCC 0.873), while region-level feature f7708 recognises intron-region annotations (MCC 0.694) (see Figure 8b). These features are the best detectors of their corresponding genomic annotations in all four annotated genomes (from species *Arabidopsis thaliana*, *Oryza sativa*, *Solanum lycopersicum* and *Zea mays*), with MCC 0.821 to 0.937 for the two splice-related features and 0.650 to 0.741 for the intron-related feature, showing that these features are general and applicable to multiple species.

Features may even be used as classifiers: a single threshold (identified on a validation set) on f586 gives a precision of 0.845 and a recall of 0.901 at a prevalence of 6 10*^−^*^4^ when used to predict “splicing acceptor” sites, while another threshold on f4131 gives 0.876 precision and 0.866 recall on “splicing donor” sites.

The best feature for “exon region” annotations reaches only MCC 0.32, despite exons carrying strong sequence signals (three-base periodicity, biased amino-acid usage, . . . ). After further investigation, we find that exon signal is split into three strong features corresponding to the three within-codon positions in exons: features f3930, f2618 and f1200 have MCCs of 0.668, 0.542 and 0.633 with the first, second and third codon bases, respectively (Figure 8b).

Conversely, no strong feature can be identified for transcription start site (TSS) and transcription termination site (TTS): the best features have MCC values of 0.035 and 0.029 respectively (Figure 8b), which is consistent with these sites having no sharp sequence consensus in plants.

#### 2.4.2 A splice feature anticipates a change in the reference annotation

During feature validation, we observe mismatches between the activation of some features and the annotations reported at the <u>AT1G65170</u> locus of *A. thaliana*. While TAIR10 and Araport11 [73, 74] annotate a splicing donor for intron 3 at position chr1:24,210,073, both top donor features (f4131 and f2634) give 0.00 activation at that position. However, they do activate 54 bp downstream at position chr1:24,210,127 (f4131 at 14.79, f2634 at 9.66) (see Figure 8e,f). Similarly, the intron-region feature f7708 reads the 54 contested bases between the annotated splicing donor and the splicing donor feature activation position at exactly 0.00, against per-intron means of 1.04 to 1.67 across the six other introns of the gene, and starts activating 2 bp downstream of the base the “splicing donor” features pick out.

Given the convergence between the predictions from all three features, we examine the annotation history of <u>AT1G65170</u>. We find that the original 1999 bacterial artificial chromosome (BAC) submission (AC007230.2, clone T23K8) [75, 76] annotated the splicing donor at chr1:24,210,127 (where our “splicing donor” features activate). This annotation was modified in 2001 (NC 003070.1, a GeneMark.hmm reannotation [77]) to chr1:24,210,073. The annotation did not change through nine RefSeq releases until 2026, when TAIR12 [78] moved it back to chr1:24,210,127 (see Figure 8g).*^*^*

No genomic annotations are used to train the Botanic1 model or the SAEs; and the mapping of SAE features with annotations, in particular this donor feature, only uses TAIR10 annotations from *A. thaliana*.

These results suggest that SAE features may be used to improve and correct genome annotations.

#### 2.4.3 Alternative splice sites are identifiable from two features

The splicing prediction problem is more complex than multi-class labelling base pairs into categories of splice sites because alternative splicing makes splice-site usage inherently context-dependent. Donor and acceptor sites are not systematically used, even within the same tissue and developmental stage. Motivated by the observation of unequal activation levels of donor-associated features among splice sites within the same gene, we ask whether these patterns might be a sign of the model capturing some representation of alternative splicing.

For this experiment we use PastDB [79] (Plant Alternative Splicing and Transcription Database), which quantifies alternative 5’ splice-site events from RNA-seq across hundreds of *A. thaliana* samples spanning tissues, developmental stages, and environmental conditions. For each splice-site one may compute the percent splice-site usage (PSU) across these reads. From these data we construct three classes (Figure 8h): constitutive donors (PSU > 95%), intermediate-usage donors (PSU between 20% and 80%) and decoys (non-annotated GT dinucleotides). We use a simple two-level decision tree over dictionary features (Figure 8i). a first feature *f_A_* separates donors from decoys, and a second feature *f_B_* separates constitutive from intermediate-usage donor sites. We evaluate this classifier on chromosome-cross-validation, fitting both features and thresholds on four chromosomes and evaluating on the fifth. The three splice site usage classes are balanced at 885 sites each, and all candidate sites contain the canonical GT dinucleotide. The two-feature classifier reaches a balanced accuracy of 0.841 [0.827, 0.855], compared with 0.787 for the best single dictionary feature, 0.643 for the same classifier using positionally weighted and Markov-model-based donor-site scores, and 0.846 for an unconstrained decision tree of depth 2 (free to choose the features, thresholds, and order of the two splits). Per-class recall is 0.955, 0.851, and 0.716 for decoy, constitutive, and intermediate-usage donors, respectively (Figure 8j). All five folds select the same pair of features, f4131 and f2634, which are also the two donor-associated features highlighted in Figure 8e.

Both features are associated with splice-site usage in a monotonic fashion, beyond the discrete classes used for training (Figure 8k). Their activations correlate strongly with PSU across donor sites (Spearman ρ = 0.753 for f4131 and ρ = 0.759 for f2634), with median activation increasing steadily across the PSU range, and dropping to zero at most decoy sites. To test whether donor/decoy discrimination relies on gene-level sequence context, we repeat the analysis using non-splice-site GT dinucleotides from the same gene as each donor, rather than decoys sampled elsewhere in the genome. Balanced accuracy decreases only modestly, from 0.841 to 0.818 [0.804, 0.832], while discrimination between constitutive and intermediate-usage donors remains nearly unchanged. As a second control, we randomly permute the PSU-derived labels among donor sites, destroying the relationship between feature activation and splice-site usage while preserving the distinction between donor and decoy sites. Performance drops to 0.637 ± 0.007.

Splice-site recognition has already been shown in prior plant genomic models [34, 33, 31] (see also our results in Section 2.3.1.2). The quantitative prediction of alternative splice-site usage was demonstrated previously by specialised models like Pangolin [80] trained on RNA-seq-derived usage measurements from four mammalian species. Also, some splice-site usage results were demonstrated by AlphaGenome [81], for which they trained a dedicated output head using observed splice-site usage data. To the best of our knowledge, our results are the first evidence that representations of a genomic language model pre-trained without supervision contain individual sparse features that encode quantitative splice-site usage.

### 2.5 Applications

#### 2.5.1 Botanic1 for zero-shot variant effect prediction

Botanic1 achieves state-of-the-art zero-shot performance for variant effect prediction. We first test whether its likelihood ratio scores capture expected relationships with allele frequency and variant class, then evaluate it on a new benchmark of experimentally validated causal variants rather than proxies for functional impact.

##### 2.5.1.1 Zero shot scores are correlated with minor allele frequency and variant classification

After confirming the expected monotonic relationship between the missense-to-synonymous ratio and allele frequency (Supplementary Figure S34), we reproduce Figure 7 of Mendoza-Revilla et al. [31] on the *A. thaliana* 1001 Genomes panel. We score all 6,491,238 biallelic single nucleotide polymorphisms (SNPs) with the three Botanic1 sizes, as well as AgroNT and GPN, the two models evaluated in the original study. First, we measure the Pearson correlation between minor allele frequency (MAF) and each model’s zero-shot LLR. LLR score calculation, variant panels, scoring windows and the block-jackknife uncertainty protocol are described in Section 4.9. The correlation increases monotonically with Botanic1 model size: r(LLR, MAF) = 0.0932 0.0985 0.1047 for Botanic1-S, Botanic1-M and Botanic1-L, all models surpassing both GPN (0.0890) and AgroNT (0.0595). On this panel the original study reports *r* = 0.05 for AgroNT and 0.07 for GPN [31]; our reproduction gives 0.0595 and 0.0890 (Section 4.9).

We next evaluate whether LLR scores can directly distinguish missense from synonymous mutations and observe the same model ordering, with the top 3 being taken by Botanic1 models (Figure 9a). Botanic1-S achieves a higher AUROC than GPN, although the difference is not significant, whereas Botanic1-M and Botanic1-L show clear improvements.

**Figure 9.**
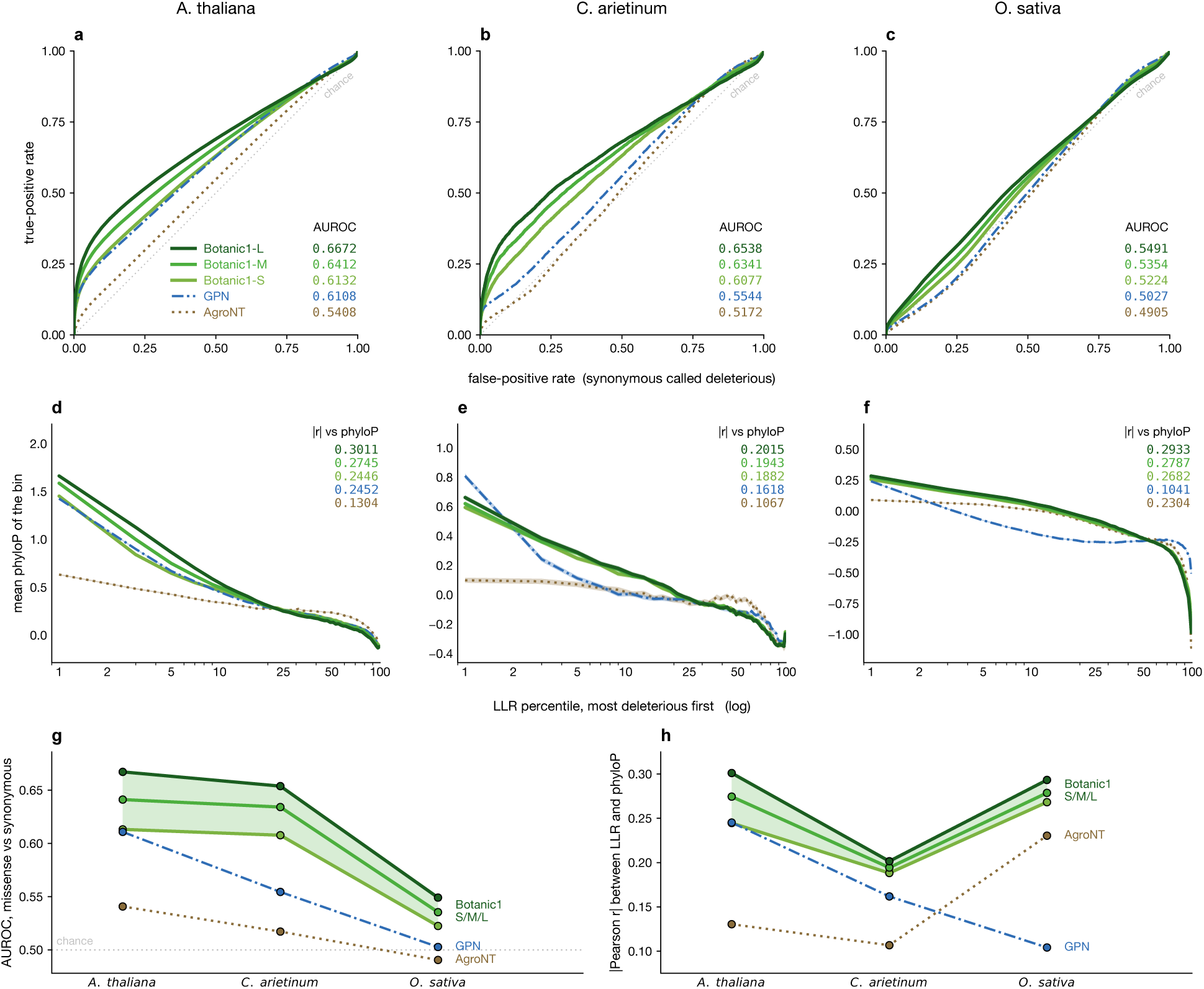
Zero-shot likelihood against variant categories and conservation scores. All panels use 6,491,238 *Arabidopsis thaliana* SNPs (1001 Genomes), 880,345 *Cicer arietinum* SNPs (CicerSeq, CDC Frontier v1) and 6,494,573 *Oryza sativa* SNPs (RiceVarMap2.0), in that order across the columns, which is increasing phylogenetic distance from the Brassicales. **a** to **c**, Each model’s LLR as a classifier of missense (957,062 variants) vs synonymous (690,484), as categorised by Ensembl VEP on *A. thaliana*: each panel shows the ROC curves on one species. **d** to **f**, Agreement between LLR and MSA-derived phyloP, with the same species order as **a** to **c**. The variants are ranked by each model’s LLR and cut into fifty equal bins, and the mean phyloP of each bin is plotted against its percentile, with a log x-axis. a model that separates deleterious from neutral variants descends from left to right; a model with no signal is flat. **g**, Summary of **a** to **c** as AUROC against species. **h**, Summary of **d** to **f** as Pearson *r* between LLR and phyloP against species. In both **g** and **h**, the Botanic1 ladder is drawn as a band from its weakest to its strongest size. phyloP compares models within a species well, but only loosely between species: the three tracks are separate PlantRegMap-derived alignments with different score distributions, covering 93.8%, 96.3% and 92.5% of each variant panel.

Following Mendoza-Revilla et al. [31], we consider a third proxy for functional impact: the multiple-sequencealignment-derived PhyloP score. Here too, Botanic1 outperforms both GPN and AgroNT, with performance improving consistently with model size (Figure 9d, Supplementary Figure S35). Our measures of AgroNT’s LLR and phyloP on *A. thaliana*, *r* = 0.130, matches the 0.13 reported in the original study (Figure 7d); for GPN we measure 0.245 against the reported 0.20. Agreement with PhyloP is substantially higher for coding than for non-coding variants (Supplementary Figure S36a), suggesting that modelling regulatory variation remains an important area for improvement.

In the original study [31], the strong performance of GPN was attributed to its training dataset being focused on the Brassicales family, and this advantage did not persist when evaluated on rice, where the more phylogenetically diverse AgroNT performed better. Botanic1 is trained on 320 species (the species of the 326-genome corpus that contribute training windows, see Section 4.2) spanning the land plants with a sequenced genome, yet still outperforms both models on *A. thaliana*. We then test whether this advantage persists outside of the Brassicales order.

We repeat the same zero-shot evaluations on chickpea (*Cicer arietinum*) and rice (*Oryza sativa*), with rice being substantially more phylogenetically distant from both Arabidopsis and chickpea than the two eudicots are from each other. On *O. sativa*, GPN’s missense-versus-synonymous AUROC drops from 0.611 in *A. thaliana* to a near-random 0.503 (Figure 9c), while its correlation with PhyloP falls from 0.245 to 0.104 (Figure 9f). Large drops for both measures are also observed with *C. arietinum*, although still giving better than random performance and surpassing those of AgroNT (Figure 9b,e). Botanic1 on the other hand, retains the same ordering with respect to model scale and remains the strongest model for both species with only a small performance drop on chickpea (Figure 9g,h). Investigating the large drop in the missense vs synonymous prediction in rice, we also measure the ratio against the MAF, and find that the enrichment differs substantially between the three species, indicating different evolutionary pressures and links to variant impact (Supplementary Figure S36b).

We observe a similar species dependence when we probe different layer depths of each model. Instead of using the LLR score, we compute embedding distances between ref and alt alleles at different depths: the signal peaks at intermediate depth in *A. thaliana* but at the output layer in *O. sativa* (Supplementary Figure S37). Thus, the depth that contains most allele-frequency information does not appear to be a fixed property of the architecture. Taken together, these results show that with sufficient model capacity and careful pre-training, broad pretraining species coverage does not exclude models from performing well on zero-shot tasks, even compared to specialist models.

##### 2.5.1.2 Zero-shot causal variant prioritisation

To further test the zero-shot capabilities of our Botanic1 models, we construct a dataset of 545 loci across 14 species, each centred on an experimentally validated causal variant (see Section 4.10 for the evaluation protocol and Section 4.11 for the dataset construction). Contrary to the previous analysis, here we are not using a proxy for functional impact: all positive labels correspond to a published study and a proven effect on some aspect of the organism’s biology. Our results show that gLMs are able to rank the documented causal variant in the top 1% or 0.1% of all SNPs in the locus at a higher rate than the bioinformatics baselines, with Botanic1 models occupying the top positions, and that performance increases monotonically with model size and plateaus when the context length reaches 4 kbp (Figure 10).

**Figure 10.**
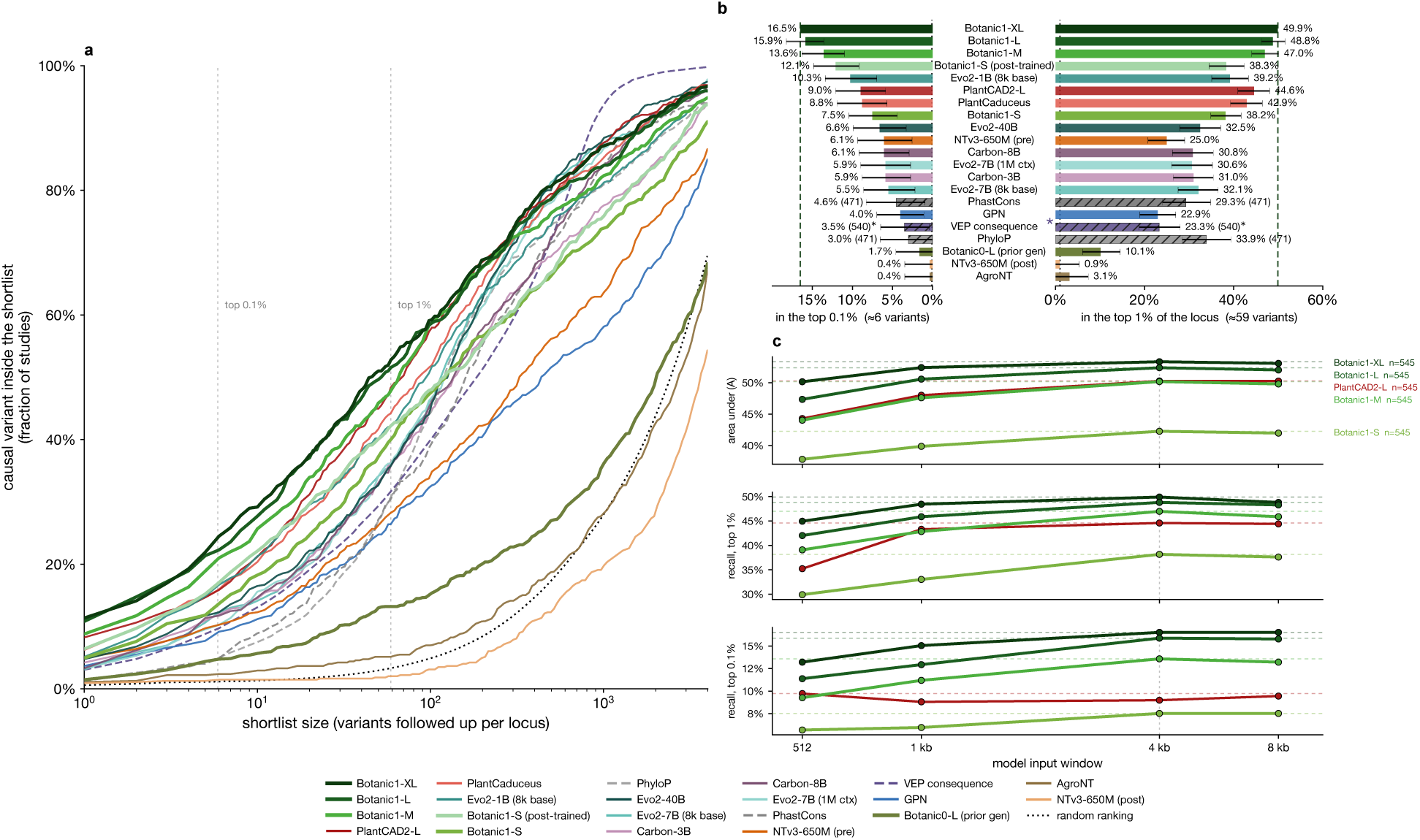
Zero-shot recovery of experimentally validated causal variants. 545 studies, one per distinct causal variant, each scored against every documented SNP in a 100 kbp region centred around the causal variant (median 5,904 candidates). All gLMs are scored on the same 545 studies with a 4,096 bp window, except PlantCaduceus-l32 and GPN, only trained on up to 512 bp windows and scored with 512 bp context. **a**, Fraction of studies whose causal variant falls inside a shortlist of the given size. Recalls are tie-aware (Section 4.10.5). Non-gLM baselines are added with dashed lines and the random baseline (shuffled ranking) as a dotted line. **b**, Recall at top 0.1% (growing left), and at top 1% (growing right). Rows are ordered by the 0.1% column. Full results including paired confidence intervals can be found in (Table 3). Hatched bars are the non-gLM baselines, with the number of studies shown in parentheses next to the score in case of partial support. The asterisk on Ensembl VEP consequence signals the fact that this score is categorical, so most of the recall shown comes from its tied block rather than from an ordering (see main text for explanations). **c**, Context length sweep for Botanic1 models and PlantCAD2-L. Dashed coloured lines correspond to the maximum metric attained by each model.

**Table 3:**
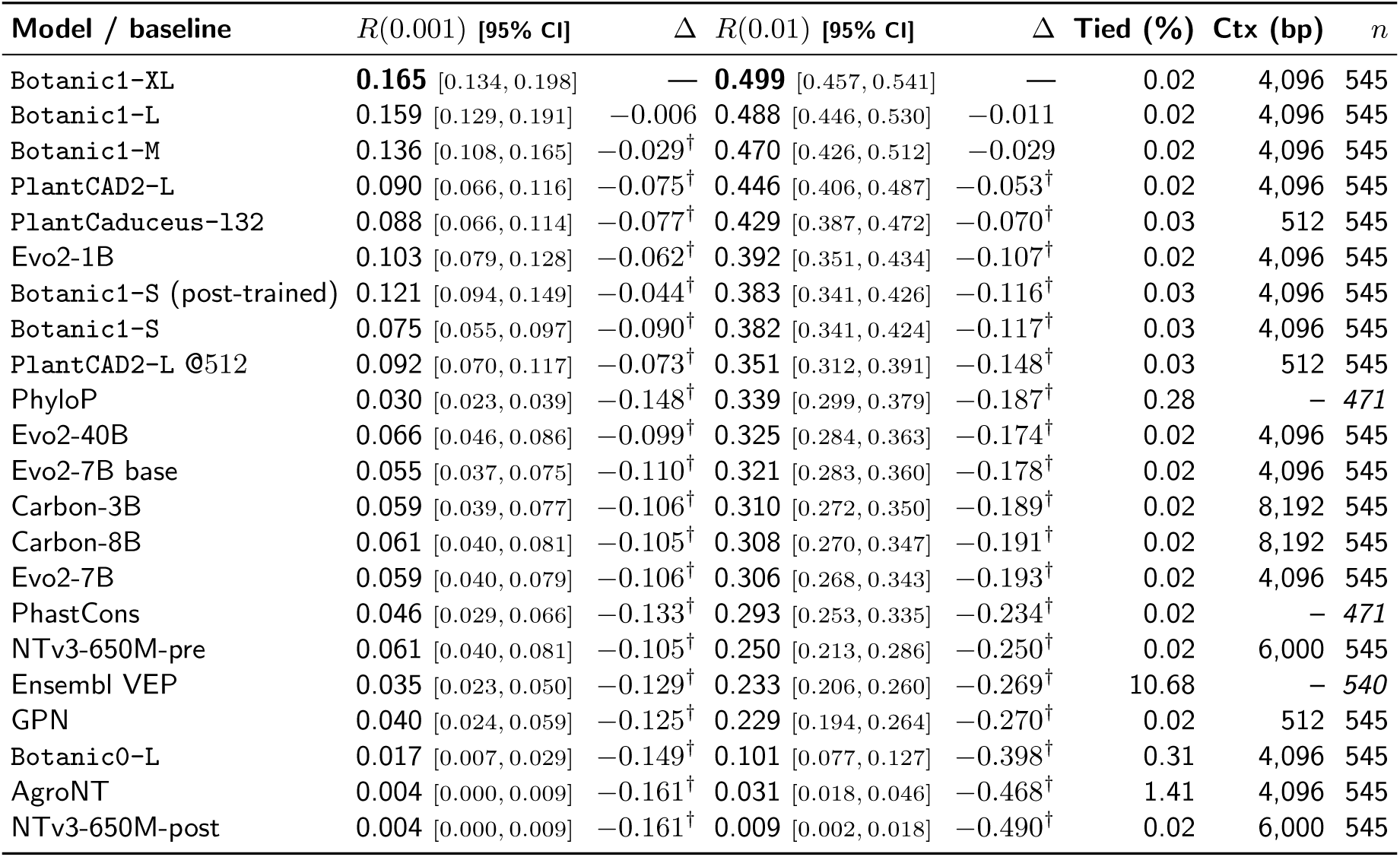
Zero-shot causal-variant prioritisation. This table reports the percentage of studies that have the causal SNP (Recall) at different thresholds for each model’s score. R(0.001) corresponds to the recall at the top 0.1% of all scored variants in the 100 kbp locus centred around the documented causal SNP. On average, R(0.001) corresponds to a shortlist of about 6 SNPs, and R(0.01) of about 60. Each row corresponds to a different scoring method and rows are sorted by the R(0.01) value. Column “Tied” gives the median percentage of candidates that are scored exactly the same as the causal variant across studies. The recalls are tie-aware: if the candidate is within a tie block in a study and the considered top fraction has to cut within the block, then the contribution of that study to the recall is neither 0 nor 1 but the proportion of the tie block that can fit in the shortlist (see Section 4.10.5). 95% Confidence intervals on recall scores are computed using bootstrap over compatible studies. Δ is the paired difference against Botanic1-XL at the endpoint to its left, with a dagger where its own 95% interval excludes zero. PhyloP, PhastCons and Ensembl VEP are paired on their respective intersection set with Botanic1-XL (471 studies and 540 studies).

All gLMs are evaluated on the full dataset of 545 studies from the reference genomes sequences only, while bioinformatics baselines require either precise annotations for Ensembl VEP or multiple sequence alignment (MSA) support for conservation scores. As a result, PhyloP and PhastCons, which are built on PlantRegMap, are only scored on a subset of 471 studies. Since the focus of this paper is comparing Botanic1 models to other gLMs, both Table 3 and Figure 10 report results with maximum coverage across methods, see Supplementary Table S14 and Section 4.10.5 for results on the strict intersection of all methods’ supported studies. Every language model is scored with min(4,096, trained context) bp context ( PlantCaduceus-l32 and GPN are fixed at 512 bp) for each LLR, putting the candidate SNP in the middle of the window.

In the table, we observe that Botanic1-XL, Botanic1-L and Botanic1-M have the highest scores on both recalls. At R(0.001) followed by an experimental Botanic1-S post-trained to predict *A. thaliana* annotations and the allele distribution of the 1,135 accessions of the 1001 Genomes project (see Section 4.10.3.2), which ranks ahead of Evo 2 and PlantCAD2-L at that threshold; at R(0.01) the same model falls behind PlantCAD2-L, PlantCaduceus-l32 and Evo2-1B and matches the plain Botanic1-S. Paired statistics (Section 4.10.5) are used to test the gaps in Table 3, Botanic1-XL reaching 0.499 at α 0.01 is significantly better than the next best non Botanic1 models, 0.446 for PlantCAD2-L and 0.429 for PlantCaduceus-l32; scoring a difference of 0.053 [ 0.090, 0.018] and 0.070 [ 0.106, 0.035] respectively, both excluding zero. Botanic1-S (post-trained) probes whether supervising on annotations and on observed allele frequencies on a single species allows to increase the zero-shot ranking signal, and whether any of those gain can transfer: post-trained only on *A. thaliana*, it outperforms Botanic1-S on 240 non-Arabidopsis studies at α 0.001 (0.096 against 0.062, paired delta with 95% CI at +0.033 [+0.004, +0.067]), but not PlantCAD2-L (0.129), and not at α 0.01. All models for which we perform a context sweep gradually improve performance until 4,096 bp, and plateau after (Table 4).

**Table 4:** Context length sweep for causal variant prediction. Each row fixes the model, and each column fixes the context length given to the model, with the candidate variant centred in the window. Displayed values are the tie-aware R(0.01). All models and all context lengths are scored on the same 545 studies.

| Model | $n$ | 512 | 1,024 | 4,096 | 8,192 |
| --- | --- | --- | --- | --- | --- |
| Botanic1-XL | 545 | 0.450 | 0.484 | 0.499 | 0.488 |
| Botanic1-L | 545 | 0.420 | 0.459 | 0.488 | 0.483 |
| Botanic1-M | 545 | 0.391 | 0.428 | 0.470 | 0.459 |
| Botanic1-S | 545 | 0.299 | 0.330 | 0.382 | 0.376 |
| PlantCAD2-L | 545 | 0.352 | 0.433 | 0.446 | 0.444 |

In this benchmark, non-gLM baselines fail in somewhat unexpected ways. Ensembl VEP is a categorical score, and usually scores the causal variant as “high impact” along with many other variants in the locus (median of 10.7% of all SNPs), that it cannot order further. At a typical locus a top 1% shortlist holds about a tenth of that group, and picks that tenth arbitrarily. In only 66 of the 540 studies covered by Ensembl VEP is the causal variant’s class assigned to a low enough number of SNPs in the locus to measure the recall at top 1% unambiguously, achieving a value of 0.923, against 0.742 for Botanic1-XL and 0.803 for Botanic1-L on those same studies. At top 0.1% the ordering reverses, 0.417 for Botanic1-L and 0.348 for Botanic1-XL against 0.272. Conservation scores also have a ceiling on their possible recall performance, due to the depth of the original MSA and each score’s formula. PhyloP is the most sensitive to this effect, assigning a tied score to a median number of 39 top variants in 87 studies, reproducing the weakness of VEP to a smaller extent. PhastCons uses contextual information and is more continuous, and does not saturate in practice. This explains the differences between the two scores for their recall at top 0.1%.

#### 2.5.2 Botanic1 is the foundational building block for agentic plant-biology workflows

We next ask whether we can expose the real biological knowledge contained within Botanic1 to a generalist coding agent. We consider a retrospective causal-variant discovery problem using as a starting point a melon sexdetermination study [52], in which a G124R substitution in the ethylene-signalling gene <u>CmEIN3</u> was established through field experimentation as a causal SNP that changes melon flowers from female to hermaphrodite. We provide a lightweight generalist LLM agent, Gemma 4 E4B [53] with the 3,061 candidate variants from the chromosome 2 locus identified by bulk segregant analysis of the two segregating populations performed by Rashid et al. [52] and ask it to rank them without access to the publication nor the internet. Across experiments, the agent and prompt are held fixed and only the tools made available to the agents are changed (Section 4.12), and we measure its ability to rank the causal variant in a shortlist of candidates.

With no bioinformatics nor gLM tools, the agent does not identify the causal variant. It reads the beginning of the file, and Gemma 4 tries to rank variants from text alone, unsuccessfully. No variant is invented and this behaviour is expected as generalist models are not trained to understand the statistical and functional constraints encoded in genomic sequences. Providing access to Gemma 4 to a conventional bioinformatics environment substantially improves its performance. The agent autonomously applies variant-quality filters, reconstructs segregation statistics and annotates candidate consequences using standard genomic resources such as Ensembl VEP that we put at its disposal, thereby reducing the search space to a small set of plausible coding variants. When being prompted naively in this new setup, the Gemma 4-powered agent suggests a variant list containing the causal variant in about 15% of the cases. Despite the variant being present in these cases, the agent does not have sufficient information to distinguish it from the other variants. When being prompted with expert guidance, the causal variant appears more often in the list, in about 35% of the tested runs. 10% of the time, Gemma 4 narrows down the list to two variants that it cannot tell apart, reaching expert bioinformatician performance.

We then replace the conventional bioinformatic tools with Botanic1-L as a directly callable zero-shot prediction model (Section 4.12). For each SNP, the agent queries Botanic1 for its log-likelihood ratio between the alternative and reference alleles conditional on the surrounding genomic sequence, providing a continuous measure of surprise correlated with causality, as evidenced in Section 2.5.1.2. Thanks to this hybridisation, the new agentic system ranks the experimentally validated <u>CmEIN3</u> mutation first across all other candidates.

To quantify robustness to stochasticity, we repeat each setting 20 times using identical prompts following recommended sampling settings (Section 4.12). The agent alone fails to recover the causal variant in the top 20 in every run (Recall@1, Recall@10 and Recall@20 = 0). Conventional bioinformatics tools improve retrieval. Naive use of the tools gives a Recall@1 of 0.00 (0.00–0.16), a Recall@10 of 0.05 (0.01–0.24), and a Recall@20 of 0.15 (0.05–0.36) while tools with expert guidance gives a Recall@1 of 0.15 (0.05–0.36), a Recall@10 of 0.20 (0.08–0.42), and a Recall@20 of 0.35 (0.18–0.57)

By contrast, giving the agent access to Botanic1 gives a Recall@1, Recall@10 and Recall@20 of 1. No fabricated variants (see definition in Section 4.12) are observed for either the tool-free or Botanic1 configurations, whereas fabricated variants occur in three and four of the 20 repetitions of the naive and expert-guided conventional bioinformatics workflows (Supplementary Table S15). These rates are measured over repetitions on a single locus, so they quantify how reliably the agent recovers this variant rather than a general recovery rate; the enrichment of experimentally validated causal variants by Botanic1 over 545 studies is reported in Section 2.5.1.2.

## 3 Discussion

In this work we introduce our agentic gLM *Model Factory* as well as its first output: the Botanic1 model family, a new set of plant-specialist gLMs based on bidirectional Mamba2 backbones.

This family demonstrates a strong and balanced profile as plant DNA sequence encoders, with state-of-the-art performance on most of the frozen-model evaluation tasks from both the PGB and the PlantCAD2 dataset suite. Even the 318M-parameter model Botanic1-S outperforms or ties all generalist and plant-specialist gLMs we test, many of which are much slower and larger. In addition to being strong sequence encoders, Botanic1 models can be fine-tuned on more difficult tasks where adaptation is needed. Botanic1 models surpass the specialised from-scratch baselines DenseNet [63] on terminator strength prediction and ChromBPNet [37] on ATAC-seq prediction. To our knowledge, Botanic1 models are also the first plant-specialist gLMs pre-trained to a 131,072 bp (128 kbp) context, twice the 64 kbp context recently reported by PlantGFM [82]. Using synthetic benchmarks, we show that Botanic1 models capture long-range interactions up to 128 kbp context. Finally, the benefit of longer context is illustrated on a real biological task: using appropriate pooling mechanisms that limit signal dilution, we demonstrate improved performance with context size up to 32 kbp for a chromatin accessibility task.

Plant gLM research is still in its infancy and AI plant researchers often fail to perform evaluations as comprehensive as in other areas of machine learning due to limited computational capabilities. We hope that our systematic evaluation of a wide array of both plant-specialist and generalist gLMs on 34 per-task-and-species metrics, 22 of which enter *S*_bal_, sheds more light on the progress and the current state of plant gLM research. Nevertheless, and although benchmarks are a powerful tool to perform comparisons across models on common grounds and to establish where the current state of the research stands in terms of performance, we find that current plant gLM benchmarks suffer from three major shortcomings. First, although gene regulation mechanisms largely rely on large-scale and long-context interactions, and models strive to extend to still larger context windows, benchmarks are often limited to short-window tasks. Second, most tasks rely on a very limited set of plant species, typically the most studied and the best-annotated ones, thus limiting their ability to really test whether models generalise across species. Finally, tasks included in benchmarks often bear little practical relevance, and to assess variant effect prediction capabilities, a skill that could arguably be one of the most useful and immediate applications of gLMs, benchmarks typically rely on proxies such as allele frequencies, which reflect selective pressure only indirectly and are confounded by population dynamics.

Long context is a major blind spot of current benchmarks, especially given that gene regulation mechanisms are known to rely on long-range interactions via enhancers and silencers, prompting recent research work to develop models that can accept increasingly longer windows as input. Most benchmark tasks are assessed on sequences shorter than 1 kbp, making statements on long-context performance poorly grounded. Even when training for a specific context target, gLMs are evaluated out of their training distribution on much shorter sequences [33]. We thus expect that if the only goal is to beat benchmarks, short-context specialists such as MarinDNA [26] (limited to 255 bp) should be the best performers. We argue the plant genomic community as a whole should strive towards building benchmarks with much longer-context tasks that carry biological meaning, in order to recognise gLMs if and only if they can capture and articulate both short and long interactions, not only in theory but also as demonstrated by reliable benchmark numbers.

In addition, benchmarks suffer from limited species coverage, heavily biased towards species with numerous available annotations such as *Arabidopsis thaliana*. This is probably one of the reasons why GPN [83], trained only on Brassicales, is such a strong performer (although this does not explain its retained performance outside of this clade). This limitation prevents us from assessing how well gLMs generalise across species, which is why we are working on extending our benchmark tasks to more species. We expect this to make Botanic1’s margin over competing models even more marked as it has the broadest pre-training plant species coverage out of all the baselines we study.

For a gLM, strong performance on standard benchmarks is not sufficient to establish practical utility, as many genomics benchmark tasks are only tangentially relevant to industry use. Our newly introduced causal-variant benchmark addresses part of this gap by using experimentally validated functional variants as a source of ground truth, instead of relying on indirect proxies such as allele frequency or variant class. Still, this new benchmark also uses some simplifications: the search is limited to a 100 kbp causal region and to a single basic type of mutation, SNPs. Besides, the task relies on the assumption that the documented causal SNP should be ranked among the candidate variants assigned the lowest likelihood by the model, an assumption that could be challenged since the surrounding unlabelled variants may include other functional or phenotypically relevant mutations. Nevertheless, although it should not necessarily be ranked first, we expect a validated causal variant to rank near the top of its locus, within the top 1% ( 60 variants) or even 0.1% ( 6 variants), which is why we choose to assess the performance on this task using the recall at these thresholds. This limitation affects the interpretation of the absolute benchmark scores, but not the relative comparison between methods, since all methods are evaluated against the same set of unlabelled variants.

The causal-variant benchmark illustrates advantages of gLMs that are not directly measured by the current evaluation suite. First, these models only require a sequence from the reference genome around the variant that one wants to score. Neither annotations nor multiple sequence alignments are needed. Second, because Botanic1 models are pre-trained on reference genomes only, their scores on any given alternate variant are not subject to linkage disequilibrium or influenced by the variant’s frequency in arbitrary populations, making them an ideal complement to association scores. Ideally, a variant-effect prediction method should also be interpretable: this is where integrating scores like the LLR with interpretability features or fine-tuned models to predict molecular phenotypes like the base-pair-level chromatin accessibility holds promise, especially if orchestrated by a capable tool-calling agent. We validate that Botanic1 models lend themselves to interpretability research: ATAC-seq performance reveals a reliance of the fine-tuned model on known regulatory motifs such as G-box. More broadly, their layers encode biology: SAEs let us open the black box of our models and shows to what extent known biological features are encoded in the weights, enabling zero-shot annotation capabilities of Botanic1 across many annotation types. Botanic1 models also integrate into complex research workflows, which are increasingly operated by LLM agents. Equipped with Botanic1 models’ skills, a local and privacy-preserving generalist Gemma 4 [53] agent is able to answer open research questions end-to-end, as demonstrated in the melon example. Botanic1 is the missing piece enabling a coding agent to tackle real genomics-based research questions.

While much is known about optimal data mixtures for LLM pre-training [54, 55, 22, 56, 23, 57, 58], the same cannot be said for gLM pre-training. We open-source the resulting pre-training corpus, the data mixture that was used to train the four Botanic1 models, at https://huggingface.co/datasets/living-models/Botanic1-pretraining, so that the community can build upon our work. We dedicated significant resources trying to improve the pre-training data pipeline, and find actionable design choices at three levels of processing: *intraspecies*, *inter-species*, and *inter-window*. Some confirm existing literature wisdom, like the functional biasing of window sampling, while others like the inter-window deduplication filter are introduced as new and valuable steps. Due to the low sample size (two seeds for each experiment), the fixed model scale and architecture, and the caveats described above regarding the evaluation suite, these conclusions should not be considered final. Nevertheless, the data factory experiments highlight the importance of evaluation design when trying to optimise for the pretraining dataset: conclusions may differ depending on whether one is using the model’s loss or downstream task performance, and apparent generalisation to “held-out” species may be confounded by the presence of closely related species in the training set. Future work is also needed to understand the effect of including other kingdoms like animals or prokaryotes (as done by models like NTv3 and Evo 2), but our results suggest that plant-specific pre-training is both more efficient and achieves a higher ceiling for plant-specific downstream tasks. Pushing the comparison between LLM and gLM further would require us to also “mid-train” [23] and “post-train” [84] our models using supervised loss against annotated data, in addition to our masked language modelling self-supervised pre-training. What those annotations and stages exactly entail for gLMs remains an open question, to which NTv3 [25] contributed its own approach. After masked language modelling pre-training, NTv3 introduces a second stage that adds supervised prediction of 15,889 functional tracks across nine species and 21 gene-structure and regulatory annotation classes across 24 species. Splice donor and acceptor sites are two of those 21 annotation classes with direct supervision on *A. thaliana* as well as on other species, so NTv3, through posttraining, trains on the test set for this task (NTv3 correctly reports uncontaminated benchmark numbers in the original paper). Scoring the pre-train-only checkpoint allows us to pinpoint the effect of post-training: on splicing NTv3-650M-pre reaches a stratified AUPRC of 0.955 (donor) and 0.946 (acceptor) against 0.979 and 0.978 for NTv3-650M-post, so the annotation stage gives only a slight performance boost of +0.02 to +0.03 using direct supervision as performance is near saturation (Supplementary Figure S15d). Botanic1-L, with no annotation supervision, reaches 0.984 and 0.980 on the same cells. However on the two zero-shot families, which score variants from the model’s own likelihood, the post checkpoint score falls sharply from 0.692 to 0.438 in mean LLR AUROC and from 0.725 to 0.496 in causal variant discovery recall AUC. This shows that supervised posttraining of this type is detrimental to evolution tasks closer to pre-training supervision. Therefore it seems that plant gLMs have not found a suitable post-training scheme that allows to both gain from annotation supervision without catastrophically forgetting evolutionary pressure learned in pre-training. However, annotations are scarce for plants, and studying variations of similar methodologies on our models would mean either shrinking the species coverage usable for subsequent training steps or generating more data to maintain the 320-species coverage used for pre-training. Unlike for LLMs, none of these stages involves reinforcement learning, since in-silico verifiable reward environments for plant genomics have yet to be built: today, collecting one reward for such a post-training would require greenhouse experiments or field trials, all of them subject to the constraints of the human world and of biology, in particular on speed, measurement noise, and effort. The results we report show that careful pre-training alone can already produce strong models usable for real genomics research, and while integrating subsequent training stages remains an open research avenue, our *Model Factory* provides the scaffolding to enable such research as the relevant data become available.

## 4 Methods

### 4.1 Pre-training

The four Botanic models share the same backbone based on the repetition of Mamba2 blocks [85] and the AdamW optimiser [86], and differ mainly in width, depth and batch size; the number of cumulative tokens and the learning rate schedule are fixed. Supplementary Table S19 gives the exact configuration details for all Botanic1 models.

#### 4.1.1 Architecture

##### Why a state-space backbone

###### Insights from the literature

It might seem counterintuitive not to use a Transformer architecture [87], which has well-tested scaling properties, especially for encoder-only models such as BERT [88]. However, genomics has different inductive biases from regular text and remains difficult to understand, which motivates our data-driven approach. One such inductive bias is locality: certain motifs, such as TATA boxes, are contiguous and short. This may also explain the strong performance of convolutional neural network (CNN) baselines trained from scratch [64, 63]. Indeed, known motifs represented as position weight matrices (PWMs) are regularly used to initialise convolutional filters when training CNNs on genomic data [64, 89]. Mamba2 architectures [85] are also well tested and have good scaling properties despite being used less often than Transformers. They contain small sequence-convolution kernels, making them suitable candidate architectures for gLMs. The work of Evo 1 [90, §2.3, Fig. 1G] showed, by analyzing more than 300 models spanning four architectures, that a strong transformer baseline, Transformer++, was significantly worse in perplexity at every compute budget than equivalent statespace and hybrid architectures. More precisely in their work, Hyena and StripedHyena scale best, ahead of Mamba: at the biggest budget they consider (8 10^19^ FLOPs), the compute-optimal perplexity is about 3.30 for Transformer++, 3.23 for Mamba and 3.13 for Hyena and StripedHyena. Interpolating each architecture’s curve with a slope gives roughly 0.16 of perplexity drop per decade of compute for Transformer++ and 0.18 to 0.20 for the three other architectures; at the biggest reported budget, Transformer++ sits at about 0.07 perplexity above Mamba and 0.17 above Hyena and StripedHyena. According to the Transformer++ line fit, this translates to Transformer++ needing roughly 3× the compute to reach Mamba’s perplexity at that budget (and roughly 10× for StripedHyena’s). a state-space backbone therefore already provides substantial efficiency gain over a well-tuned Transformer on genomics data. a second line of evidence is the strong performance of PlantCAD2-L (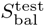 = 0.752) [33] and PlantCaduceus (0.751, Figure 3) [34], which are both Mamba-family encoders.

Reaching StripedHyena 2’s optimal throughput requires Savanna, a pre-training stack of custom kernels and a three-dimensional data-tensor-context parallelism mesh built specifically for multi-hybrid models [91, 92]. Adopting it would therefore require maintaining a second training system alongside our existing codebase, so we decide against it. In addition, performing Evo 2 inference, which uses a StripedHyena 2 architecture trained with Savanna, requires Hopper GPUs, whereas our training cluster contains only Blackwell and Ada GPU architectures. In contrast, Mamba2 [85] has context-scaling properties similar to those of Hyena [93] and provides kernels that integrate readily into a standard PyTorch stack. We therefore obtain improved scaling efficiency on genomic data at modest implementation cost. Contrary to the recent trend in Evo 2 and Carbon [24, 94], we use a bidirectional backbone. Although autoregressive models are studied more extensively because of their many applications in generative artificial intelligence, encoder-only models appear more data-efficient for classification tasks using frozen embeddings, as recent studies exemplify [95, 96].

###### Replicating findings from the literature on our problem

We also test whether the findings from the literature replicate for encoder-only models and our plant-specific tasks. a 352M-parameter encoder-only Transformer (rotary position embeddings, RMS normalisation, query-key normalisation) is pre-trained using the same general pre-training scheme: same pre-training data, tokeniser, 8,192-token windows, peak learning rate and schedule shape as Botanic1-S (318M), on a 500-B-token horizon. On 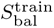 it needs about six times more tokens than Botanic1-S to reach its maximum, about eight times more at 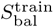 = 0.70, and its 500B endpoint stays below the Botanic1-S checkpoint’s value at 314.6B tokens (Supplementary Figure S9). On a per-task basis it trails in eight of the nine *S*_bal_ components and ties on causal variant discovery (subset) (Supplementary Figure S10). The comparison is robust: fourteen other BiMamba2 runs at similar sizes using different hyperparameters outclass a 1.0B Transformer trained for up to 105B tokens, as well as two other Transformers. The pair has similar training loss at matched token counts, so training loss is insufficient to distinguish the two architectures.

Every model is a stack of BiMamba2 blocks. a block instantiates two Mamba2 modules, one per reading direction. Inputs are tokenised one token per nucleotide over a vocabulary of nine entries: the four bases, N, and the <unk>, <pad>, <mask> and <cls> special tokens. Embedding and output head are untied. All four models are pre-trained on sequences of exactly 8,192 tokens, a <cls> token followed by 8,191 bases.

#### 4.1.2 Training objective

Pre-training uses single-position MLM. Each eligible position (all positions except special tokens) in the sequence is selected independently with probability 0.15. a selected position is then replaced by <mask> with probability 0.8, replaced by a token drawn uniformly from the four nucleotide tokens or N with probability 0.1, or left unchanged with probability 0.1. Special tokens and N tokens are excluded from both masking and scoring. For all other positions that are not selected for masking, the target label is set to 100 so that position does not contribute to the loss. The resulting loss is the mean cross-entropy between predicted and ground-truth tokens over selected positions, weighted per position to down-weight soft-masked repeats as in Evo 2 [24] (see Section 4.2): a position of repeat fraction *f* receives the weight 1 (1 w)f with *w* = 0.1, so a repeat base contributes a tenth of what a normal base contributes.

#### 4.1.3 Optimisation

We use AdamW (β_1_ = 0.9, β_2_ = 0.95, ɛ = 10*^−^*^8^) with weight decay 0.05 and gradients clipped at a global L_2_ norm of 1.0; per-block gradient norms are additionally logged for diagnostics. The learning rate follows a warmup-stable-decay (WSD) [97] schedule shared by all four sizes: a warmup over the first 3.93B tokens to a peak of 10*^−^*^4^, a constant phase, then a decay over the last 10% of the 314.6B-token horizon down to a tenth of the peak. Training runs are performed in bf16 under fully sharded data parallelism with a compiled model graph. Batch size is set so that Botanic1-S, Botanic1-M and Botanic1-L see the same 192 sequences, or 1.573M tokens, per step, and reach the 314.6B-token horizon in 200,000 steps. As in [32], we define tokens seen as token tensors transferred to the GPU and processed by the model, without correcting for the fact that only 15% of their positions contribute to the loss (see Section 4.1). Botanic1-XL doubles the tokens per step to 3.146M and reaches the same horizon in 100,000 steps, so its warmup (1,250 steps against 2,500 for the other three) and decay window match the others in tokens. All four reported checkpoints are taken at the end of the horizon, fully decayed.

#### 4.1.4 Compute

All four models are pre-trained on nodes of eight NVIDIA B200 GPUs, on one node for Botanic1-S, two nodes for Botanic1-M and Botanic1-L, and four nodes for Botanic1-XL, one data-parallel replicate per node (Supplementary Table S19). Jobs run through SLURM and resumed from their latest checkpoint when preempted. The training time we report is the wall-clock time spent training towards the reported checkpoint, measured from the per-step throughput logged every ten steps, excluding queue time and the segments discarded when a job resumes from an earlier checkpoint. Botanic1-S trains in 77 hours (616 GPU-hours), Botanic1-M in 83 hours (1,334 GPU-hours), Botanic1-L in 202 hours (3,228 GPU-hours) and Botanic1-XL in 128 hours (4,103 GPU-hours), so the four released models together cost about 9,300 B200 GPU-hours, about 48 node-days on our training cluster. This excludes the data ablations, the architecture comparisons, the context-extension and evaluation.

Model FLOPs utilisation (MFU) [98] is the training FLOP rate divided by the 2.25 PFLOP/s dense BF16 peak specification of one B200. The FLOP count per token uses the 6N rule for the parameter matrix multiplications, with N the non-embedding parameter count, plus the state-space scan, which is not included in the 6N. Per token, per layer and per scan direction, the chunked SSD algorithm of Mamba2 [85] costs H [2Q(N*_s_* +P)+4N*_s_*P] FLOP in the forward pass, with H heads of dimension P = 64, state size N*_s_* = 128 and chunk length Q = 128: the first term is the two intra-chunk matrix products account and the second term is the chunk-state and state-to-output products. Counting the backward pass at twice the forward pass and the two scan directions, the training cost is F = 6N + 60 L d_inner_N*_s_* = 6N + 15,360 L d_model_ FLOP per token, where the scan adds 20% to 6N at d_model_ = 1,024 and 12% at 1,792. Using this formula the four models reach 15.3%, 15.5%, 17.9% and 20.7% MFU (Supplementary Table S19); under the bare 6N rule the figures are 12.8%, 12.9%, 15.8% and 18.5%. Utilisation rises with width because the projections of the wider models fill the tensor cores better, and the scan itself, which is memory-bound, weighs less in the total. The single-nucleotide vocabulary makes the embedding and output layers negligible.

#### 4.1.5 Context extension training

The context-extended models of Section 2.2.3 start from a 318M BiMamba2 backbone with the Botanic1-S configuration, pre-trained at 8,192 bp for 500B tokens under the above schedule, decayed to a tenth of the peak learning rate at the end of its horizon. Four extension stages follow, doubling the maximum input window in the training mix at each stage to 16,384, 32,768, 65,536 and finally 131,072 bp. Every stage trains for the same 34,679 steps, with the tokens per step doubling at the 65,536 bp stage and unchanged at 131,072 bp, so the four stages contribute 50, 50, 100 and 100B tokens. Each stage warm-restarts the learning rate and repeats the warmup-stable-decay shape of the base schedule at half its peak: a warmup over the first 2,500 steps to 5 10*^−^*^5^, a constant phase, then a decay over the final 10% of the stage to 5 10*^−^*^6^.

We additionally continue the training of each context-extension stages and the base stage up to the same number of cumulative tokens as the last context extension stage, which isolate the effect of context extension itself from the confounding factor that models from earlier stages would have instead received less tokens in total. More specifically, each of those models (all models from each stage except the one after the last stage) undergo CPT by branching out before the learning-rate decay phase of the ongoing stage and extending it to reach 300B tokens the amount of additional tokens cumulatively by the last context extension. The per-stage data mixtures are described in Section 4.2.8 and the variable-length packing in Section 4.1.6.

#### 4.1.6 Batching and data loading

Our training codebase builds on the code from [32], itself a fork of Meta’s Lingua [99] research pre-training library. We retain Lingua’s general structure: a chain of nested iterators, each yielding a sample together with the random state immediately after its random operations, so that training can restart from the exact next sample. The dataloader state is saved to disk alongside the weights and optimiser state when the job receives a specific signal (SIGUSR2). Lingua’s chunk assignment, weighted source sampling, token packing and prefetch shuffling are mostly unchanged. The earlier code [32] added support for genomic data and masked language modelling. Here we further add efficient variable sequence lengths for context extension, multi-threaded prefetch for larger prefetch sizes or sequence lengths, and case sensitivity. Case sensitivity exposes a metadata vector containing repeat information for each token or base pair, allowing repeats to be down-weighted correctly in the loss. Supplementary Figure S38.

##### Sources and workers (Supplementary Figure S38a)

Each data block of Section 4.2 corresponds to a single source and is stored as K*_s_* pre-shuffled Zstandard-compressed JSONL chunks (see Methods in [32]). Chunks are distributed to the data-parallel workers, and workers sharing a chunk read it at interleaved line offsets, so every worker consumes a disjoint stream and no line is seen twice in a pass. Each worker draws samples from a source using fixed weights, then consumes the next line of that source. The four Botanic1 training runs in Supplementary Table S19 draw from a single ≈ 8 kbp source detailed in Section 4.2.1; data mixtures are used only during context extension (Supplementary Figure S42), where up to five data sources of different window lengths are sampled per stage.

##### Per-sequence transforms (Supplementary Figure Figure S38b)

Each JSONL line holds one stored genomic window plus its margin (Section 4.2). The loader draws a start offset uniformly at random following [31, 32], far enough from the end of the line to leave enough space for the target number of bases after it. It reads the lower-case pattern of that text into a binary repeat mask before upper-casing destroys the soft-mask signal, and maps any remaining ambiguity code to N. With probability 0.5, the sequence is reverse-complemented during training and its repeat mask is reversed alongside its tokenisation. Although all models trained here use single-nucleotide tokenisation (SNT), our codebase supports tokenisers that aggregate repeat annotations across the bases of each token and produce a fractional repeat score. With SNT, we map bases to a precomputed set of identifiers and prepend a <cls> token following [88].

##### Packing (Supplementary Figure

The strategy used to pack tokenised sequences into rows is configurable. The four base runs presented here use the padded strategy, where the tokeniser trims each sequence to the row length, and the packing strategy places one sequence per row. In this case, no padding is required, as all sequences have the same length due to the use of the single-base tokeniser. When sequence lengths differ during context extension, the packed flat packing strategy instead concatenates sequences back-to-back into a single flat chunk of mbs tokens tokens, (90,112 at the 16 kbp stage, 180,224 at 128 kbp), following the same approach that Lingua uses to pack a text stream. a sequence that extends beyond the end of a chunk is split, with its remaining portions placed at the beginning of the next chunk, ensuring that no tokens are ever dropped. In either case, the cumulative sequence lengths and the length of the longest sequence in the batch are emitted alongside the batch, and the backbone uses them to keep each sequence independent of its neighbours through the use of varlen kernels.

##### Batches (Supplementary Figure S38d)

Rows are masked at the rates of Section 4.1.2 as they enter a prefetch buffer that holds in our experiments 32 batches. Once the buffer is full it is shuffled, and batches are emitted as slices of the buffer, so that neighbours in a batch are not neighbours in packing order; this decorrelates the long sequences that packing tends to group together. Each consumed slice is refilled immediately and the buffer is permuted again on every full cycle, a blockwise shuffle buffer (see [100]). Lingua builds its targets by stacking copies of the row shifted by one token, the “views” of next-token prediction; masked language modelling needs a single view, so we set that count to one and the second channel of the emitted array carries the labels instead. Alongside the batch the loader returns the per-position repeat fraction that will be used to weigh the loss, the cumulative sequence lengths, the longest sequence in the batch, and its own state (random state and indexing). Supplementary Figure S39 displays the evolution of the masked-language-modelling training loss (Supplementary Figure S39a) as well as the evolution of the species-balanced downstream score 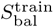 (Supplementary Figure S39b) for the four model sizes against the number of tokens seen. a star marks each model’s checkpoint at the common 314.6B-token horizons. The corresponding scaling results are reported in Section 2.2.2.

### 4.2 Genomic data sourcing and preprocessing

A model factory is only as strong as the data it trains on, and data selection research for genomic language model training is a relatively new and under-explored field at least compared to the immense body of work considering traditional LLMs. While recent efforts have somewhat converged on a few key preprocessing steps, we make the conscious effort to explore different ways to prepare the pre-training data, drawing inspiration from both LLM pre-training and bioinformatics best practices.

When building a pre-training corpus, the objective should be to maximise signal and minimise noise and redundancy. Concretely, the corpus is a set of windows: fixed-length contiguous stretches of genomic sequence, each tokenised into a single training example. Broadly, there are three major levels at which signal and redundancy can be controlled: *intra-species*, *inter-species*, and *inter-window*.

Our previous effort, Botanic0, reproduced AgroNT’s [31] genome-wide uniform sampling and did not apply any of those controls, resulting in particular in the large majority of the windows being sampled on intergenic regions full of repeated nucleotide patterns leading in turn to poor performance and training efficiency. converging on sampling preferentially for functional sequence: GPN [83] enriches for genic and promoter windows, PlantCAD2 [33] trains on gene-centred windows, and Evo 2 [24] and the Nucleotide Transformer v3 [25] weight short-context pre-training towards gene-proximal, functional regions; GPN, PlantCAD2, and Evo 2 further down-weight repetitive sequence in the loss. The concurrent MarinDNA [26] takes this to its limit in mammals, drawing every training window from a mixture of five functional region classes (coding sequence, upstream, downstream, ncRNA and enhancer) and centring each window on an individual element. It does so at a context of only 255 bp, shorter than any other model in this list and shorter than the genes, promoters and accessible regions one would want to localise in a newly assembled genome. The other two levels (*inter-species*, and *inter-window* ) remain comparatively unexplored. Interspecies control (i.e. how the token pool is divided across species) is the genomic counterpart of the data-mixture problem in language model pre-training, where data mixtures are often reweighted by domain, or by language in the multilingual case, rather than sampled in proportion to raw volume [101, 102, 103, 104, 105, 106]. Here we can build on the large body of work on phylogenetic distances, from classical tree-based sequence weighting [107] to phylogeny-aware model training [108]. Inter-window control is complementary to the first two levels: genes and functional regions are near-atomic units that can be highly conserved, even across highly divergent genomes (e.g. orthologs [109]). Finally, a corpus balanced across species and enriched for function can still include many near-identical copies of the same locus, which the model can easily memorise during training. This is the genomic counterpart of deduplication in LLM pre-training, where removing exact and near-duplicate documents have been shown to improve accuracy per training step while sharply reducing verbatim memorisation [110], memorisation grows log-linearly with the number of times an example is duplicated [111], and repeated data disproportionately damages the internal structures associated with generalisation [112]. Embedding-space deduplication [113] is the closest analogue to ortholog detection here, since near-identical loci are related by descent rather than exact string identity.

We design the data pipeline to be as modular as possible so that we can ablate precisely the influence of the different controls presented above as well as more easily allow the addition of potential other data sources for the next generations of our models in the spirit of the model factory. The main steps of our data pipeline are thus (Figure 2a): filtering assemblies on quality and annotation, controlling redundancy between species, preferentially sampling windows within species based on functional annotations, and down-weighting repetitive sequence in the loss.

#### 4.2.1 Source catalogues and assembly quality control

The initial assembly catalogue is sourced from Ensembl Plants release 62 and the NCBI Assembly resource [114, 115]. From Ensembl, we retain species present in Ensembl’s peptide-level comparative-genomics analyses and download the soft-masked, top-level genome FASTA and its GTF annotation. For NCBI accessions, we retain only complete assemblies at scaffold level or above. We keep the latest major release we prefer the RefSeq GCF assembly over its paired GenBank GCA assembly, and we exclude assemblies flagged as partial or contig-level. When the two catalogues contain the same species, we choose the Ensembl record.

Some quality controls are applied on assemblies before window sampling. First, we retain only embryophytes (land plants) and remove the non-embryophyte algae present in the source catalogues (Five species spanning green and red algae). For the 8 kbp artefact, a species remains eligible when the highest contig N50 (defined as the largest contig length such that half the assembly lies in contigs that long or longer) among its candidate assemblies is at least 50 kbp, and the smallest assembly length is lower than 10 Gbp. The longer-context artefacts raise the contig-N50 threshold to roughly eight times the window length: 130 kbp, 260 kbp, 520 kbp, and 1.05 Mbp for the 16, 32, 64, and 128 kbp windows respectively, so that a window fits comfortably within a single contig. Contig N50 is computed directly from the FASTA files for the 118 Ensembl assemblies whose source metadata is missing. We exclude annotation files containing fewer than 3,000 or more than 300,000 gene records, as well as files containing no exon or CDS records. Assemblies failing these checks are treated as unannotated; after sequence-similarity pruning, species without any valid annotated assembly are removed. For the 128 kbp data mix, only species with at least one assembly recorded as long-read are retained.

We provide the resulting full assembly list as supplementary material in Section 4.2.6.

#### 4.2.2 Species redundancy and representative assemblies (inter-species control)

To control the inter-species redundancy, we compute a whole-genome distance matrix with sourmash [116]. Every available assembly is sketched, with lowercase soft-masked bases converted to N, so that annotated repeats do not contribute k-mers. Sourmash sketching is done using *k* = 31 and a scaled value of 5,000. The pipeline estimates assembly-level average nucleotide identity (ANI) from maximum containment and defines distance as 1 ANI. For two species represented by multiple assemblies, their inter-species distance is the 10th percentile of all pairwise distances between assemblies of the two species.

These parameters are chosen by a grid search that maximises the agreement between the resulting tree topology and the TimeTree reference phylogeny [117], so that the distances track known divergence times while being computed directly from the assemblies used to build the pre-training corpus.

We use the resulting inter-species distance matrix in several ways. First, we remove disconnected distance outliers. a single-linkage threshold is swept until the largest connected component contains at least 99.75% of species, and all species outside that component are discarded. This removes five of the 2,023 species that pass the preceding quality filters and occur in the distance matrix: *Aquilegia reuteri*, *Isoetes taiwanensis*, *Ceratopteris richardii*, *Adiantum capillus-veneris*, and *Acrolejeunea sandvicensis*. The first corresponds to an erroneous metagenome bin, at maximal distance from every other species, the remaining four are isolated early-diverging lineages: a lycophyte, two ferns and a liverwort. We then group the remaining 2,018 species by complete linkage at a low ANI-distance threshold of 0.05. The resulting 1,344 clusters include distinct but closely related species; and we select one candidate species per cluster in order to avoid giving more importance to densely sampled lineages. Instead of hard-sampling a single species per cluster, we also explore using GSC sampling weights [60] derived from the pairwise distance matrix, but we find that this does not improve the downstream score and it is disabled for the sake of simplicity.

To select which species are included in each group, we design four priority tiers. We first prefer species having both reference status (defined as possessing an NCBI reference assembly or originating from Ensembl Plants release 62) and at least one valid annotation (Tier 1). We then prefer species with reference status (even without a valid annotation) (Tier 2), followed by annotated non-reference species (Tier 3) and, finally, species having neither property (Tier 4). Only two clusters have reference-first ranking select an unannotated “reference” species over an annotated alternative. Among the closely related highest tier species, ties are broken using contig N50.

After this clustering and selection step, we now have one assembly/genome per species, so we can use the terms interchangeably. For all artefacts using functional-biased window sampling, we then remove all species without a valid annotation, leaving 326 genomes for the 8 kbp corpus. The pipeline can relax this requirement for repeat-density sampling, which does not require annotations. Longer-context artefacts have more demanding contig-N50 thresholds or other quality controls and thus contain fewer species. Train and test splits are performed at the species level, and 3 out of 4 test species are the same across all context lengths (the 4th species being removed from long context test due to its low contig N50).

Supplementary Figure S40 summarises the phylogenetic composition of the training set on a dated tree. Although the corpus spans 48 orders and 102 families, its coverage is strongly concentrated within angiosperms: 310 of the 320 species belong to the angiosperm crown group, with 241 eudicots and 61 monocots. Earlier-diverging land-plant lineages are sparsely represented, with only ten species across liverworts, mosses, lycophytes, and gymnosperms. This imbalance is driven primarily by the availability and quality of genome annotations. Ferns provide an extreme example: although seven of the 18 fern genomes in the source catalogue pass the contig-N50 threshold, only two also pass annotation QC, and both are subsequently removed as outliers, leaving no ferns in the final corpus. Finally, species abundance does not translate directly into training-data abundance because of functional-balanced window sampling, and genome-length weighting modifies each species’ contribution: for example, Poales represent 11% of species but 14% of training windows, whereas Fabales represent 10% of species and 8.7% of windows.

#### 4.2.3 Functional-balanced window construction (intra-species control)

We now describe the *intra-species* sampling: how windows are drawn from each genome.

First, sequence records identified as mitochondrial, chloroplast, or plastid sequences are excluded. Primary chromosomes are always retained; unplaced or unlocalised scaffolds and contigs are kept only when the annotation describes at least one feature on them, and dropped otherwise. Within each retained record, an *unambiguous interval* is a maximal contiguous run of A, C, G, or T (upper case or lower case), any ambiguity code breaks the interval.

For functional windows, our window sampler constructs a set from all exon and CDS features together with strand-aware regions extending 1 kbp upstream of each transcript. These regions are merged, expanded by half a window length on each side to provide local genomic context (4,096 bp at 8 kbp and 8,192 bp at 16 kbp), and intersected with the *unambiguous intervals*. For intergenic and non-coding windows, the sampler then draws random intervals from the same genome as a background component.

The CDS-anchored window sampling scheme described above assumes that the chosen window is mostly made up of annotated loci. This is usually true for the shorter windows (8 and 16 kbp), but at 32 kbp and beyond, it is much more common to end up sampling sparsely annotated genomic regions due to one isolated gene. For these long-context artefacts we replace the gene anchor strategy with an annotation-free sampling that uses repeat fraction as a proxy for functional content. Each candidate window is scored by rewarding both low repeat (lowercase) fraction and long contiguous stretches of unique (uppercase) sequence. The “functional” set is the top-scoring half of a genome’s candidate windows, and is also complemented by random background windows drawn from the same genome. On annotated genomes, this soft-mask score rises with gene-annotation density (Spearman ρ = +0.63 over 45,780 pooled 128 kbp windows; Supplementary Figure S41).

##### 4.2.3.1 Genome-length weighting

To limit the impact of large genomes on the final training mix, windows are also down-sampled relatively to their species length. With D*_s_* denoting the genome length (taken as the number of non-N bases), we apply the following weights, normalised so that the largest is one and the floor is 0.5:

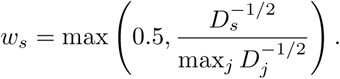

The expected window count therefore scales as √*D_s_* instead of D*_s_* until the floor is reached, penalising species with exceptionally large genomes, such as the ~10 Gbp tetraploid wheat *Triticum turgidum* [118], while no species is down-sampled below half of its global pool (Figure 2a). The smallest genome in the corpus, *Arabidopsis thaliana* (0.12 Gbp), receives a weight of 1; the weight decreases with genome length until it reaches the floor of 0.5 at 0.48 Gbp, after which all larger genomes receive the same weight. Thus, for example, a 0.5 Gbp genome and the 11.9 Gbp genome of *Vicia faba* receive the same correction despite having widely different lengths. Among the 306 of the 320 contributing species whose assembly size is resolvable from the released metadata, 208 sit at the floor, so only the remaining 98 smaller genomes are reweighted differently by this scheme.

As a result, the correction reduces the tendency for larger genomes to contribute more windows but without erasing it completely. Empirically, doubling genome length increases a species’ share of the 8 kbp windows by about 39% on average (D^0.47^), compared with 48% without correction. Species with genomes larger than 2 Gbp therefore still occupy a larger median fraction of the corpus (0.50%) than species below 0.48 Gbp (0.19%), whereas perfectly uniform sampling across these 306 species would assign 0.33% to each.

#### 4.2.4 Window deduplication and species-length weighting (inter-window control)

Finally, we also use window-level and *inter-window* controls.

Windows containing more than 20% N bases are removed. Near-duplicate windows are then filtered *within each selected assembly*, removing near-copies of the same window within species, but not orthologous sequence between species. Two windows are compared using the fraction of k-mers they have in common (Jaccard similarity) estimated with a per-window MinHash sketch (128 smallest hash values, each k-mer is canonicalised with its reverse complement so that strand orientation alone does not make two windows different). a window is discarded when its estimated Jaccard similarity to any window in the current set reaches 0.3. This removes roughly 15–25% of the sampled windows depending on the species pool and recipe; on the full 330-species ablation pool it removes 20.6% (65.6 → 52.1 Gbp). We use *k* = 8 at 8 kbp and *k* = 16 for all longer contexts; the threshold remains fixed at 0.3 through 16 kbp and is lowered to 0.25 at 32 kbp and 0.2 at 64 and 128 kbp. This 0.3 threshold is not comparable to the ≈ 0.9 typically used to remove near-duplicate documents in the LLM literature: with *k* = 8 there are only 4^8^/2 canonical 8-mers, the same order as the number of 8-mers in a single window, so two unrelated windows already share a non-negligible fraction of their sketch which impacts the threshold. This is also why the longer contexts move to *k* = 16.

For each artefact, we store the training window plus a small margin, a 100 bp margin at 8 kbp, widening the margin to 500, 1,000, 2,000, and 3,900 bp for the 16, 32, 64, and 128 kbp windows. Eligible intervals are traversed exhaustively with a 50 bp overlap between successive stored windows as in [31, 32]. During training, a substring of the target length is selected at a uniformly random offset within this margin as data-augmentation [31, 32]. The substring is itself reverse-complemented with probability 0.5 (Section 4.1.6).

#### 4.2.5 Serialisation and metadata

Each resulting window is then serialised as one jsonline object with the sequence in a text field and the species identifier retained in species id. The FASTA letter case is preserved: lowercase marks repeats and is later used for loss down-weighting (Section 4.1). All windows are shuffled and written to eight Zstandard-compressed JSONL training shards (replicating the number of training GPUs in each node). For each final data artefact, a manifest stores the resolved source, filtering, sampling, split, and output settings used; the pipeline revision and working-tree state; the realised per-species count; and a SHA-256 digest to uniquely identify every output file.

#### 4.2.6 Genome assemblies in the pre-training corpus

Supplementary Table S8. lists every species in the champion pre-training corpus (the functional balanced recipe with GSC phylogenetic reweighting disabled and the genome-size correction of Section 4.2 enabled), together with the assembly retained as the species representative and the number of 8,192 bp windows it contributes. The corpus is assembled from 5,074 candidate assemblies by the filters of Section 4.2, leaving 322 training and 4 held-out genomes (one per species); 320 of the training species contribute at least one window, for 7,207,506 windows and 59.0 Gbp of training sequence, the 8,192 bp row of Supplementary Table S20. Window counts are read from the pipeline manifest for that corpus build; species are ordered alphabetically.

#### 4.2.7 Ablation protocol

When performing ablations isolating each control in Section 2.2.1, we use a fixed model size (300M parameters) and token budget, scoring every configuration on the downstream score 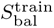. The unique-token pool size is also fixed for each comparison (trimmed down to the lowest common denominator among the compared configurations), effectively fixing the number of trained “epochs”. Each cell is run for two independent seeds, controlling randomness in both the data preprocessing and model training.

##### 4.2.7.1 Window-sampling ratios

The functional-balanced recipe draws functional and background windows in equal proportion (1:1). The functional-heavy variant raises the ratio of functional to background windows to 4:1, both are compared with uniform sampling on each species pool (Figures 2c and 2g).

##### 4.2.7.2 N-content filter sweep

The sweep of Figure 2h varies the maximum fraction of N bases a window may contain: 0 (any window containing an N is discarded), 0.05, 0.20, and no filter at all, on the 48-species phylogenetically diverse pool under functional-balanced sampling. The final pipeline uses a threshold of 0.20.

##### 4.2.7.3 Data specialisation

We test on four additional 300M models how the phylogenetic proximity of the pre-training species pool affects the performance on a target species, here Arabidopsis (Supplementary Figure S3). We follow the two-seed, iso-step and iso-token protocol above, share the same configuration, and change only the species pool: a specialist pool of 48 Brassicales genomes (the clade of Arabidopsis that is used in GPN) and a phylogenetically diverse pool of 48 species selected with the whole-genome distance matrix described above, each with or without Arabidopsis. Each model is scored using two different aggregations of the 28 skill metrics logged at ablation time (this suite predates the 34-metric suite of Section 4.3): the Arabidopsis performance metric averages the seven Arabidopsis metrics (TIS, TTS, genomic region classification, chromatin accessibility, LLR, and splice donor and acceptor), and the other is an average over the remaining 21 metrics.

#### 4.2.8 Context extension data preparation

To expose the model to longer contexts, we create five datasets with window sizes 8,192, 16,384, 32,768, 65,536, and 131,072 bp. Their sizes are powers of two but for simplicity we instead refer to them as 8, 16, 32, 64, and 128 kbp. Those are produced by re-running our data pipeline (Section 4.2) at the target window length and increasing the contig-N50 floor accordingly. This limit rises faster than assembly contiguity improves, causing fewer species to qualify at each step: 326 species at 8 kbp, then 285, 273, 261, and 238 at 16, 32, 64, and 128 kbp (Supplementary Table S20). The 128 kbp dataset requires at least one long-read assembly per species, which is why it has the fewest species. Total base pairs per dataset range between 59 and 116 Gbp: fewer but longer windows offset the shrinking species pool. Window sampling also changes with window length: while the shorter-window (8 and 16 kbp) datasets use coding-region-biased sampling (Section 4.2), longer windows (32, 64, and 128 kbp) almost always contain coding regions, so biasing by repeat content becomes more informative; this also lets more assemblies qualify even without precise genomic region annotations, which are no longer needed.

### 4.3 Building the supervisory signal

A model factory requires an evaluation signal that can be measured repeatedly and compared reliably across training runs. a particularity of gLM evaluation is that such a signal should necessarily span multiple species in order to measure both in-data mix species and performance on previously unseen species. It must also contain sufficient genuine biological signal to distinguish meaningful improvements in the learned representations from gains driven by simple sequence statistics or benchmark-specific artefacts.

To construct this signal, which we call *S*_bal_, we start from the two supervised benchmark suites which are becoming standard in plant genomics: the PGB released with AgroNT [31] and the datasets suite released with PlantCaduceus [34]. From these suites we keep the tasks that are sufficiently fast to measure while still giving strong signal about representation quality. This removes every fine-tuning task, which is too costly at checkpoint frequency, and the datasets whose class imbalance is so extreme that the only applicable subsampled estimator is biased and noisy (Section 4.3.5). Regression targets such as promoter and terminator strength are handled separately similarly as requiring fine-tuning [32]. We therefore leave them out of *S*_bal_ and report them separately under parameter-efficient fine-tuning in Section 2.3.2.1. We then de-duplicate the tasks that recur across the two suites, keeping a single copy of the splice-site capability. Finally as said above, two of the remaining tasks cannot be scored exhaustively: translation initiation and termination on the three large cross-species genomes as they reach 11.3 and 29.5 million candidate positions for a few thousand true sites, so we score them on a class-stratified sample with a prevalence correction on the AUPRC estimation. That estimate is too imprecise to be averaged into the aggregate score, so those metrics are measured but reported in their own section Section 2.3.1.2 (see Section 4.3.5). We add to those benchmarks a series of genomic region classification tasks derived from GPN [83], which measures the annotation discrimination of embeddings by averaging an embedding and the embedding of its reverse complement pooled over a short window (the central 100 bp of a 512 bp context), which we extend to other species than *Arabidopsis thaliana* (more details can be found in either [83] or [32]). Finally, we complement the result with multi-species and zero-shot evaluations, including a new causal variant discovery task described in Section 4.10 as well as an LLR evaluation task described in Section 4.3.2, to test whether improvements generalise across species, genomic contexts, and sources of biological supervision.

We measure over 34 per-(task, species) scalars, of which 22 enter *S*_bal_. Averaging the 22 metrics directly would give artificially more importance to well-represented tasks, in particular because the number of species available per task depends on how the benchmark was created and should be irrelevant: chromatin accessibility alone contributes seven of the 22 tasks, and *A. thaliana* recurs in seven families as this species is an emblematic test species in plants which has been annotated extensively. We therefore report *S*_bal_, which first averages species within each of the nine task families and then averages the nine families, so that every specific biological capability enters with equal weight:

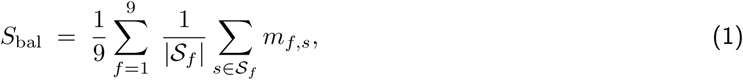

where *_f_* is the set of species evaluated for family *f* and *m_f,s_* the corresponding metric. The nine families, and the species evaluated for each, are listed in Supplementary Table S4. Seven are supervised probes trained on frozen representations; the remaining two are zero-shot and score variants directly from the model’s likelihood.

We use two variants of the same score: S^train^, which we use as a training signal in our *Model Factory*, and 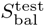, which follows a protocol more aligned with the literature (Section 4.3.6), scoring splicing on centre-token embeddings with the prevalence-corrected stratified AUPRC (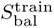 pools the window mean and scores splicing with the prevalence-corrected stratified AUROC on the same stratified draw, Supplementary Table S5) and, for auto-regressive models, performing two forward passes, one on the normal strand and one on its reverse complement, which allows centre-pooling with auto-regressive models without losing or diluting information. Every leaderboard value and model comparison in this article uses 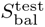, training trajectories, ablations and checkpointfrequency figures plot use 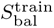. Both aggregate the same tasks and differ only in the exact scoring protocol, so we write *S*_bal_ without superscript when a statement holds for both protocols.

We test whether this, somewhat arbitrary, averaging leads to other conclusions than other equally valid aggregation choices. We re-score every model under alternative aggregations (Supplementary Figure S43a). Out of 24 models, 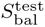 and the unweighted mean of the 22 metrics agree almost perfectly (Spearman ρ = 0.97), as do a nine-task subset taking one highest-signal task per family (ρ = 0.99), a greedy five-task panel (ρ = 0.98), and the first principal component of the score matrix (ρ = 0.90). Two out of four aggregations select the same 3 Botanic1 models as top performers as 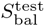; the greedy five-task panel and the principal component keep the same top two Botanic1 models and have PlantCaduceus and PlantCAD2-L respectively move to third place. This shows that our choices are both defendable and close to harmless in terms of final ranking.

Using 79 BiMamba2 runs between 100M and 700M parameters and fitting a Principal Component Analysis (PCA) shows that one principal component explains 91% of cross-model variance and the effective dimensionality of the score distribution is ≈ 1.2. On the contrary using the diverse set of baselines reported in Figure 3, which vary in architecture, training objective, model size and training data, the first principal component explains only 60% of the variance and the effective dimensionality rises to 2.4 (Supplementary Figure S43b). The single best metric (conservation in maize) still correlates with 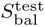 at ρ = 0.94, but the principal component itself no longer recovers the full top three. We consequently report cross-model comparisons on the full score together with the per-capability breakdown (Supplementary Figure S5).

This score is also motivated by the size of our evaluation cluster, which readily supports 22 metrics on all our checkpoints. However, the analysis shows that when compute is limited, reporting two to three well-chosen metrics should be sufficient to track model progress and rankings.

The second principal component has mean weights of 0.35 on LLR and 0.37 on causal variant discovery, the two zero-shot families, against positive weights on chromatin accessibility and PRO-seq. It therefore separates models that score variants well from models that are good at providing sequence-level embeddings. Families also differ markedly in how widely they discriminate the models and how closely their orderings follows the ordering of 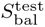; Section 4.3.3 quantifies both properties per family. We keep the low-variance families in *S*_bal_ for exhaustiveness.

#### 4.3.1 Are leaderboard differences significant?

*S*_bal_ is a mean over nine task families, so when observing a performance difference between two models there are two questions to be asked: would this difference survive an equally defensible choice of tasks and species, and, given this way of aggregating scores, is this difference statistically significant? We answer each of those questions with a paired bootstrap with 20,000 replicates; the text and Supplementary Table S7 report both, and the figures report only the example-level bootstrap, against PlantCAD2-L in Figure 3e as well as for every pair of leading models in Supplementary Figure S18.

##### Bootstrapping at the family level

For each pair of models we take the nine per-family differences, resample the nine families with replacement and report the 95% percentile interval of the resampled mean.

##### Per-sample bootstrap

The second bootstrap test resamples the test examples within each of the 22 tasks and species: the rows of the different test probes, the variants of the three LLR cells, and the 29 studies of the causal-variant task. Every model is scored on the same drawn rows: each difference between two models is paired at the example level, and the per-replicate cell scores are aggregated into *S*_bal_. Every example-level interval is the bootstrap’s 95% percentile interval overlaid with the reported value, and every error bar reported either on Figure 3d, Supplementary Figure S17 or on the per-family and per-metric size and token panels (Supplementary Figure S5 to Supplementary Figure S11) is the 95% CI.

The same example-level bootstrap is run on the twelve metrics held out of *S*_bal_ (Section 4.3.4), the crossspecies splice donor, splice acceptor, TIS and TTS cells, each resampled over its 5,000 stratified rows with the prevalence weights, with paired differences to PlantCAD2-L on the same draws (Supplementary Figure S8). Their intervals are on average (median) two and a half times wider than those of the *S*_bal_ cells, which is the reason these metrics were not included in the aggregate score.

##### Results

Botanic1-L and Botanic1-XL are significantly better than PlantCAD2-L under both tests, and Botanic1-XL is significantly better than both PlantCAD2-L and PlantCaduceus. Botanic1-M is not significantly better from PlantCAD2-L with the family-level test (+0.012, [ −0.002, +0.030]) but is using the per-sample test ([+0.004, +0.022]). Botanic1-S ties with PlantCAD2-L on both tests. Within the Botanic1 family, the step from Botanic1-S to Botanic1-M is significant under both tests (+0.006), the steps from Botanic1-M to Botanic1-L and from Botanic1-L to Botanic1-XL are not (+0.004 and +0.002).

A capability-level reading of the Botanic1-S against PlantCAD2-L comparison, and the comparison outside the tasks of *S*_bal_, are given in Section 2.3.1.

##### A note about guiding research decisions using part of the target objective

All models are trained with self-supervised MLM; therefore, pre-training does not directly optimise the downstream tasks. In addition, 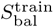 is never used to select a specific checkpoint within a training run. As shown in Figure 3a, some intermediate Botanic1 checkpoints score above the checkpoints we report at 314.6B, mainly because of optimisation noise. Steering data-mixture decisions with a downstream benchmark suite is standard in LLM pre-training literature [22, 56, 23, 58], and we follow the same process deliberately. However, this creates a potential asymmetry between the fixed competitor models and the multiple training runs used to improve Botanic1 candidates through choices such as data mixtures, batch sizes and learning rates (Section 4.3.6). We expect competitors to have followed similar best practices, although perhaps less systematically and without targeting the exact task set we choose. In practice, we expect the effect to be small relative to the margins in Supplementary Table S7. Reporting the per-capability breakdown (Supplementary Figure S5), the aggregate and many benchmarks that are not optimised helps reduce this ambiguity.

#### 4.3.2 The nine task families

Supplementary Table S4. gives the nine task families the 22 metrics are grouped into, with the species evaluated for each. Twelve metrics are held out of *S*_bal_: the six cross-species splice metrics and the six cross-species TIS and TTS metrics (both in Supplementary Figure S8a), discussed in Section 2.3.1.2. All twelve are scored on a stratified sample of test sets too large to evaluate exhaustively; at their positive rates the metric must be a prevalence-corrected AUPRC, and that estimate is too imprecise to enter the aggregate score (Section 4.3.5).

Seven families are supervised probes trained on top of frozen representations. They mostly focus on annotation tasks, which might help bioinformaticians in their day-to-day workflows: where genes begin and end (TIS, TTS), which regions are transcriptionally active (PRO-seq) or accessible (chromatin), which positions are evolutionarily constrained (conservation), where introns are spliced (splicing), and which genic region a window belongs to (genomic region classification). The remaining two families are zero-shot scoring using the log probability ratio of reference versus alternate allele described at length in the literature, notably in GPN [83].

Species evaluated, by family: chromatin accessibility on *A. thaliana*, *Oryza sativa* (MH63 and ZS97), *Setaria italica*, *Brachypodium distachyon*, *Sorghum bicolor* and *Zea mays*; genomic region on *A. thaliana*, *O. sativa*, *Z. mays* and *Solanum lycopersicum*; conservation on *S. bicolor* and *Z. mays*; splicing, TIS and TTS on *A. thaliana*; PRO-seq on *Manihot esculenta*; and LLR on *A. thaliana*, *Z. mays* and *S. lycopersicum*. Causal variant discovery is scored on a fixed subset of 29 curated studies from four species (13 *A. thaliana*, 11 *O. sativa*, 4 *Triticum aestivum*, 1 *S. bicolor* ), each causal SNP ranked by signed LLR against 100 background SNPs of its locus, and contributes a single recall-AUC scalar; the full 545-study benchmark (Section 4.10), which ranks every documented SNP of the locus by LLR , is evaluated separately in Section 2.5.1.2. We plot each task with a measure of its complexity in Supplementary Figure S44. Supplementary Table S5 defines each family precisely: source dataset, task type, window length, pooling and the metric under each protocol. Supplementary Table S6 complements it with, for every logged metric, the original train and test split sizes with their positive counts and the exact subsampling applied under 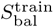 and 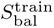.

#### 4.3.3 Which task families separate models?

Not every task family included Supplementary Table S4 contributes equally to ranking the models. One can measure two different informative properties for a task family: how widely a family spreads the models, and whether the ordering it produces agrees with 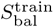. Supplementary Figure S19 highlights those properties for each of the 9 task families. Translation termination spreads the 24 models the more strongly (standard deviation 0.142), while PRO-seq, genomic region, chromatin accessibility and conservation barely separate models at all (0.039 to 0.059); Seven families of tasks follow 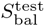 ordering closely (ρ = 0.69 to 0.94), but the two zero-shot families do not agree a lot with it (ρ = 0.56 for LLR and 0.55 for causal variant discovery). This complexity explains why we report the per-capability breakdown (Supplementary Figure S5) alongside 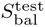 rather than relying on it alone.

#### 4.3.4 Why remove twelve frozen probe metrics from *S*_bal_?

Twelve of the 34 metrics the suite logs do not enter *S*_bal_: translation initiation and termination, and splice donor and acceptor recognition, each on rice, sorghum and maize. All twelve are scored on a class-stratified sample of test sets that are too large to evaluate exhaustively, and the resulting estimates are too imprecise to average into the aggregate score (Section 4.3.5). Supplementary Figure S8a shows the same qualitative picture as 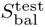: Botanic and PlantCAD2 separate from the rest across the twelve metrics, the models trained beyond plant genomes (Carbon, Evo 2) stay well below the plant-trained encoders even under the two-strand centre protocol described in Section 4.3.6.1, and performance is largely flat in parameter count above ∼ 10^8^ (Supplementary Figure S6); the lower rice TTS score of the largest Botanic model lies inside its 95% interval (Section 2.3.1.2). Translation initiation and termination are evaluated on all four species: *A. thaliana* on its full test set, and the three larger genomes by sampling. The maize test sets contain 11,268,672 and 29,539,071 candidate positions with only 3,098 true sites each, a positive rate of 0.01% to 0.03%. Scoring them exhaustively is intractable. In addition a raw AUPRC would be uninformative because it would reflect only the majority class. We therefore sample each test set to 500 positives and 4,500 negatives and report the prevalence-corrected metric, stratified AUPRC, whereas *A. thaliana*, whose positive rate is orders of magnitude higher, keeps hundreds of positives under a uniform subsample and is scored with the plain AUPRC (the sampling of every metric is detailed in Supplementary Table S6). The splice donor and acceptor tasks are evaluated similarly: their datasets are the largest test sets in the suite, with 34.8 and 40.3 million candidate positions for maize donor and acceptor, respectively, and 18 to 22 GB of raw sequence per split. They contain ∼ 24,000 positives per species, about eight times as many as the translation initiation and termination sets.

#### 4.3.5 Precision of the stratified cross-species metrics

Twelve of the metrics are scored on a class-stratified sample of 500 positives and 4,500 negatives which is reweighted to the true prevalence. Reweighting is important because their test sets hold between 1.4 and 40 million candidate positions and scoring them exhaustively for every model in Figure 3 would cost on the order of 10^4^ NVIDIA H200 GPU-hours, extrapolated from the per-model embedding throughputs measured on the evaluation cluster. We want to know whether that estimate is precise enough to enter *S*_bal_ so every task-species is re-scored with all positives kept and the negative sample scaled from 45,000 to 2 million reaching the original number of positives on some datasets.

##### 4.3.5.1 The estimator is consistent, but the estimate is positively biased

At a positive rate near 10*^−^*^4^ even a half-million-row uniform sample holds fewer true sites than the 500 the estimate uses by construction so scaling positives does not help resolve the score. On the contrary, scaling negatives converges to the true estimate. On translation initiation in rice, the one pool small enough to score exhaustively, the positive-complete estimate lands within 0.001, 0.013 and 0.024 of the exact AUPRC for GPN, PlantCAD2-L and Botanic1-L. The published 500-positive values are another matter: they sit a median 0.088 from their converged value, 33 of the 36 beyond 0.03, and 28 of the 36 above it (one-sided binomial *p* = 6 × 10*^−^*^4^). The estimate is therefore noisy and systematically optimistic.

##### 4.3.5.2 The estimator is too noisy to be included in *S*_bal_

*S*_bal_ is an unweighted mean of nine families, so a family carrying that much error contributes 0.006 to 0.010 of uncertainty to the aggregate score, against a median 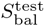 gap of 0.0043 between neighbouring models in the top eight and 0.0020 between Botanic1-XL and Botanic1-L. Admitting these metrics would inject as much uncertainty as the gap between models.

##### 4.3.5.3 Standard parametric model fitting does not explain the measured noise level

Calibrating a bi-normal distribution, the standard model for binary classification scores (negatives pinned to N(0, 1) and positives to N(θ, 1), where θ is estimated by Brent’s method from the exact AUPRC without sampling), and replaying a 500-positive, 4,500-negative draw from that simple model predicts both the upward bias of the estimator and its dependence on prevalence: from 0.003 at the highest prevalence in the suite (0.58%) to between 0.07 and 0.14 at the lowest (0.010%). However, this simple model predicts only about half the observed magnitude, a median of 0.047 against 0.088. We use the “normal-deviate axes” trick to replot the 12 ROC curves after passing both coordinates through the inverse of the standard normal CDF, the probit function Φ*^−^*^1^: instead of the false-positive and true-positive rates (FPR, TPR) we plot (Φ*^−^*^1^(FPR), Φ*^−^*^1^(TPR)). So a rate of 0.5 maps to 0, 0.84 maps to +1, 0.977 maps to +2, and so on: the axes are now in units of standard deviations (“normal deviates,” i.e. z-scores). Suppose the scores are bi-normal: negatives drawn from N(µ*_−_*, σ^2^ ) and positives from

N(µ_+_, σ^2^ ). At threshold t, we have:

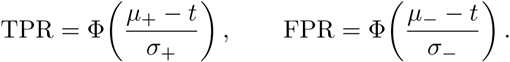

Apply Φ*^−^*^1^ to both and eliminate t, and one obtains an exact straight line:

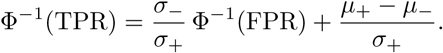

So with those new axes, bi-normal scores produce a straight line, and the slope of that line is σ*_−_*/σ_+_, the ratio of the two spreads. Curvature means the parametric fit is not accurate; a straight line with slope 1 means equal spreads; a straight line with slope below 1 means the positives are more spread out than the negatives. In this new frame, we observe pronounced curvature and a median slope of 0.61, indicating that positives are about 1.6 times as dispersed as negatives. Substituting that measured dispersion into the model lowers the prediction to 0.030, even further from the observed metric. The distributional assumptions are therefore not met; the most plausible cause is the non-independence of test rows, which is difficult to measure. The AUROC cannot disambiguate the cause because all models have scores in a narrow band between 0.961 and 0.997, whereas the AUPRCs span 0.07 to 0.85; one fixed AUROC admits almost any AUPRC under standard score families. Supplementary Figures S13 and S14 show the fit results.

##### 4.3.5.4 The stratified estimator of the splicing tasks in *S*_bal_ is well-behaved

The two splicing metrics within *S*_bal_, donor and acceptor recognition in *A. thaliana*, are scored on a stratified subset drawn to match the target class prevalence of the full dataset (2,400 positives against 97,600 negatives for donor, 3,500 against 96,500 for acceptor), using the prevalence-corrected stratified AUPRC metric as a score under 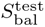 (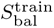 scores the same draw with the prevalence-corrected stratified AUROC). These draws keep respectively 26% and 40% of the full test datasets (377,873 and 250,084 rows, prevalences 2.3% and 3.5%), when the cross-species estimator, excluded from *S*_bal_, keeps only under 0.4% of datasets of 1.4 to 40 million rows at target prevalences near 10*^−^*^4^ for the minority class. To measure the quality of the estimator directly we re-score the full test datasets for six models spanning the experiment settings, and replay 10,000 fresh prevalence-matched draws per task (Supplementary Figure S16). The estimator is unbiased to within 10*^−^*^4^ of corrected AUPRC; its standard deviation is 0.002 to 0.003 for the five models above 0.97 and 0.006 for AgroNT; the published result is within 0.004 of the exact full-dataset number for every model except AgroNT on acceptor, which is at 0.006, about one standard deviation away (Supplementary Table S3). The splicing family used in *S*_bal_, contributes at most 4 10*^−^*^4^ of uncertainty to *S*_bal_, and under 2 10*^−^*^4^ for the leading models, an order of magnitude below the 0.0020 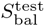 gap between Botanic1-XL and Botanic1-L. In paired draws the family ordering of two models reverses in more than 5% of draws only when their full-dataset gap is below 0.002, so only for actual ties at the precision of the table.

#### 4.3.6 Competitor models

Every baseline evaluated in this report can be downloaded from HuggingFace (HF). Supplementary Table S1 details the evaluated models. More precisely, PlantCaduceus-l20, -l24, -l28 and -l32 [34] are downloaded from the HF repositories kuleshov-group/PlantCaduceus l20, l24, l28 and l32; PlantCAD2-S, -M and -L [33] from kuleshov-group/PlantCAD2-Small-l24-d0768, PlantCAD2-Medium-l48-d1024 and PlantCAD2-Large-l48-d1536.

NTv3-100M-pre, NTv3-650M-pre, NTv3-100M-post and NTv3-650M-post [25] are taken from InstaDeepAI/NTv3 100M pre, NTv3 650M pre, NTv3 100M post and NTv3 650M post; AgroNT [31] from InstaDeepAI/agro-nucleotide-transformer-1b; GPN [83] from songlab/gpn-brassicales. Carbon-500M, -3B and -8B [94] are taken from HuggingFaceBio/Carbon-500M, Carbon-3B and Carbon-8B, PlantGFM [82] from hu-lab/PlantGFM, and Evo 2-1B, -7B, -20B and -40B [24] from arcinstitute/evo2 1b base, evo2 7b, evo2 20b and evo2 40b. MarinDNA-1B [26] is taken from marin-dna/marin-dna-exp135-m5.1 and PlantBiMoE [30] from plant-llms/PlantBiMoE. The Plant-DNA-LLM suite [62] contributes 22 checkpoints, all under the zhangtaolab/ namespace: plant-dnabert-6mer, -singlebase and -BPE; plant-dnamodernbert-singlebase and -BPE; plant-nucleotide-transformer-6mer, -singlebase and -BPE; plant-dnagpt-6mer, -singlebase and -BPE; plant-dnagemma-6mer, -singlebase and -BPE; plant-dnamamba-2mer, -3mer, -4mer, -5mer, -6mer, -singlebase and -BPE; and plant-dnamamba2-BPE.

##### 4.3.6.1 Embedding layer and pooling

As in Botanic0 [32], we probe the default embedding exposed by each model’s implementation, which for almost every backbone is the last hidden layer. We do not grid-search over layers even though we know from [119, 24] the best layer of a model is task-specific. Embeddings are precomputed once per (model, task) and reused for every probe trained on that task. Two pooling rules are used: for the point-feature tasks (conservation, TIS, TTS and splicing) we take the embedding of the token at the centre of the window following [34] and [32]; for the window-level tasks (PRO-seq, chromatin accessibility) we take the mean over all tokens. The Genomic region classification task follows the GPN protocol [83]: the central 100 bp of the 512 bp context is first mean-pooled and the forward and reverse-complement embeddings are then averaged for encoders, while decoder-only models concatenate them, following the same reasoning as for the point-feature tasks. Under single-nucleotide tokenisation the centre token is nucleotide 255 of a 512 bp window; under k-mer tokenisation it is the k-mer containing that nucleotide; the k-mer tokenisers used here advance by a fixed stride, so that index stays sequence independent even though a small fraction of windows carry N bases (below 0.2% of rows in every evaluated split). However when using byte-pair encoding (BPE) the centre-token index depends on each sequence, so it is derived from the tokeniser’s offset mapping and cannot be vectorised as easily.

For decoder-only backbones (Carbon, Evo 2, PlantGFM and the causal Plant-DNA-LLM models) pooling the centre token of a single forward pass would make models blind to the right half of the window, so on most of the sequence-level tasks with a central motif we run two forward passes, one on the sequence and one on its reverse complement, we take the centre token of each strand and concatenate the two embeddings, following the Evo 2 [24] methodology; on the window-level tasks they are mean-pooled like every other backbone. This two-strand protocol gives all decoder-only models two forward passes per sequence and injects the prior that both strands carry the same information. Scoring splicing with a mean over its whole 398 bp window with a single forward pass instead costs 0.75 to 0.91 of corrected stratified AUPRC for an encoder and an autoregressive model alike, an order of magnitude more than any sampling choice (Supplementary Figure S15). It is debatable whether or not the centre-token pooling protocol used in the literature is realistic; we discuss that assumption in Section 2.3.1.

##### 4.3.6.2 Frozen-embedding probe training

All sequence-level tasks using frozen embeddings use gradient-boosted decision-tree classifier probes (XGBoost [120]) trained on the frozen embeddings from each model. Probe hyperparameters are fixed per task family and pooling protocol, and every model scored under a given pooling protocol uses an identical probe configuration (XGBoost hyperparameters). TIS, TTS and conservation reuse the configuration released with PlantCaduceus [34]: maximum tree depth 6, learning rate 0.1, L2 regularisation 1.0, and up to 1,000 boosting rounds with early stopping after 10 rounds without improvement; under this configuration our probes reproduce the published PlantCaduceus and GPN values within 0.01 AUPRC on TIS and TTS. PRO-seq, chromatin accessibility and splicing keep the same hyper-parameters with a learning rate of 0.05 and no L2 regularisation for encoders, while decoder-only models on splicing, scored under the RC-concatenated centre pool, keep the learning rate of 0.1 and the L2 regularisation of 1.0, and the multi-class genomic region task uses a learning rate of 0.3 with at most 100 rounds (early stopping after 3 rounds without improvement). Every probe fits XGBoost using the following hyperparameters: tree method=hist on GPU, max depth = 6, min child weight = 1, gamma = 0, alpha = 0 and max delta step = 0. The same hyperparameters were used in Botanic0 [32].

##### 4.3.6.3 Model-specific considerations

For NTv3, we follow the protocol of [32] and target the nucleotide-level embedding from the deconvolutional tower using NTv3 650M post, consistent with how the authors perform their fine-tuning experiments. We additionally score the pre-train-only checkpoint NTv3 650M pre, using its last transformer layer as described in the reference implementation; scoring the pre/post pair isolates the contribution of the post-training stage in both directions (Supplementary Figure S15d and Section 3). The post variant additionally requires the species name as a string argument; the plant species in our tasks are not all in its supported list, so we pass *Arabidopsis thaliana* when the species is not recognised. For PlantCaduceus and PlantCAD2, we use the same reverse-complement averaging scheme as the original code, resulting in RC-equivariant embeddings and, for zero-shot scoring, logits are averaged over both strands. The Plant-DNA-LLM models are trained on capped contexts, particularly DNABERT as a cap at 512 tokens: when a task’s tokenised window exceeds the budget, we crop a window centred on the labelled position instead of truncating from the right, which would discard the site being classified. For MarinDNA, which is trained on only 255 bp sized windows, we score only the tasks that fit natively at 255 bp: genomic region classification on the four species, the three LLR species and causal variant discovery recall. The remaining tasks would require cropping windows to half their length or less, which would make the comparison unfair.

##### 4.3.6.4 Evo 2 inference

Evo 2 embeddings are read from intermediate layers: blocks.21.mlp.l3 for the 1B and 20B models, blocks.28.mlp.l3 for 7B and blocks.43.mlp.l3 for 40B. These layer choices follow the Evo 2 recommendations for its 7B sized model and use it as a reference across model depths to identify a hopefully equivalent layer at the same 87.5% relative depth in all model sizes. We also test the final post-normalisation layers, which sharply degrade performance. Evo 2 is the only model for which we had to study several layers because our first implementation using post-normalisation embeddings showed near-random performance and therefore warranted further investigation. Evo 2 inference uses the dedicated vortex implementation and its Transformer Engine kernels; its input projections use FP8 on GPUs of compute capability 8.9 or above. We evaluate it on H200s (Hopper) with FP8 enabled. Disabling FP8 or using different hardware raises no error but can degrade the output to near-random. Because this failure is silent, in our code every Evo 2 evaluation must pass the tests committed by the Arc Institute in the evo2 repository.

#### 4.3.7 Training-token accounting for competitors

Figure 3d (right) and Supplementary Figures S7 and S11 report, for every model evaluated, the total base pairs seen by this model during training. However, some ambiguity remains because accounting methods differ and original publications contain missing information or conflicting statements. To resolve that ambiguity, we use the numbers reported by the target paper or otherwise rederive them from hyperparameters: steps × global batch × sequence length, or pre-training size × reported epochs, multiplied by the tokeniser’s bases per token. Every value is verified against the model’s publication, supplementary material or model card, and Supplementary Table S2 documents each count with its derivation, source location and caveats. We will update this report if original authors spot inaccuracies.

Native token counts are not directly comparable across tokenisers: one token covers one base for singlenucleotide and byte tokenisers, approximately six bases for the 6-mer models (AgroNT, Botanic0, Carbon) and about seven for the BPE variants of the Plant-DNA-LLM suite. The axes therefore plot the base-pair conversion, and Supplementary Table S2 keeps the original counts in tokens. This adds another potential source of error. NTv3’s post is counted as 12.1T (the sum of its stated stage budgets including preand post-training); the three Plant-DNA-LLM checkpoints without a stated budget (the two ModernBERT variants and DNAMamba2) are omitted from the token panels; and Evo 2-20B is drawn at 9.3T, the budget of its Evo 2-40B parent from which it originates through model surgery, which does not require retraining [121].

#### 4.3.8 Fine-tuning hyperparameters

The fine-tuning experiments of Table 1 select hyperparameters using exhaustive grid search, independently for each task backbone fine-tuning adaptation regime. Every combination in Supplementary Table S9 is first trained and evaluated on a validation split, and the combination with the best validation metric is retrained from scratch on the full training set to produce the reported performance. The hyper-parameters grid is not shared across backbones: as compatible learning-rate ranges differ widely between models. The two classification tasks additionally search the classification head’s learning rate and the pooling operator.

##### 4.3.8.1 Parameter Efficient Fine-tuning methods

LoRA [35] and IA^3^ [36] were introduced originally for Transformers and are defined on the attention projections: LoRA adds a low-rank update to the query, key and value matrices, and IA^3^ rescales the keys, the values and the input of the second feed-forward matrix with learned vectors. Botanic1-S and PlantCAD2-L have no attention layers so it is not clear what LoRA or IA^3^ means for any of those models. Both Botanic1-S and PlantCAD2-L replace attention with Mamba2 modules [85]. Every Mamba2 module has two dense linear layers: an input projection, which maps the input block (d) to the gate branch and the state-space input (2d each), the input-dependent B and C matrices and the step sizes, and an output projection, which maps the 2d state-space output back to d. Following recommendations from the state-space model fine-tuning literature for LoRA [122, 123], LoRA adds a rank-8 update to these two projections of both directions of every block; the recurrence parameters (A, D, the depthwise convolution, the step-size bias and the norms) stay frozen. For IA^3^, we rescale the output of the input projection and the input of the output projection by analogy with the recommendations above for LoRA. For AgroNT, an ESM-style Transformer, we use LoRA normally on query, key and value, IA^3^ on keys, values and the second feed-forward matrix. Adapters are implemented using the PEFT library [124], and in every fine-tuning experiment, including the two parameter-efficient ones, the task heads are fully trained.

For every LoRA run reported here, when it is not specified, we use rank 8, a scaling factor α/r = 1 (α = 8) and no adapter dropout.

The selected configurations follow, verbatim, one file per task. Levels are backbone regime subtask: for promoter strength, poly(A) and lncRNA the grid search runs per subtask. Terminator strength trains one head to predict both systems and therefore has a single configuration per backbone × fine-tuning regime across the two systems. PlantCAD2-L fine-tuning has severe instabilities and leads to NaN gradients with standard configurations on some tasks including poly(A). Therefore it requires an fp32 classification head (head fp32) and a low backbone learning rate (1–3 10*^−^*^6^): in bf16 its head-logit projection overflows and every species collapses to constant output, whereas the fp32 head recovers five of six species (see Table 1 note *^a^*).

##### 4.3.8.2 Per-species tuning

Table 1 reports a per-species number alongside the global number for classification tasks. The first number in the table uses hyperparameters tuned once across all species of a task. The second uses hyperparameters tuned separately for each species and reports the better of the two highest-ranked validation configurations after retraining for that species (top-two runoff).

##### 4.3.8.3 From-scratch baselines

The CNN column of Table 1 is not retrained. Both task-specific models’ authors, Gorjifard et al. [63] and Jores et al. [64], publish their test-set prediction values per sample, which allows us to recompute any unreported metrics. We recompute the reported metrics from those released predictions using the same metric implementation that we use elsewhere in the article.

### 4.4 Long-context evaluation

#### 4.4.1 Pseudo-perplexity evaluation

We score PPPL [67] under a paired masked-position protocol. For each of 1,251 sequences we subsample 30 positions and mask each in turn. We report the corpus PPPL, the exponentiated mean negative log-likelihood over all scored positions of all sequences. Each scored position carries the same per-position repeat weight the training loss uses, 1 (1 w)f with *w* = 0.1 and *f* the soft-mask repeat fraction of the position (Section 4.1), so that the metric scores the sequence content the model is trained to model rather than the repeat fraction of whichever window a length happens to draw. For every pPPL we report we use this same weighting. Bootstrap distributions samples sequences with replacement (10,000 draws), using the same draws for every checkpoint so that bootstrapped models performances can be aligned and substracted. Comparing *k* checkpoints against the reference, we form each paired differences, studentise each difference by its bootstrap standard error, and take the distribution of the maximum absolute studentised difference as the null hypothesis; a family-wise-corrected p-value is the fraction of that null hypothesis at or above the observed statistic. We evaluate on both training and held-out data from the same sources at fixed length, and add a nested-context evaluation so that scored positions can be paired across lengths. We take held-out 2^17^ bp sequences and restrict the scored positions to the central 2^13^ bp subwindow, then grow the window symmetrically by doublings up to 2^20^ bp.

#### 4.4.2 Long-context retrieval (NIAH)

One way to run NIAH on DNA is Evo 2’s categorical-Jacobian score [24, 125], but it is expensive: it requires four forward passes per scored position, and its logit-based score must be null-normalised before it can be compared across models, doubling the budget. Thus, we adopt here a key-value variant analogous to LLM NIAH benchmarks [68]: a key-value pair is inserted at a controlled depth within a random haystack and repeated at the end of the context with its value masked (Figure 6b). Taking advantage of the ability of an MLM to decode multiple tokens of a span in parallel, we recover the whole value in a single forward pass, which makes the probe two orders of magnitude cheaper. All our NIAH evaluations use a 16 bp value, since decoding it in one pass matches decoding 1 bp at a time (0.83 vs. 0.83) and it is the longest value span still decoded reliably; beyond it performance collapses (0.55 at 32 bp, 0.22 at 64 bp, Supplementary Figure S28b). We use 100 bp keys because shorter keys degrade retrieval, the score falling from 0.83 at 100 bp to 0.69 at 25 bp (Supplementary Figure S28a). a grid spans ten depths (0 to 90%) and haystack lengths from 2^10^ to 2^17^ bp. The haystack’s L bases are drawn uniformly at random. One cell of the grid fixes a haystack length and the relative depth at which the needle is inserted: writing *s* for the key and value lengths combined, the needle sits at an offset of round(depth (L 2s)) and the query (the repeated key and the masked value) occupies the final *s* positions. The generator is seeded by the key length, value length and repeat index. a repeat scores the fraction of value positions recovered, a cell the mean over 8 repeats; tokenisation being single-nucleotide, chance level corresponds to an accuracy of 1/4. To summarise a grid as one number we rescale each cell’s accuracy so that chance maps to 0 and perfect recovery to 1, clip below at 0 and average.

### 4.5 Base-resolution ATAC-seq prediction

Our pre-trained Botanic1 models are trained in a self-supervised manner, so to predict the ATAC-seq signal at base resolution we add a task-specific head to the backbone and fine-tune the backbone and head on ATAC-seq data from two plant species, *Arabidopsis thaliana* and *Zea mays*.

#### 4.5.1 Datasets and preprocessing

We download and reprocess the ATAC-seq libraries used in the fine-tuning of NTv3 [25] following ENCODE processing standards [126]. In short, we obtain sequencing reads from the European Nucleotide Archive (ENA) and trim adapters and low-quality bases with fastp [127]. We discard reads shorter than 15 bp after trimming. We align the trimmed reads to TAIR10 and Zm-B73-REFERENCE-NAM-5.0, for *A. thaliana* and *Z. mays* respectively, with Bowtie 2 [128], using a maximum fragment length of 2,000 bp (-X 2000) and reporting one alignment per read (-k 1). We filter aligned reads with samtools [129] to retain properly paired reads (-f 2) with a minimum mapping quality of 30 (-q 30) and to remove unmapped reads, secondary and supplementary alignments, and PCR/optical duplicates ((-F 1804)). We correct mate information with samtools fixmate, quantify and remove mitochondrial and plastid reads, and mark and remove duplicates with Picard MarkDuplicates [130]. We convert the filtered, de-duplicated alignments to ENCODE tagAlign format, applying the standard +4/-5 shift. We call peaks with MACS3 [131] (--nomodel --shift -75 --extsize 150 --keep-dup all) at a false discovery rate of *q* < 0.01, using a per-genome effective genome size computed as the non-organelle, mappable (non-N) sequence length. We generate +4/-4 shifted Tn5 insertion tracks directly from the filtered, de-duplicated alignments to use as the training target. We verify these tracks are byte for byte identical to those generated by the ChromBPNet pipeline. We evaluate the quality of each library on fraction of reads in peaks (FRiP), TSS enrichment, library complexity, duplication rate, mitochondrial and chloroplast read fractions and fragment length distribution. For each species, we train ChromBPNet and perform fine-tuning on the library with the highest TSS enrichment for which ChromBPNet’s Tn5 shift detection succeeds. For *Z. mays* this is SRX21046141 (TSS enrichment 6.150). For *A. thaliana*, the library with the highest TSS enrichment (SRX21812658) is excluded because ChromBPNet’s automatic shift detection fails, most likely due to the high AT content of the *A. thaliana* genome, so we instead use the next-highest, SRX8571616 (TSS enrichment 2.77).

#### 4.5.2 ChromBPNet baseline

ChromBPNet [37] is a base-resolution convolutional model of chromatin accessibility that takes a 2,114 bp input and predicts the signal over the central 1 kbp. It is composed of two BPNet models [132]: a Tn5 bias model, which captures Tn5’s sequence-preference bias and is trained on background (non-peak) windows, and an accessibility model. ChromBPNet adds the frozen bias model’s output to that of the accessibility model, yielding a sum equal to the observed ground truth signal. The model has two heads: a count head, which predicts the total read coverage over the input window that represents the total magnitude of accessibility, and a profile head, which predicts the base-resolution shape of the signal as a multinomial distribution over positions.

We train ChromBPNet with its official pipeline on the same ATAC-seq data used for fine-tuning, for both species, and evaluate it with the same metrics as our models.

#### 4.5.3 Input windows and data splits

For each species, the data-preprocessing step (Section 4.5.1) provides base-resolution ATAC-seq signal tracks in BigWig format. We construct 2,114 bp input windows centred at peak summits, evaluating performance over the central 1 kbp while 557 bp flanking regions on each side are ignored in the loss and provide context only.

Peak windows are centred on the summit, which places the maximum signal at a fixed index in every training example. To remove that dependency, we jitter the training windows following ChromBPNet: each training window is randomly shifted by up to 500 bp, so the summit no longer sits at a fixed position. Jitter is applied to the training split only; validation and test windows are not jittered.

Each window is either a peak or a background window. Peak windows are the called ATAC peaks; background (non-peak) windows are the non-peak regions used to train ChromBPNet bias model. Botanic1-S and ChromBPNet are fine-tuned on the same peak and background windows in each species. Every epoch uses all of the peak windows. The background windows are resampled each epoch; we draw a fresh random subset of them, its size set by a negative sampling ratio of 0.1, one tenth of the number of the peak windows. Although evaluation is over the peak set alone, background windows are included in training, as for ChromBPNet, so the model learns to separate accessible from inaccessible sequence rather than only fitting variation within peaks.

We discard windows whose total count falls outside ChromBPNet’s outlier bounds, below the 0.0001 quantile or above the 0.9999 quantile of the per-window total-count distribution over the pooled peak and background windows (the composition used to set λ). We apply no normalisation to the BigWig tracks; the signal is the raw per-base read count.

We split the data by chromosome. For Arabidopsis, chromosomes 1, 3, and 5 are used for training, chromosome 2 for validation, and chromosome 4 for testing. For maize, chromosomes 1 and 9 are held out for testing, chromosome 4 for validation, and the remaining chromosomes (2, 3, 5, 6, 7, 8, and 10, with unplaced scaffolds) for training.

#### 4.5.4 Prediction heads, training objectives, and fine-tuning regime

For ATAC-seq prediction with our pre-trained Botanic1-S backbone, we attach a prediction head to it. Following ChromBPNet’s head design, our dual-head design has two heads: a profile head that predicts the per-position logits through a single layer projection, and a count head that takes Botanic1-S’s last hidden states, applies mean pooling over the scored region and projects them to the log1p of the total signal.

The single head uses a linear projection layer followed by a softplus, which predicts the per-base signal on the log1p scale (so it is non-negative). The total read count over the scored region is then derived from these per-base predictions: we map each back to counts, sum them, and take log1p, that is, log 1 + *_i_*(e*^si^* − 1) for per-base log1p signal *s_i_*, so it has no separate count head. This follows the NTv3 [25] head design.

The dual head is trained with a profile term and a count term (Equation (2)). The profile term is the multinomial negative log-likelihood of the observed per-base counts under the softmax of the profile logits over the scored region, and the count term is the mean-squared error on the log1p total:

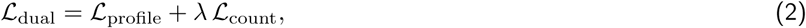

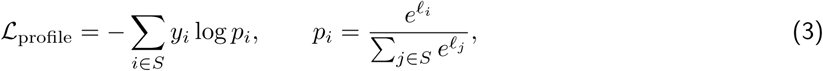

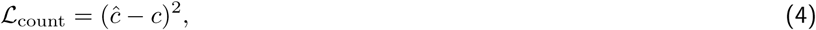

where *S* is the scored region, ℓ*_i_* and *y_i_* are the predicted logit and observed count at position i, and ĉ and *c* = log(1 + *_i∈S_* *y_i_*) are the predicted and observed log1p totals. The count weight λ is computed per species following ChromBPNet’s implementation, as the median total count over the pooled train and validation windows (after trimming at the 0.9999 quantile), divided by ten and floored at one, which gives λ = 120.5 for Arabidopsis and λ = 27.2 for maize.

What motivates the single-head design is the burden of recomputing λ for every species as an extra hyperparameter in the dual-head design. Because a single per-base head predicts both the shape and the magnitude of the signal, the total can be derived from it, so no count weight is needed. Its loss is a single masked mean-squared error between the predicted per-base signal and the observed counts. The predicted signal is on the log1p scale, so the observed raw counts are log1p-transformed to the same scale before the error is computed:

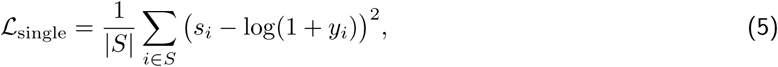

where *s_i_*is the predicted per-base signal on the log1p scale at position *i*.

We fully fine-tune the backbone and head together, with no frozen weights, and train each head design under two backbone initialisations: the pre-trained Botanic1-S weights and a randomly initialised backbone (from scratch). Comparing these two initialisations helps assess the contribution of pre-training. Each configuration is trained once with a single random seed, so we do not assess variability across training runs. In all cases we optimise with AdamW (β_1_ = 0.9, β_2_ = 0.999) under a differential learning rate, 10*^−^*^3^ for the head (for maize Botanic1-S from scratch single-head the learning rate is set to 3 10*^−^*^4^) and 10*^−^*^4^ for the backbone (a 0.1 backbone multiplier), an effective batch size of 64 (per-device batch 8 with 8 gradient-accumulation steps), and a linear learning-rate decay without warmup. We train for up to 15 epochs on Arabidopsis and 20 on maize, evaluate once per epoch, stop early after 5 epochs without an improvement in validation loss, and keep the checkpoint that has the best validation loss.

#### 4.5.5 Evaluation metrics and significance testing

##### 4.5.5.1 Peak counts

We assess prediction magnitude over peak windows using count Pearson and Spearman correlations. Count Pearson is the Pearson correlation between predicted and observed log1p total counts. Spearman measures the corresponding rank correlation and therefore tests whether predicted accessibility ranks the peaks in the same order as the observed signal.

##### 4.5.5.2 Peak profile

Profile predictions are evaluated using the Jensen-Shannon distance (JSD), in both raw and normalised forms. For both measures, we report the median over the peak set.

For a peak window, let *p* = softmax(***ℓ***) be the predicted profile over the scored region, *q* the observed profile (per-base counts normalised to sum to 1) and *u* the uniform distribution over that region. Raw JSD (d_raw_) and normalised JSD (d_raw_) are defined as follows:

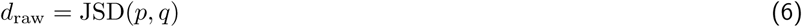

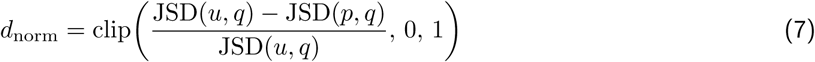

Raw JSD measures the distance between the predicted and observed profile distributions, and lower values indicate that the two profiles are similar. Normalised JSD compares this distance with the distance between the observed and a uniform profile and rescales the result to [0, 1]. a value of 1 indicates that the observed and predicted profiles are identical, while 0 indicates that the predicted profile is no better than a uniform distribution.

##### 4.5.5.3 Uncertainty

We use a bootstrap over the held-out test peaks to estimate the uncertainty of each metric and test whether the differences between models are significant.

In each replicate, we resample the test peaks with replacement and recompute each metric: count Pearson and Spearman from the resampled observed and predicted totals, and raw and normalised JSD as the median of the resampled per-peak values. We repeat this 1,000 times and for each metric, we use the 2.5th and 97.5th percentiles of the resulting distribution as estimates of the bounds of the 95% confidence interval (Table 2).

To compare two models, we use a paired bootstrap (the same resampled set of peaks is provided to both models in each replicate). For each metric, the difference between the two models can thus be measured per peak. We estimate the 95% confidence interval with the same method as above. The difference between the two models is considered statistically significant if 0 is not in this interval.

A fixed random seed is used for reproducibility.

#### 4.5.6 Base-level attribution by in silico mutagenesis

##### 4.5.6.1 Procedure

To determine which nucleotides in the input sequence drive the model’s ATAC-seq predictions, we perform in silico mutagenesis (ISM). We mutate each base to the three alternative nucleotides and re-evaluate the resulting sequence using the model. The change in prediction quantifies the contribution of each base: important nucleotides for the model yield substantial changes, whereas irrelevant ones produce minimal effect.

For each locus, we take the 2,114 bp input window and perform ISM across the central 1 kbp region. At each position within this region, we substitute the reference nucleotide with each of the three alternative bases and evaluate the modified sequence using the best-validation Botanic1-S (dual-head, pre-trained) model. For each locus, this requires 3,001 forward passes (1,000 positions × 3 alternatives, plus the reference). ISM relies exclusively on forward passes; therefore, no activations are stored. We run inference in float32 precision using a batch size of 64.

We select ISM over gradient- or reference-based attribution methods [133, 134] because standard attribution rules cannot be easily applied to BiMamba2 architectures. In the BiMamba2 selective state-space block, several input-dependent branches interact, including the *y* silu(z) gating and the dynamic B, C, and Δt terms. This makes methods such as DeepLIFT [133] which rely on defined rules for propagating contributions through nonlinear operations, difficult to apply directly. In addition, rule-based attribution requires access to intermediate operations, which the fused selective-scan implementation does not expose. We therefore use ISM, which only requires forward passes on mutated inputs and can be applied in the same way to both BiMamba2 and our convolutional baseline. The single-nucleotide tokenisation gives each sequence position a corresponding token, so we can introduce a mutation by directly replacing that token without computing positional offsets. Rather than hard-coding nucleotide token IDs, we retrieve them directly from the tokeniser at runtime.

We calculate contributions separately for each head because they produce different output types. The count head predicts a single scalar value representing the log1p total accessibility over the scored region. a mutation’s contribution is defined as the difference between predictions for the mutated and reference sequences. The profile head produces a probability distribution over positions in the scored region. a mutation can therefore alter the entire distribution, precluding a direct scalar difference. Following ChromBPNet, we reduce the change in the profile to a single contribution score. At each position, we multiply the change in probability by the corresponding probability in the reference profile, then sum these values across positions. This gives greater weight to changes at positions with higher predicted signal in the reference profile.

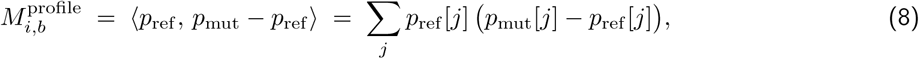

with *p* = softmax of the profile logits over the scored region. We use probability differences rather than log-probability differences so that the contribution score can take either sign. In log-probability space, the corresponding weighted difference is:

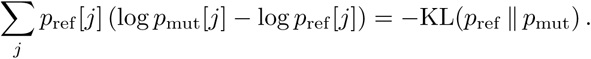

which is non-positive for any mutation. The probability-space inner product, by contrast, can be either positive or negative, allowing mutations with opposing effects on the reference-weighted profile to be distinguished.

Contributions are mean-normalised following the BPNet/ChromBPNet convention. Writing M*_i,b_* for the score shift when position *i* is set to base b, we take 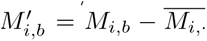*_·_* and report the value at the reference base, 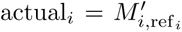. Since M*_i,_*_ref_ = 0 by construction, this quantity is proportional to the negative mean contribution of the three alternative nucleotides, 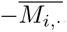. The reference nucleotide therefore does not need to be re-evaluated. Under this convention, a positive contribution indicates that mutations away from the reference decrease the prediction, and thus that the reference nucleotide supports the predicted signal. We retain both the hypothetical contribution matrix of shape (*L*, 4) where *L* is the number of scored positions (the central 1 kbp), and the projected contribution vector of shape (*L*, ). The former is used as input to motif-discovery tools.

##### 4.5.6.2 Motif selection

Motifs are taken from JASPAR CORE plants (2024, non-redundant position frequency matrices) [135]: 805 matrices, of which 601 are at least 8 bp. Shorter matrices are excluded because they match by chance every few hundred bases, making occurrence in a window uninformative. To scan for a motif, we convert its JASPAR position frequency matrix (the per-position counts of A, C, G and T) into a log-odds scoring matrix, which scores a candidate site by how much more probable its sequence is under the motif than under a background base composition. Each position’s probabilities are first mixed 99:1 with the background so that a zero count cannot produce an infinite score, and scores are taken as log_2_ of the resulting ratio. The background is estimated from the sequences themselves, all test-split windows, peaks and background alike, over the full 2,114 bp context of each (Arabidopsis: a 0.326, C 0.175, G 0.174, T 0.324), rather than assumed uniform at 0.25. Plant genomes are often AT-rich, so using equal background frequencies for all four bases can give too much weight to a and T matches and increase the scores of AT-rich motifs.

When scanning a sequence, the scoring matrix assigns a score to each position. Positions with score above the chosen threshold are considered motif matches. We define this threshold using a per-placement p-value. a threshold of *p* = 10*^−^*^6^, for example, means that under the background model, a randomly generated placement has probability 10*^−^*^6^ of achieving a score at least as high as the threshold. Converting this p-value to a score cutoff requires the null distribution of motif scores, which we compute exactly using dynamic programming over the binned score distribution rather than by sampling. We use a threshold of 10*^−^*^6^ rather than the conventional 10*^−^*^4^, to reduce the number of chance motif matches. Scanning a 1 kbp window on both strands involves approximately 2,000 motif placements. At a threshold of 10*^−^*^4^ scanning about 2,000 positions gives an average of 2,000 10*^−^*^4^ = 0.2 chance matches per window, corresponding to an approximately 18% probability of at least one chance match under an independence approximation. Such background matches can attenuate peaks-vs-background enrichment towards 1. For example, DOF1.5 shows 1.23× enrichment at 10*^−^*^4^, compared to 2.21× at 10*^−^*^6^.

Both strands are scanned, and overlapping above-threshold placements are merged into a single motif occurrence to avoid counting the same candidate binding site multiple times at adjacent offsets.

Enrichment is defined as the ratio of fraction of peak windows containing exactly one motif occurrence to the corresponding fraction of background windows. We compute this ratio over the complete test split (4,646 peaks and 4,646 background windows for Arabidopsis) rather than over a subsample. At stringent thresholds, motif occurrences in background windows are rare, making estimates from subsampled data unstable.

All 601 matrices are ranked by enrichment. Of these, 206 yield at least 50 usable windows and 35 show at least fivefold enrichment after excluding low-complexity motifs. However, these 35 matrices represent only 28 distinct consensus sequences and approximately four sequence families, as motifs for paralogous factors are catalogued separately. We therefore select one representative per family rather than by choosing the highest-ranked motifs. Otherwise, the selection would be dominated by near-identical G-box variants, providing largely redundant tests of sequence attribution. We analyse HY5 (G-box, TGCCACGTGGCA, 15.4× enriched) and TCP22 (GTGGGCCCCAC, 9.0 ), from unrelated families, together with HHO3 (AAGATTCT, 1.00×) as a negative control: a matrix that is neither enriched nor depleted in accessible chromatin, and not a repeat. a motif that occurs at a similar frequency in peaks and background provides a cleaner negative control. a depleted motif is itself associated with accessibility, although in the opposite direction, so its expected contribution is not necessarily zero. HHO3 is suitable as a control for two additional reasons. At 8 bp, HHO3 is the shortest matrix retained. Its two terminal positions have information contents of only 0.56 and 0.53 bits, so most of the motif information is carried by the six internal bases. HHO3 also cannot reach the 10*^−^*^6^ scanning threshold. The highest possible matrix score corresponds to *p* = 3.6 10*^−^*^5^, which gives about a 7% chance of a background match within a window; for the other two motifs, this probability is 0.2%. These additional background matches tend to reduce the measured contribution, and we therefore treat the HHO3 result conservatively. Low-complexity matrices are excluded requiring that no single nucleotide constitute more than 60% of the consensus and that consensus contain at least three distinct nucleotides. The second criterion excludes dinucleotide repeats that can satisfy the first.

For each motif, we select 50 peak windows. Each window is eligible if it overlaps a called peak, contains exactly one occurrence of the motif entirely within the scored region, and shows no evidence of amplification or repeat artefacts; specifically we exclude windows in which a single base accounts for more than 10% of the observed reads. From the eligible windows, we select the 50 with the highest observed signal. When multiple copies of a motif are present in a window, they can be redundant or interact with each other, making it difficult to attribute the ISM contribution to an individual motif occurrence. Therefore, windows where the motifs appear multiple times are discarded from this analysis.

##### 4.5.6.3 Effect quantification

Contribution is compared within each window rather than across windows, because their magnitude can vary several-fold between loci. For each window, we compute the mean contribution across motif’s positions and compare it with the mean contribution across flanking positions in the same window. We exclude a 10 bp buffer on either side of the motif to avoid including attribution signal immediately adjacent to the motif in the flanking estimate. The per-window effect is

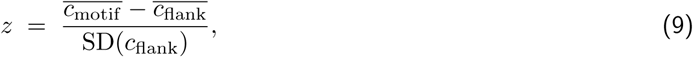

where 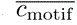 and 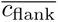 represent the mean contributions at motif and flanking positions, respectively, and SD(c_flank_) is the standard deviation of flanking contributions within the same window. We report the mean z-score across the 50 windows. *z* is dimensionless and this allows us to compare the count and profile heads, even though their raw contribution values have different scales: predicted log(1 + x) total for the count head and probability distribution for the profile head. We record the proportion of windows in which the motif’s mean exceeds that of flanking positions to measure how consistently the effects appear across loci.

Minus-strand motif occurrences must first be reoriented to match the reference motif direction. The per-base contributions are then averaged across windows. Position 0 therefore always refers to the first base of the motif. Without this reorientation, corresponding positions on the two strands would not be aligned, making it difficult to detect the position-specific patterns we are looking for within a motif.

Concentration within a motif is defined as the mean absolute contribution over the motif’s core positions divided by the mean absolute contribution over its two outermost positions. Cores are defined from the JASPAR matrix, with each core position required to carry at least 1.5 of the maximum 2.0 bits of information. For HY5, this corresponds to positions 4–9 of MA0551.2 (CACGTG, the G-box), and for TCP22, to positions 3–5 together with 7–8 of MA1288.2 (GGG and CC). Position 6 in TCP22 is excluded because the matrix carries only 0.136 bits at this position. No core is defined for HHO3. We use the absolute value of the terminal contribution in the denominator because the terminal mean can be close to zero or negative. a value close to zero makes the ratio unstable, while a negative value changes its sign.

We use a two-sided Wilcoxon signed-rank test for the paired motif-to-flank differences and calculate the pvalue using the normal approximation rather than the exact distribution (Supplementary Table S18). We specify this because, at *n* = 50, the two methods give p-values that differ by several orders of magnitude. The negative control also gives a significant p-value despite having a very small effect. With 50 paired windows, even a small but consistent difference can be statistically significant. We therefore consider the effect size and consistency across windows together with the p-value.

### 4.6 Transcription factor family binding

#### 4.6.1 Labels and chromosome split

We use the 568 DAP-seq and ampDAP-seq peak files released for the *Arabidopsis thaliana* cistrome [65], which resolves motifs and peaks for 529 TFs, several of them assayed under both protocols. Assayed TFs are grouped into 46 DNA-binding families. TAIR10 is tiled into non-overlapping 250 bp label windows. For each window and each family, a positive label is assigned if a family contains a 201 bp peak for which the window covers at least 70%, corresponding to at least 141 bp. Windows in which fewer than 90% of bases are A, C, G or T are excluded from scoring. Chromosomes 1 to 3 are assigned to training, chromosome 4 to hyperparameter and checkpoint selection, and chromosome 5 to final evaluation. The retained sets contain 293,591, 74,327, and 107,851 windows, respectively. The primary metric is macro average precision: average precision computed separately for each of the 46 families, then averaged across families without weighting. a window is a negative for every family that has no peak covering it, and the positive prevalence is only 0.047 averaged across families.

#### 4.6.2 Pre-trained checkpoints

The Botanic1-S arm fine-tunes the context-extended Botanic1-S backbone of Section 4.1.5, taken at the end of the final (131,072 bp) curriculum stage, that is at step 34,679 of that stage and 800B cumulative tokens, with 317,896,201 parameters. The Botanic1-M comparison fine-tunes the released Botanic1-M checkpoint at 314.6B tokens (687,526,665 parameters), pre-trained at 8,192 bp only.

#### 4.6.3 Fine-tuning

All gLMs are trained on 2,048 bp windows centred on the 250 bp labelled window. The model’s sequence embeddings are pooled over the central window and passed to a 46-output head. All weights are updated using asymmetric multi-label loss with positive focusing parameter 0, negative focusing parameter 2, and negative probability clip 0.05. We use reverse-complement augmentation with probability 0.5, reverse-complement logit averaging at evaluation, effective batch size 32, AdamW with weight decay 0.01 and maximum gradient norm 1.0, and no warmup. Foundation-model fine-tuning uses bfloat16. DeepCistrome receives 250 bp inputs and is trained under its optimised published procedure [66], with the same chromosome roles and windows as the foundation models.

Each gLM fine-tuning is tested on three different learning-rate settings across three seeds, resulting in nine runs per model. a setting fixes a backbone rate and a head rate together; the head rate is not screened independently of the backbone rate. The settings, as (backbone, head), are (3 10*^−^*^6^, 10*^−^*^4^), (10*^−^*^5^, 3 × 10*^−^*^4^) and (3 10*^−^*^5^, 10*^−^*^3^) for Botanic1-S and AgroNT; (10*^−^*^5^, 3 × 10*^−^*^4^), (3 10*^−^*^5^, 10*^−^*^3^) and (10*^−^*^4^, 3 × 10*^−^*^3^) for Botanic1-M; (3 10*^−^*^5^, 10*^−^*^4^), (10*^−^*^4^, 3 × 10*^−^*^4^) and (3 10*^−^*^4^, 10*^−^*^3^) for NTv3-652M; (3 10*^−^*^5^, 10*^−^*^3^), (10*^−^*^4^, 3 × 10*^−^*^3^) and (3 10*^−^*^4^, 10*^−^*^2^) for NTv3-106M; and (3 10*^−^*^7^, 3 × 10*^−^*^5^), (10*^−^*^6^, 10*^−^*^4^) and (3 10*^−^*^6^, 3 × 10*^−^*^4^) for PlantCAD2-L (Supplementary Figure S20). Some of these settings make training diverge, so a setting is eligible only when all three seeds converge during training. We evaluate performance on chromosome 4 once per epoch for early stopping: the learning rate is reduced by a factor of 0.3 after two evaluations without improvement, and training stops after nine evaluations without an improvement of at least 0.0005, with a cap of 18 epochs. We select the best setting with three finite seeds using mean chromosome 4 macro average precision. Means differing by less than 0.002 are resolved by lower seed standard deviation, then fewer median optimiser steps.

#### 4.6.4 Fine-tuning compute and label efficiency

For the compute efficiency analysis (Figure 5a), we plot the chromosome 4 average precision recorded for each epoch across the three seeds against a compute estimation as 6ND, where N is the model parameter count and D is the number of tokens processed. Each optimiser update contains 32 sequences of 2,048 tokens, and one epoch contains 9,175 updates. Parameter counts are 317.9M for Botanic1-S, 651.8M for NTv3-652M, 106.5M for NTv3-106M, 1B for AgroNT, and 1.39B for PlantCAD2-L. PlantCAD2-L has 694.4M trainable parameters, but its reverse-strand Mamba modules reuse the forward-strand input and output projections, so each token passes through those 692.5M shared weights twice and the compute estimate counts them twice. Training loss continues to decrease across all epochs (data not shown), but the evaluation average precision eventually plateaus and declines for all models, as shown in Figure 5a.

For the label-efficiency analysis (Supplementary Figure S21), we subsample 1%, 5%, 20%, and 100% of the unique training windows and train Botanic1-S, NTv3-652M, and NTv3-106M with their best learning rate setting, across three seeds and using the same early-stopping scheme as in Section 4.6.3 with a 30-epoch limit.

#### 4.6.5 Pre-training control

The random-initialisation control uses the same architecture, tokeniser, inputs, training windows, loss, augmentation, evaluation schedule, stopping rule, and three seeds as Botanic1-S. The pre-trained and random-initialisation models each receive their own three-setting learning-rate screen. The control’s settings are (3 10*^−^*^4^, 10*^−^*^4^), (10*^−^*^4^, 3 × 10*^−^*^5^) and (3 10*^−^*^5^, 10*^−^*^5^), this time choosing larger values for the backbone. Both are evaluated on chromosome 4 as shown in Figure 5b.

#### 4.6.6 Canonical 6-mer enrichment

For the canonical 6-mer enrichment analysis, we examine 5 TF families with known motifs: WRKY, bZIP, bHLH, MYB, and NAC. We select all positive windows (from the ground truth labelling) and order them by averaged model score across 3 seeds for each model and each TF family. The comparison is performed on the highest-scoring vs the lowest-scoring fifths of those windows annotated as binding. a 6-mer counts once when it or its reverse complement is present anywhere in a window. We retain all 6-mers present in at least 2% of the two fifths combined, and calculate log_2_ (+1) enrichment. The six most enriched Botanic1-S 6-mers are shown in Supplementary Figure S22.

### 4.7 Accessible chromatin region prediction

To test whether longer contexts capture informative long-range regulatory signal, we assess how the performance of Botanic1 on chromatin accessibility prediction varies with input context length.

#### 4.7.1 Task

We reproduce and extend the maize cell-type task of Figure 5G in [33]: a multi-label classification in which each ATAC-Seq-peak-centred genomic window is scored for accessibility in 92 maize cell types. Peak coordinates, labels, and the train/test split are retrieved from the HuggingFace release accompanying the paper [136]. All models are trained with binary cross-entropy against these 92 accessibility labels. We report the distribution of the AUPRC across cell types measured on the test set.

#### 4.7.2 Context window extension

The published dataset contains 600 bp windows centred on each ATAC-seq peak. To assess how context length affects prediction, we symmetrically widen the window around the same peak up to values ranging from 150 bp up to 128,000 bp, keeping the peak set, labels, and split fixed. Some samples in the train and test sets correspond to sequences close to chromosome extremities; for those rows, extending to longer context requires us to pad the sequences with N nucleotides. However, padding stays limited even at the longest contexts (Supplementary Table S10), with around 2% of sequences concerned at 128,000 bp context.

#### 4.7.3 Models

We compare four backbones: the small and medium PlantCAD2 models [33, 136], PlantCAD2-S and PlantCAD2-M, and two variants of Botanic1-S, one pre-trained at 8,192 bp and the other extended to 131,072 bp of genomic context (denoted Botanic1-S-8k and Botanic1-S-128k, respectively).

#### 4.7.4 Training strategies

We train for the task using four distinct recipes. The first, *Full-window LoRA fine-tuning*, is the closest to the fine-tuning strategy used in [33]. The second, *Central-window LoRA fine-tuning*, adjusts the pooling step to focus on the tokens at the central peak region. The third, *Frozen-backbone probing*, freezes the backbone and trains a simple head on hidden states mean-pooled at various central-window widths, and is also the most compute-efficient. The fourth, *Frozen-backbone probing with learned pooling*, replaces the fixed central window with per-position weights learned jointly with the head. Each recipe is described in more detail below.

##### Full-window LoRA fine-tuning

Each backbone is fine-tuned with LoRA [35] using the PEFT library [124], with LoRA rank = 8, α = 8, dropout = 0, targeting the *x* proj, in proj and out proj modules. Training uses the HuggingFace Trainer with a nominal learning rate of 10*^−^*^4^, an effective batch size of 128, BF16 precision, a linear learning-rate schedule with 50 warm-up steps, and one training epoch. Chromosomes 9 and 10 are held out as the test set; the remaining chromosomes are used for training. Hidden states from the last layer are averaged over the whole input window and passed to a two-layer ESM-style classifier head (dropout → dense → tanh → dropout → linear). The gradient-accumulation factor is set per (model, context) cell to keep the effective batch size at 128; due to a bug in our fine-tuning pipeline that was discovered a posteriori, the effective learning rate seen by the optimiser is the nominal learning rate multiplied by the gradient-accumulation factor for that cell. Refer to Supplementary Table S11 for the effective learning rates used across the experiments. With this strategy, all models are tested at input contexts 150, 300, 600, 1,600, 2,600, 4,600 and 8,000 bp; Botanic1-S-128k is tested at additional contexts of 16,000 and 32,000 bp.

##### Central-window LoRA fine-tuning

To separate the effect of a growing input window from the effect of diluting the peak signal at the pooling step, we repeat the same LoRA fine-tuning of Botanic1-S-128k with the same hyperparameters, but replace full-window pooling with mean pooling over a fixed 101 bp window centred on the ATAC-seq peak. We test context lengths of 150, 300, 600, 1,600, 2,600, 4,600, 8,000, 16,000, and 32,000 bp.

##### Frozen-backbone probing

For a lighter comparison, and to investigate the capability of non fine-tuned backbones to encode long-distance interactions, we freeze each backbone and train only a classifier on hidden states mean-pooled over a peak-centred window, sweeping across the following pool widths: 1, 11, 25, 51, 101, 151, 201, 251, 301, 351 and 401 bp and across input contexts (from 150 up to 128,000 bp). The experiments are run with a single linear head and with the same ESM-style head as strategy 1 above (*Full-window LoRA fine-tuning* ). We use the same train/test split as in [33].

##### Frozen-backbone probing with learned pooling weights

Instead of arbitrarily choosing a pooling window width or pooling weights by hand, we let the head learn the pooling weights along the input window itself. We introduce a trainable logit vector **a** ∈ ℝ*^L^* (one parameter per token position of the tokenised input of length L) and pool the frozen per-position hidden states as 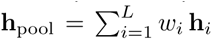, before passing **h**_pool_ to the classifier head. Two initialisations of **a** are reported: *uniform*, with all logits equal so that every position starts at exactly 1/L, and *random*, meant to introduce more randomness at initialisation with each position’s relative weight drawn independently from *U* [0.5, 1.5] before taking logarithms.

Pool logits **a** are optimised with a separate learning-rate multiplier relative to the head, which we sweep over 1×, 10× and 100×. Only 100×’s weight profile has converged by the end of the epoch (measured by the leastsquares slope over the final quarter of training). Every learned pooling experiment is run at three training seeds, controlling both the initialisation draw and the batch order. Another control is *reverse complement* learning, in which every input sequence is reverse-complemented while labels are left untouched.

All other hyperparameters (effective batch 128, BF16, one epoch) match strategy *Frozen-backbone probing* above. We report the frozen-backbone instantiation, with the same ESM-style head as strategy 1.

#### 4.7.5 Statistical annotations

For all strategies, we compare the per-cell-type AUPRC of each model between consecutive context lengths using two-sided Wilcoxon signed-rank tests paired by cell type on the 92 per-class AUPRC values, with Holm-Bonferroni correction applied within each panel for the violin figures and within each pooling-width column for the heatmap figures.

### 4.8 Sparse autoencoder training and feature scoring

#### 4.8.1 SAE training

We train a collection of BatchTopK SAEs [39]. This architecture consists of two dense layers (encoder and decoder) and a BatchTopK activation between them, trained for reconstruction of original signal (see Figure 8a). At inference time, this lets us decompose the model’s representation at any token into a combination of about *k* *features*, using only the encoder and the activation layers. We apply it to the residual stream after four different layers (normalised depths 0.25, 0.50, 0.67 and ~0.9) from each of Botanic1-S and Botanic1-M models, obtaining overall eight dictionaries. The number of features is taken to be eightfold the dimension of the model’s residual stream (therefore, into d_sae_ = 8,192 features for each of the dictionaries), and the number of active units per token we set to *k* = 64. The model itself is frozen throughout, so the dictionary is an instrument bolted onto the network rather than a probe fitted to a label (Figure 8a). For training we use 80,000 windows of size 8,191 over 320 plant species, the same corpus used to pre-train the Botanic1 models (Section 4.2).

Although several dictionaries train healthily and produce results, for simplicity we report the results of only one dictionary, for Botanic1-M at layer 35 of 52 (normalised depth 0.67). So, every feature index and every number in this section comes from one dictionary. Feature indices are specific to that dictionary and do not carry across depths or across models; moreover, their order carries no additional information, since the SAE training procedure is equivariant with respect to this order.

#### 4.8.2 Finding features for biological concepts

After training we use several annotated databases of genome data to find features related with different biological concepts. For transcription and splicing data we use data of four species, *Arabidopsis thaliana*, *Oryza sativa*, *Solanum lycopersicum* and *Zea mays*, carrying region, site and codon-position labels from the Ensembl Plants gene annotations [114] of the assemblies those genomes are read from (TAIR10, IRGSP-1.0, Zm-B73-REFERENCE-NAM-5.0 and SL3.0). To establish the relationship between genomic annotations and features we collect feature activations for 32,000 coordinate-anchored windows of 8,191 bp each.

We make a chromosome-disjoint split: *selection*, 16,510 windows used to score features against concept labels; *validation*, 8,277 windows to adjust thresholds and select the best features; *test*, 7,213 windows to report the scores (see Supplementary Table S16).

Basic extracted annotations include region (cds/utr/intron/exon/intergenic), site (donor/acceptor/tis/stop/tss/tts) and codon position. We score every feature against every label as a single-threshold classifier over a grid of thresholds and rank them by the MCC, see Equation (10) where TP, TN, FP, FN mean the numbers of true positives, true negatives, false positives, and false negatives respectively [137].

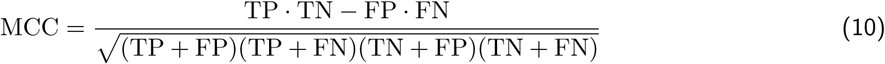

We find that this statistic remains informative even at low positive-label frequencies (e.g. 6 10*^−^*^4^ for a splice site). The features and thresholds reported in Figure 8b and Section 2.4 are found as giving the best MCC for corresponding labels on validation split. Additional results: MCC for per-species best features corresponding to genomic concepts in both reading directions, are given in Supplementary Table S17.

For cis-regulatory elements Figure 8d the relation of features to known cis-elements is established from both directions. First, from motif to feature, we find occurrences of JASPAR motifs in the corpus and collect features activating exclusively near these sites. Then for found features we take the top-32 activations across the corpus and construct their own consensus-motifs.

#### 4.8.3 Two-feature tree for alternative splice-site usage

We extract alternative splicing events from PastDb data [79] based on RNA-seq reads of *A. thaliana*. We keep only sites having at least 10 reads. We define PSU for a site (donor) as a fraction of reads using this site to the whole number of this gene’s reads. From it we derive the concepts of intermediate-usage and constitutive donors: PSU between 20% and 80% and over 95% respectively. From the total of 26,603 donor sites only 885 fall into the intermediate-usage class, so we sample exactly the same amount of constitutive donors and add the same amount of decoys: GT dinucleotides from the genes. The two-level tree shape is pre-defined; the features and thresholds are chosen to maximise the balanced accuracy. For this experiment we also use a validation-test split by chromosomes.

Balanced accuracy, the confusion matrix and the thresholds are all computed on these 2,655 sites. For Figure 8k we use a wider set of donors, to cover the whole range of PSU, which is divided into twelve bins of up to 250 donors (some bins have fewer). In total this gives 3,254 donors scored against the same 885 decoys. The added donors are not used in feature selection and threshold fit, only the Spearman correlations between feature activation and PSU are computed over all 3,254.

### 4.9 Zero-shot variant effect prediction on population variants

#### 4.9.1 Variant panels

In Section 2.5.1, we reproduce Figure 7 of Mendoza-Revilla et al. [31], originally conducted on the *Arabidopsis thaliana* variants extracted from the 1001 Genomes Project [138]. For every species, we keep biallelic SNPs and filter on call rate, the fraction of accessions with a called genotype at the site. The threshold is given per species below.

##### 4.9.1.1 Arabidopsis thaliana

Variants are extracted from the 1001 Genomes GMI-MPI v3.1 release (1,135 accessions) [138]. We find that the number of SNPs and call rate filter reported by Mendoza-Revilla et al. [31] are inconsistent: the authors report 6,494,574 SNPs by applying a call rate filter of 95%, yet we only obtain 4,135,872 SNPs by applying the same filter. In contrast, we reproduce the authors’ count under their 99% threshold to within one SNP (286,671 against their reported 286,672). We thus decrease the threshold to 88.37% to obtain a number of biallelic SNPs (6,491,238) close to what was reported for the original figure; this is the set that we score and that we report in our results. This choice has little effect on the missense/synonymous ratios by MAF class (Supplementary Figure S34).

##### 4.9.1.2 Oryza sativa

Like in Mendoza-Revilla et al. [31], the RiceVarMap2.0 panel (4,726 accessions) [139] is used to extract SNP variants. Following the original study, we keep biallelic SNPs with a call rate of at least 99%, leaving 6,494,573 SNPs. While Mendoza-Revilla et al. [31] report that they evaluated a 50% subsample of the SNPs passing this filter (3,128,064), we score all 6,494,573 SNPs. This count differs by exactly one variant from the *A. thaliana* SNP count reported by Mendoza-Revilla et al. [31] (6,494,574) and that we cannot reproduce under their stated filter. This coincidence is consistent with the two species’ counts having been swapped in the original report, although we cannot confirm it.

##### 4.9.1.3 Cicer arietinum

We use the CicerSeq cultivated panel (3,171 accessions) [140, 141], published as per-chromosome HapMap tables, which we convert to VCF against the CDC Frontier v1 assembly (RefSeq GCF 000331145.1): 167 of 2,470,880 records (0.007%) are dropped due to a mismatch in the reference allele. We also drop 52,014 (2.11%) SNPs on unanchored scaffolds, leaving 2,418,699 SNPs. Of these, 880,345 pass a 95% call rate threshold. Unlike RiceVarMap2.0, this panel is not imputed, so the 99% threshold used for rice would retain only 56,826 SNPs (2.3%); we therefore keep a lower threshold.

Some models cannot score all variants; this information is reported in the figures where relevant (Section 4.9).

#### 4.9.2 Model coverage across sizes

The supporting panels for Section 2.5.1 (Supplementary Figure S34 and Supplementary Figure S37) are computed on the same 6,491,238-variant *A. thaliana* set as the figures from the main text.

Every panel of this benchmark reports Botanic1-S, Botanic1-M and Botanic1-L only. The Botanic1-XL results here use an earlier checkpoint of the same 3.2B configuration, trained on a longer token horizon whose learning rate had therefore not decayed at the 314.6 billion token reference point, so those results are not comparable with the other three sizes and are not drawn.

#### 4.9.3 Scoring

To score SNPs, we employ both the LLR (defined below in Section 4.9.4) and embedding-based scores (Section 4.9.5). LLR scores and embedding-based scores are computed on a 6,001 bp window centred on the variant.

#### 4.9.4 LLR score

SNPs are scored using the LLR, the masked-marginal approach introduced for protein language models [19, 20] and carried over to genomic language models in [83, 142].

For Masked Language Models, it is:

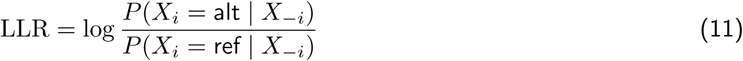

where X*_i_* is the nucleotide at the variant position, *X_−i_* the surrounding sequence, and alt and ref the alternate and reference alleles, respectively. Variants are ranked by LLR, so that a higher value indicates a more disruptive substitution. For multiple-bases tokenisers, the reported LLR is a ratio between two 6-mer token logits, each reflecting the entire alternate or reference 6-mers rather than the substituted base.

Autoregressive or causal models cannot output *P*(*X_i_|X_−i_*): they predict the sequence from left to right, so each position can only attend to the preceding context. Brixi et al. [24] define the delta-likelihood score as the difference between two sequences’ likelihood, which in this case differ in only one position:

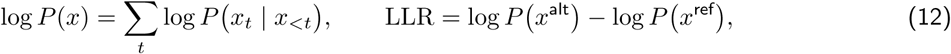

where x^ref^ and x^alt^ are the two versions of the window and LLR keeps the orientation of Equation (11), so the same ranking by −LLR applies. The log-likelihood is summed over positions, not averaged, for precision concerns.

#### 4.9.5 Embeddings-based score

The embedding-based score is taken as the dot product between the embeddings of the reference and alternative sequences. It is extracted at four layers per model, matched by relative depth (Supplementary Figure S37).

#### 4.9.6 PhyloP score

PhyloP [143] scores are read from the per-species bedGraph tracks published by PlantRegMap [144] at each variant’s coordinate.

#### 4.9.7 Uncertainty

Neighbouring SNPs are correlated by linkage disequilibrium. We thus compute confidence intervals using blocks of SNPs instead of individual SNPs. To choose a suitable block size, we consider the reported LD decay scale in *A. thaliana* [145] (~10 kbp) which we multiply by 100. The blocks are thus obtained by cutting the genome into 1 Mbp blocks (121 blocks for *A. thaliana*). The confidence intervals are computed by resampling blocks via a paired leave-one-block-out jackknife. We confirm that this confidence interval estimation is correct by performing bootstrap on the same blocks: a 2,000-replicate block bootstrap agrees with the jackknife to within 7 10*^−^*^4^ on every interval.

### 4.10 Zero-shot causal variant prioritisation benchmark

#### 4.10.1 Task and ground truth

Zero-shot evaluation of mutations has been used since the first biological language models, starting with protein sequences [19, 20] and later expanded to gLMs, theoretically able to score regulatory as well as coding variants. New genomic language model publications usually include a benchmark of zero-shot mutation scoring, using as a ground truth ranking either the annotations and algorithmic classification to categorise the variants into consequence classes much like Ensembl VEP [146], or using the population-level frequency of each variant when available. Both act as a proxy for functional impact, and positive correlation with zero-shot scores is interpreted as evidence that a model has learned functional constraint [142]. As a proof of concept, Zhai et al. [34] then showed that in a real use case, the zero-shot score of their model was able to single out a well-studied causal sweet corn mutation in the Su1 locus, where the GWAS p-value alone could not. This use-case recreates a common situation in genetics, where a locus is identified through a GWAS or BSA study, but the exact mechanism-carrying causal variant cannot be pinpointed because of linkage disequilibrium or association power.

This example illustrates the utility of the zero-shot score over bioinformatics baselines, but to our knowledge no benchmark with any significant scope has been developed. To address this gap, we design a benchmark of 545 loci across 14 species, each centred on an experimentally validated causal variant.

#### 4.10.2 Candidate loci and coordinate quality control

This new dataset is constructed starting with agent-driven literature search first, and pairs each variant (defined by its genomic coordinates, reference and alternate alleles) with a publication reporting an experimental characterisation of its impact on a plant phenotype or molecular function. We restrict our analysis to direct mutagenesis, transgenic complementation or reporter assays, biochemical characterisation of the substitution, or genetic mapping that resolves the exact allele and includes functional follow-up. We exclude tagging SNPs (variants reported only through linkage disequilibrium), structural alleles, and variants without a verifiable reference coordinate. Positive labels are experimentally or genetically supported causal nucleotide, rather than the most significant marker in an association study or allele frequency. Details and the complete eligibility rubric and literature-curation protocol are given in Section 4.11.

Each causal variant is placed at the centre of a 100 kbp locus and complemented with all documented SNPs in this region, retrieved from public population-variation VCFs for the relevant species, thus constituting the set of candidate variants. We use documented SNPs as we assume that most will be largely neutral, but we acknowledge the positive-unlabelled nature of the task (Section 4.10.4). If the original publication uses a different reference genome, the causal variant coordinates are lifted, as are the panel variants if necessary, to the most recent assembly.

#### 4.10.3 Ranking candidate SNPs

Every SNP in the set of candidate variants is scored by each method, using either the LLR formula for zero-shot scoring, Ensembl VEP categorisation, or MSA-derived conservation scores.

##### 4.10.3.1 gLM scoring

Each candidate SNP is scored using the LLR already defined in Section 4.9.4, and candidates are ranked by its absolute value LLR because it does not require determining which is the reference and which is the alternate allele, which is not obvious when the allele in the reference genome is not frequent in the population. LLR is computed on a window of length W centred on the candidate variant, with W = min(4,096, trained context): 4,096 bp for the Botanic models, PlantCAD2-L, AgroNT and Evo 2, and 512 bp for GPN and PlantCaduceus-l32. Only biallelic SNPs are scored. When a variant lies within W/2 of a chromosome end, the window is truncated with the variant kept as close to centre as possible.

Using the signed LLR requires choosing a reference and alternate sequence, something that is not always straightforward. For example, we might decide that the reference should correspond to either the ancestral allele, the major allele, or the reference genome’s nucleotide, and all three might not agree. Using the absolute LLR in this benchmark asks whether the causal variant is functionally constrained at this position compared to background SNPs in the locus. Here we do not pretend to say that either ref or alt is more deleterious, something that is not needed for this benchmark.

The causal variant discovery family of *S*_bal_ (Section 4.3.2) is a lighter and faster protocol used for evaluation purposes. It differs from the full benchmark by scoring only 29 studies from four species (13 *Arabidopsis thaliana*, 11 *Oryza sativa*, 4 *Triticum aestivum*, 1 *Sorghum bicolor* ), chosen to balance variant classes. Another difference is that in this benchmark, the causal SNP is ranked against 100 background SNPs from its locus instead of all published SNPs: the 100 background SNP are chosen as the 50 closest in allele frequency and the 50 nearest by position when the population VCF carries allele frequencies, otherwise the 100 nearest by position. Finally in this benchmark candidates are ranked by the signed LLR of Equation (11), with the reference genome allele as reference, instead of LLR due to legacy choices. Therefore the numbers from this benchmark are not directly comparable to Table 3.

##### 4.10.3.2 The post-trained Botanic1-S model

Botanic1-S (post-trained) is an experimental model starting from the pre-trained 318M Botanic1-S checkpoint before context extension, and further trained to predict genomic annotations on *A. thaliana* only. It adds a rank-64 low-rank adapter [35] on the Mamba2 input and output projections for 3,000 steps, supervised at masked positions by four per-position classifiers. Instead of using the reference genome only for the MLM loss, we also use a soft cross-entropy against the full distribution of alleles observed in the 1,135 accessions of the 1001 Genomes project [138].

Let *M* be the masked positions of a batch, and *M_h_ ⊆ M* those where classifier *h* ᗴ 1, . . . , 4 has a label. The objective is

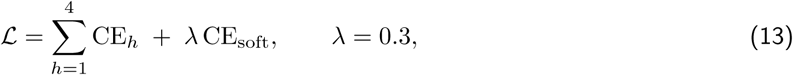

where CE*_h_*is the class-balanced cross-entropy of classifier *h* against its label over M*_h_*, and CE_soft_ is the crossentropy of the frozen language-modelling head against π*_i_*, the empirical distribution of bases observed across the called accessions at position i, weighted towards positions of low across-accession entropy.

The four classifiers use the following labels:

- minor-allele-frequency class from the 1001 Genomes [138] (monomorphic, ≥ 5%, and < 5%).
- conserved-noncoding-sequence clade depth from Conservatory [147], graded on four classes from unconserved to deepest clade.
- Ancestral/derived allele class using another Brassicaceae species as an outgroup, *Capsella grandiflora*, at all orthologous positions (about 16% of segregating sites). Four classes combine ancestral status and allele frequency: ancestral fixed, ancestral rare, derived common and derived rare.
- base-pair-level TAIR annotation [73]: intergenic, 5*^′^* UTR, 3*^′^* UTR, coding exon, intron, splice donor, splice acceptor, TIS, TTS, intergenic.

Each prediction contributes a cross-entropy loss for each position with a valid label (for the derived-allele class, a missing position label means that the corresponding loss is excluded, rather than assuming a default value).

The classifiers are then dropped, the adapter is merged into the backbone, and the model is scored exactly like every other Botanic model.

##### 4.10.3.3 Baseline scoring

Three non-gLM baselines are also used to rank all SNPs in the window.

Ensembl VEP [146] returns a categorical consequence per transcript. We take the most severe consequence across all transcripts of a variant and score it by its rank in the Sequence Ontology hierarchy, from transcript ablation down to intergenic variant. There are 32 different consequence terms, but only 22 can apply to SNPs, and less than 20 are observed in practice for most loci.

PhyloP [143] and PhastCons [148] are read from the per-species bedGraph tracks published by PlantRegMap [144] at each variant’s coordinate, under 08-download/ Species /sequence conservation/ Code PhyloP and the matching PhastCons path.

Negation is applied if needed so that all signs follow the same convention: a higher value means a more disruptive variant.

#### 4.10.4 Assumptions and limits

In this benchmark we only score SNPs and abstain from scoring other potential causal mutation like indels and structural variants. We also assume that the published, validated causal SNP should rank at the top of the locus, or in other words, that every other SNP in the region is a negative. Another particularity of this benchmark comes from comparing naturally occurring background variants with potentially engineered causal variants. None of the gLMs in our comparisons are trained on pangenomes or any VCF-derived population data, so all variants are theoretically equal regardless of their origin.

#### 4.10.5 Aggregated statistics

For study *s* and model *m* we define the top fraction of causal variants *q_s,m_* as *q_s,m_* = *r_s,m_*/N*_s_*, where *r_s,m_* is the rank of the causal variant from study *s* according to the model *m* and N*_s_* is the number of candidate variants for study s. Recall at search fraction α for model *m* is the proportion of studies whose causal variant is inside a shortlist of the top K*_s_*(α) = αN*_s_* candidates. We compute it tie-aware, as the expectation over a uniformly random order inside the block of candidates sharing the causal variant’s exact score:

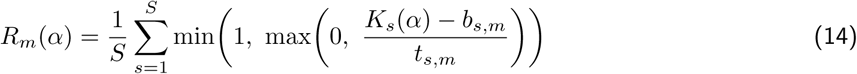

where *b_s,m_* is the number of candidates with scores strictly better than the causal variant and *t_s,m_* the number of candidates with the same score. In the absence of ties, R*_m_*(α) = 1[q*_s,m_* α].

When variants are tied, no further sorting is applied, and with the tie-aware recall definition above (Equation (14)), each study where the causal variant is in a tie contributes to R*_m_*(α) with the share of the tied block that can fit into the corresponding shortlist size K*_s_*(α). Some scores (especially discrete ones like Ensembl VEP) yield many ties, which is why we also report the median percentage of candidate SNPs assigned the exact same score as the causal variant across studies (Table 3 and Supplementary Table S14).

The macro recall AUC for a given model is then defined as the area under R*_m_*(α) curve, and in the absence of ties, it equals the mean of 1 − *q_s,m_* across studies:

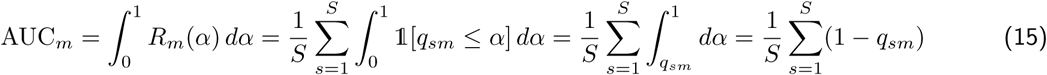

Three metrics are reported throughout the results in Section 2.5.1.2: recall at α = 0.01, recall at α = 0.001 and the recall AUC. One caveat of AUC is that it also encompasses what happens at high α values, which is of little practical interest (whether the variant is among the top 50% or among the top 80% does not matter much when one is trying to prioritise variants for further testing). This is why we prefer to focus on recall at α = 0.01 and at α = 0.001 when possible.

For several species in the benchmark, no PlantRegMap conservation track exists, and the 74 corresponding studies cannot be scored with PhyloP and PhastCons. Table 3 reports each model’s recall scores and statistics on the set of studies it successfully scores (up to 545 studies), while Supplementary Table S14 reports the results on the 471 studies that are scored by all models.

Pairwise comparisons between models are only performed on set of studies that both models successfully score. We report the paired difference in recall against Botanic1-XL at α = 0.01 and α = 0.001, together with a 95% confidence bootstrap interval as described below.

Each bootstrap replicate is constructed by drawing papers with replacement: at each draw we pick a paper and add all of its studies to the replicate. We resample papers rather than studies because two studies from the same paper are rarely statistically independent. On every replicate, recalls are computed from the same list of resampled studies for both models and the difference is recorded.

### 4.11 Causal variant dataset construction

We construct the dataset from the ground up by doing a literature search of experimentally validated causal variants, reviewing and filtering abstracts and full text papers, extracting and validating genomic coordinates against a reference genome, and homogenising the results in a common format, all performed by AI agents. Starting from a broad literature search, we apply progressively stricter filters with progressively more capable (and more expensive) agents to build the dataset. Supplementary Table S12 gives an overview of the pipeline at different stages, which we describe now in more details. We also report some implementation details in Supplementary Table S13.

#### 4.11.1 Search and abstract screening

Our search for papers is a two-stage process: first retrieve titles and abstracts from PubMed, OpenAlex and the bioRxiv archive, and then apply regex filters on the resulting list.

PubMed is queried with a Boolean expression joining two groups of MeSH terms: one group for focusing on relevant species (Plants and 36 named plant taxa), and the second for the object of the study, a causal mutation (Genetic Variation, Quantitative Trait Loci, Genome-Wide Association Study, Genes, Plant, Chromosome Mapping, Cloning, Molecular, Alleles, Mutation and Quantitative Trait, Heritable). We search OpenAlex with text queries plant GWAS causal SNP. bioRxiv .meca archives from January 2020 to January 2026 are downloaded and filtered locally by regex filter as described next.

The resulting list of documents is then filtered with regular expressions.Titles and abstracts are screened for presence of a plant expression (at least one of 603 scientific, abbreviated and common names derived from the NCBI plant genome list) and a variant expression (16 strings, including SNP, mutation, variant, allele, GWAS and QTL). Selected records are deduplicated by DOI and by PMID, in case of duplication the PubMed version is preferred.

In a second step, we select only documents with abstracts and screen them with a lightweight agent (claudehaiku-4-5-20251001) to scan for a plant species, a specific causal gene or variant, and evidence of functional validation. The agent’s output is a valid JSON formatted response of at most 256 tokens. We do not require exact coordinates or a specific evidence type at this screening phase, as these are checked later. Out of 36,256 documents with a screenable abstract, this screen keeps 4,543 (Supplementary Table S12).

#### 4.11.2 Eligibility checks

For every paper that passes the title/abstract screen, we then try to retrieve the full text if the original search does not already return it. We query PMC BioC for records with a PMID, bioRxiv JATS for 10.1101 DOIs, and the open-access URL returned by OpenAlex from the DOI. If the full text cannot be found by any means, the following steps use only the abstract.

From the full text, documents are assessed once with claude-sonnet-4-6 which is asked to parse the paper and summarise the information in another structured record containing the species, assembly, gene, variant description and class, coordinates, alleles, nature of the experimental evidence, and a confidence level. An optional field is reserved for additional information such as association results (GWAS, BSA or QTL mapping), reference genotype panels or phenotypic data. Finally, we ask the agent to fill the binary eligible flag, which allows the study to continue to the tool-assisted verification. The eligibility criterion is based on four conditions:

- The organism is a plant species (Viridiplantae)
- The variant is a SNP in the nuclear genome
- The paper provides functional validation for this exact SNP. Accepted evidence includes transgenic complementation or reporter assays, site-directed mutagenesis, saturation mutagenesis, biochemical assays, and BSA or GWAS fine-mapping followed by functional validation.
- The variant is located precisely in the reference assembly

We reject tagging SNPs, structural alleles and gene-level perturbation if they do not also include specific evidence for a causal SNP. Eligible documents are passed on to the coordinate verification step.

#### 4.11.3 Coordinate verification and genomic quality control

Rather than relying on text parsing alone, we also require the documented variant to be checked with bioinformatics tool calling. We give claude-opus-4-6 the structured JSON output from the last stage and the following list of tools:

- ensembl gene lookup to retrieve Ensembl gene annotations;
- ensembl xref search to map gene symbols and external identifiers to Ensembl stable identifiers;
- ncbi gene lookup as a fallback when a species or gene cannot be resolved through Ensembl;
- cds position lookup to convert reported amino-acid substitutions into candidate genomic substitutions;
- ensembl sequence lookup to retrieve genomic sequence and check the reported reference allele;
- lookup known panels to retrieve the registered target assembly, reference FASTA, population VCFs and required liftover chains;
- register species to record the normalised Ensembl species identifier used by the subsequent reference check; and
- web search as a fallback for identifiers or resources that cannot be resolved through any structured database.

Augmented by bioinformatics tools, this is where the agent is able to map the paper’s information (e.g. amino-acid coordinates, gene relative position) to a verified genomic coordinate in a reference assembly. When the causal variant is a documented SNP, the alternative allele is also verified. This is not necessarily the case though, for example mutagenesis studies produce variants that are not in reference panels.

The result of this stage is the final structured output with the species, target assembly, precise genomic coordinates, forward-strand reference and alternative alleles, target FASTA, population VCF and any required liftover steps.

#### 4.11.4 Final cohort

Finally, all eligible studies with a complete final record are checked programmatically to ensure that all reference alleles at the causal SNP position resolve to the correct content in the correct assembly, using Ensembl. If the reported allele matches the complement in Ensembl, both alleles are converted to the forward genomic strand.

The pipeline produces 591 verified unique studies, each associated with a causal SNP. Some studies refer to the same variant, after removing duplicates the cohort consists of 572 unique variants.

Finally, as a last verification step we use Codex with a two-pass filter: GPT Luna 5.6 on all studies and then GPT Sol 5.6 on rejected papers, both enabled with their full suite of tools (including the full text source, web search and bioinformatics tools). 545 studies are selected with validated causal variants. The full audit, including the first- and second-pass decisions and a summary for each variant, is available on request, and a future manuscript will present the complete dataset. We give below an example from this audit:

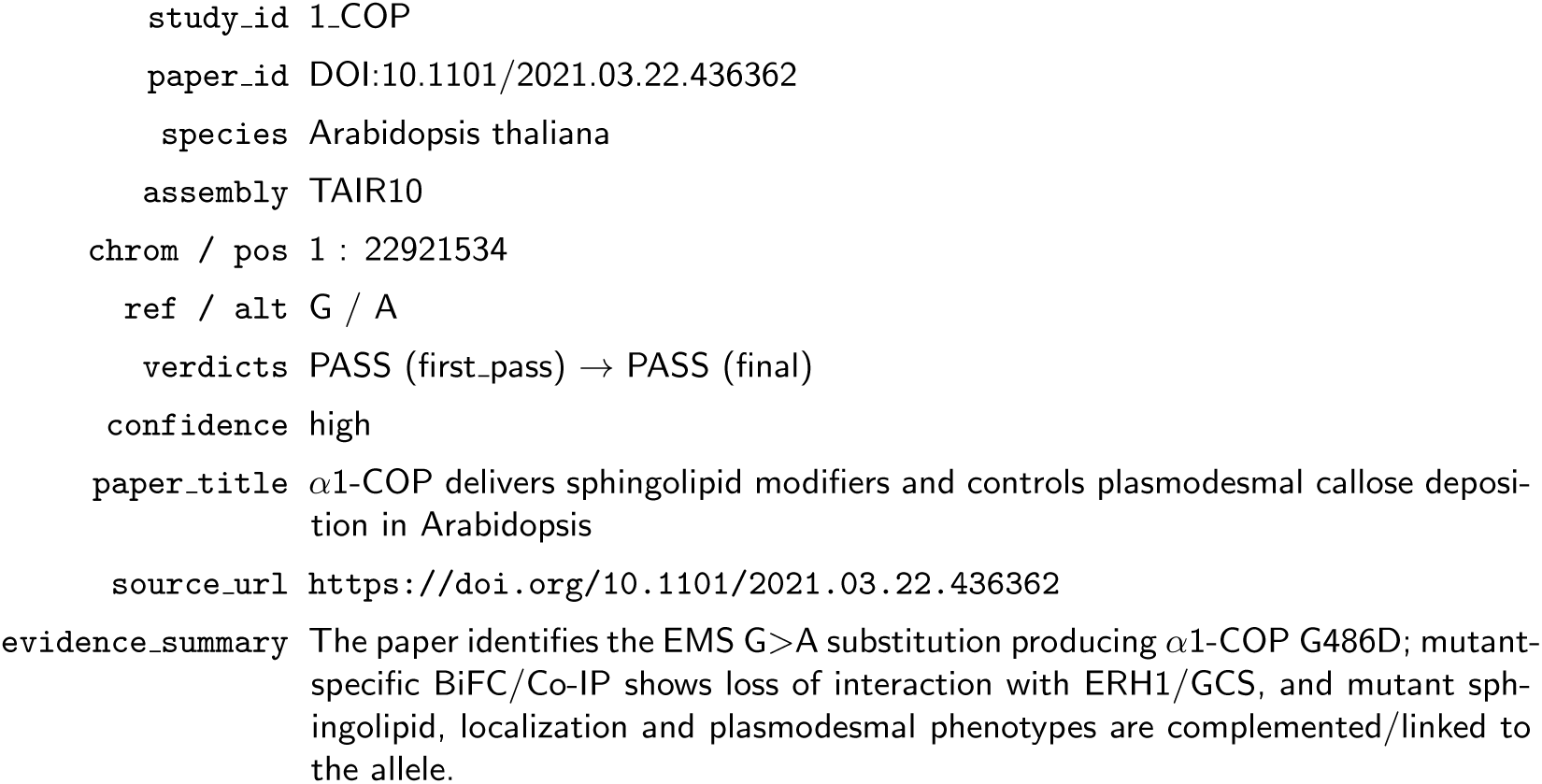

The final benchmark dataset is therefore made up of 545 unique causal SNPs across 14 plant species (Supplementary Table S12).

No stratification or post-selection is applied, the final dataset reflects the distribution of published and verifiable plant functional variants rather than a curated and balanced set.

### 4.12 Agentic causal variant discovery case study

#### 4.12.1 Study and candidate set

The case study of Section 2.5.2 is a retrospective reanalysis of a published melon (*Cucumis melo*) sex determination study [52], in which a G124R substitution in the ethylene signalling gene <u>CmEIN3</u>, flipping flowers from female to hermaphrodite, was experimentally established as causal. Starting from the raw sequencing data of the two segregating populations, bulk segregant analysis yields 47,492 candidate variants spread over the twelve melon chromosomes. The segregation signal, summarised by the ΔSNP index over 1 Mbp windows, localises the trait to chromosome 2 and narrows the candidates to 3,061 variants. This chromosome 2 candidate list, with each variant’s reference allele, alternative allele and sequencing metadata, is the input we provide to the agent across all experimental conditions.

Restricting the variant list, to chromosome 2 variants is a substantial simplification of the problem. In realworld settings, the starting point would be the full set of variants, across the entire genome. In that case, neither bioinformatics tools nor Botanic1 can recover the causal variant on their own. However, when combined, they can.

#### 4.12.2 Agent configuration

We use Gemma 4 E4B [53], instruction-tuned, run in bf16 and served with Ollama on a single NVIDIA L4 GPU node, so the whole loop runs on self-hosted infrastructure. The agent orchestrates its tool use freely. The prompt is identical across conditions, sampling uses the recommended temperature of 1.0 with thinking enabled

The agent is given two turns in the <u>No tools</u> and <u>Botanic1</u> conditions and four turns in the <u>Conventional</u> <u>bioinformatics tools</u> condition (see Section 4.12.3). In the <u>Conventional bioinformatics tool</u>, the follow-up questions (turn 2 to 4) are fixed messages bioinformatician experts could send to further guide the agents in the completion of their work. Namely: apply read-depth filters, double-check tool invocation, look at programs documentation, verify intermediate files before answering, etc. We report agent performance after the first message and after the follow ups, as a way to see what it can achieve on its own and under bioinformatician guidance.

Each condition is repeated 20 times to quantify robustness to sampling stochasticity.

#### 4.12.3 Three tested conditions

The agent is tested under three different conditions, which only differ in the sets of tools the agent can use to solve the task.

**Condition 1: No tools.** The agent receives only the candidate file and ranks the variants using its own internal knowledge.

**Condition 2: Conventional bioinformatics tools.** The agent receives a sandbox in which it can store intermediate results. a standard genomic toolset is installed in this sandbox (e.g. bcftools, samtools), together with Ensembl VEP, and the agent can retrieve publicly available reference files (genomes and annotations) if needed. In practice, the agent applies quality control (keeping biallelic SNPs and filtering low-confidence calls), computes allele frequency in between the mutant and wild-type bulks, checks variant functional impact using Ensemble VEP, ranks the variants, and annotates the SNPs that pass quality control for predicted consequence.

**Condition 3:** Botanic1. The agent receives the candidate file and a skill describing how to use Botanic1-L on a genomic sequence and specifically its log-likelihood ratio to score variants Section 4.10. No other tool is available in this condition. The agent chooses to discard the 567 candidates that are not single nucleotide substitutions (indels and rearrangements) and scores the remaining 2,494.

#### 4.12.4 Performance metrics

We assess the performance of the agent on two axes: whether it can identify the true variant, and whether it follows the rules and only ranks the variants we provide. For each of the 20 repetitions, we retrieve the agent’s final ranking and extract the rank of the experimentally validated (true) SNP. We then compute Recall@1 (the fraction of repetitions where this rank is 1), Recall@10 (rank ≤ 10), and Recall@20 (rank ≤ 20). We also count how often the output contains fabricated variants that are not in the list of candidates it is provided with. Supplementary Table S15 summarises those metrics.

## Data and model availability

All artefacts are released for research use under the Living Models research licence and are accessible from https://huggingface.co/spaces/living-models/botanic1-report.

- **Models.** The four Botanic1 checkpoints are available in the Hugging Face collection https://huggingface.co/collections/living-models/botanic1: https://huggingface.co/living-models/Botanic1-S, https://huggingface.co/living-models/Botanic1-M, https://huggingface.co/living-models/Botanic1-L and https://huggingface.co/living-models/Botanic1-XL.
- **Pre-training data.** The 8 kbp corpus that was used to train the four models, the one selected by the ablations of Section 2.2.1 (326 species, 7,207,506 windows, 59 Gbp; Supplementary Table S20), is released with its sequence case, augmentation margins, train/test split, source manifests and assembly metadata at https://huggingface.co/datasets/living-models/Botanic1-pretraining.
- **Sparse autoencoder.** The BatchTopK dictionary analysed in Section 4.8, trained on the residual stream of Botanic1-M at layer 35 (8,192 features, *k* = 64), is also released at https://huggingface.co/living-models/Botanic1-M-sae-k64-codebook8192.
- **Causal-variant benchmark.** Finally, the full curation audit of the 545 causal variants, including the first- and second-pass decisions and a summary for each variant (Section 4.10), is available on request; a future manuscript will present the complete dataset.

## Acknowledgements

We thank Benjamin Trom for his help in improving the pre-training stack and in using coding agents efficiently, and Adnane Boualem for providing the data and reviewing the melon use case.

## Author contributions

A.B., V.C., J.O.d.T. and A.R. contributed equally to this work as co-first authors; T.J., G.K. and Z.S. contributed equally as second authors.

**Conceptualisation, supervision and writing.** A.B., V.C., J.O.d.T. and A.R. conceived the study, supervised the research and led the writing of the manuscript (original draft; review and editing). V.C. and G.A. supervised the ATAC-seq experiments, and J.O.d.T. supervised the long-context extension work.

**Methodology, software and investigation.** J.O.d.T. contributed the initial pre-training stack. V.C., A.R. and J.O.d.T. were in charge of the pre-training experiments and managed the associated infrastructure (resources). A.R. led the interpretability experiments, with the help of V.C. and A.B. G.K. carried out the long-context extension and the associated synthetic-data evaluation (validation). T.J. and L.S. developed the agentic use-case illustration with Gemma, supervised by V.C. and J.O.d.T.

**Data curation.** V.C. built the data pipeline and performed the data ablation experiments with support from G.A., J.O.d.T., A.R. and L.S. G.A. developed the bioinformatics pipeline for ATAC-seq preprocessing.

**Formal analysis and validation.** A.B. was in charge of the long-context biological evaluation. Z.S. performed the ATAC-seq experiments under the supervision of V.C. and G.A.

**Funding acquisition and project administration.** L.S. and C.V. sponsored and enabled the work.

## A Supplementary Material

**Supplementary Figure S1.**
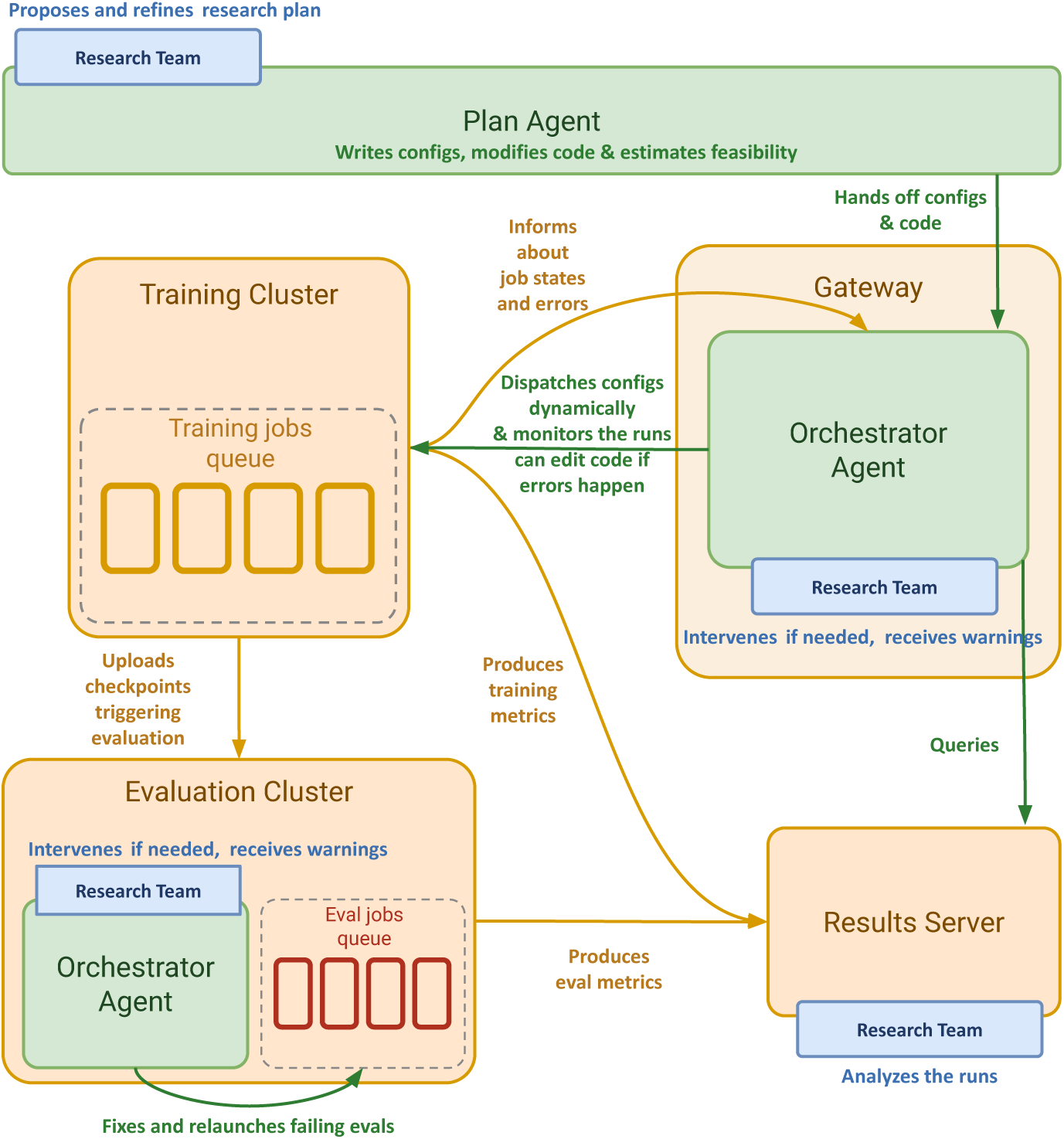
Agentic orchestration of the experimentation loop. A plan agent turns a research plan proposed by researchers into training configurations and potentially code changes and estimates their feasibility in a given timeframe, then hands them to an orchestrator agent on a gateway node, which dispatches jobs to the training-cluster queue, monitors the runs, and edits code when errors occur. Training runs produce checkpoints on disks, which in turn trigger evaluation of those checkpoints on a separate cluster, where a second more lightweight orchestrator agent potentially fixes and relaunches failing evaluations. Training and evaluation metrics are written to a results server that the agents query and that researchers analyse in parallel. The researchers intervene at each stage as needed and receive warnings from both orchestrators.

**Supplementary Figure S2.**
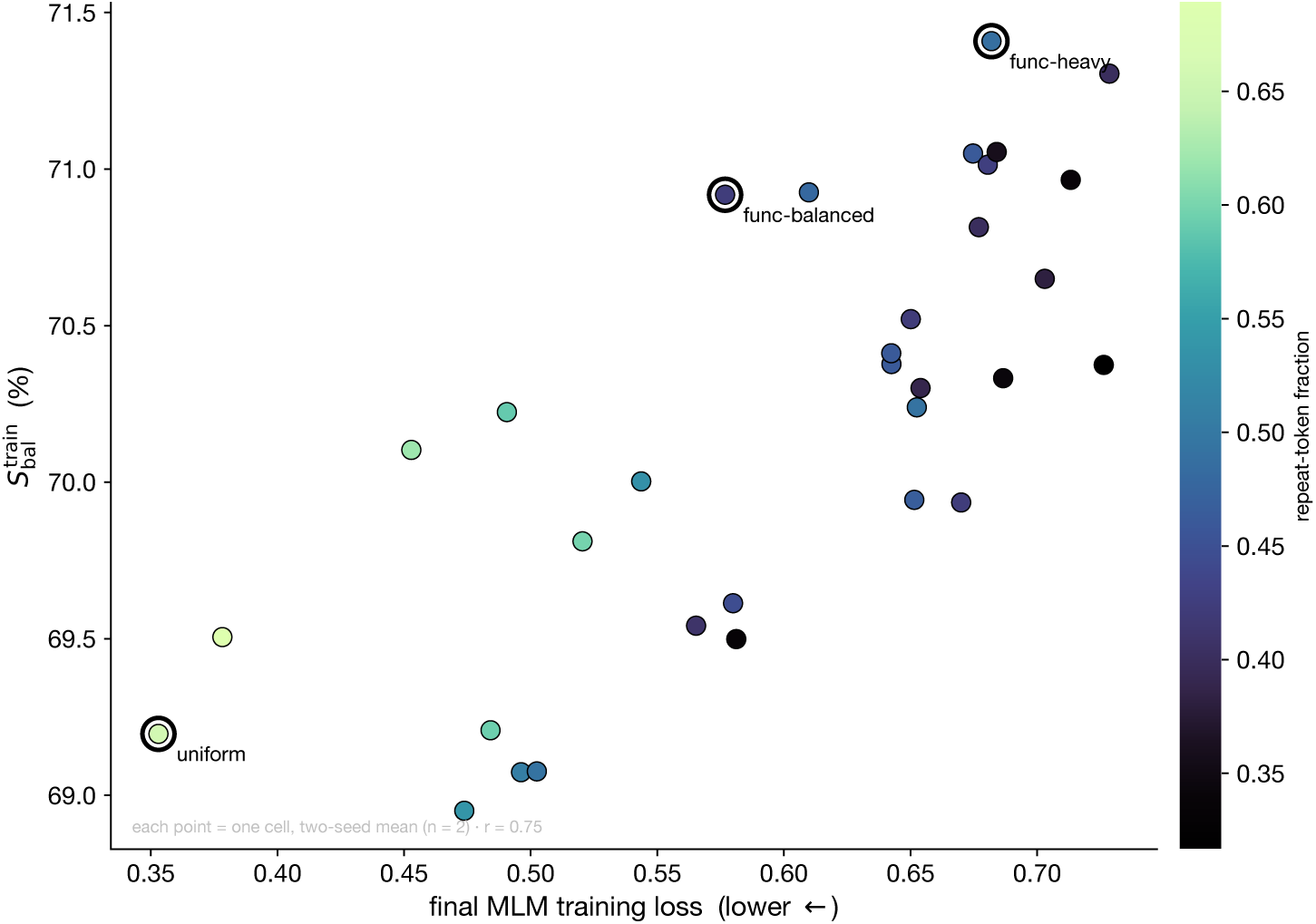
Training loss, downstream score and token repeat fraction. Each point is one data ablation experiment (averaged across the two seeds), on the x-axis we plot the final MLM training loss against its matching 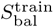 on the y-axis. The colour is the fraction of training tokens on annotated repeats. Circled points correspond to the window sampling strategy sweep results on the crop-centric species pool (see Figure 2c). Loss is measured on each cell’s own training distribution.

**Supplementary Figure S3.**
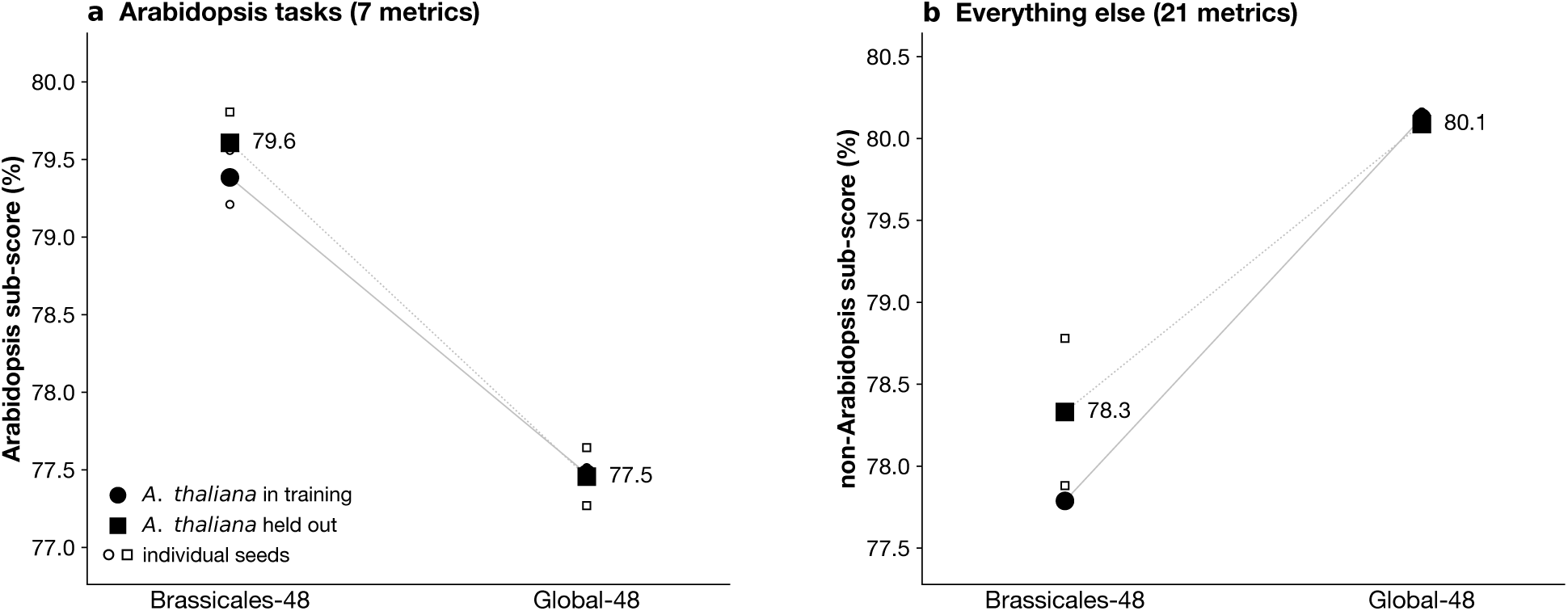
Phylogenetic distance drives dataset leaks. Following the data ablation experiments, 300-M parameter models are trained at iso-step and iso-token with the same configuration except for two settings: global species pool and inclusion or not of Arabidopsis in the pool. Brassicales-48 corresponds to a specialist dataset of Brassicales genomes, Global-48 is a phylogenetically diverse using minHash pairwise distances. Each species pool is trained with or without Arabidopsis. We report the results on a subset of the evaluation suite defining *S*_bal_ (see Supplementary Figure S6). Filled markers are two-seed means; small open markers are the individual seeds. **a**, Average score on Arabidopsis tasks. **b**, Average score on non-Arabidopsis tasks.

**Supplementary Figure S4.**
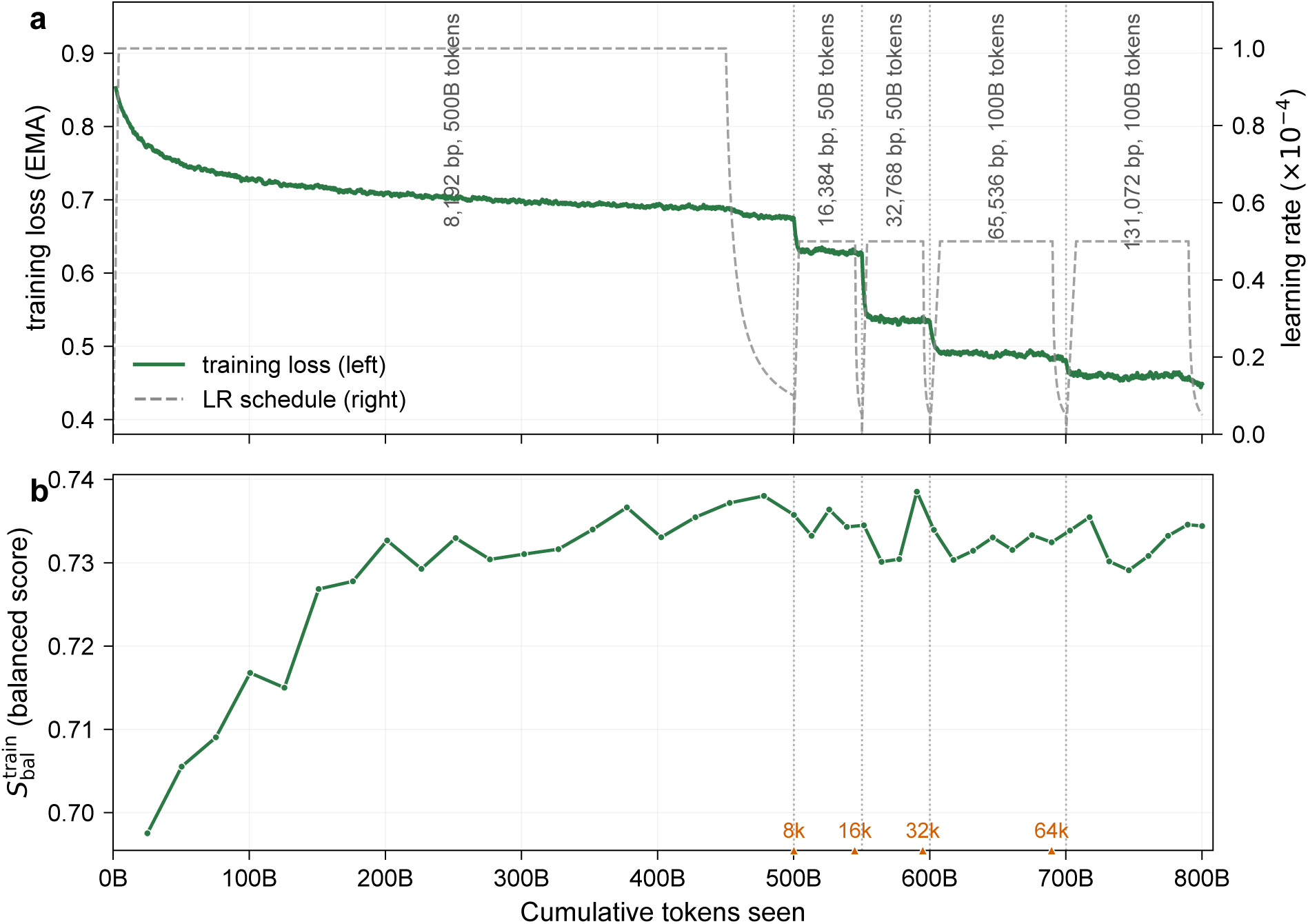
Staged context-window extension of the 318M Botanic backbone. From the 8,192 bp checkpoint (pre-trained for 500 billion tokens), the input window is progressively doubled to 16,384, 32,768, 65,536, and 131,072 bp (dotted lines mark stage boundaries), with a learning-rate warm restart at each stage. Every extension stage runs the same 34,679 optimiser steps: 50, 50, 100, and 100 billion tokens for the four stages, 300 billion in total on top of the 500-billion-token base. **a**, Masked-language-modelling training loss (EMA, left axis) and learning-rate schedule (dashed, right axis) against cumulative tokens. **b**, Evolution of S^train^, evaluated checkpoints shown at a uniform spacing in tokens. The carets mark where the checkpoints evaluated in Figure 6 leave this run: the 8,192 bp rung from the base, the other three from step 31,000 of their parent stage. Each then continues pre-training at its own window until its tokens past the base reach 300 billion, the same total the 131,072 bp arm accumulates along the chain.

**Supplementary Figure S5.**
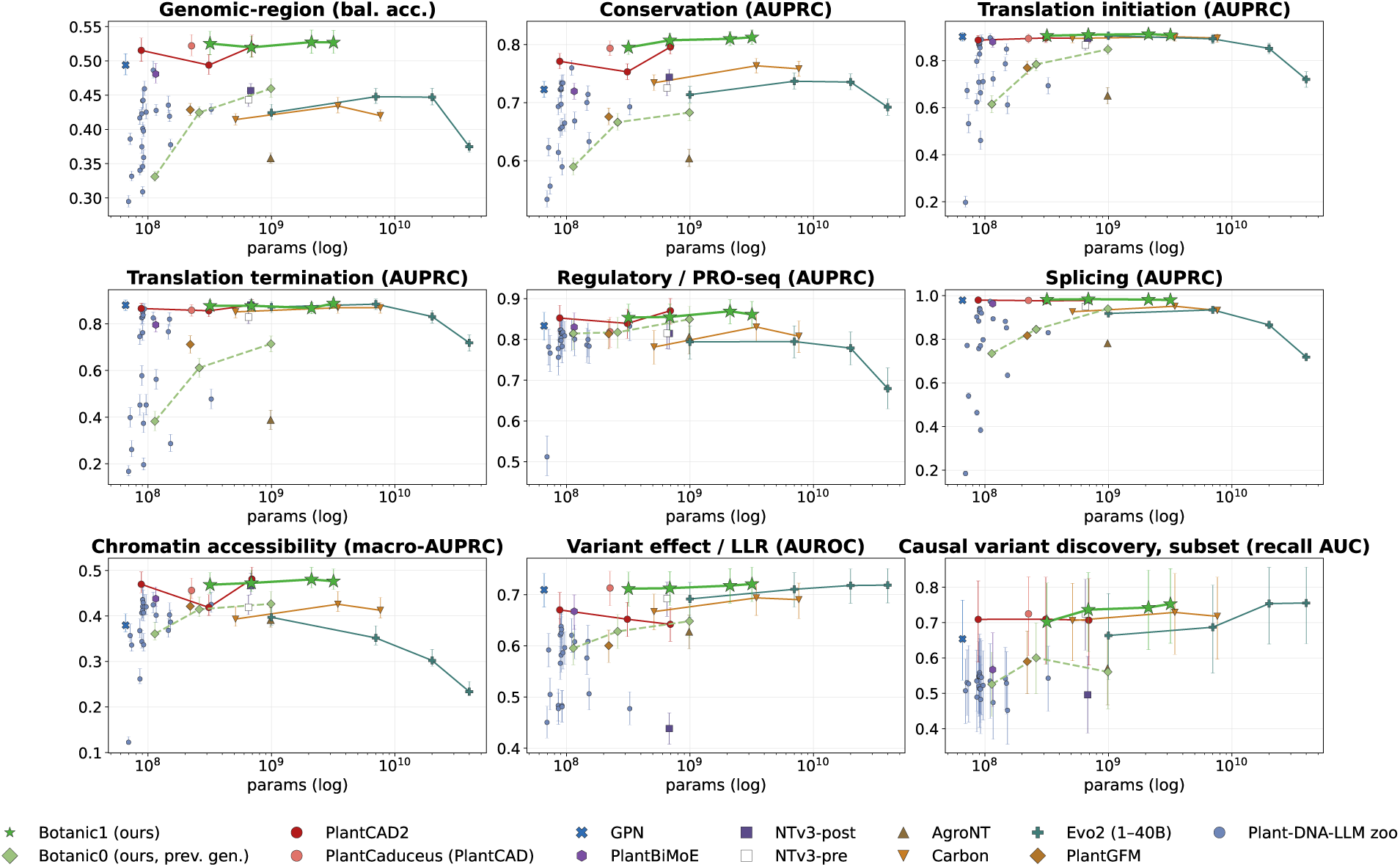
Short-context suite performance versus model size, by capability sub-category. Species-balanced mean for each of the nine capability families (the constituents of *S*_bal_) against trainable parameters (log x), per model family. Trait lines connect multi-size families (Botanic1, Botanic0, PlantCAD2, Carbon, Evo 2, and the Plant-DNA-LLM zoo). Vertical bars are 95% confidence intervals of each family mean under resampling of the test examples within its cells (Section 4.3.1), drawn about the reported value.

**Supplementary Figure S6.**
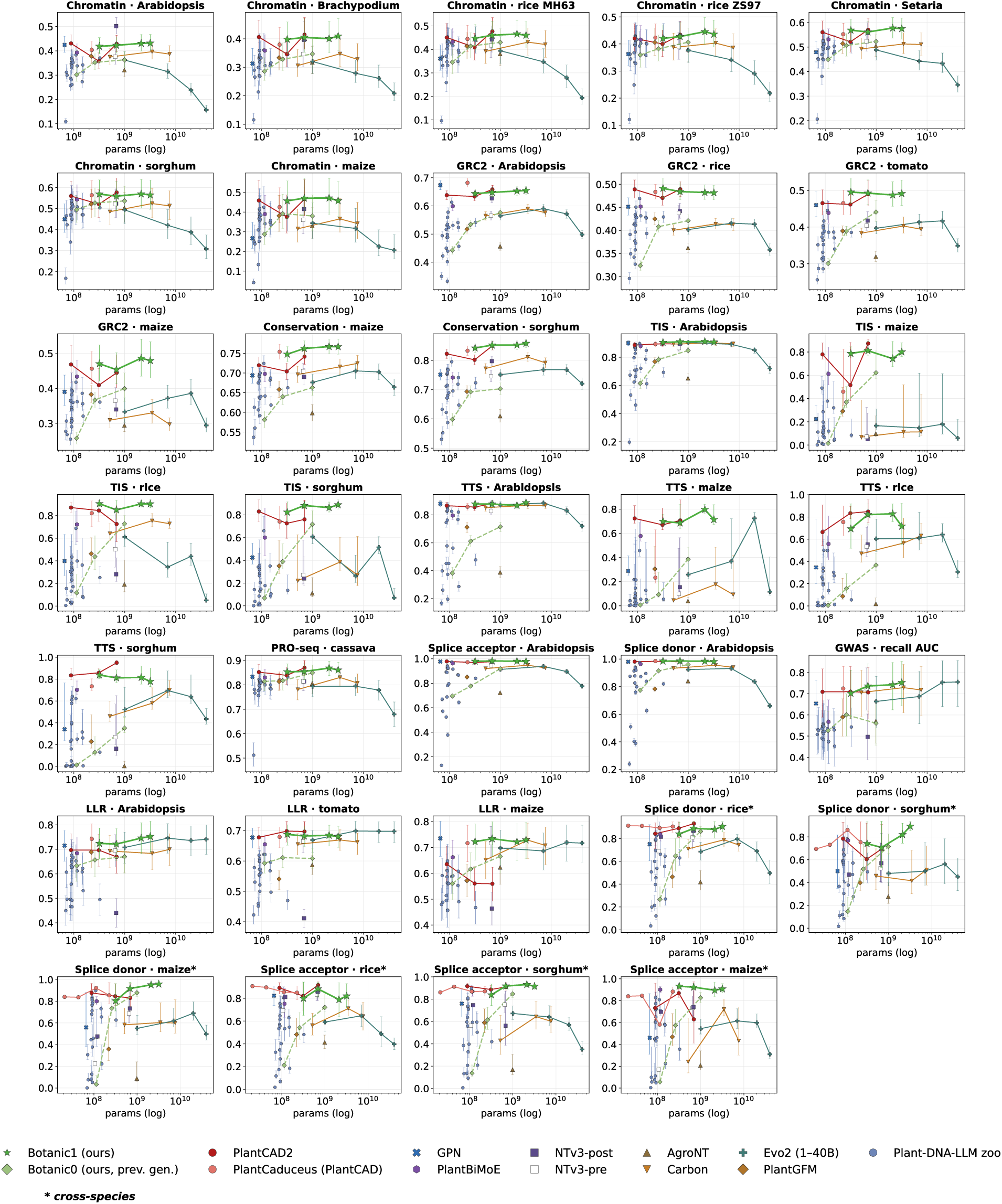
Short-context suite performance versus model size, per individual metric. Zoom in on each of the 34 tasks from frozen model evaluation: the per-species downstream tasks, the three LLR species, causal variant discovery recall (subset), and the six cross-species splice cells and TIS/TTS. Filled squares are used for the released NTv3-post, open squares for the pre-train-only NTv3-pre. Vertical bars are 95% confidence intervals under resampling of the test examples of each cell (Section 4.3.1), drawn about the reported value, for the 22 *S*_bal_ cells and the twelve cross-species cells alike.

**Supplementary Figure S7.**
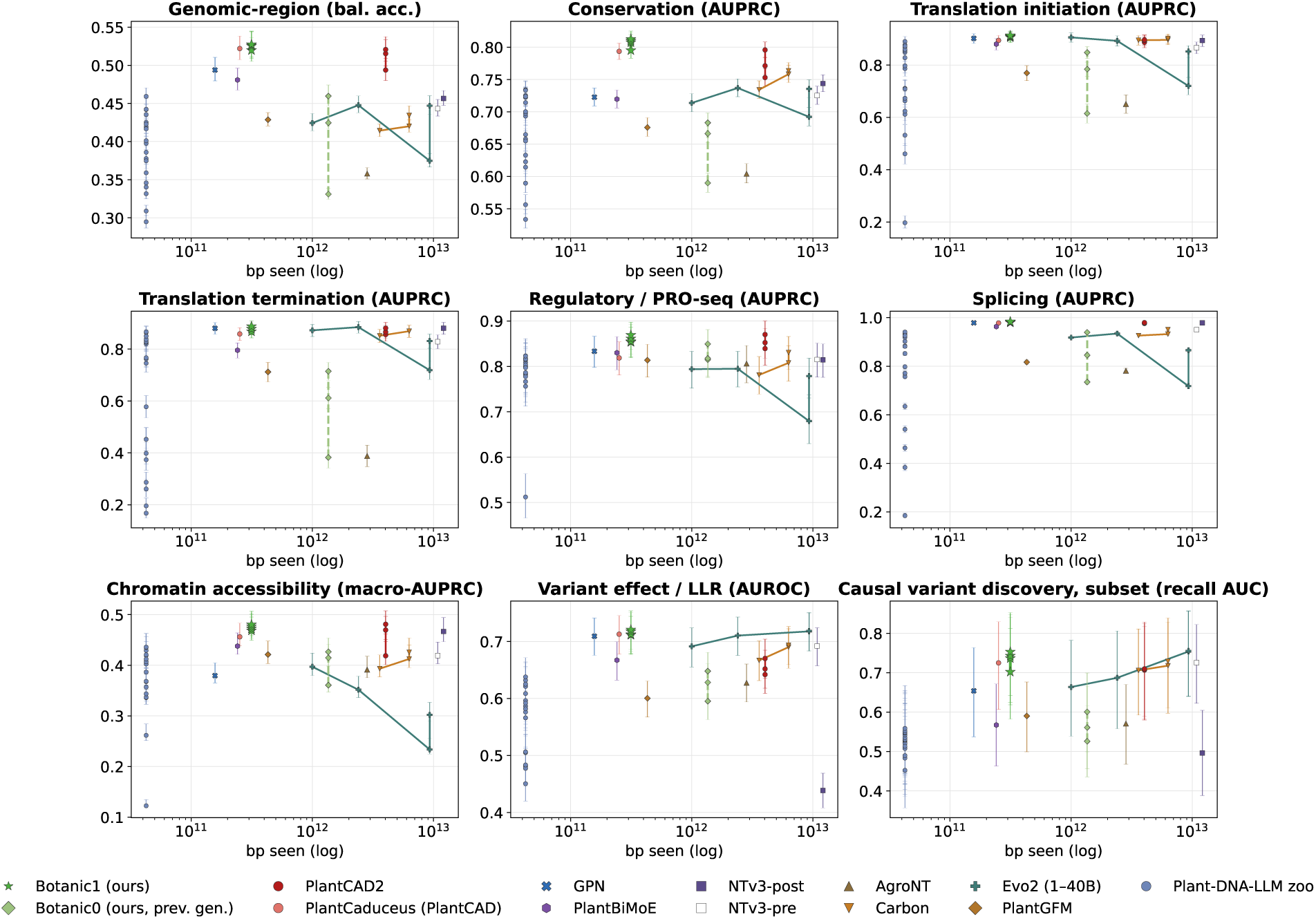
Short-context suite performance versus training base pairs seen, by capability sub-category. Species-balanced mean for each of the nine capability families against the total base pairs seen during training (log x), per model family: the tokens seen converted at each tokeniser’s bases per token. Token counts and provenance are extracted from competitor publications (Supplementary Table S2). The three Plant-DNA-LLM variants released after the publication state no token budget and are absent from these panels. Vertical bars are the confidence intervals of Supplementary Figure S5.

**Supplementary Figure S8.**
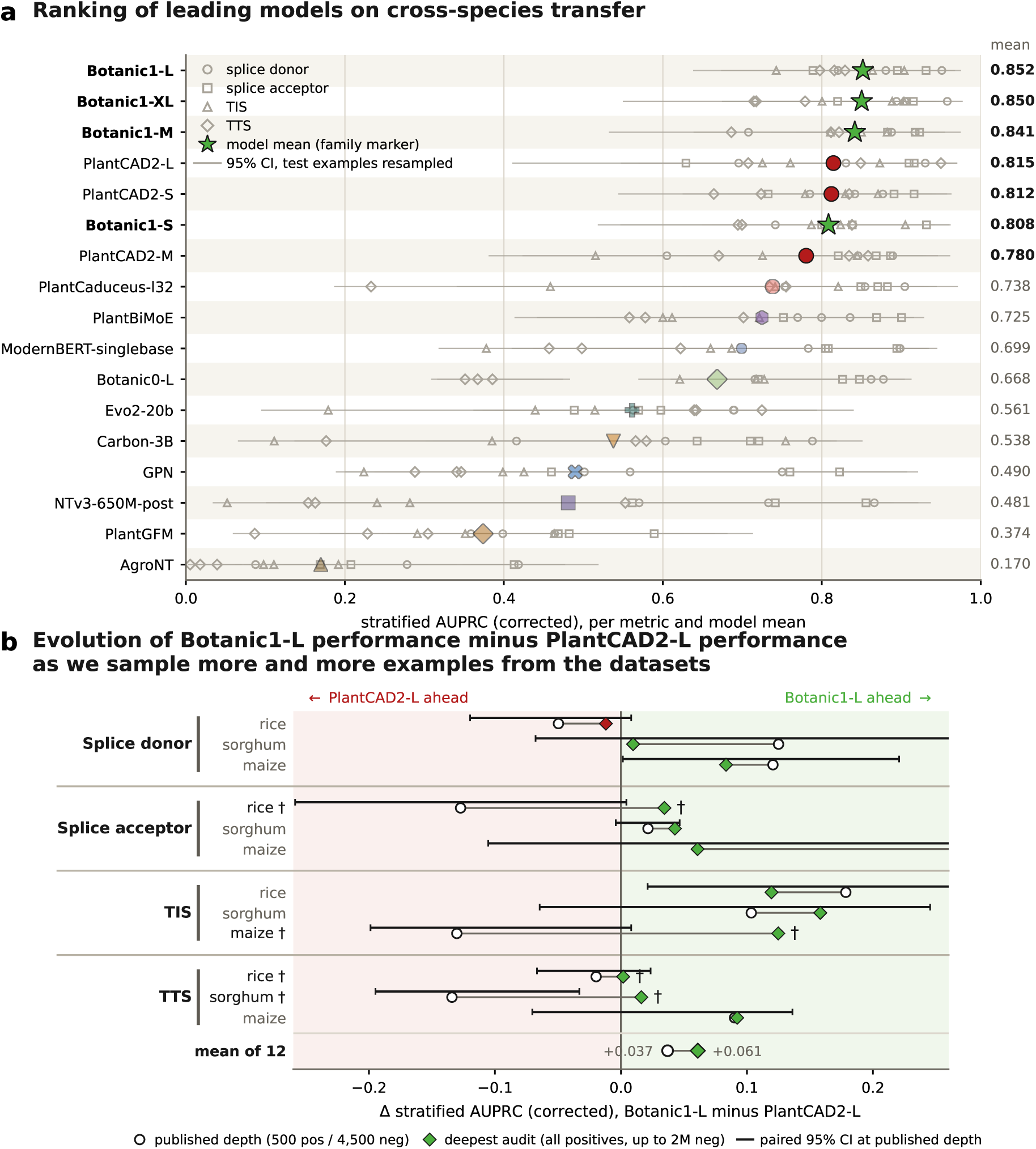
Cross-species transfer of splice-site, TIS and TTS recognition, and the precision of its estimator. Twelve held-out metrics, prevalence-corrected stratified AUPRC for splice donor, splice acceptor, TIS and TTS on rice, sorghum and maize; probes are frozen embeddings trained on *Arabidopsis thaliana* only, and all twelve metrics are held out of *S*_bal_. Every model is scored under the test evaluation protocol 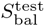: centre token for encoders, two-strand centre pooling for the autoregressive models. **a**, The twelve metrics per model (small open marks, one glyph per task) and their mean (family marker), one row per model: the four Botanic sizes, the three PlantCAD2 sizes and the strongest member of each remaining family, sorted by mean. The thin whisker behind each mark is its 95% confidence interval under resampling of the test examples of that cell (Section 4.3.1). **b**, Botanic1-L against PlantCAD2-L. For each metric, the difference between the two at the published sampling depth (500 positives, 4,500 negatives; open circles) and at the deepest audit (every positive kept, up to 2 million negatives; filled diamonds) (Section 4.3.5). The black bar above each row is the paired 95% interval of the published-depth difference. a dagger (*†*) marks the four metrics where the ordering of the pair reverses at the deepest depth. Only this pair of models is audited.

**Supplementary Figure S9.**
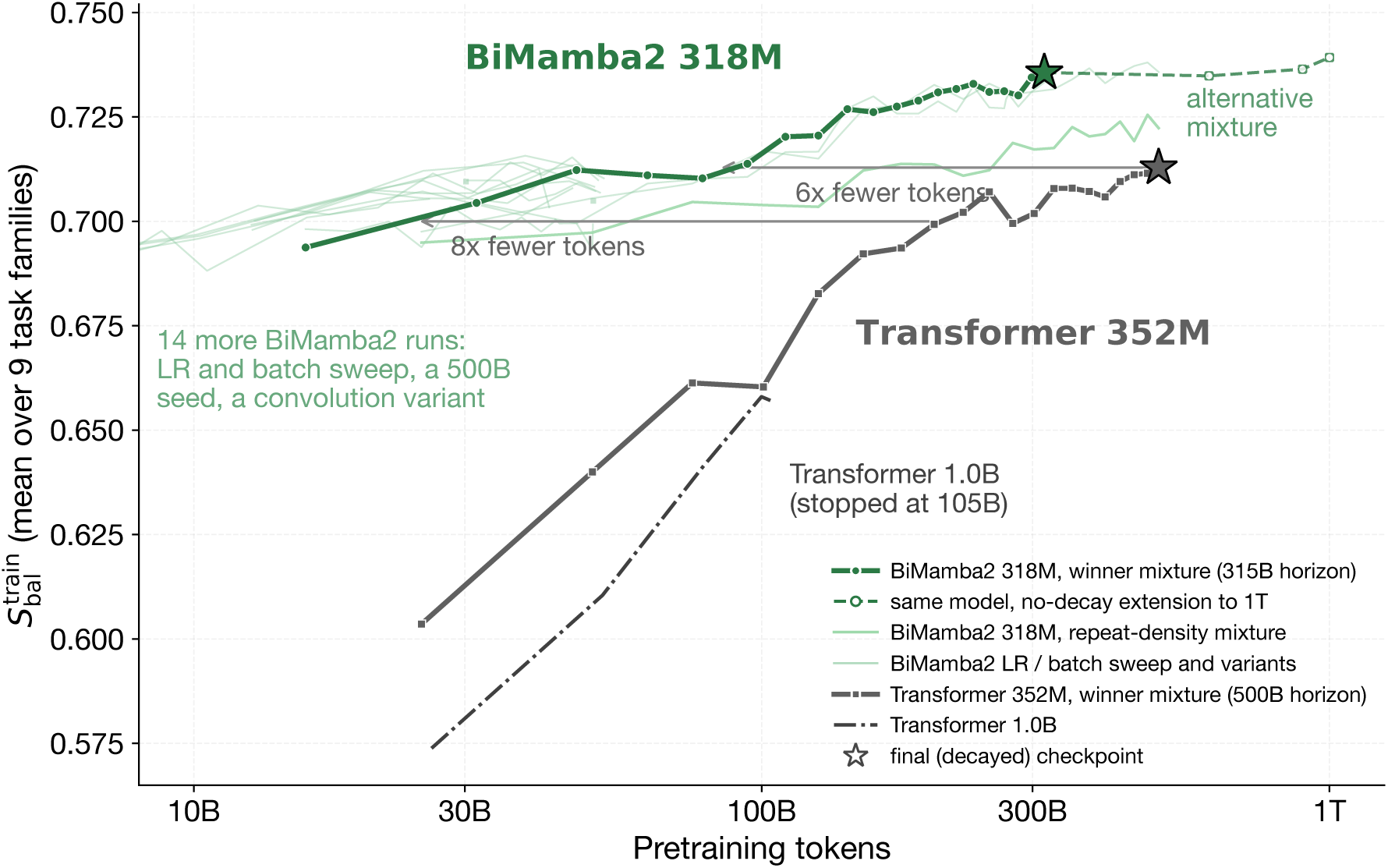
Architecture comparison at a matched budget: BiMamba2 against a Transformer encoder on 8,192-token contexts. 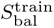 against tokens seen for Botanic1-S (318M BiMamba2 parameters) and a 352M Transformer encoder pre-trained with the same data, tokeniser, window length, peak learning rate and learning rate schedule on a 500-billion-token horizon. Grey lines connect iso budget scores: the Transformer needs about six times more tokens to reach its own final value and about eight times more to reach 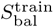 = 0.70. Thin green lines are other fourteen BiMamba2 runs (a learning-rate and batch-size sweep, a second 500B seed on an alternative mixture, and a convolution-augmented variant); the dash-dotted line is a 1.0B Transformer using the same pre-training scheme whose training stops at 105B tokens. The dashed continuation is the beginning of a training extending Botanic1-S to one trillion tokens at constant learning rate. Stars mark final decayed checkpoints. Evaluation noise per point is about ±0.003.

**Supplementary Figure S10.**
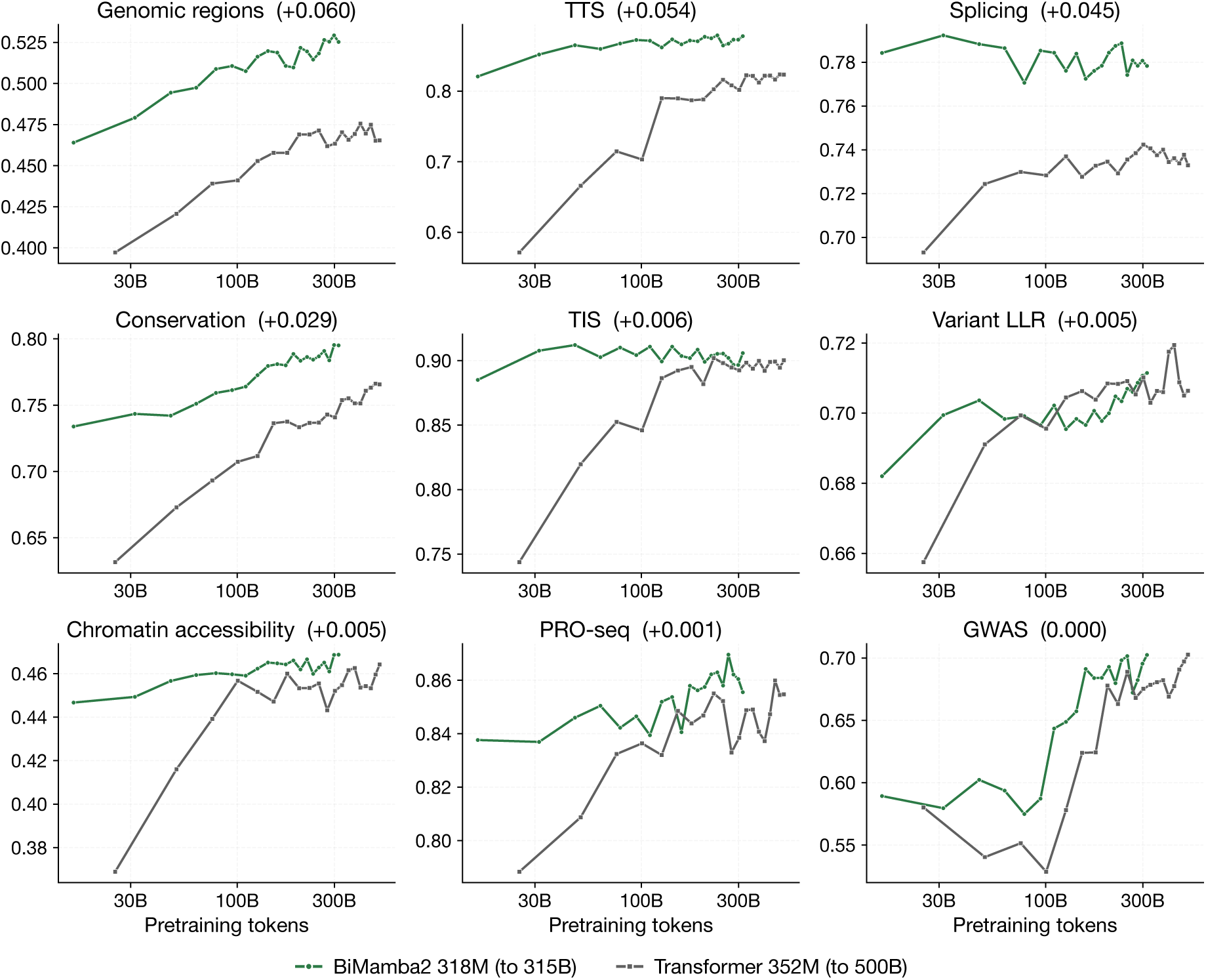
The same architecture pair per task family. Species-balanced family means for the nine task families included in *S*_bal_, scored under the 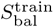 protocol, for Botanic1-S (to 314.6B tokens) as well as a 352M Transformer encoder, against tokens seen. The value in each panel title is the remaining difference at the two models’ final checkpoints.

**Supplementary Figure S11.**
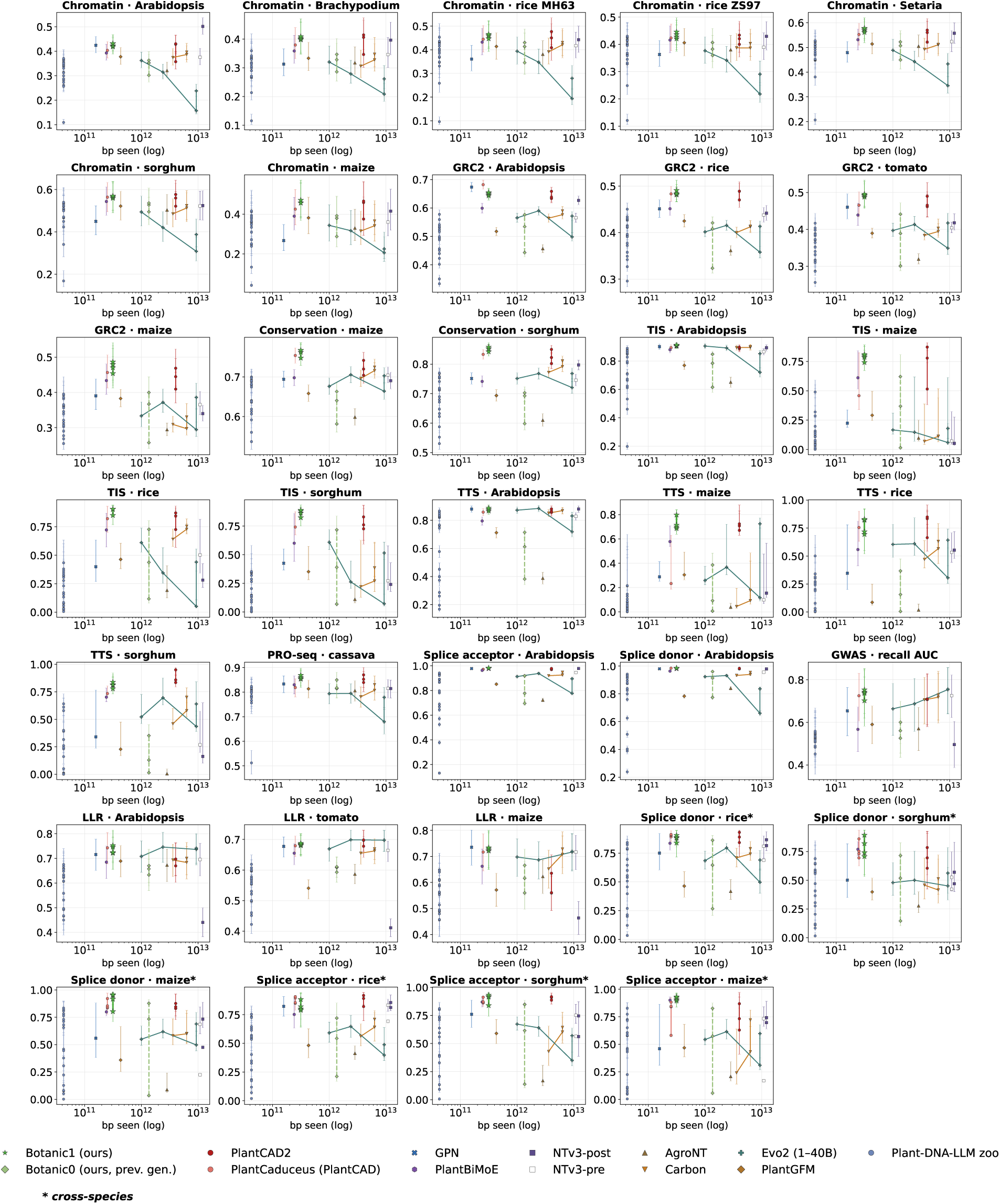
Short-context suite performance versus training base pairs seen, per individual metric. Individual 34 metrics against the total base pairs seen during training (tokens seen converted at each tokeniser’s bases per token). Token counts and their provenance are given in Supplementary Table S2. Vertical bars are the confidence intervals of Supplementary Figure S6.

**Supplementary Table S1:**
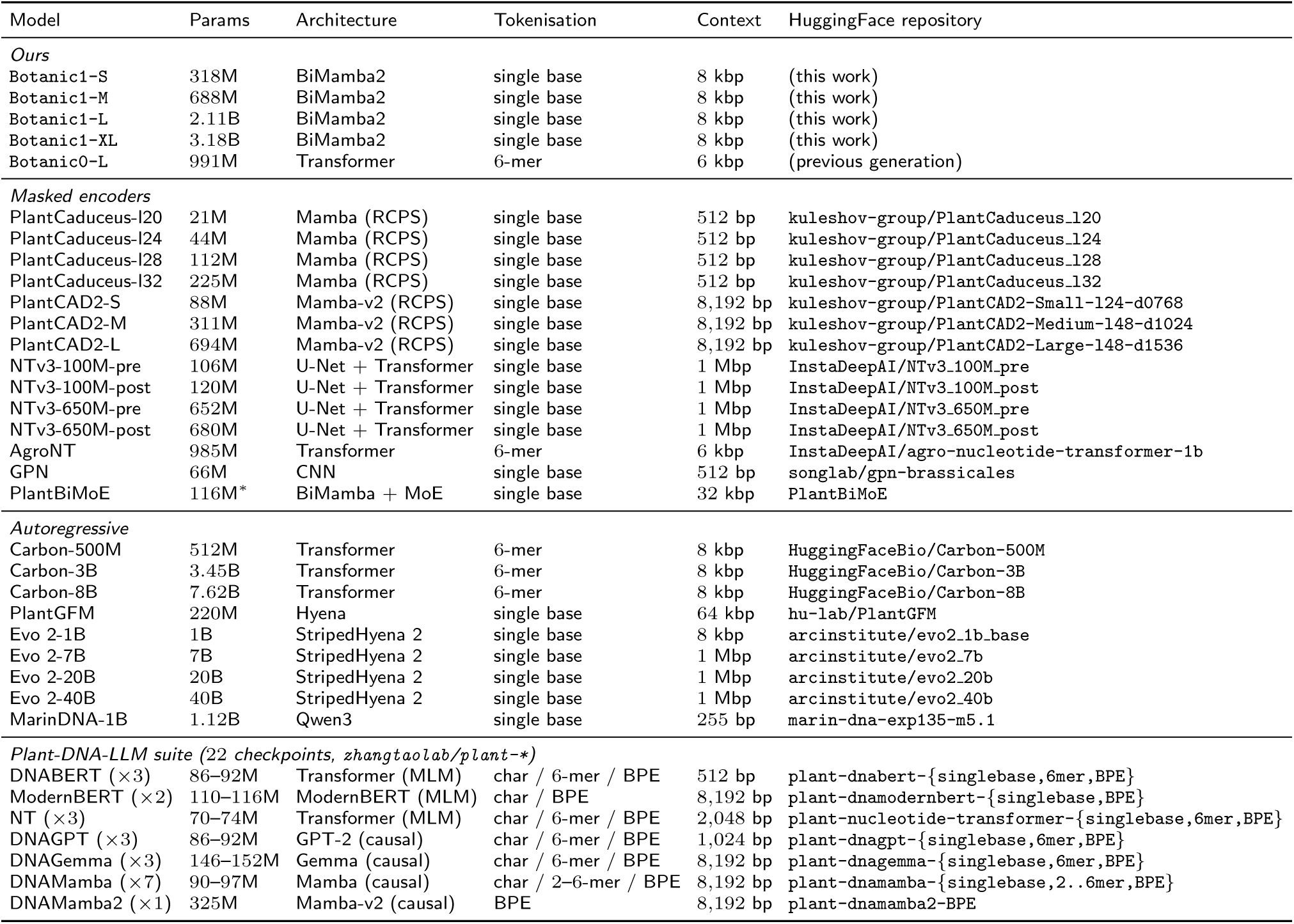
Baseline backbones evaluated in this report. Parameter counts are trainable parameters as declared by each model’s configuration and code. Context is the maximum the released checkpoint can accept. RCPS stands for reverse complement parameter sharing, a design used in the PlantCaduceus and PlantCAD2 families. The Plant-DNA-LLM suite is 22 checkpoints with four architectures and three tokenisations; they are grouped by architecture, with per-checkpoint parameter counts available in Supplementary Figure S6.

**Supplementary Figure S12.**
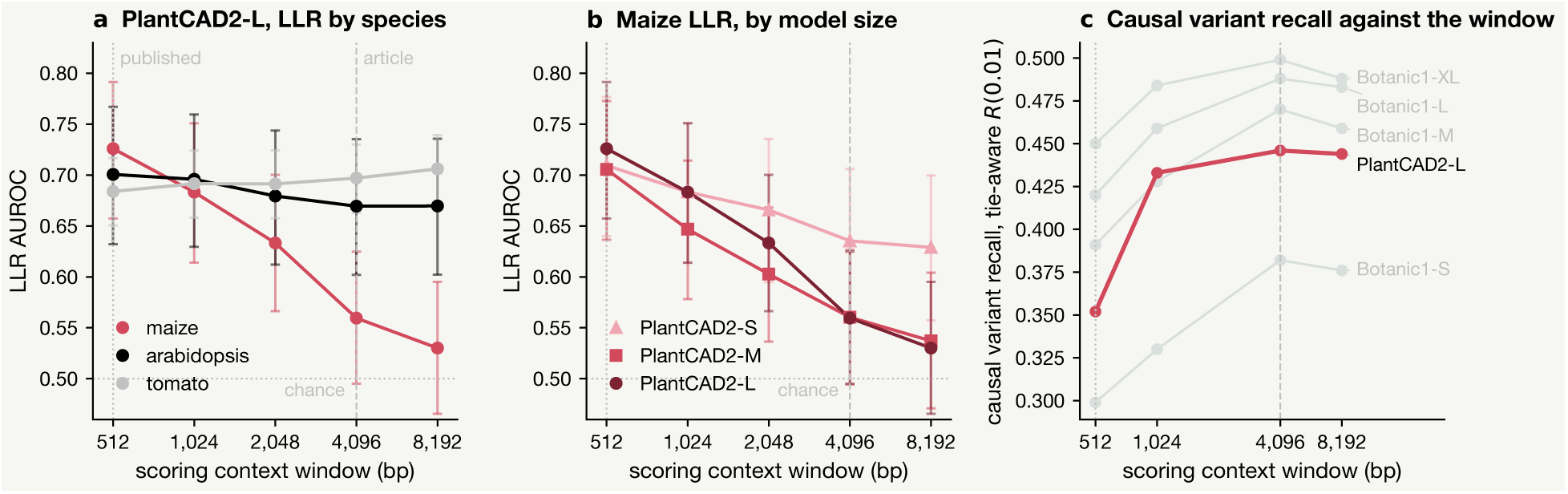
Zero-shot scores against the scoring context window, for the PlantCAD2 family. W = min(4,096, trained context) puts PlantCAD2 at 4,096 bp; GPN, PlantCaduceus and the Plant-DNA-LLM suite, trained on short contexts, are scored at 512 bp (Supplementary Table S5). **a**, PlantCAD2-L on the three LLR species, from 512 to 8,192 bp. Tomato score rises (+0.022) and arabidopsis score is nearly flat ( −0.031) while maize score falls 0.196, from 0.726 to 0.530. **b**, Maize alone, per size: maize is the most repeat-rich of the three genomes. **c**, The context sweep for causal variant discovery follows an opposite trend as above: PlantCAD2-L rises from 0.352 at 512 bp to 0.446 at 4,096 and plateaus at 8,192. Causal-variant recall over the 29 studies gains +0.063, +0.032 and +0.030 for S, M and L on moving to 4,096 bp. Dotted and dashed rules mark 512 and 4,096 bp. Interval bars in **a** and **b** are 95% percentile bootstraps over the variants of each cell, which is why maize, with 376 variants against tomato’s 1,799, has the widest interval.

**Supplementary Table S2:**
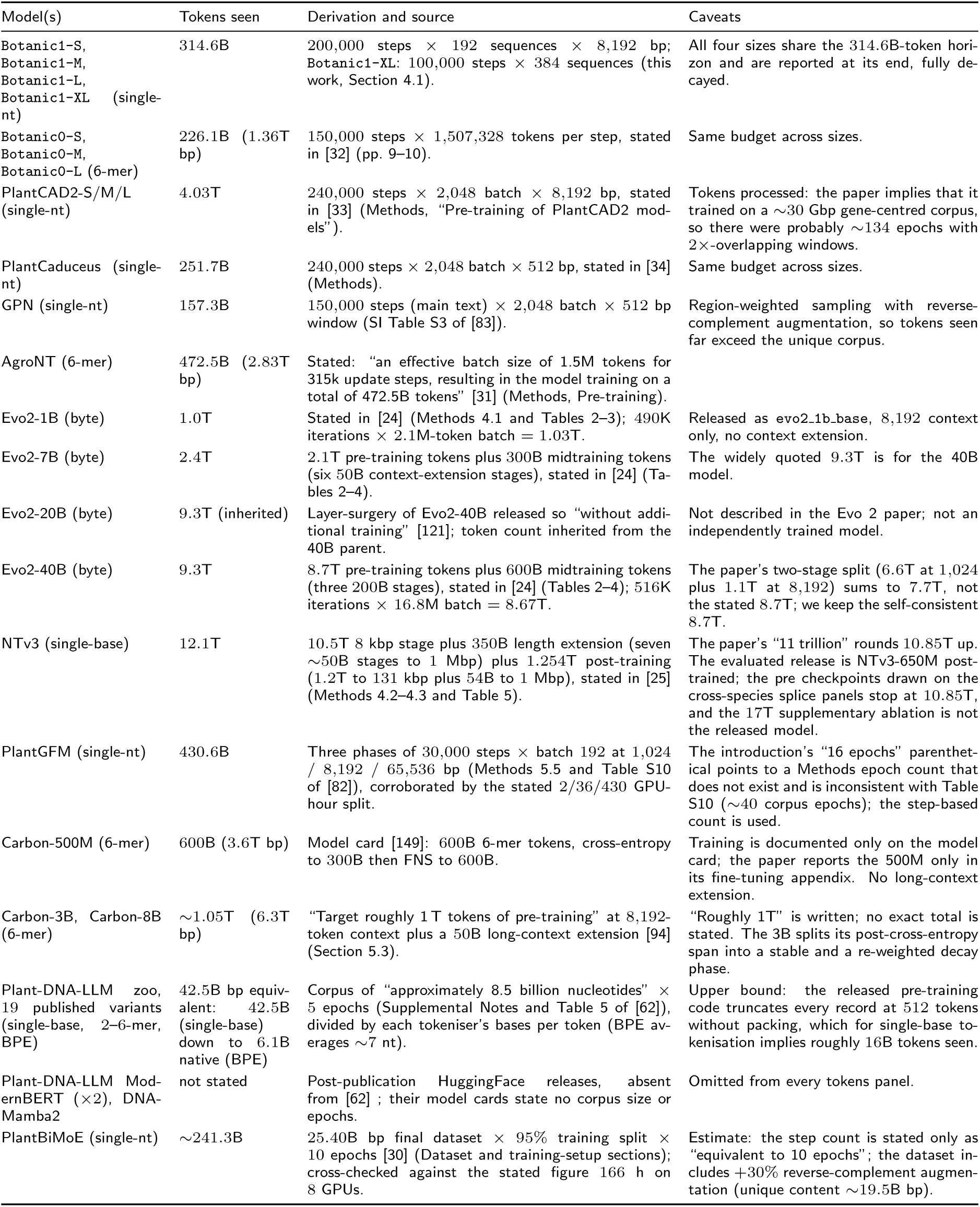
Per-model training-token accounting. “Tokens seen” is the number of tokens processed during training: what is reported in the original source, or if not present steps × global batch × sequence length, or if not present corpus × epochs. Original tokens values cannot be used as each tokeniser uses a different number of base pairs per token. One token spans one base for single-nucleotide and byte tokenisers, *k* bases for non-overlapping k-mer tokenisers, and about seven bases for the Plant-DNA-LLM BPE variants. Every value is verified against the cited source; the location of the supporting statement is given.

**Supplementary Figure S13.**
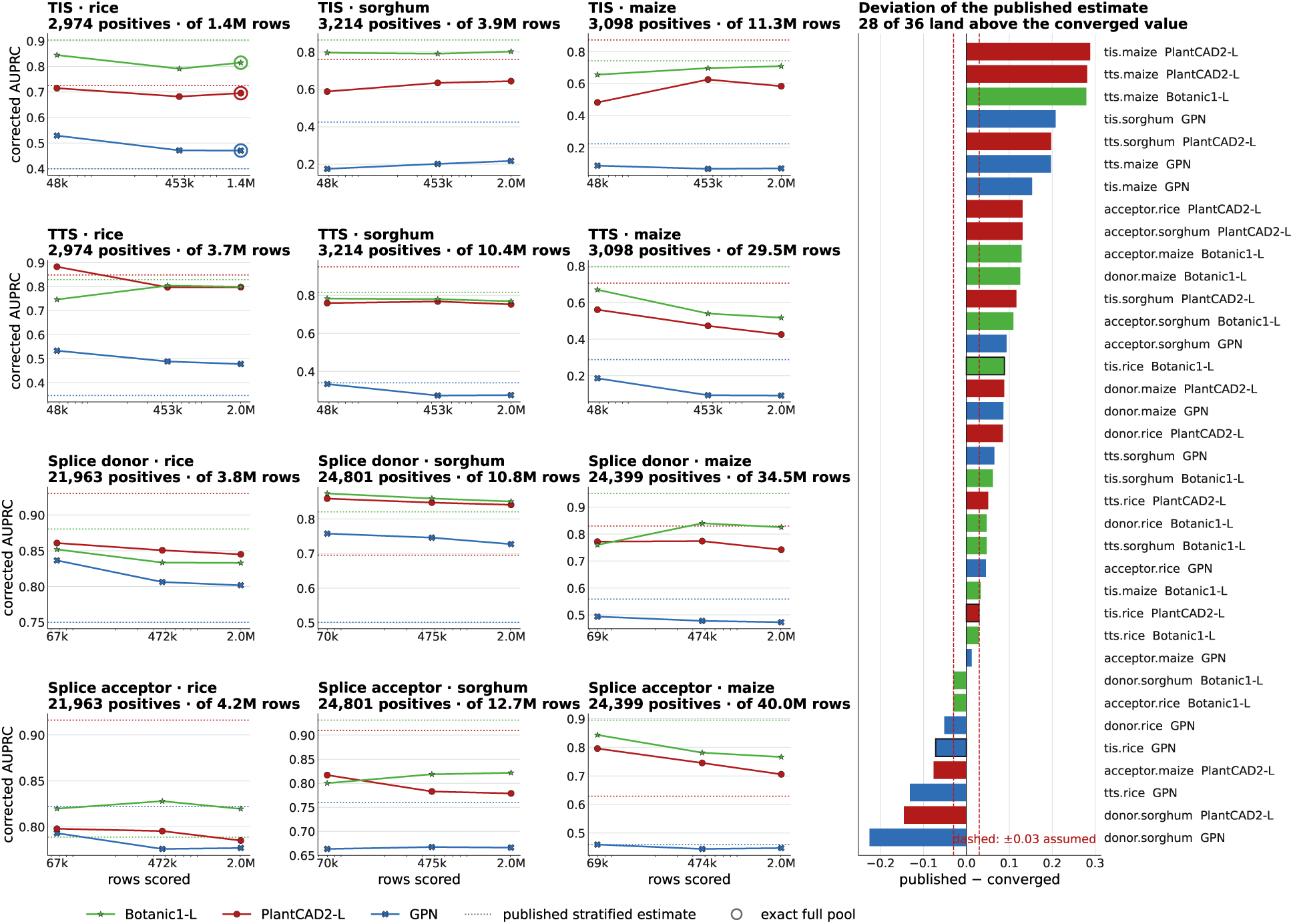
Precision of the stratified estimator used for the cross-species metrics. Each point keeps every positive and varies the number of negatives included, while applying prevalence correction: one panel per task and species, each showing the positive-complete estimate at 45,000, 450,000 and 2 million negatives for the three models. We report the reported estimator value as a dotted line and the exact full-dataset value with a ring when the dataset is small enough to reach that mark. Panel titles shows the number of positives kept, the range of rows being scored and the total dataset size. Every task-species displays all three models result to 2 million negatives, or to the full pool of 1,396,105 for rice TIS, so each comparison is at matched depth. The right-hand column shows the finite estimator value reported minus the converged estimate for every task-species and model measured, against the ±0.03 from the parametric model; bars with a bold edge are the three comparisons where the converged value is exact. Deviations reach ~ 0.3, and 28 of the 36 reported values are above their converged value. On eight of the twelve metrics the ordering of the three probes is unchanged; on splice acceptance in rice, TIS in maize and TTS in rice and sorghum the ordering of two models reverses with more negatives samples.

**Supplementary Figure S14.**
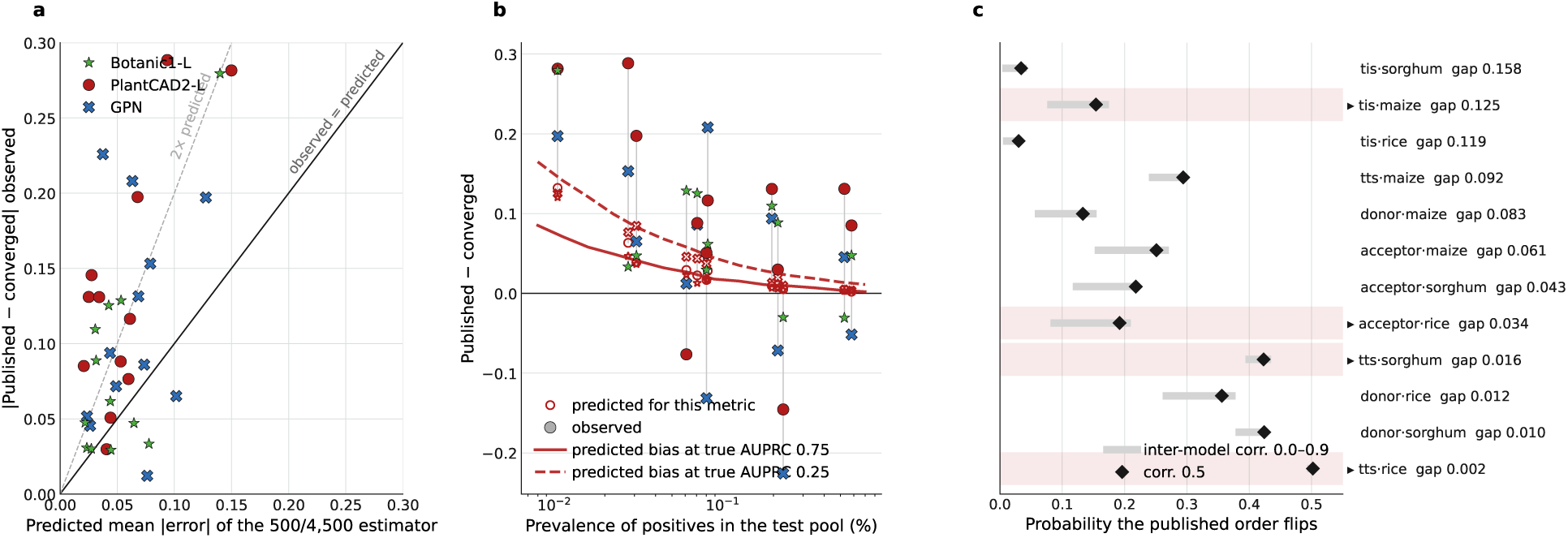
A simple parametric model explains only part of our observations. Score distributions are calibrated so that their true AUPRC at the measured prevalence is equal to the measured converged value, and the 500-positive, 4,500-negative draw is then drawn from the simple simulated model. Nothing is fitted to the observed deviations. **a**, Predicted mean absolute error of our model against the deviation observed: the sampling term accounts for roughly half of the measured magnitude. **b**, Our estimator is always too optimistic. Hollow markers are the bias predicted by the simple model for each metric, filled markers what is observed, and the red curves are each prevalence points. Estimated average precision is biased upward increasingly with positives getting rarer, and 28 of 36 values we report are above their converged true value. **c**, Probability that the published ordering of Botanic1-L against PlantCAD2-L flips, simulated with both models scored on the identical rows and inter-model score correlation from 0 to 0.9. The model predicts 2.3 to 3.1 reversals; four are observed by increasing the number of negatives. We highlight those tasks and species in the panel.

**Supplementary Table S3:**
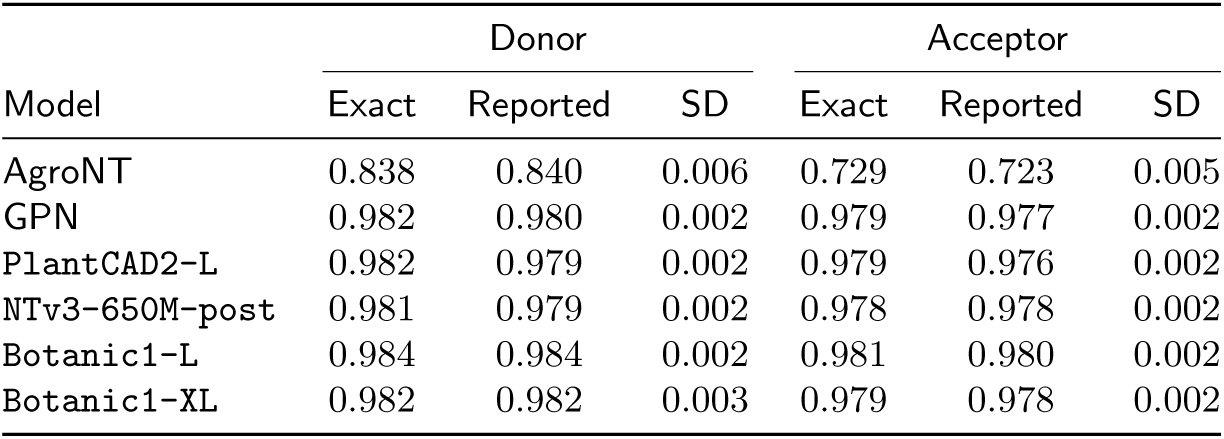
Precision of the stratified estimator behind the *S*_bal_ splicing metrics. Prevalence-corrected stratified AUPRC under the frozen-embedding protocol. Exact means the full-dataset value (377,873 rows for donor, 250,084 for acceptor); Reported means the published leaderboard value; SD is the standard deviation of the estimator over 10,000 prevalence-matched draws of the same size, shared across models.

**Supplementary Figure S15.**
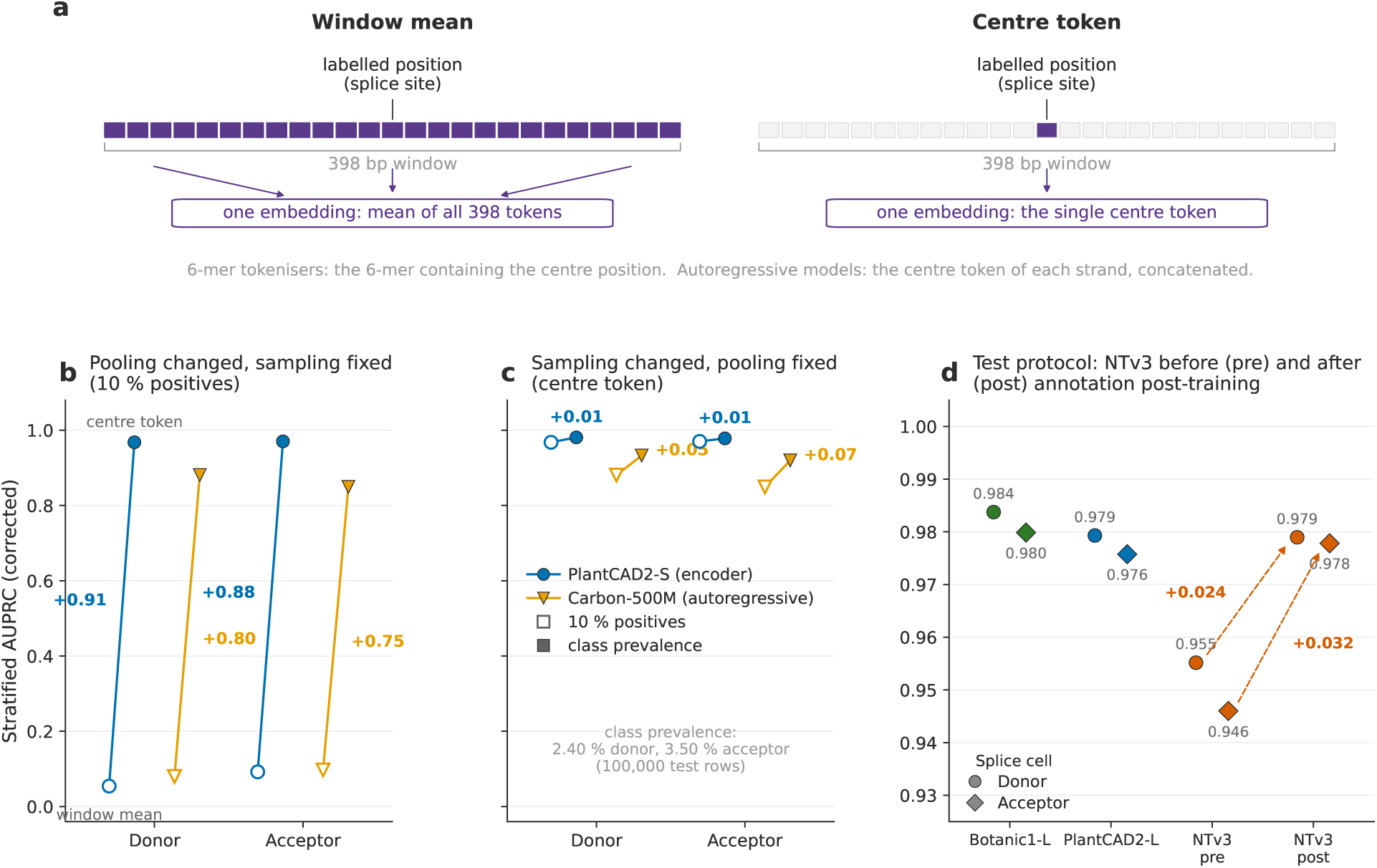
Effects of varying pooling, test sampling and training strategy. **a**, The two pooling rules applied to the 398 bp splicing windows. Window-mean pooling averages the embeddings of all tokens in the window; centre-token pooling keeps the embedding of the single token at the labelled position (for 6-mer tokenisers, the 6-mer containing it; autoregressive models take that token on each strand and concatenate the two, Section 4.3.6.1). **b**, Prevalence-corrected stratified AUPRC of one encoder (PlantCAD2-S) and one autoregressive model (Carbon-500M) on *Arabidopsis thaliana* splice donor and acceptor recognition, from window-mean (open) to centre-token (filled) pooling on the same stratified test draw of 500 positives and 4,500 negatives. The change is +0.75 to +0.91. **c**, The same two models under centre-token pooling when the test draw moves from 10% positives to the measured class prevalence (2,400 donor or 3,500 acceptor positives among 100,000 rows), on the same axis as **b**. The prevalence correction makes both draws estimates of the same quantity; the change is +0.01 to +0.07. **d**, Splicing under the 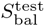 protocol (centre token, prevalence-matched draw) for Botanic1-L, PlantCAD2-L, NTv3-650M-pre and NTv3-650M-post; circles are the donor cell and diamonds the acceptor cell. The dashed arrows mark the gain of the post-trained NTv3 checkpoint over the pre-training-only one on each cell (Section 3).

**Supplementary Figure S16.**
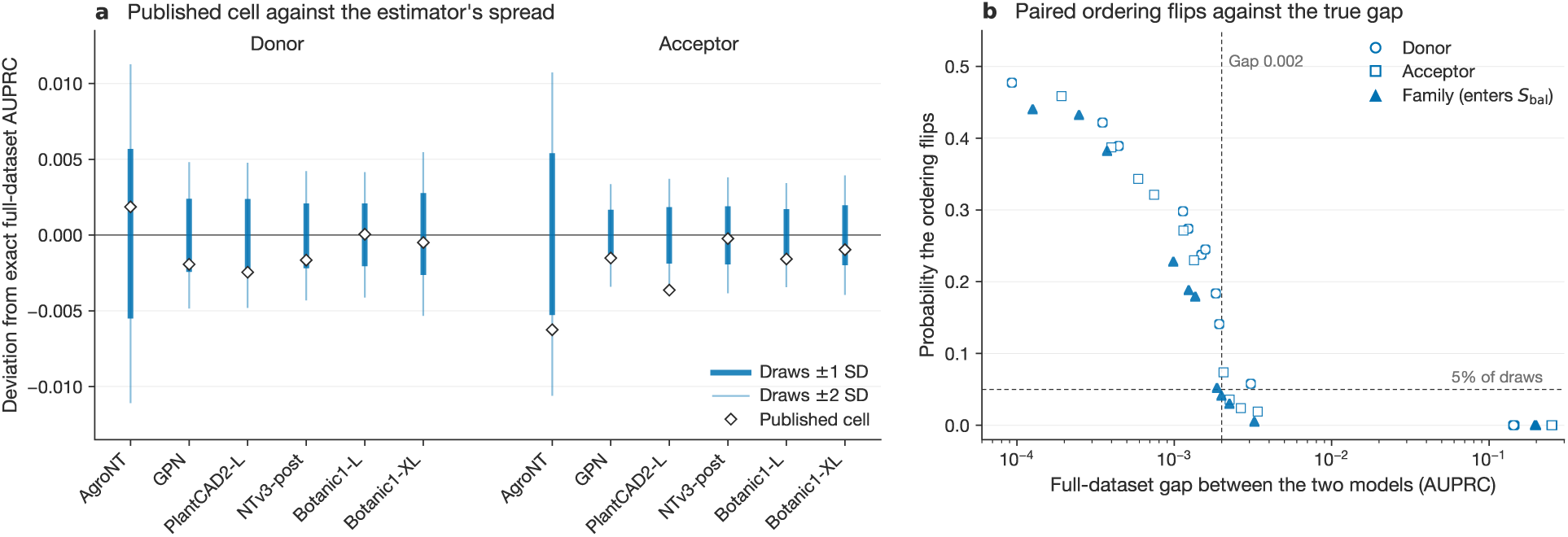
Precision of the stratified estimator behind the *S*_bal_ splicing metrics. Six models are re-scored on the full test datasets keeping all rows; the corrected stratified AUPRC estimator is then replayed 10,000 times in order to estimate the uncertainties behind the sampling. **a**, Deviation of the published leaderboard metrics from the exact full-dataset value, against the *±*1 and *±*2 standard-deviation spread of the bootstrapped estimator (bands centred on the draw mean; the bias is below 10*^−^*^4^ everywhere). **b**, Probability that the ordering of two models reverses under the estimator, against their full-dataset gap; draws are shared across models.

**Supplementary Figure S17.**
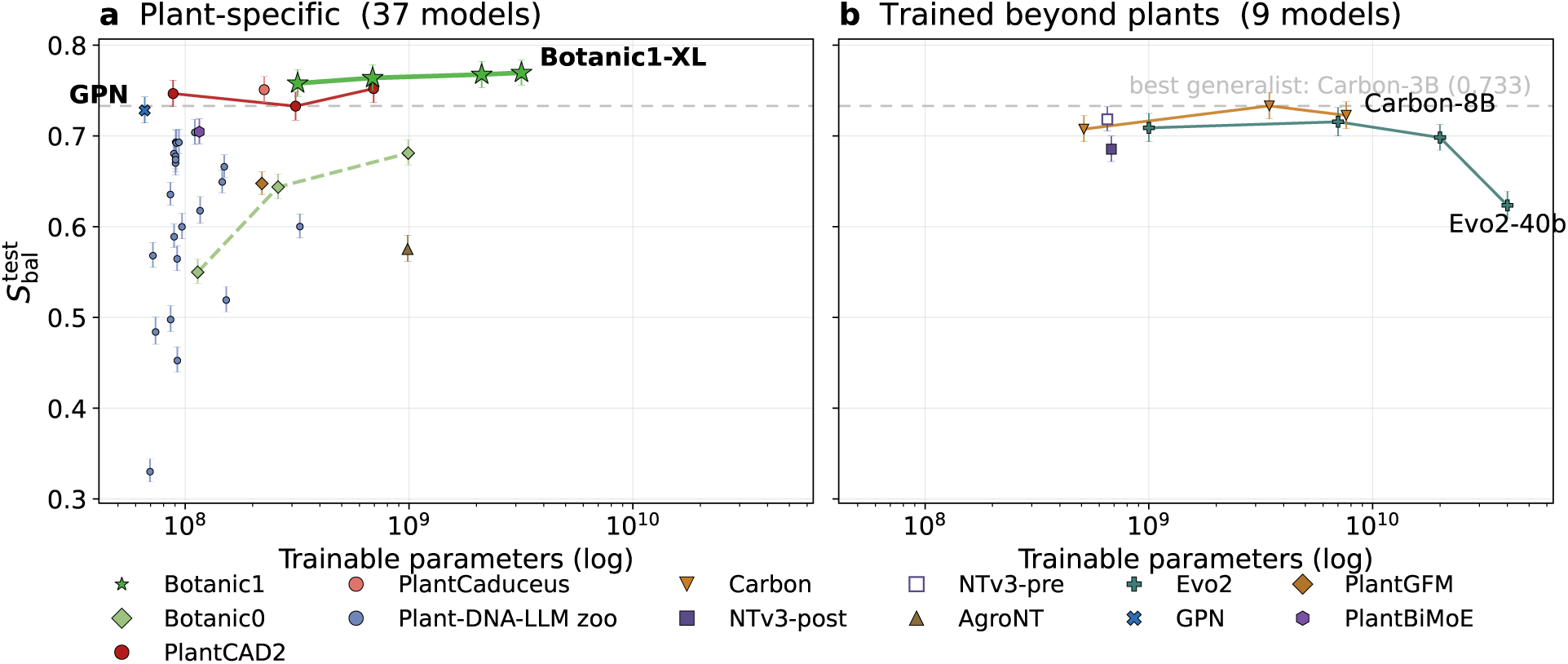
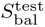 against model size, split by pre-training domain. **a**, Models pre-trained on plant genomes only. **b**, Models trained across broader domains of life: the Nucleotide Transformer v3, Carbon and Evo 2. Both panels share axes, and the dashed rule marks the best generalist score (Carbon-3B, 0.733). Vertical bars are 95% confidence intervals under resampling of the test examples within each cell (Section 4.3.1), drawn about the reported score.

**Supplementary Table S4:** The nine task families used in 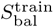 and 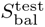. Each family contributes one ninth of the score irrespective of how many species it is evaluated on: for instance chromatin accessibility alone supplies seven of the 22 metrics and would otherwise dominate an unweighted mean. Supervised probes (XGBoost) are trained on frozen representations; the two zero-shot families score variants using the model’s likelihood directly. Splicing contributes two metrics from one species (donor and acceptor sites), each scored on a stratified sample drawn at the target class prevalence (2,400 positives against 97,600 negatives for donor, 2.33% true prevalence; 3,500 against 96,500 for acceptor, 3.53%), so the prevalence-corrected stratified AUPRC is precise enough to enter *S*_bal_ (Section 4.3.5). Causal variant discovery is scored on a fixed subset of 29 curated studies from four species (13 *Arabidopsis thaliana*, 11 *Oryza sativa*, 4 *Triticum aestivum*, 1 *Sorghum bicolor* ), each causal SNP ranked by signed LLR against 100 background SNPs from its locus, and contributes a single recall-AUC scalar; the full 545-study benchmark of Section 2.5.1.2, which ranks every documented SNP of the locus by |LLR|, is reported separately. The rows therefore sum to the 22 metrics of *S*_bal_. The suite logs twelve further metrics that are not used within *S*_bal_: TIS and TTS on rice, sorghum and maize and splice donor and acceptor on the same three species (Supplementary Figure S8a), all scored with a stratified, prevalence-corrected AUPRC, totalling 34 frozen models metrics.

| Family | What it measures | Species | Metric |
| --- | --- | --- | --- |
| *Supervised probes* |  |  |  |
| Chromatin accessibility | open versus closed chromatin | 7 | macro-AUPRC |
| Genomic region | genic compartment of a window | 4 | balanced acc. |
| Conservation | evolutionarily constrained positions | 2 | AUPRC |
| Splicing | donor and acceptor site recognition | 1 | strat. AUROC ($S_{\text{bal}}^{\text{train}}$) /<br>strat. AUPRC ($S_{\text{bal}}^{\text{test}}$) |
| TIS | translation initiation site | 1 | AUPRC |
| TTS | translation termination site | 1 | AUPRC |
| PRO-seq | nascent transcription | 1 | AUPRC |
| *Zero-shot* |  |  |  |
| LLR mutation effect | deleterious versus benign variants | 3 | AUROC |
| Causal variant discovery | experimentally supported variant recovery (29 studies) | 4 | recall-AUC |

**Supplementary Table S5:** Task definitions and scoring protocol behind 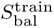 and 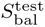. One row per task family; the species included within each family are listed in Supplementary Table S6. Window is the sequence length given to the model. The two “probing-protocol” columns explains how exactly is the embedding obtained for encoders-only (masked language models) and for decoder-only models (Carbon, Evo 2, PlantGFM and the causal Plant-DNA-LLM checkpoints); where the two protocols differ, the 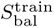 value is given first and the 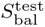 value second, and the decoder-only column is always the 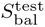 protocol. 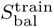 is a training signal only: it is used to steer data and architecture decisions for Botanic1, which is an encoder-only model. Every number compared across models in this article is 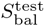. Centre-token pooling takes the embedding of the token at the centre of the window; centred-window pooling takes the mean over a central 100 bp window; mean pooling takes the mean over every token of the window. Encoders-only models use one forward pass, except on genomic region classification, where the sequence and its reverse complement are both embedded and averaged (RC-averaged), following a protocol described in GPN. Decoder-only models see only the left context of a token, so for centre-token and centred-window pooling we use one forward pass per strand and the two centre embeddings are then concatenated (RC-concatenated, Section 4.3.6.1); mean pooling is one forward pass for every model. The two zero-shot families do not use probes: encoders score a variant by the log-likelihood ratio (LLR) of the alternate against the reference allele at the masked variant position, decoder-only models by the difference of the whole-sequence log-likelihoods of the two alleles; causal variant discovery ranks by the signed LLR (Section 4.10). The scoring window is W = min(4,096, trained context) centred on the variant: 4,096 bp for every model except GPN, PlantCaduceus and the Plant-DNA-LLM suite, scored at 512 bp (480 bp for its single-base checkpoints). *^a^* Checkpoints with byte-pair tokens are centre-pooled here like every other checkpoint. a base-pair-derived centre does not map onto a token boundary for those models, so both span endpoints are resolved from the tokeniser’s offset mapping rather than from a fixed bases-per-token ratio. *^b^* Masked models with byte-pair tokens cannot mask a single position, since the two alleles tokenise differently; they mask in turn every token overlapping a ±12 bp region centred on the variant and take the mean per-token log-probability.

| Family | Source | Task type | Window (bp) | Probing protocol | | Metric ($S_{bal}^{train}$ / $S_{bal}^{test}$) | Task definition |
| --- | --- | --- | --- | --- | --- | --- | --- |
| | | | | Encoders ($S_{bal}^{train}$ / $S_{bal}^{test}$) | Decoder-only ($S_{bal}^{test}$) | | |
| Chromatin accessibility | PGB [31] | multi-label (9 to 19 tracks) | 1,000 | mean | mean | macro-AUPRC | open chromatin, one label per accessibility track |
| Genomic region | GPN-derived [83], built here | multi-class (7 classes) | 512 (central 100 scored) | centred-window, RC-averaged<sup>a</sup> | centred-window, RC-concatenated<sup>a</sup> | balanced accuracy | annotation class: CDS, intron, 5' or 3' UTR, ncRNA, repeat, intergenic |
| Conservation | PlantCaduceus suite [34] | binary | 512 | centre-token | centre-token, RC-concatenated | AUPRC | is the centre position evolutionarily conserved |
| Splicing | PGB | binary | 398 | mean / centre-token | centre-token, RC-concatenated | strat. AUROC corr. / strat. AUPRC corr. | is the centre position a splice donor (acceptor) site |
| TIS | PlantCaduceus suite | binary | 512 | centre-token | centre-token, RC-concatenated | AUPRC (*A. thaliana*); strat. AUPRC corr. (cross-species) | is the centre position a translation initiation site |
| TTS | PlantCaduceus suite | binary | 512 | centre-token | centre-token, RC-concatenated | AUPRC (*A. thaliana*); strat. AUPRC corr. (cross-species) | is the centre position a translation termination site |
| PRO-seq | PGB | binary | 1,000 | mean | mean | AUPRC | is the window transcriptionally active in PRO-seq |
| LLR mutation effect | this work | zero-shot ranking | $W$ context | masked LLR<sup>b</sup> | causal LLR | AUROC | separate deleterious from neutral variants |
| Causal variant discovery | this work (Section 4.10) | zero-shot ranking | $W$ context | masked signed LLR<sup>b</sup> | causal signed LLR | recall-AUC | rank the validated causal SNP among 100 background SNPs of its locus |

**Supplementary Table S6:** Dataset sizes and subsampling behind every metric of the suite. One row per logged metric: the 22 metrics entering *S*_bal_ and the six cross-species TIS and TTS metrics held out of it (Section 4.3.4). Original rows are the full dataset splits as released (positives in parentheses; the PGB splits from the benchmark release [31], the TIS, TTS and conservation splits from the PlantCaduceus dataset card [34]). For TIS, TTS and splicing the *Arabidopsis thaliana* probes train on *A. thaliana* and every species column scores the same probe, so the train split is repeated across the species rows of a family. The two probe-sampling columns give the rows presented to the probe, as train / test: <u>uni.</u> is a uniform random subsample at the stated cap, <u>strat.</u> a stratified draw of 500 positives and 4,500 negatives re-weighted to the true prevalence, and <u>prev.</u> the prevalence-matched splicing draws of Section 4.3.5 (2,400 positives and 97,600 negatives for donor, 3,500 and 96,500 for acceptor). Caps that exceed the split size leave it complete (PRO-seq). *^a^* genomic region datasets tile each genome into non-overlapping 512 bp windows fully covered by a single annotation class and split train and test by holding out chromosomes, so row counts follow from the annotation rather than the protocol; *^b^* multi-label task, positives are per track; *^c^* documented Ensembl variants classified by VEP consequence into deleterious and neutral, with three frequency-matched neutral variants drawn per deleterious variant, all scored zero-shot; *^d^* a fixed subset of 29 curated studies from four species (13 *A. thaliana*, 11 *Oryza sativa*, 4 *Triticum aestivum*, 1 *Sorghum bicolor* ) drawn from the curated causal-variant benchmark (Section 4.10): each causal SNP is ranked by signed LLR against 100 background SNPs of its 100 kbp locus (50 closest in allele frequency and 50 nearest by position when allele frequencies are available, otherwise the 100 nearest) and the family contributes a single recall-AUC scalar.

| Family | Species (test split) | Train rows (pos.) | Test rows (pos.) | Probe rows $S_{\text{bal}}^{\text{train}}$ | Probe rows $S_{\text{bal}}^{\text{test}}$ | In $S_{\text{bal}}$ |
| --- | --- | --- | --- | --- | --- | --- |
| Genomic region | *A. thaliana* | —<sup>a</sup> | —<sup>a</sup> | 20,000 / 5,000 uni. | 20,000 / 5,000 uni. | yes |
|  | *O. sativa* | —<sup>a</sup> | —<sup>a</sup> | 20,000 / 5,000 uni. | 20,000 / 5,000 uni. | yes |
|  | *Z. mays* | —<sup>a</sup> | —<sup>a</sup> | 20,000 / 5,000 uni. | 20,000 / 5,000 uni. | yes |
|  | *S. lycopersicum* | —<sup>a</sup> | —<sup>a</sup> | 20,000 / 5,000 uni. | 20,000 / 5,000 uni. | yes |
| Conservation | sorghum (chr. 10) | 858,086 (429,043) | 38,060 (19,030) | 20,000 / 5,000 uni. | 20,000 / 5,000 uni. | yes |
|  | maize | 858,086 (429,043) | 1,923,999 (947,769) | 20,000 / 5,000 uni. | 20,000 / 5,000 uni. | yes |
| TIS | *A. thaliana* (chr. 5) | 198,591 (24,711) | 57,825 (7,311) | 20,000 / 5,000 uni. | 20,000 / 5,000 uni. | yes |
|  | rice | 198,591 (24,711) | 1,403,089 (2,974) | 20,000 / strat. | 20,000 / strat. | no |
|  | sorghum | 198,591 (24,711) | 3,940,933 (3,214) | 20,000 / strat. | 20,000 / strat. | no |
|  | maize | 198,591 (24,711) | 11,268,672 (3,098) | 20,000 / strat. | 20,000 / strat. | no |
| TTS | *A. thaliana* (chr. 5) | 245,564 (25,112) | 71,826 (7,461) | 20,000 / 5,000 uni. | 20,000 / 5,000 uni. | yes |
|  | rice | 245,564 (25,112) | 3,721,003 (2,974) | 20,000 / strat. | 20,000 / strat. | no |
|  | sorghum | 245,564 (25,112) | 10,448,744 (3,214) | 20,000 / strat. | 20,000 / strat. | no |
|  | maize | 245,564 (25,112) | 29,539,071 (3,098) | 20,000 / strat. | 20,000 / strat. | no |
| PRO-seq | *M. esculenta* | 16,852 (49%) | 812 (49%) | full / full | full / full | yes |
| Splicing | *A. thaliana* donor | 2,588,034 (2.3%) | 377,873 (2.3%) | 20,000 / strat. | 200,000 / prev. | yes |
|  | *A. thaliana* acceptor | 1,704,844 (3.4%) | 250,084 (3.5%) | 20,000 / strat. | 200,000 / prev. | yes |
| Chromatin accessibility | *O. sativa* MH63 | 5,120,000<sup>b</sup> | 14,848<sup>b</sup> | 20,000 / 5,000 uni. | 20,000 / 5,000 uni. | yes |
|  | *O. sativa* ZS97 | 5,120,000<sup>b</sup> | 14,848<sup>b</sup> | 20,000 / 5,000 uni. | 20,000 / 5,000 uni. | yes |
|  | *S. italica* | 5,120,000<sup>b</sup> | 19,968<sup>b</sup> | 20,000 / 5,000 uni. | 20,000 / 5,000 uni. | yes |
|  | *A. thaliana* | 5,120,000<sup>b</sup> | 9,984<sup>b</sup> | 20,000 / 5,000 uni. | 20,000 / 5,000 uni. | yes |
|  | *B. distachyon* | 5,120,000<sup>b</sup> | 14,848<sup>b</sup> | 20,000 / 5,000 uni. | 20,000 / 5,000 uni. | yes |
|  | *S. bicolor* | 5,120,000<sup>b</sup> | 29,952<sup>b</sup> | 20,000 / 5,000 uni. | 20,000 / 5,000 uni. | yes |
|  | *Z. mays* | 6,400,000<sup>b</sup> | 79,872<sup>b</sup> | 20,000 / 5,000 uni. | 20,000 / 5,000 uni. | yes |
| LLR mutation effect | *A. thaliana* | — | all variants<sup>c</sup> | none (zero-shot) | none (zero-shot) | yes |
|  | *Z. mays* | — | all variants<sup>c</sup> | none (zero-shot) | none (zero-shot) | yes |
|  | *S. lycopersicum* | — | all variants<sup>c</sup> | none (zero-shot) | none (zero-shot) | yes |
| Causal variant discovery | 4 species<sup>d</sup> | 29 studies<sup>d</sup> | 101 candidate SNPs per study<sup>d</sup> | none (zero-shot) | none (zero-shot) | yes |

**Supplementary Table S7:** Performance difference between model pairs under two paired bootstraps. ΔS^test^ gives the difference in the reported score 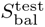 between the “Model” and the “Reference”. Both confidence intervals are 95% percentile intervals over 20,000 paired replicates (Section 4.3.1): “families” resamples the nine task families, “examples” bootstraps the test examples themselves within each of the 22 tasks and species. An asterisk marks an interval that excludes zero. “Families ahead” counts the task families in which the “Model” outperforms the “Reference”.

| Model | Reference | $\Delta S_{\text{bal}}^{\text{test}}$ | 95% CI, families | 95% CI, examples | Families ahead |
| --- | --- | --- | --- | --- | --- |
| Botanic1-S | PlantCAD2-L | +0.0058 | [−0.0063, +0.0237] | [−0.0029, +0.0159] | 4/9 |
| Botanic1-M | PlantCAD2-L | +0.0118 | [−0.0018, +0.0295] | [+0.0039, +0.0217]* | 5/9 |
| Botanic1-L | PlantCAD2-L | +0.0155 | [+0.0016, +0.0336]* | [+0.0078, +0.0252]* | 6/9 |
| Botanic1-XL | PlantCAD2-L | +0.0175 | [+0.0026, +0.0364]* | [+0.0094, +0.0273]* | 7/9 |
| Botanic1-XL | PlantCaduceus | +0.0187 | [+0.0110, +0.0270]* | [+0.0123, +0.0253]* | 9/9 |
| Botanic1-XL | Botanic1-S | +0.0117 | [+0.0042, +0.0226]* | [+0.0067, +0.0171]* | 8/9 |
| Botanic1-M | Botanic1-S | +0.0060 | [+0.0002, +0.0141]* | [+0.0005, +0.0117]* | 7/9 |
| Botanic1-L | Botanic1-M | +0.0037 | [−0.0007, +0.0076] | [−0.0008, +0.0081] | 7/9 |
| Botanic1-XL | Botanic1-L | +0.0020 | [−0.0027, +0.0076] | [−0.0040, +0.0081] | 4/9 |
| Botanic1-S | Carbon-8B | +0.0353 | [+0.0149, +0.0578]* | [+0.0249, +0.0460]* | 8/9 |
| Botanic1-S | Evo 2-7B | +0.0424 | [+0.0187, +0.0677]* | [+0.0292, +0.0564]* | 8/9 |

**Supplementary Figure S18.**
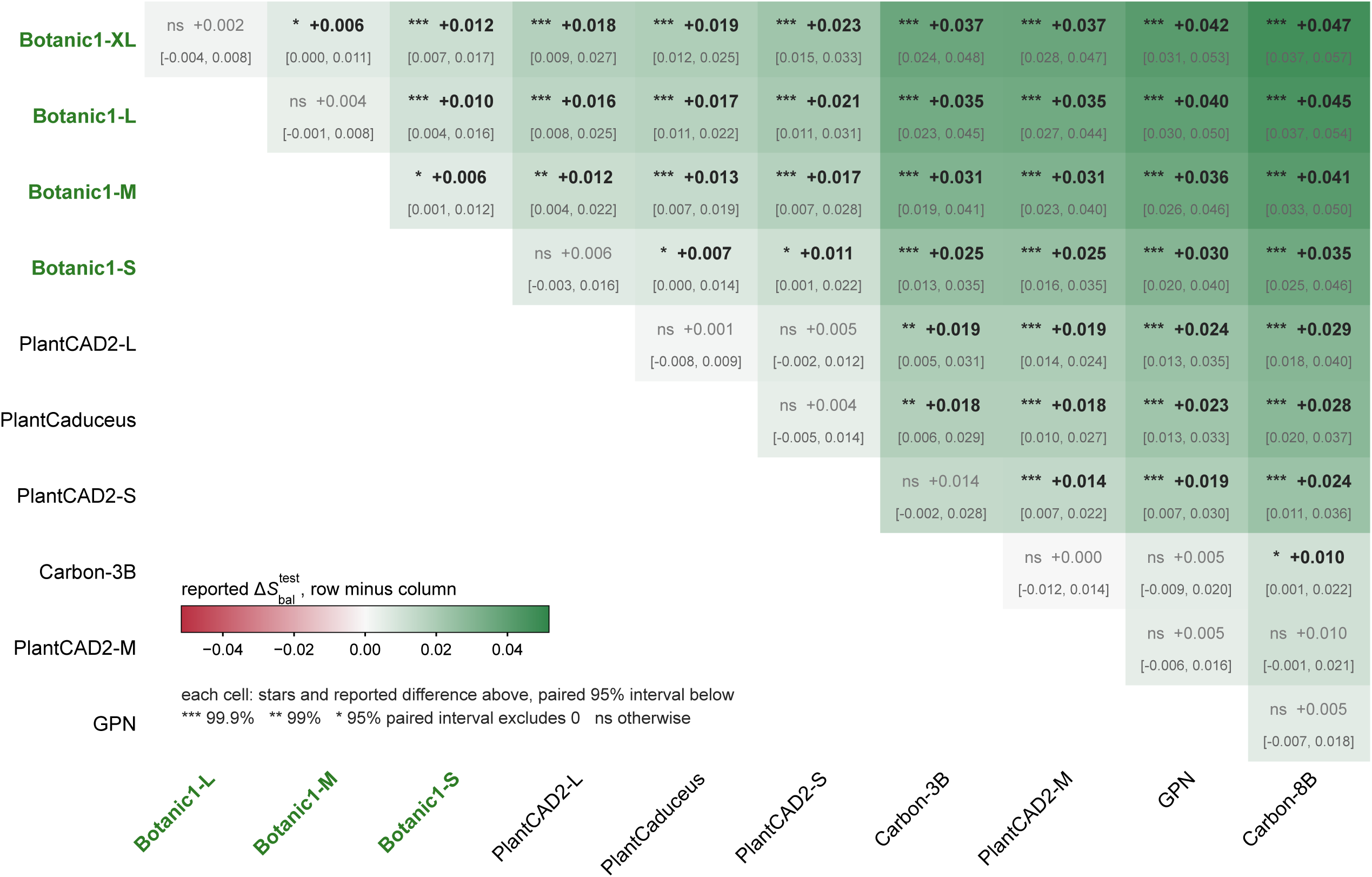
Every pair of the leading models under the per-example paired bootstrap. The 11 models with a reported 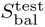 0.72, in leaderboard order. Each cell gives the difference in 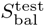 between the row model and the column model, the paired 95% interval and the significance level of the difference (*^∗^* 95%, *^∗∗^* 99%, *^∗∗∗^* 99.9%; ns otherwise). The shade is the reported difference. Figure 3e shows the column of this matrix against PlantCAD2-L, extended to every named model.

**Supplementary Figure S19.**
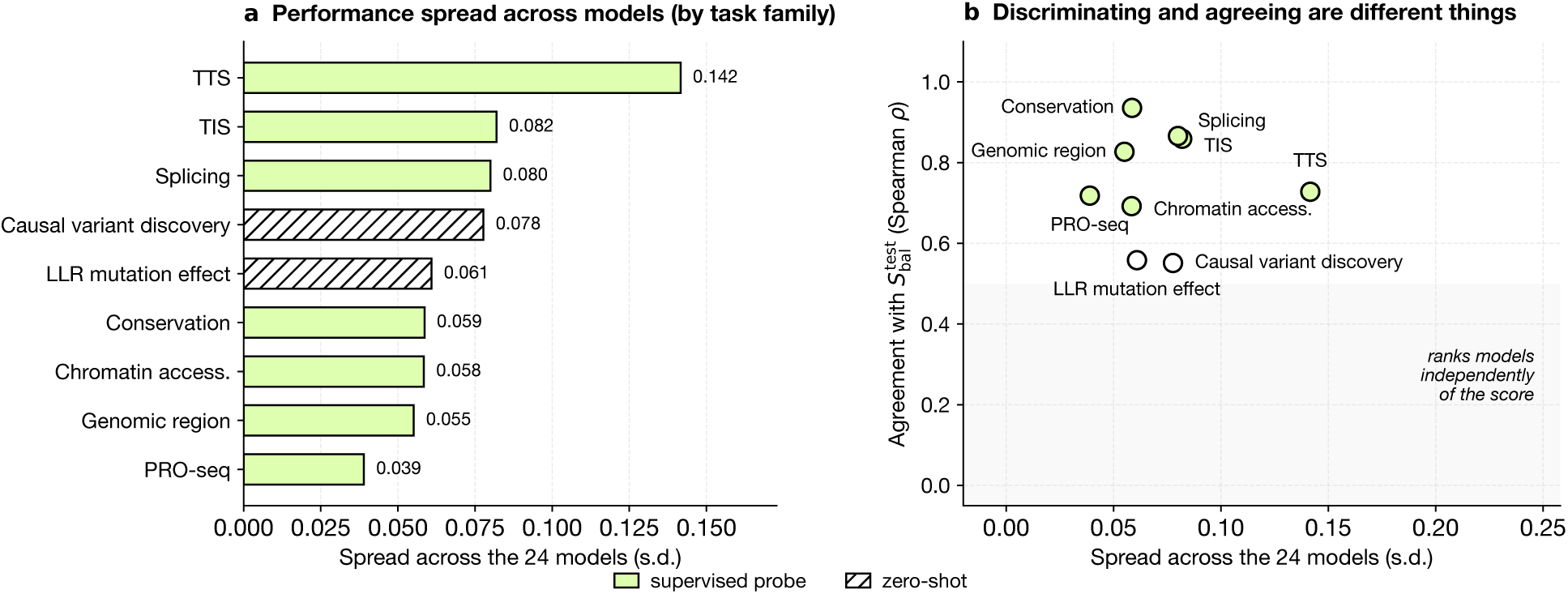
How much each task family separates models, and whether it agrees with 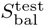. Every family is reduced to its species-mean per model, its contribution to *S*_bal_ under Equation (1), across the 24 reported models; MarinDNA-1B is excluded, its 255 bp context admitting only 8 of the 22 metrics. **a**, Standard deviation of that family score across the cohort: how far apart the family places models. **b**, The same spread against Spearman ρ between the family’s ordering and 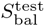. The shaded band marks families whose ordering is largely independent of the score. Frozen embeddings probes are filled, the two zero-shot families hatched in **a** and open in **b**. LLR and causal variant discovery (subset) have ordinary spread but the lowest agreement of any family.

**Supplementary Figure S20.**
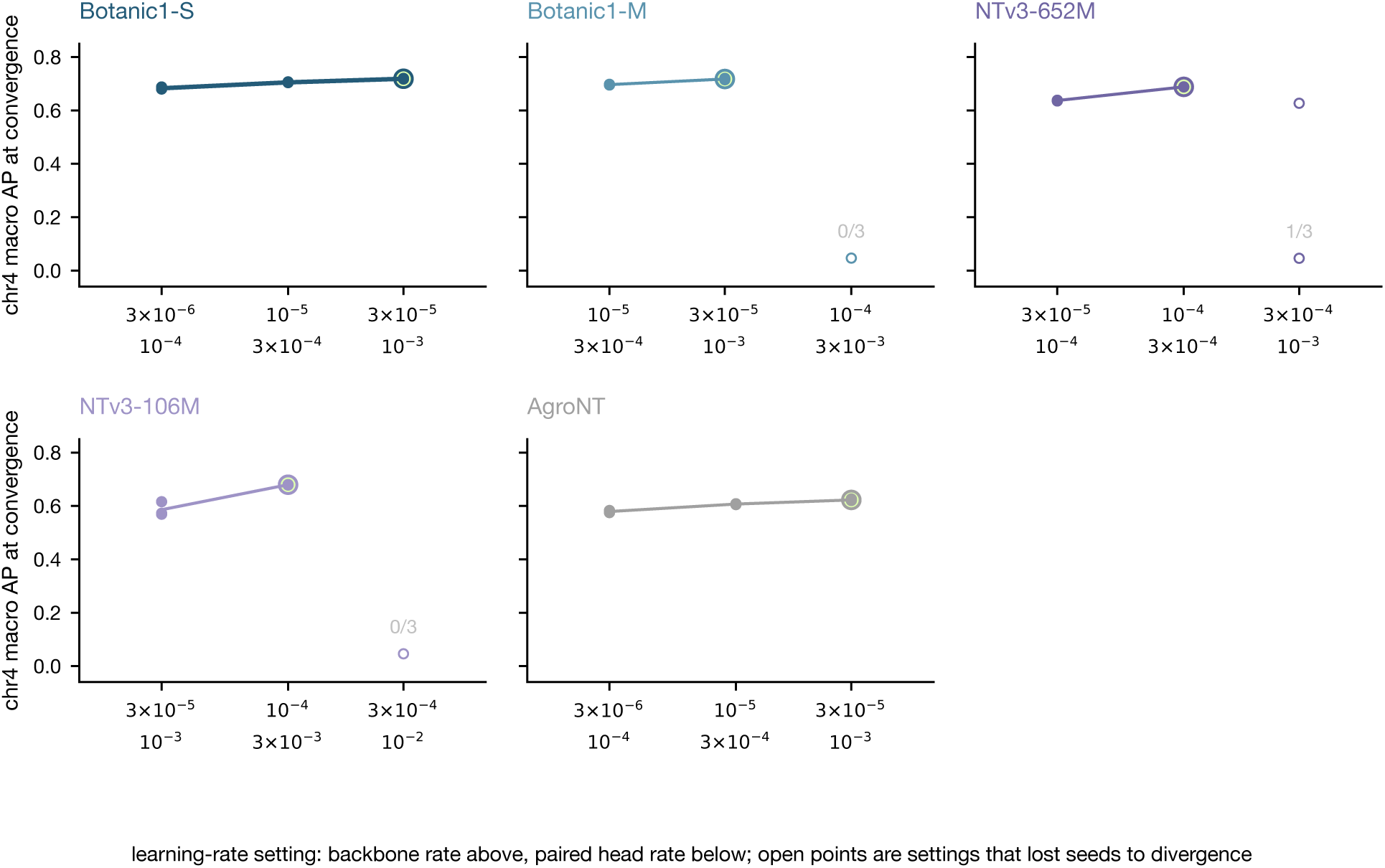
Transcription factor binding prediction learning-rate screen. All nine runs per model: three learning-rate settings at seeds 17, 29, and 43. The X ticks give one setting each, with the backbone rate above and its paired head rate below. Filled points are the three seeds of an eligible setting, the ring marks the selected setting. Open points are settings with which some seeds do not converge, with the annotation to show the number of seeds that stay finite. a diverged run reaches the mean family prevalence, 0.047.

**Supplementary Figure S21.**
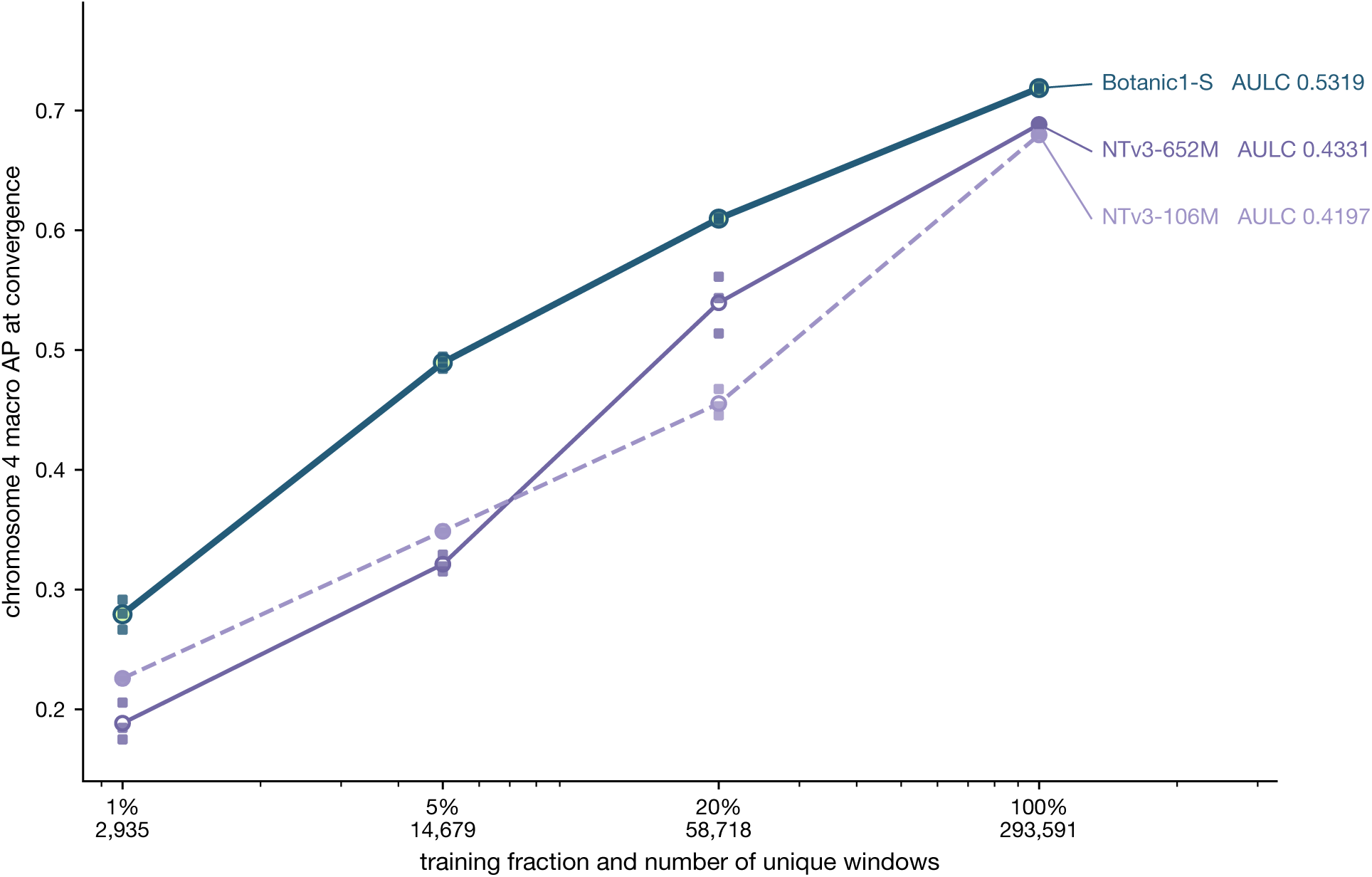
Transcription factor binding performance as labelled training data increases. Each point is a seeded experiment plus their average, each trained until convergence (see Section 4.6). AULC is the trapezoidal area on a log_10_ data axis, divided by the axis range.

**Supplementary Figure S22.**
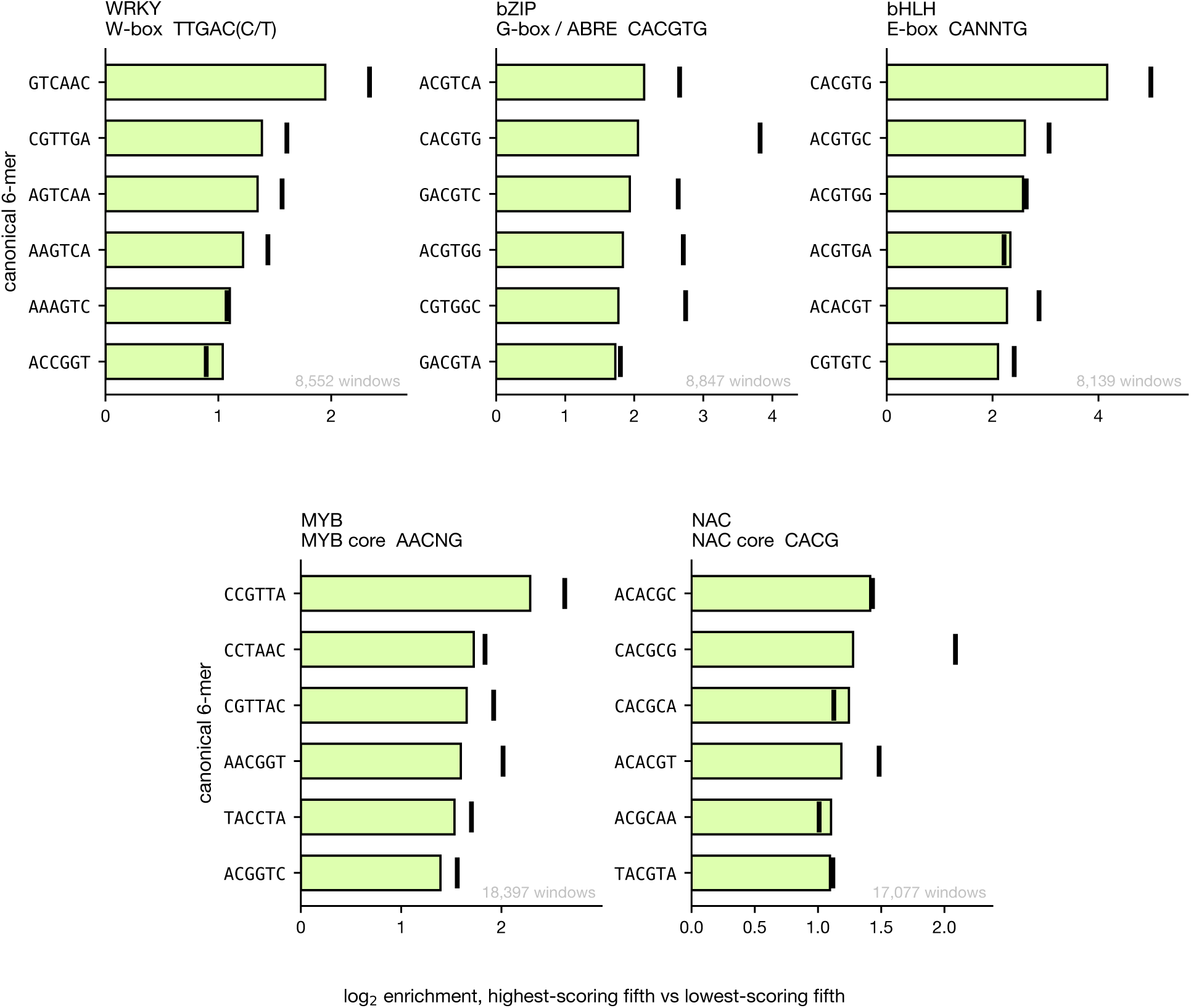
Canonical 6-mers enrichment in weakly vs strongly positive windows. Within each family’s positive-labelled windows, the highest-scoring fifth is compared with the lowest-scoring fifth on canonical 6-mer presence, ranking by each model’s score averaged over three seeds. Bars show the enrichment for Botanic1-S, and ticks show DeepCistrome enrichment. Each 6-mer is counted in either orientation and labelled with the first of the two in lexicographical order (for example, GTCAAC is the reverse complement of GTTGAC and so contains the WRKY W-box core). Below each TF family we show the main canonical binding element.

**Supplementary Table S8:**
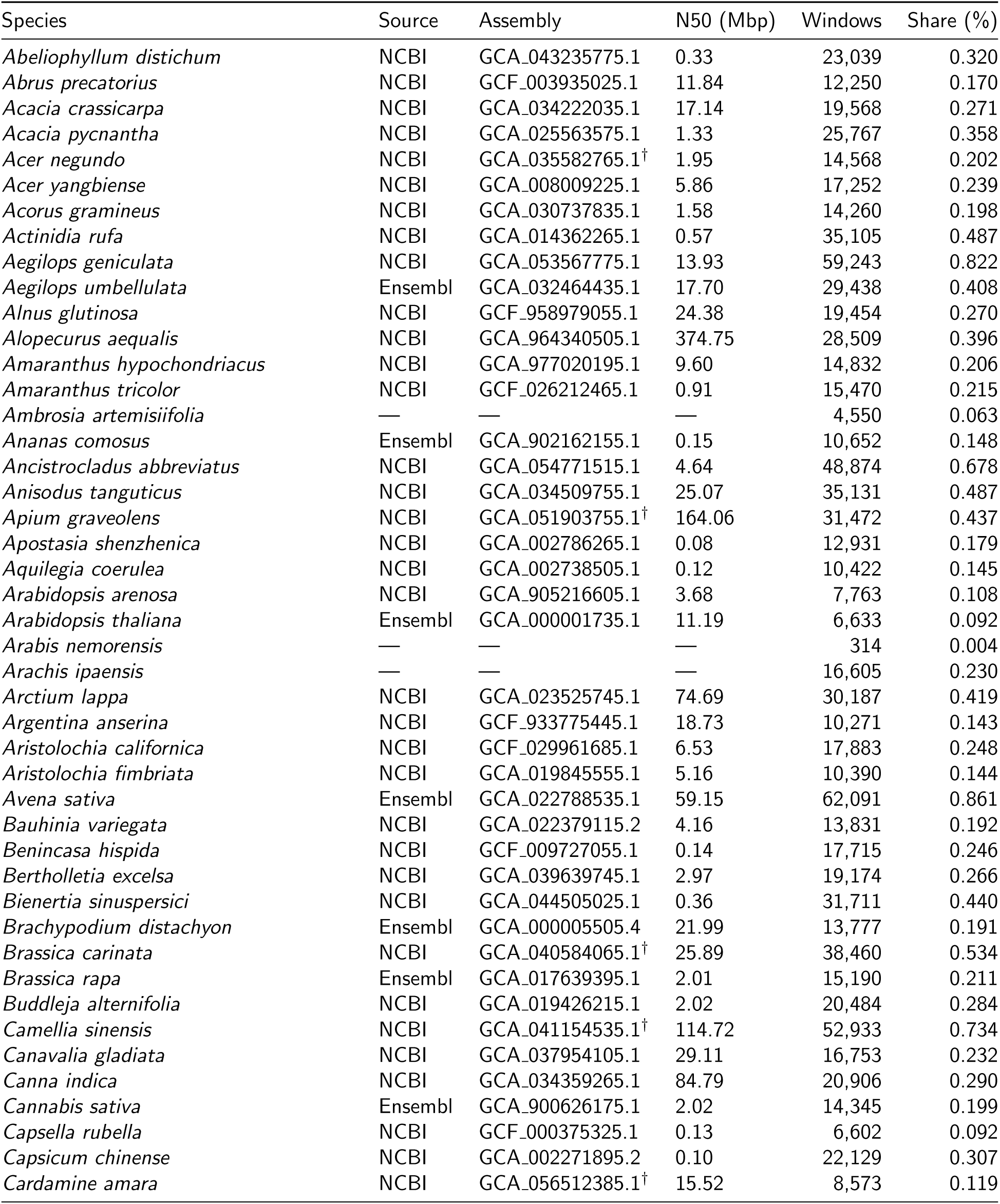
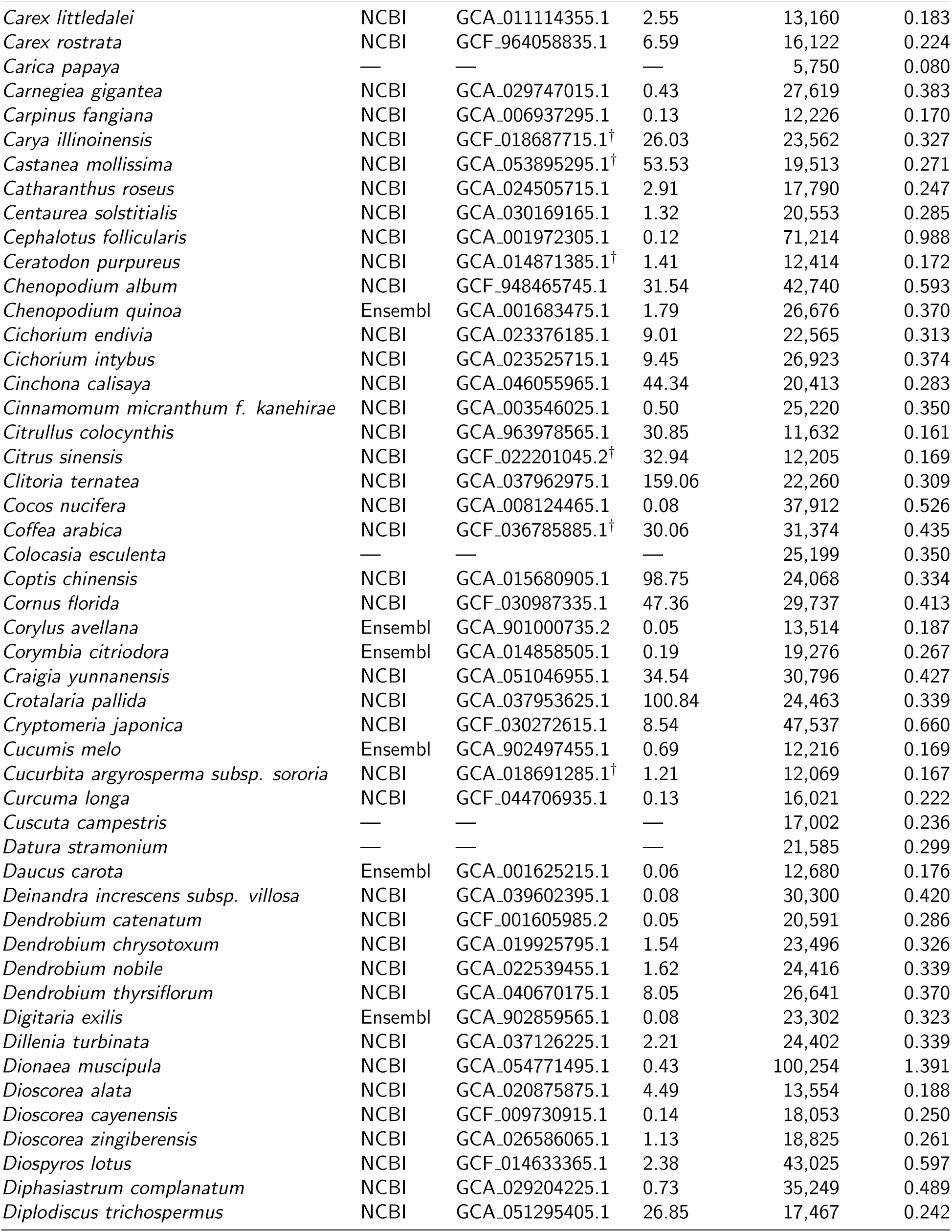
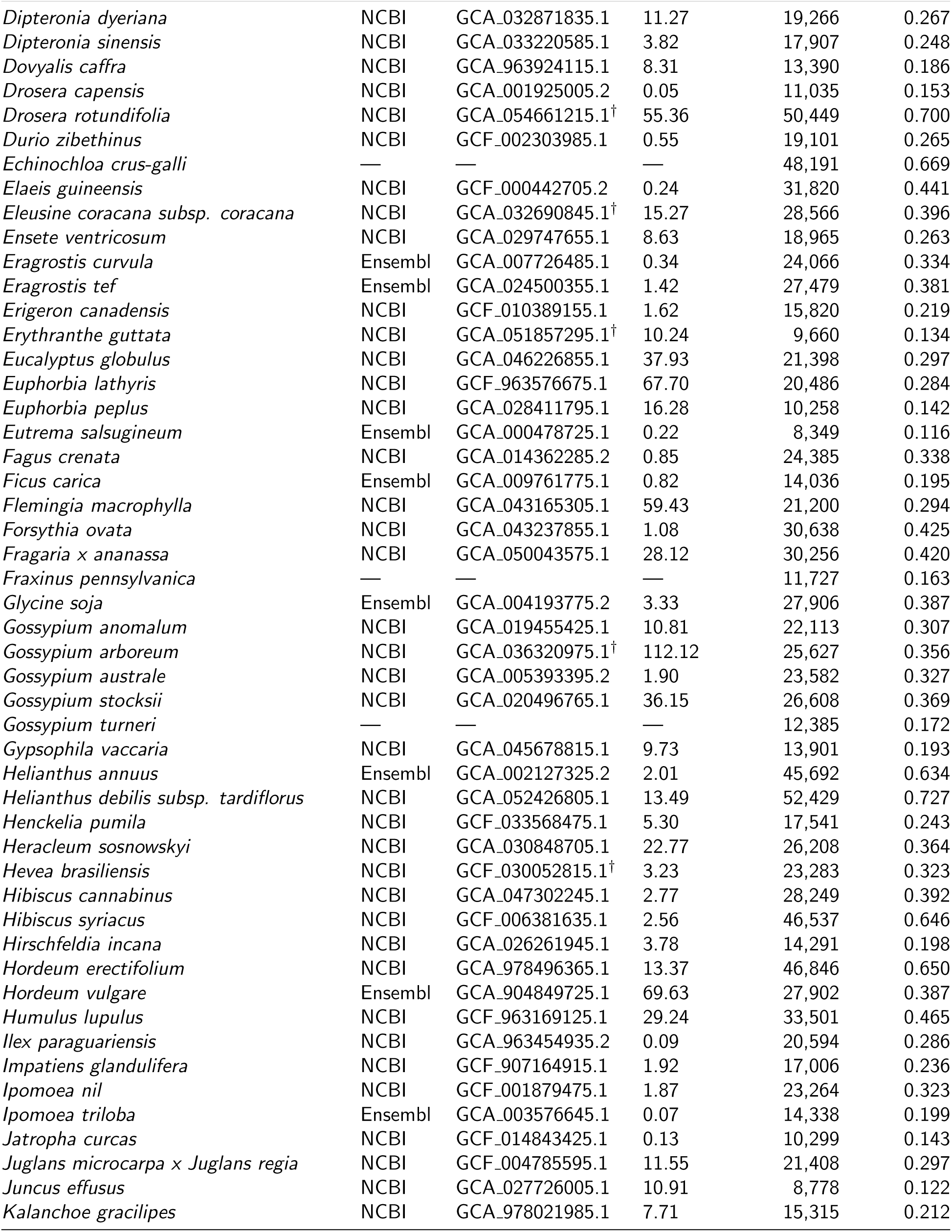
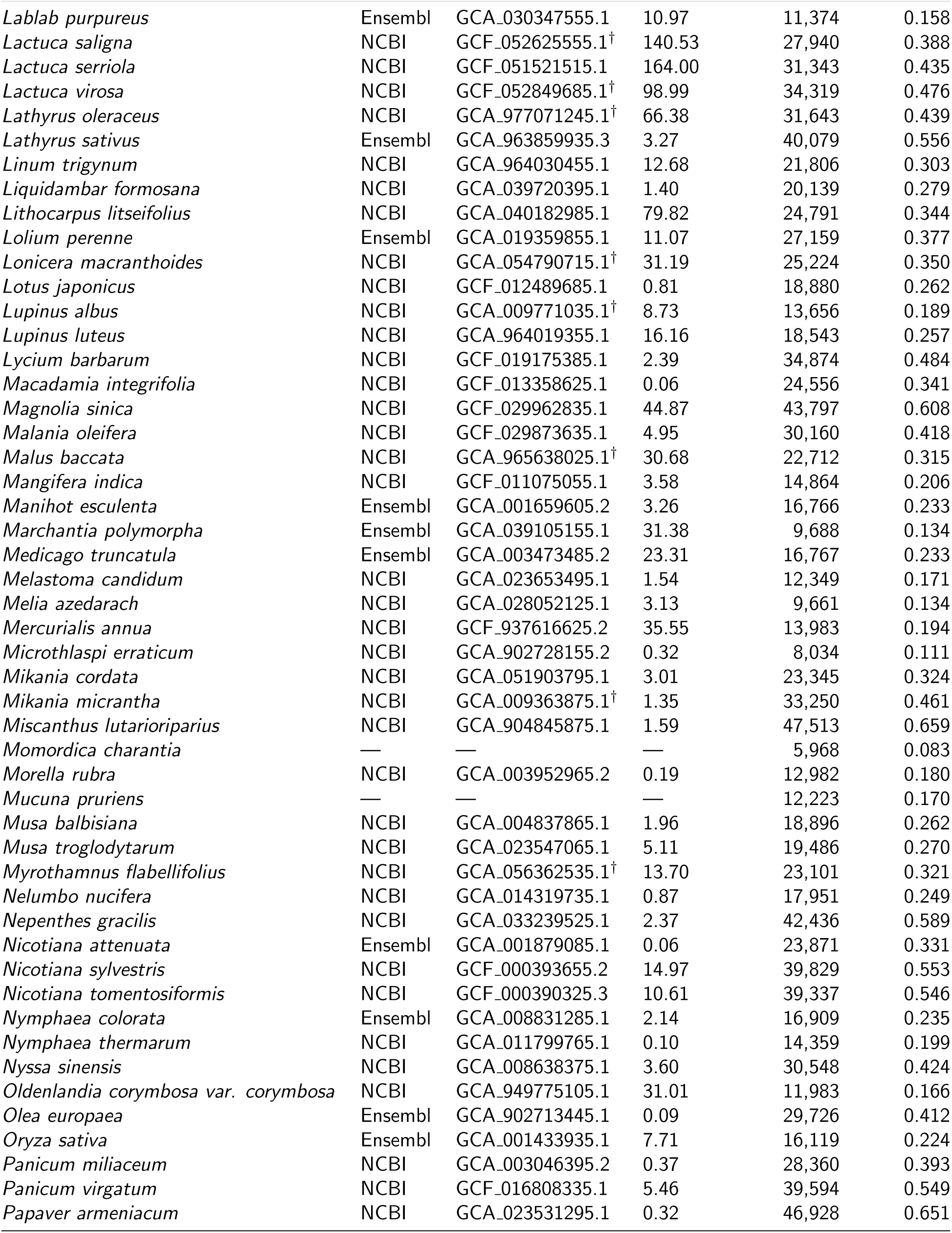
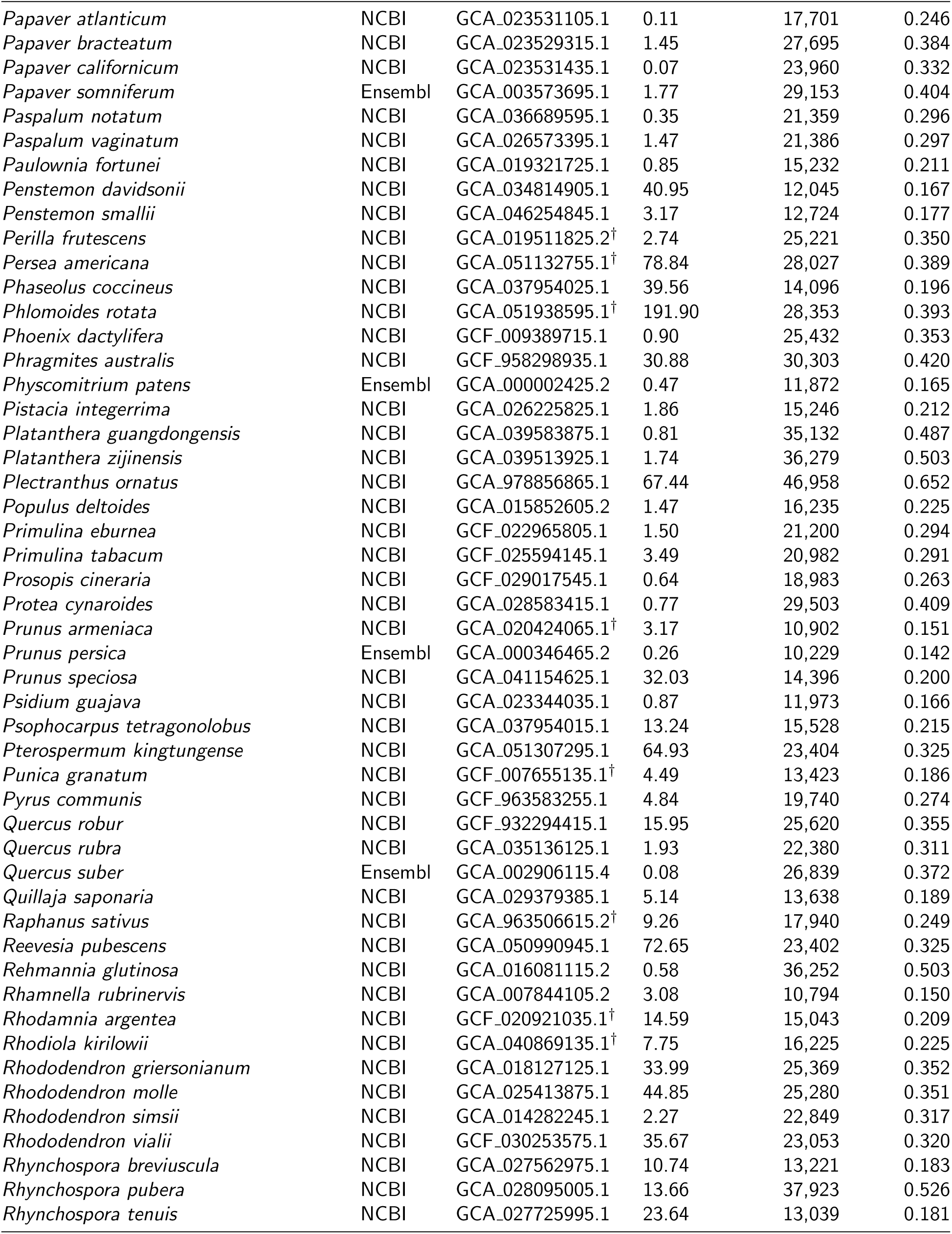
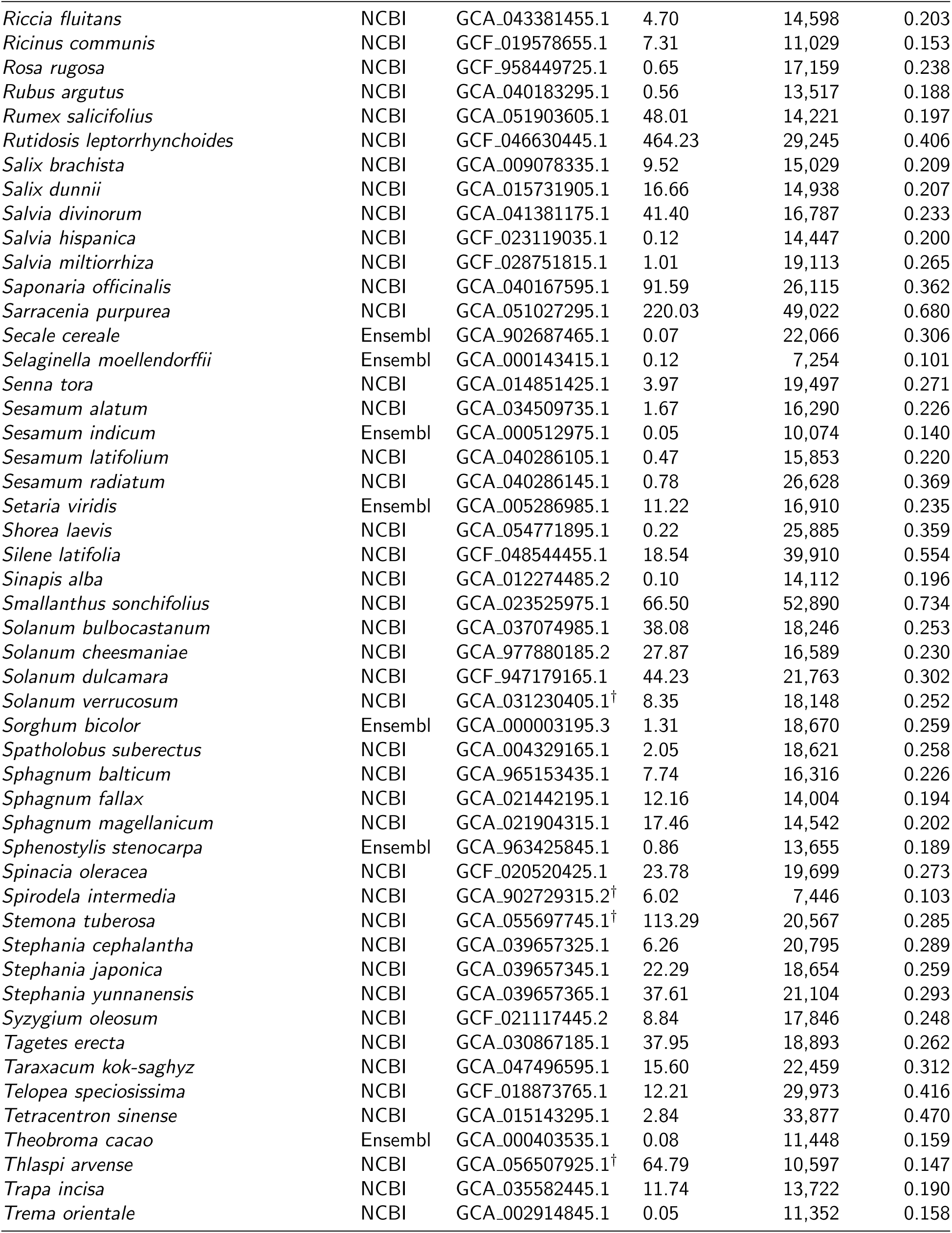
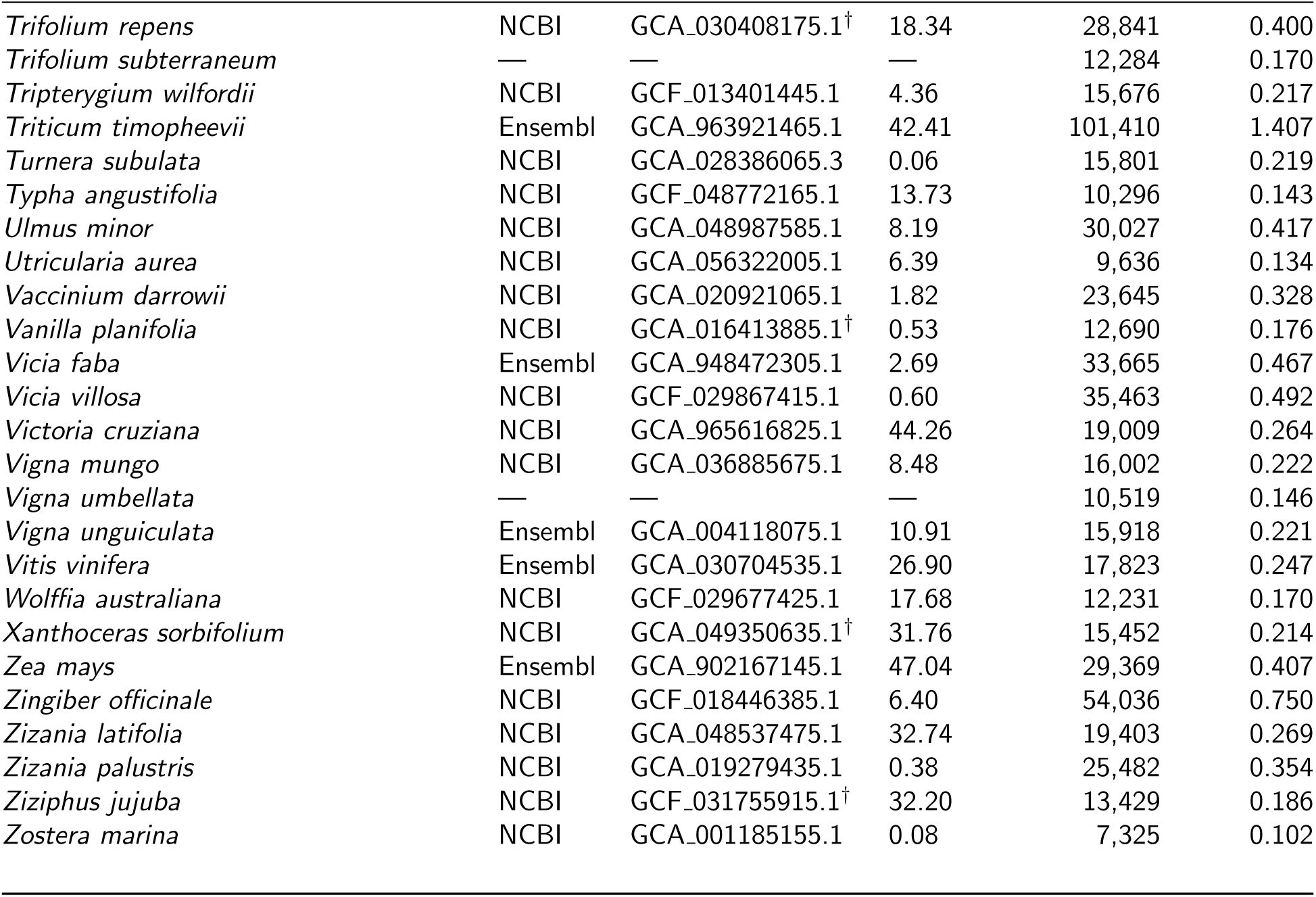
Genome assemblies in the champion pre-training corpus. Contig N50 in Mbp; “Windows” is the number of 8,192 bp training windows drawn from that species and “Share” its percentage of the 7,207,506-window corpus. Entries marked *^†^* have more than one assembly passing the quality filters, and the representative shown is the annotated assembly with the highest contig N50, matching the selection rule used by the pipeline. Dashes mark species whose representative assembly cannot be resolved from the released metadata.

**Supplementary Table S9:** Fine-tuning hyperparameter grid. Values searched for each task backbone adaptation regime. Every combination shown is run and scored on a validation split; the selected combination is the one reproduced in the YAML configs of Section 4.3.8. a dash means the hyperparameter is held fixed for that cell rather than searched. Held fixed throughout: weight decay 10*^−^*^4^, warmup ratio 0.05, no class weighting, early stopping on validation loss. Epochs are 4 for the two regression tasks and 3 for the two classification tasks.

| Task | Backbone | Regime | learning rate | head LR | batch | grad accum | pooling |
| --- | --- | --- | --- | --- | --- | --- | --- |
| Terminator strength | BiMamba2 | full-FT | {1e-05, 3e-05, 0.0001, 0.0003} | — | 8 | {1, 4} | all |
|  | BiMamba2 | LoRA | {0.0001, 0.0005, 0.001, 0.003} | — | 8 | {1, 4} | all |
|  | BiMamba2 | IA<sup>3</sup> | {0.001, 0.003, 0.005, 0.01} | — | 8 | {1, 4} | all |
|  | AgroNT | full-FT | {1e-05, 3e-05, 0.0001, 0.0003} | — | 8 | {1, 4} | all |
|  | AgroNT | LoRA | {0.0001, 0.0005, 0.001, 0.003} | — | 8 | {1, 4} | all |
|  | AgroNT | IA<sup>3</sup> | {0.001, 0.003, 0.005, 0.01} | — | 8 | {1, 4} | all |
|  | PlantCAD2-L | full-FT | {1e-06, 3e-06, 1e-05, 3e-05, 0.0001, 0.0003} | — | 8 | {1, 4} | all |
|  | PlantCAD2-L | LoRA | {1e-05, 3e-05, 0.0005, 0.001, 0.003} | — | 8 | {1, 4} | all |
|  | PlantCAD2-L | IA<sup>3</sup> | {0.0001, 0.0003, 0.003, 0.005, 0.01} | — | 8 | {1, 4} | all |
| Promoter strength | BiMamba2 | full-FT | {1e-05, 3e-05, 0.0001, 0.0003} | — | 8 | {1, 4} | all |
|  | BiMamba2 | LoRA | {0.0001, 0.0005, 0.001, 0.003} | — | 8 | {1, 4} | all |
|  | BiMamba2 | IA<sup>3</sup> | {0.001, 0.003, 0.005, 0.01} | — | 8 | {1, 4} | all |
|  | AgroNT | full-FT | {1e-05, 3e-05, 0.0001, 0.0003} | — | 8 | {1, 4} | all |
|  | AgroNT | LoRA | {0.0001, 0.0005, 0.001, 0.003} | — | 8 | {1, 4} | all |
|  | AgroNT | IA<sup>3</sup> | {0.001, 0.003, 0.005, 0.01} | — | 8 | {1, 4} | all |
|  | PlantCAD2-L | full-FT | {1e-06, 3e-06, 1e-05, 3e-05, 0.0001, 0.0003} | — | 8 | {1, 4} | all |
|  | PlantCAD2-L | LoRA | {1e-05, 3e-05, 0.0001, 0.0005, 0.001, 0.003} | — | 8 | {1, 4} | all |
|  | PlantCAD2-L | IA<sup>3</sup> | {0.001, 0.003, 0.005, 0.01} | — | 8 | {1, 4} | all |
| poly(A) site | BiMamba2 | full-FT | {1e-05, 3e-05, 0.0001, 0.0003} | 0.005 | 8 | {1, 4} | attn |
|  | BiMamba2 | LoRA | {0.0001, 0.0005, 0.001, 0.003} | 0.005 | 8 | {1, 4} | attn |
|  | BiMamba2 | IA<sup>3</sup> | {0.001, 0.003, 0.005, 0.01} | 0.005 | 8 | {1, 4} | attn |
|  | AgroNT | full-FT | {1e-05, 3e-05, 0.0001, 0.0003} | — | 8 | {1, 4} | cls_only |
|  | AgroNT | LoRA | {0.0001, 0.0005, 0.001, 0.003} | — | 8 | {1, 4} | cls_only |
|  | AgroNT | IA<sup>3</sup> | {0.001, 0.003, 0.005, 0.01} | — | 8 | {1, 4} | cls_only |
|  | PlantCAD2-L | full-FT | {1e-06, 3e-06, 1e-05} | 0.001 | 8 | {1, 4} | attn |
|  | PlantCAD2-L | LoRA | {1e-05, 3e-05, 0.0001, 0.0005, 0.001, 0.003} | 0.001 | 8 | {1, 4} | attn |
|  | PlantCAD2-L | IA<sup>3</sup> | {0.001, 0.003, 0.005, 0.01} | 0.001 | 8 | {1, 4} | attn |
| lncRNA | BiMamba2 | full-FT | {1e-05, 3e-05, 0.0001, 0.0003} | — | 8 | {1, 4} | all |
|  | BiMamba2 | LoRA | {0.0001, 0.0005, 0.001, 0.003} | 0.005 | 8 | {1, 4} | all |
|  | BiMamba2 | IA<sup>3</sup> | {0.001, 0.003, 0.005, 0.01} | 0.005 | 8 | {1, 4} | all |
|  | AgroNT | full-FT | {1e-05, 3e-05, 0.0001, 0.0003} | — | 8 | {1, 4} | all |
|  | AgroNT | LoRA | {0.0001, 0.0005, 0.001, 0.003} | — | 8 | {1, 4} | all |
|  | AgroNT | IA<sup>3</sup> | {0.001, 0.003, 0.005, 0.01} | — | 8 | {1, 4} | all |
|  | PlantCAD2-L | full-FT | {1e-06, 1e-05, 3e-05, 0.0001, 0.0003} | — | 1 | {4, 8, 16, 32} | all |
|  | PlantCAD2-L | LoRA | {1e-05, 0.0001, 0.0005, 0.001, 0.003} | 0.001 | 1 | {4, 8, 16, 32} | all |
|  | PlantCAD2-L | IA<sup>3</sup> | {0.001, 0.003, 0.005, 0.01} | 0.001 | 1 | {4, 8, 16, 32} | all |

**Supplementary Figure S23.**
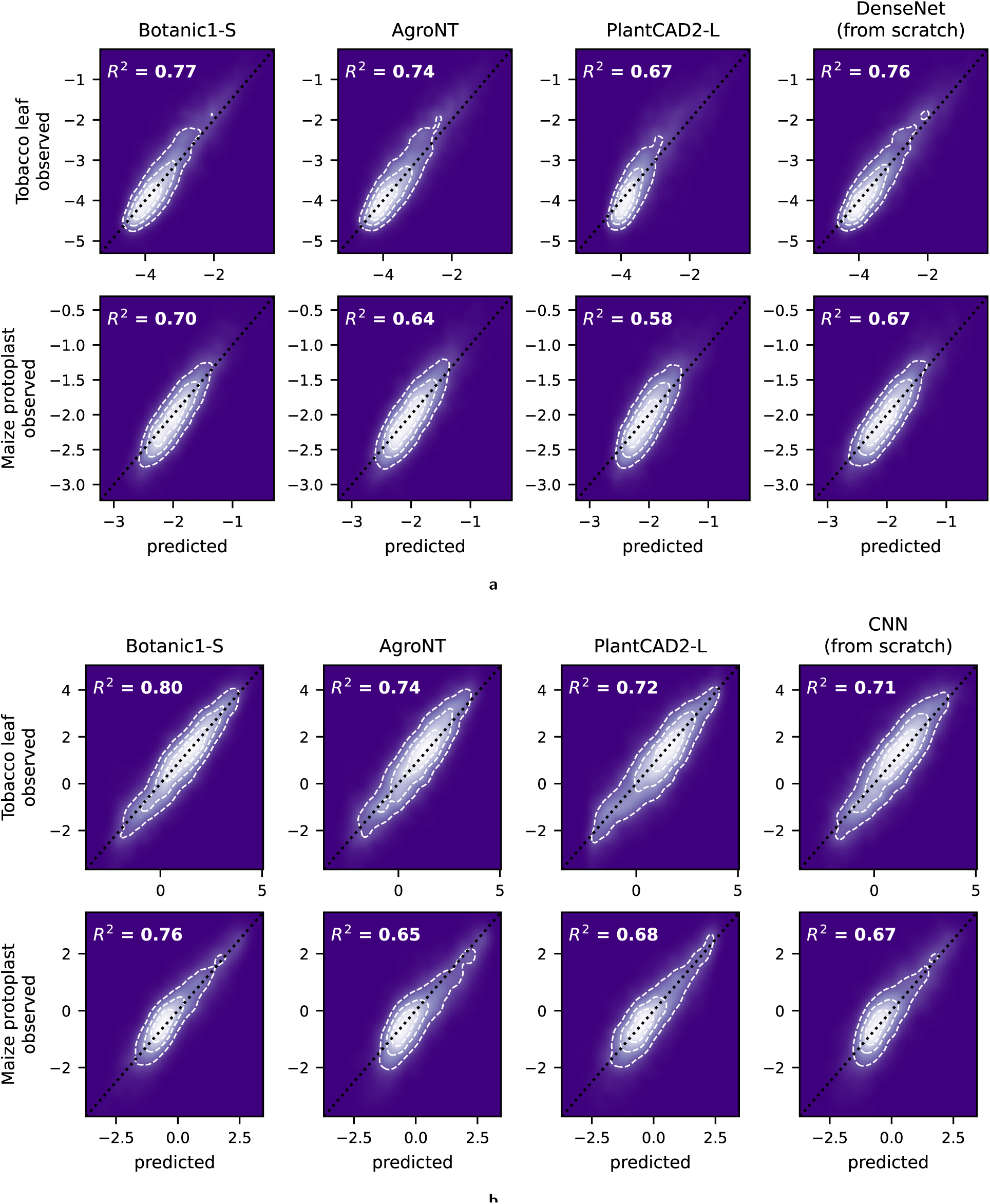
Predicted versus observed values. Predicted activity against observed activity under full fine-tuning and for the from-scratch model, as a smoothed density with white quartile contours, the dotted identity line *y* = *x*, and each panel’s R^2^. **a**, Terminator, 5,309 test sequences, one model predicting two outputs. **b**, Promoter, trained split per tissue, so the two rows are separate models on non-overlapping test sets (7,154 tobacco leaf, 7,595 maize protoplast). Every model is well calibrated mid-range and regresses towards the mean at the strong end.

**Supplementary Figure S24.**
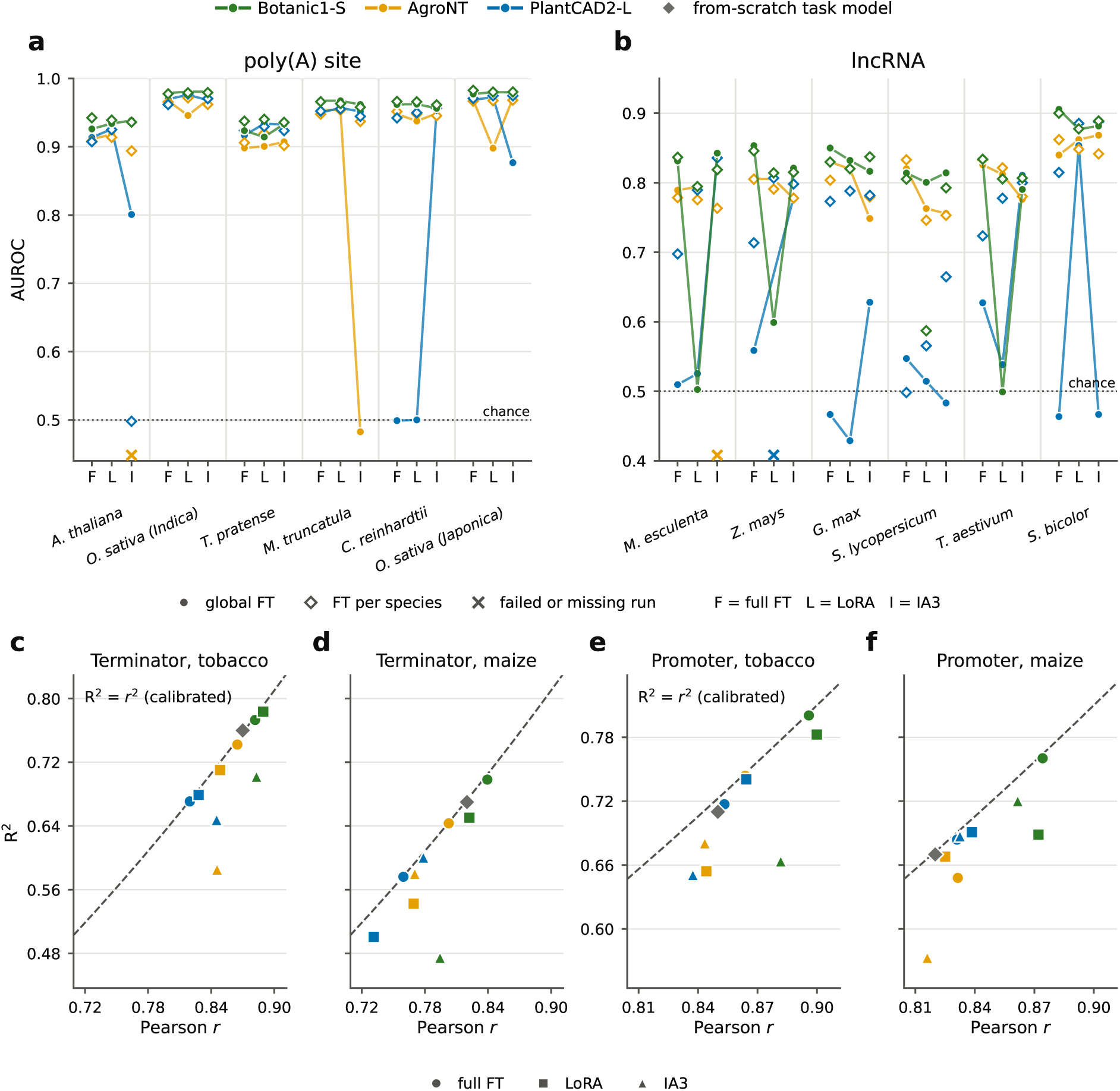
Fine-tuning across all four tasks. Held-out test performance per backbone *×* adaptation regime. **a**,**b**, The two classification tasks per species: held-out AUROC for each backbone across the three regimes (full fine-tuning, LoRA, IA^3^), with filled dots for global fine-tuning, open diamonds for per-species tuning, and crosses for failed or never-run configurations; the Botanic1-S series is drawn in front. **c** to **f**, The two regression tasks, one panel per task and assay system: Pearson *r* against R^2^, one point per backbone and fine-tuning regime, with the from-scratch task model as a grey diamond. The dashed curve marks R^2^ = *r*^2^; vertical distance below or above is the calibration gap.

**Supplementary Figure S25.**
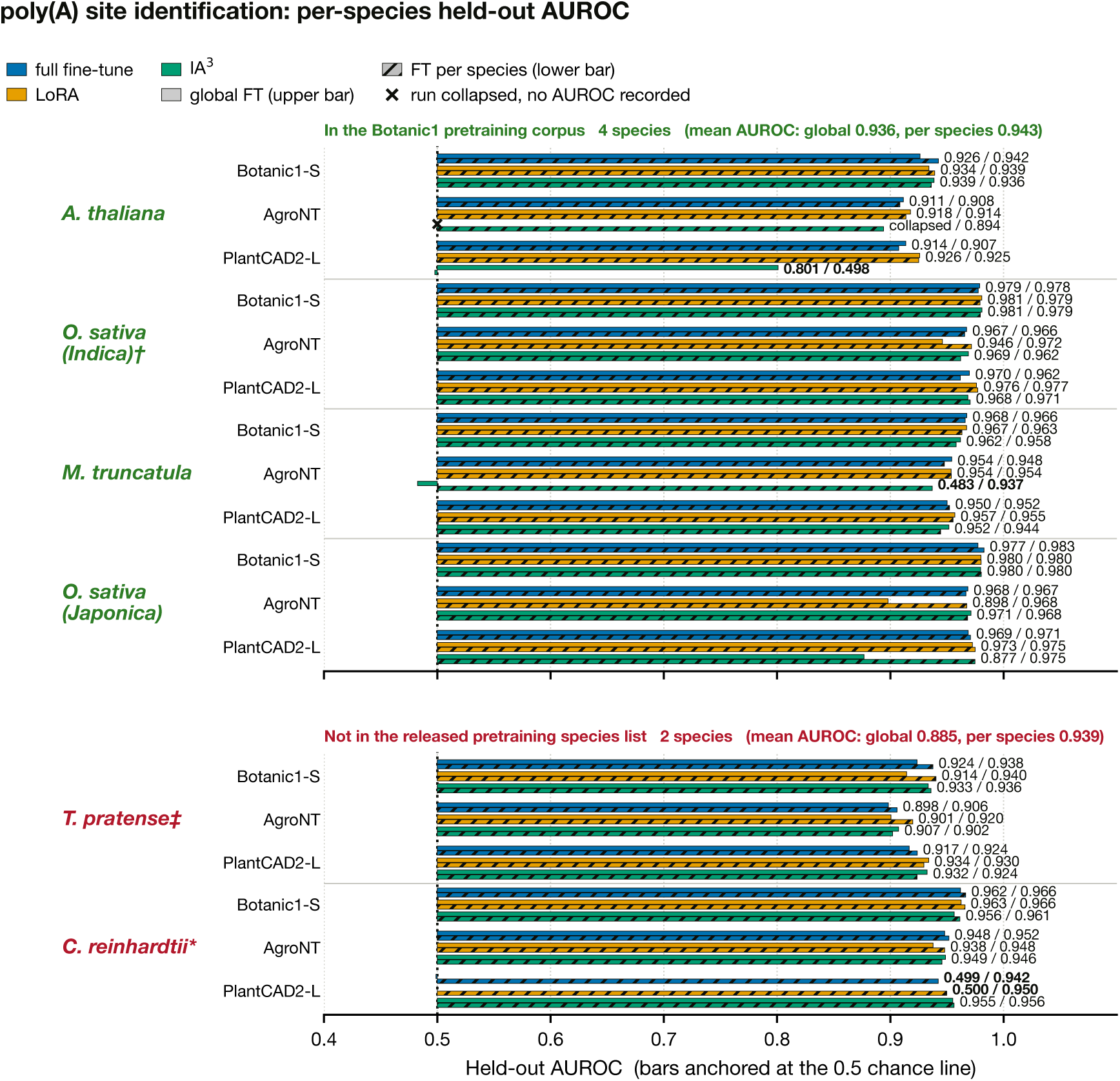
poly(A) site prediction per species. Held-out AUROC for each backbone × fine-tuning adaptation regime on each of the six species. Each fine-tuning regime shows two stacked bars: the upper solid bar is global FT (one shared hyperparameter set across species) and the lower hatched bar is FT per species (the hyperparameters are tuned for that species). Bars start at the 0.5 chance line, so a sub-chance performance is reported on the left of the vertical bar; cells with no AUROC are drawn as a cross at chance. In the underlying release table each poly(A) subtask is anonymised as s0 to s5, corresponding to the subtask indices of the Plant Genomic Benchmark (PGB); the manuscript names no poly(A) species, so the mapping from index to species rests on that ordering convention alone and is inferred rather than externally verified. Species names are printed in green when the species is in the pre-training corpus and in red when it is absent from the released pre-training species list. Symbols on species names flag partial matches to the pre-training corpus: † marks a different subspecies of a species that is in the corpus (*Oryza sativa*, GCA 001433935.1, the Japonica reference); ‡ marks a species absent from the corpus but represented by a congener (*Trifolium repens*, *T. subterraneum*); * marks *Chlamydomonas reinhardtii*, a chlorophyte alga excluded by the corpus’s embryophyta-only filter and therefore the one certain non-member. PlantCAD2-L full fine-tuning on *Medicago truncatula* has four rows in the source table with no recorded metric. “Not in the released pre-training species list” means absent from the 320-species stage manifest; 6 of the 326 corpus genomes (4 held out and 2 left windowless by filtering) are not enumerated in that manifest, so absence from the list is not proof of exclusion from training.

**Supplementary Figure S26.**
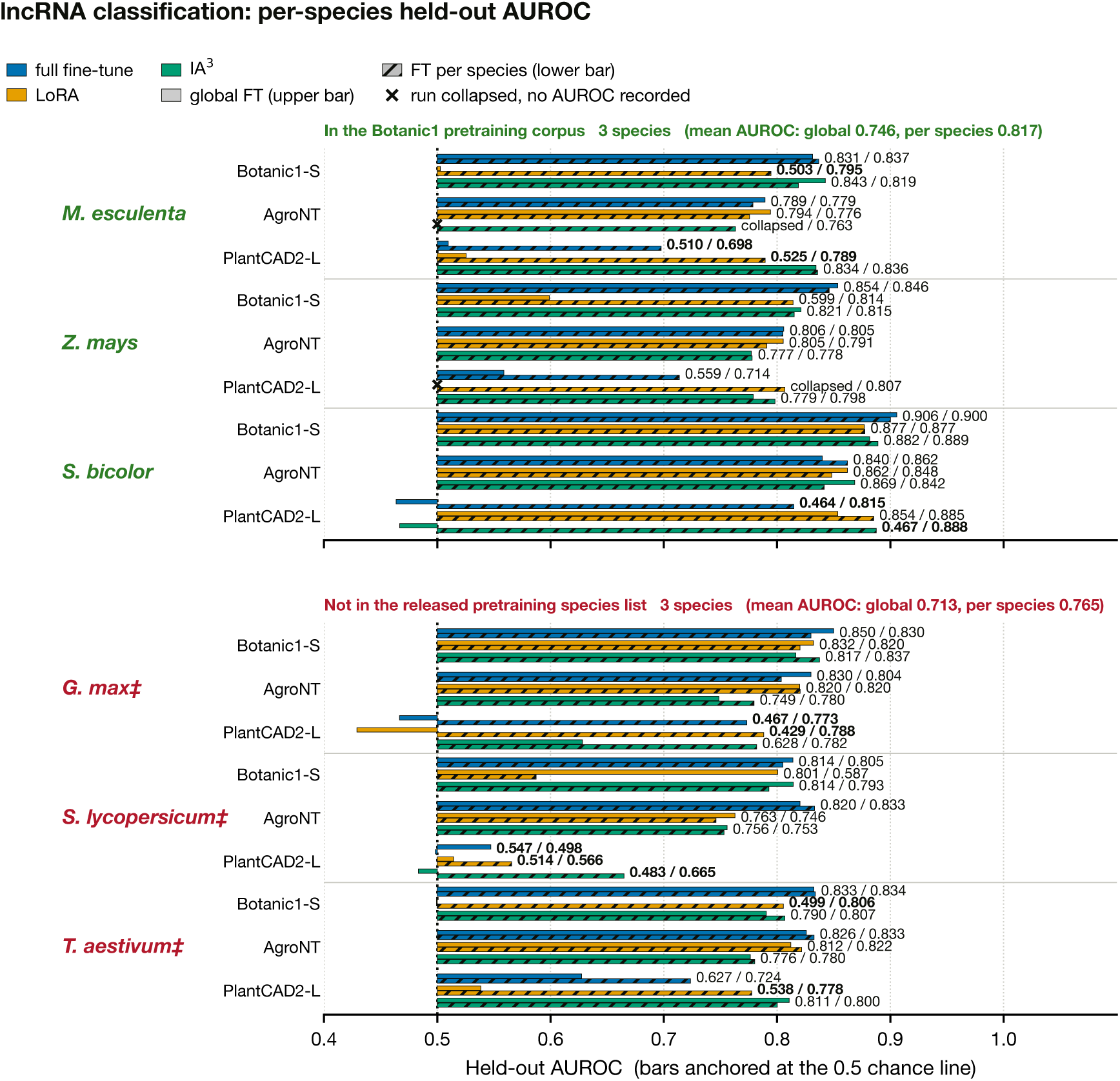
lncRNA classification, per species. Held-out AUROC for each backbone × adaptation regime on each of the 6 species. The upper solid bar of each regime is global FT and the lower hatched bar is FT per species. The in-corpus species hold a small edge (0.746 versus 0.713 mean AUROC over all cells under global FT). The one non-finite cell (PlantCAD2-L LoRA on *Zea mays*) is the collapse discussed in the main text; per-species tuning lifts PlantCAD2-L markedly on this task. In the underlying release table the lncRNA subtasks are anonymised as s0 to s5, corresponding to the subtask indices of the PGB; this mapping is crosschecked against four per-species statements in the main text (the non-finite collapse on maize; PlantCAD2-L with IA^3^ on *Manihot esculenta* and *Triticum aestivum*; LoRA on *Sorghum bicolor* ), each of which is consistent with, and only with, the PGB order used here. Species names are printed in green when the species is in the pre-training corpus and in red when it is absent from the released pre-training species list. Species names flagged with ‡ are absent from the corpus but represented by a congener (*Glycine max* : *G. soja*; *Solanum lycopersicum*: four other *Solanum*species; *T. aestivum*: *T. timopheevii* and *Aegilops*); absence of the exact species is therefore not the same as phylogenetic novelty. “Not in the released pre-training species list” means absent from the 320-species stage manifest; 6 of the 326 corpus genomes (4 held out and 2 left windowless by filtering) are not enumerated in that manifest, so absence from the list is not proof of exclusion from training.

**Listing 1:**
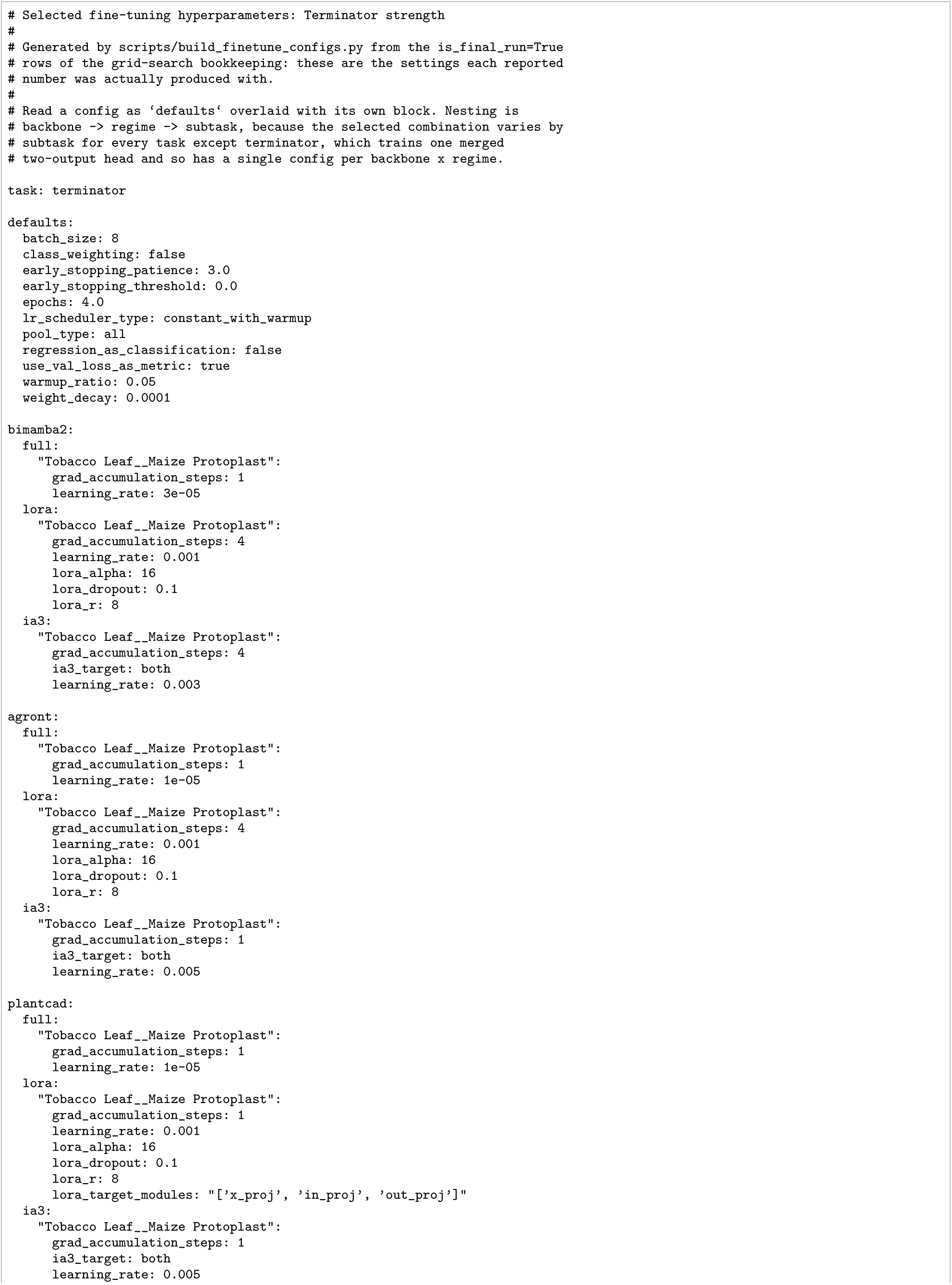
Selected fine-tuning hyperparameters for the terminator strength task.

**Listing 2:**
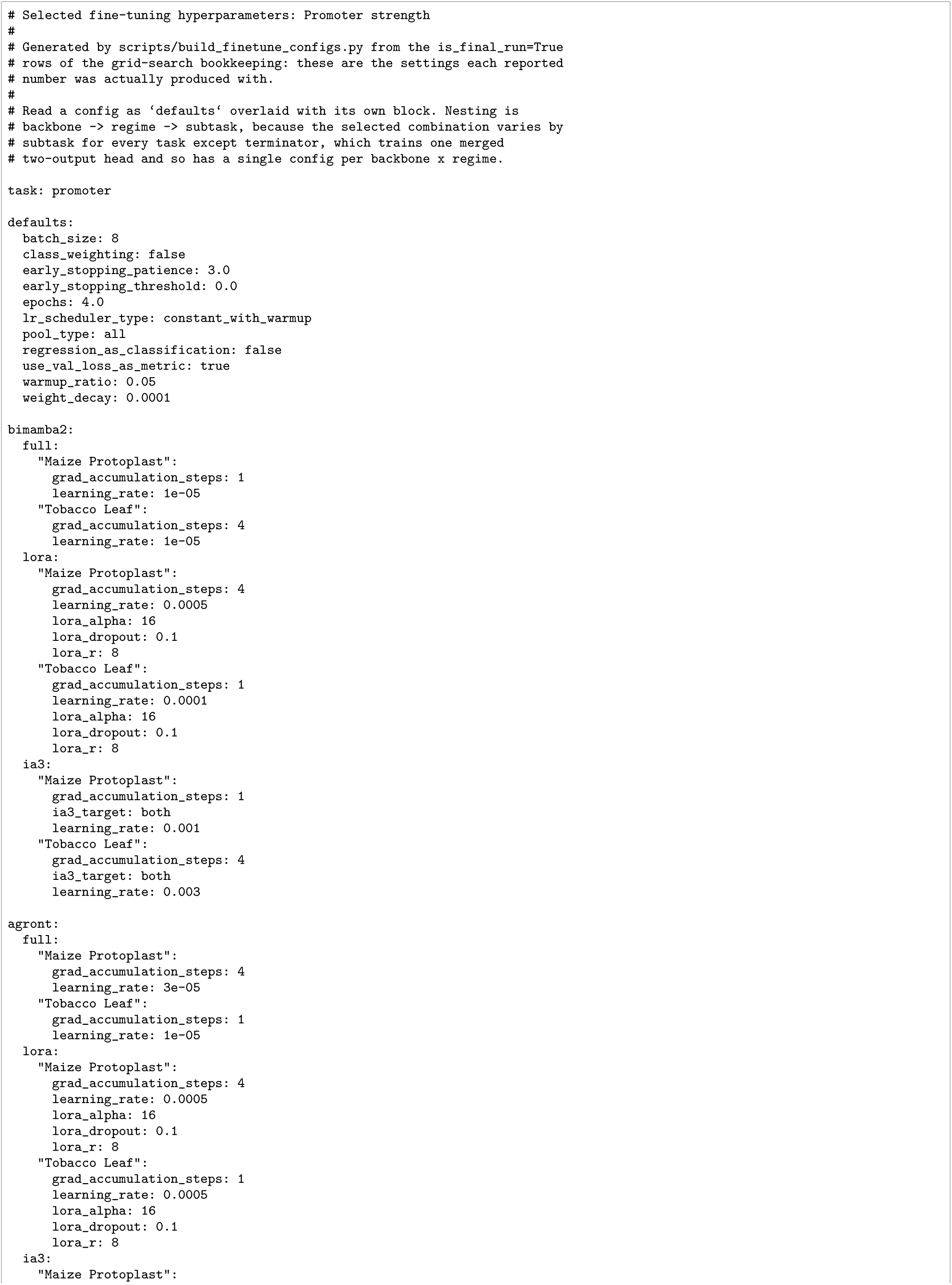
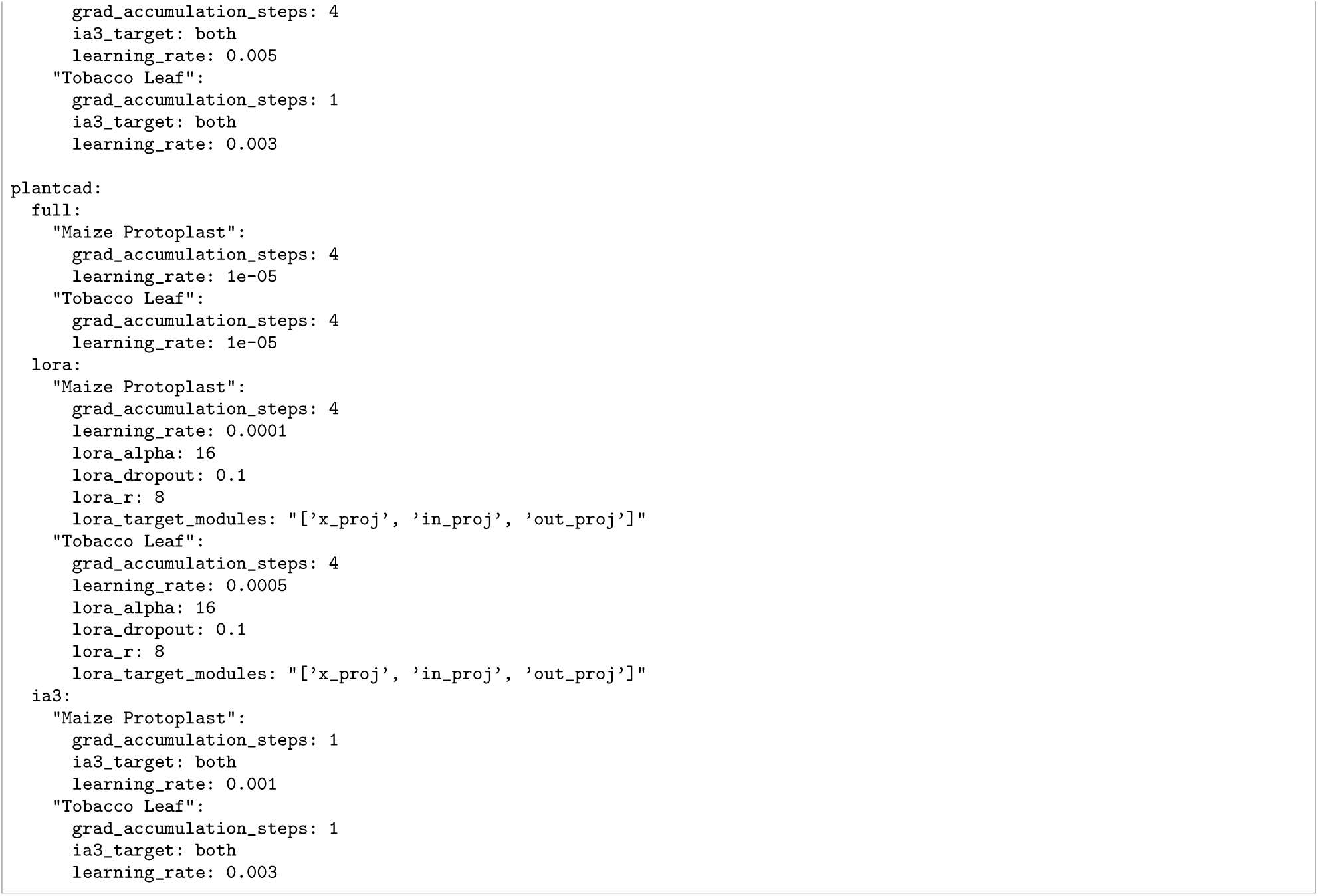
Selected fine-tuning hyperparameters for the promoter strength task.

**Listing 3:**
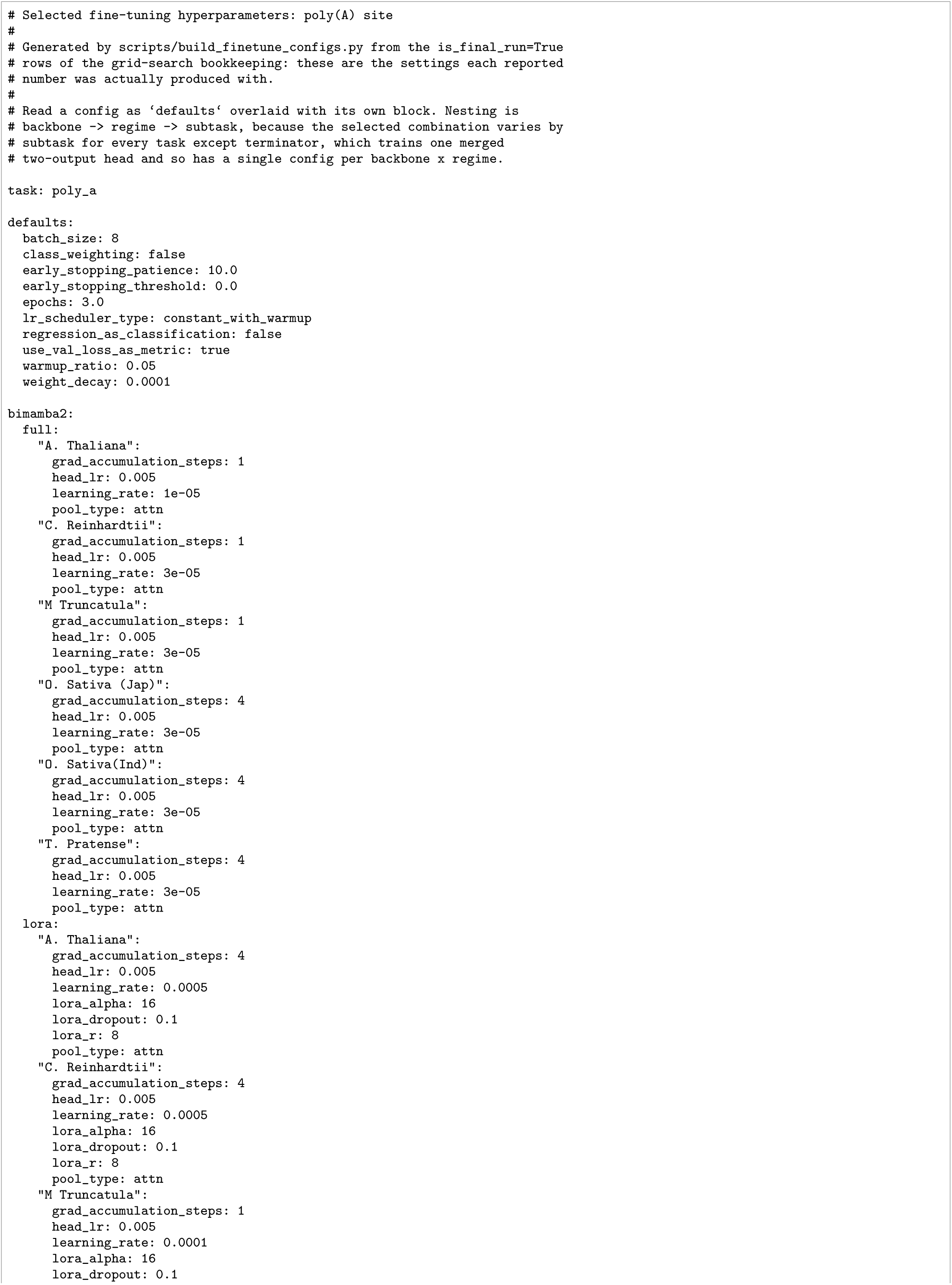
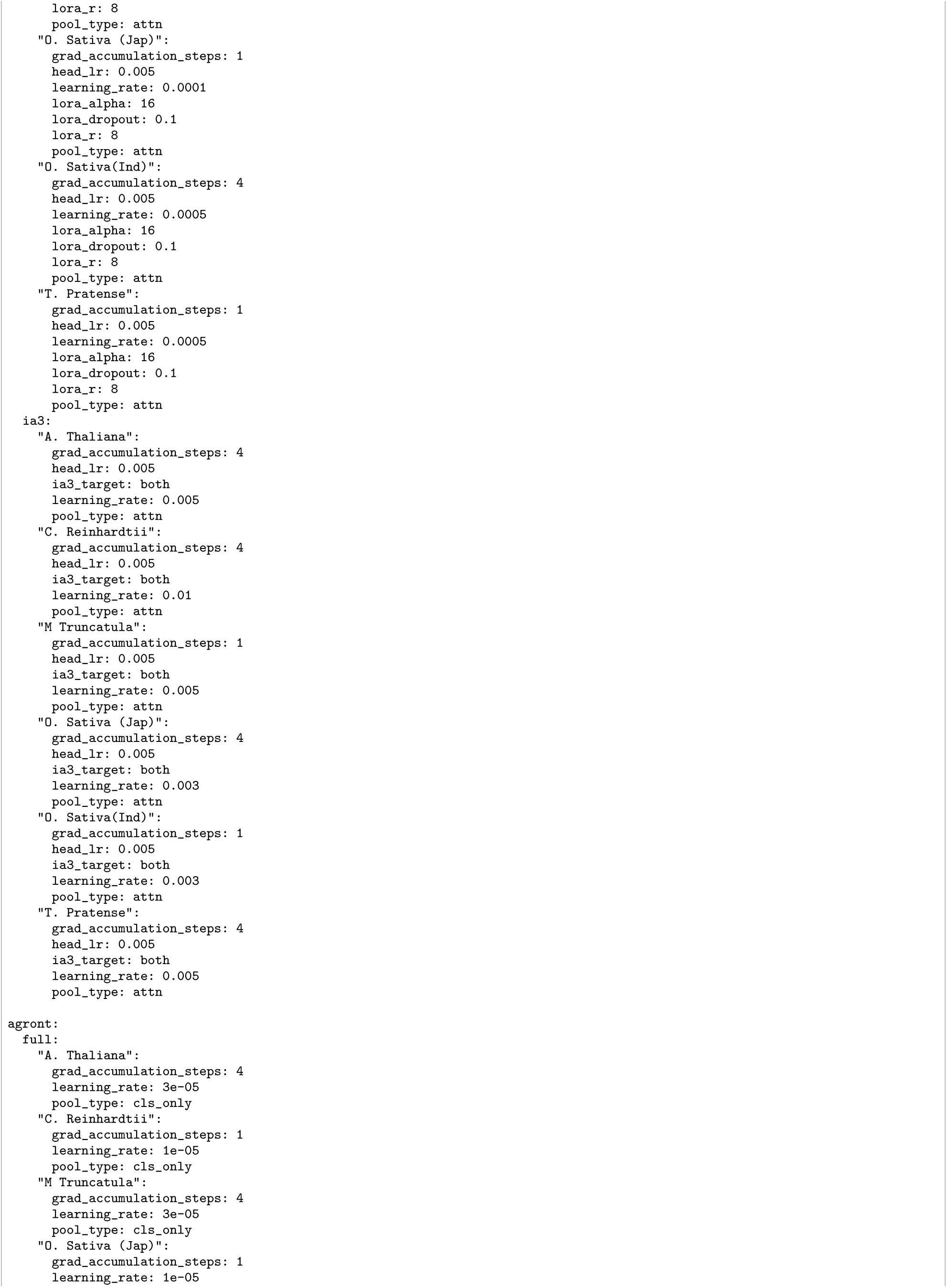
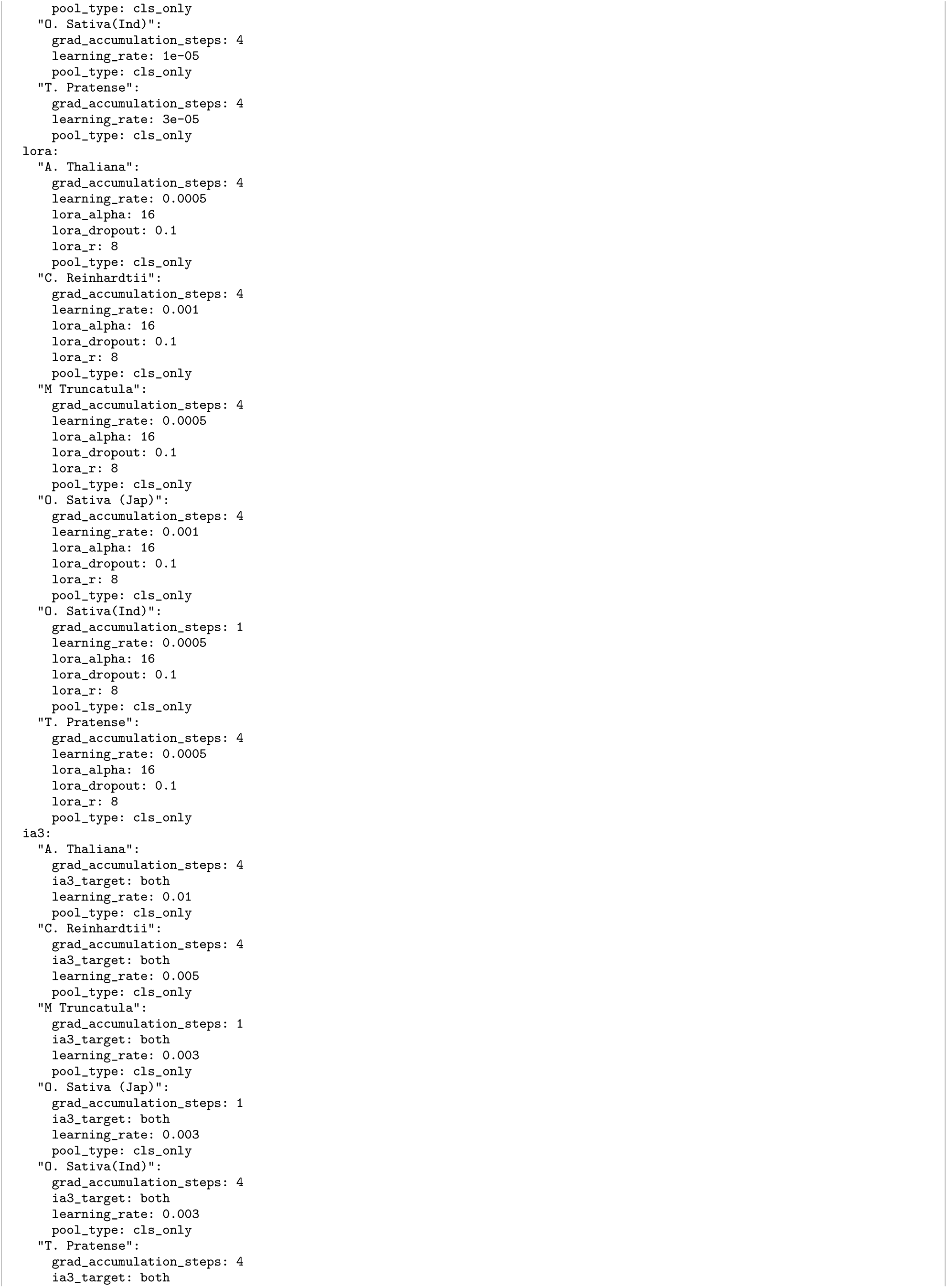
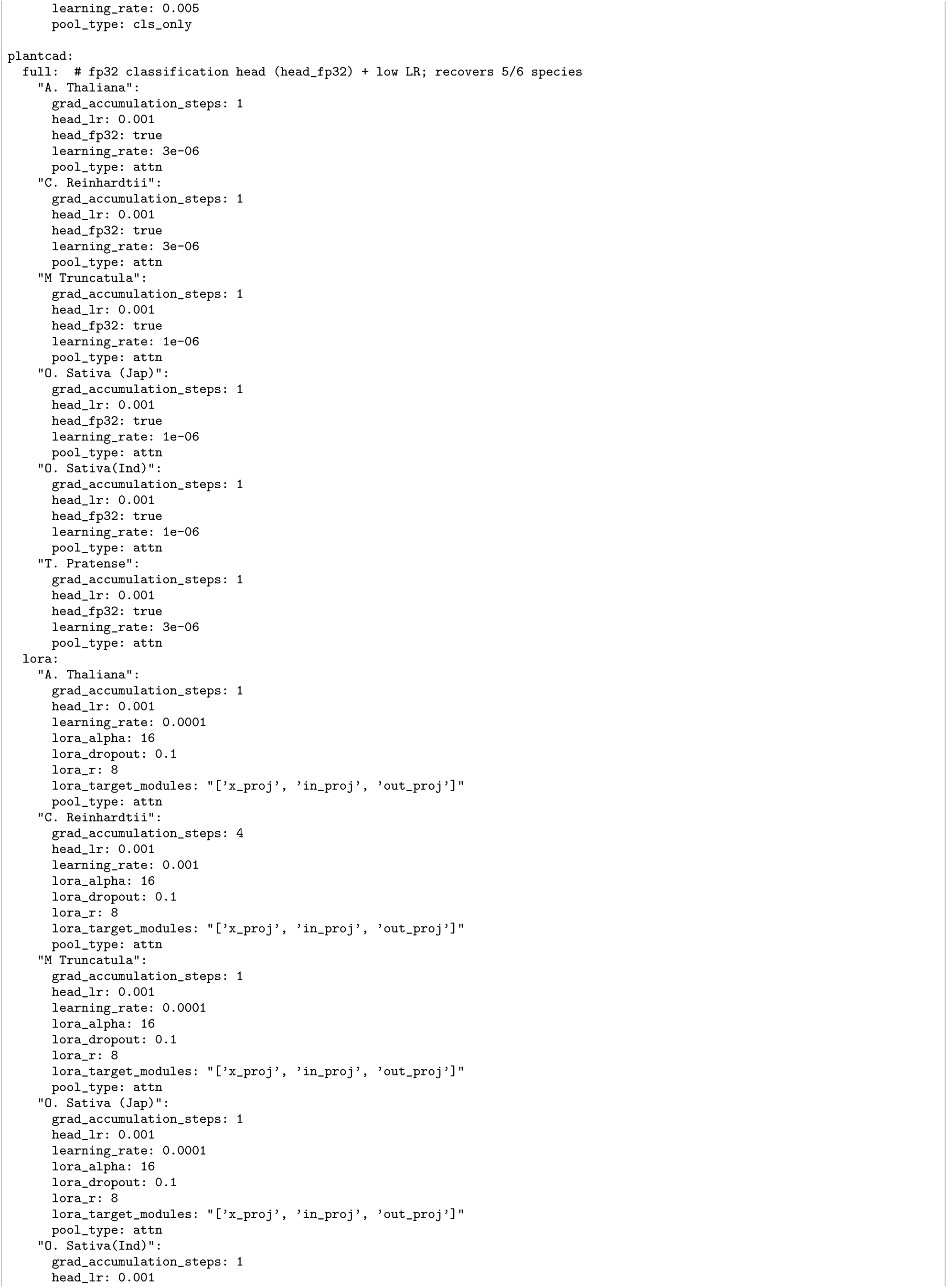
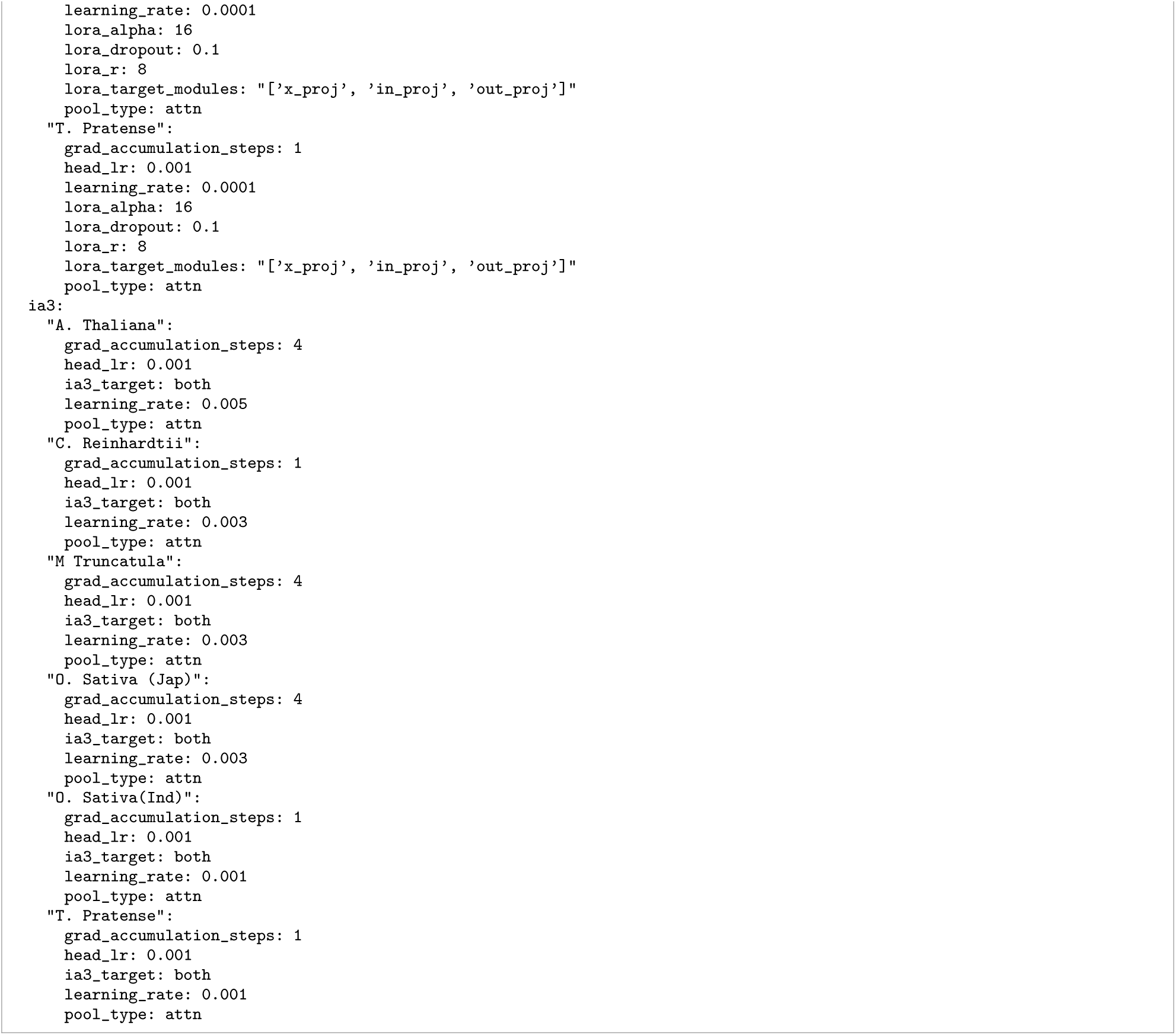
Selected fine-tuning hyperparameters, for the poly(A) site task.

**Listing 4:**
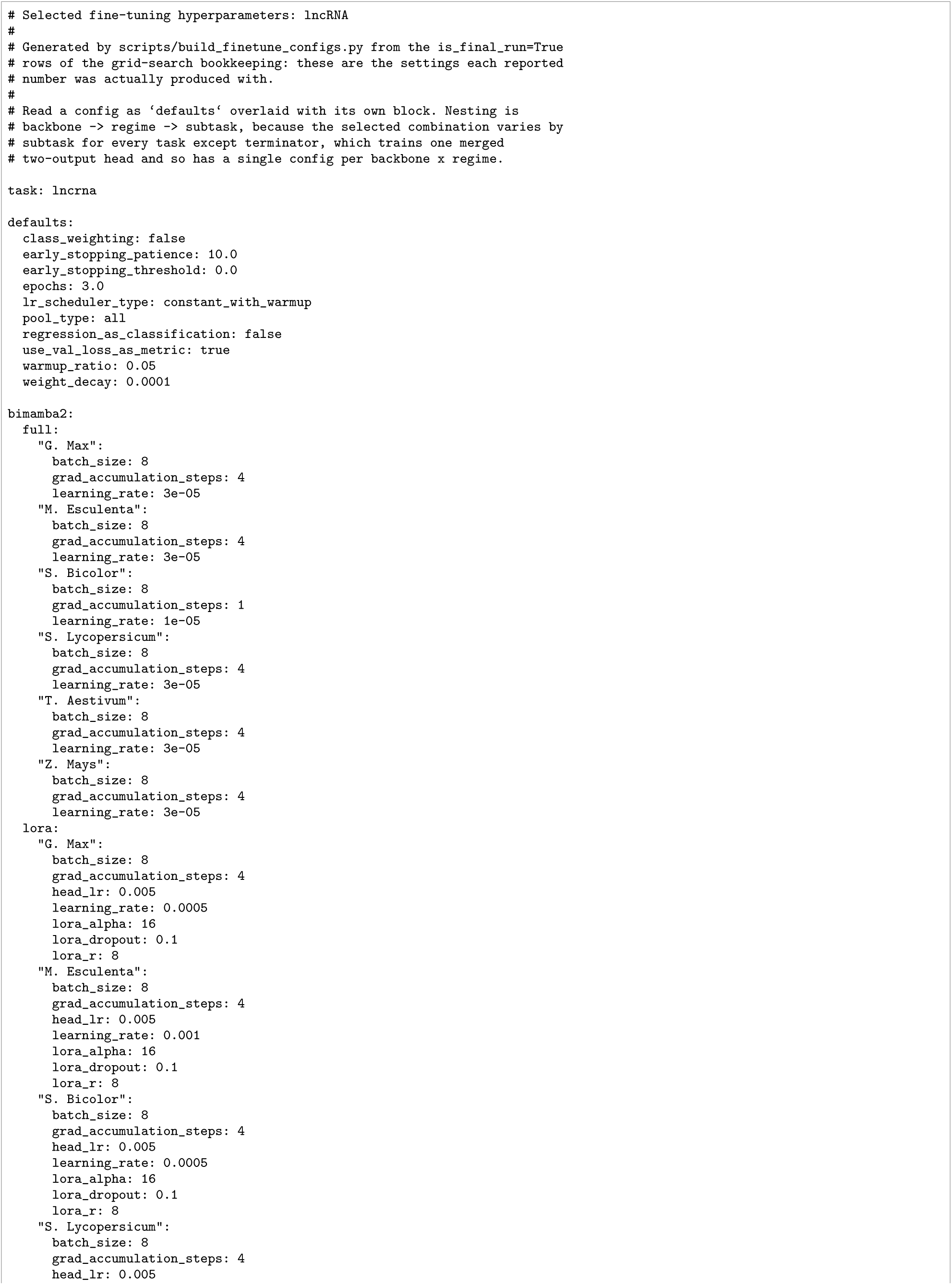
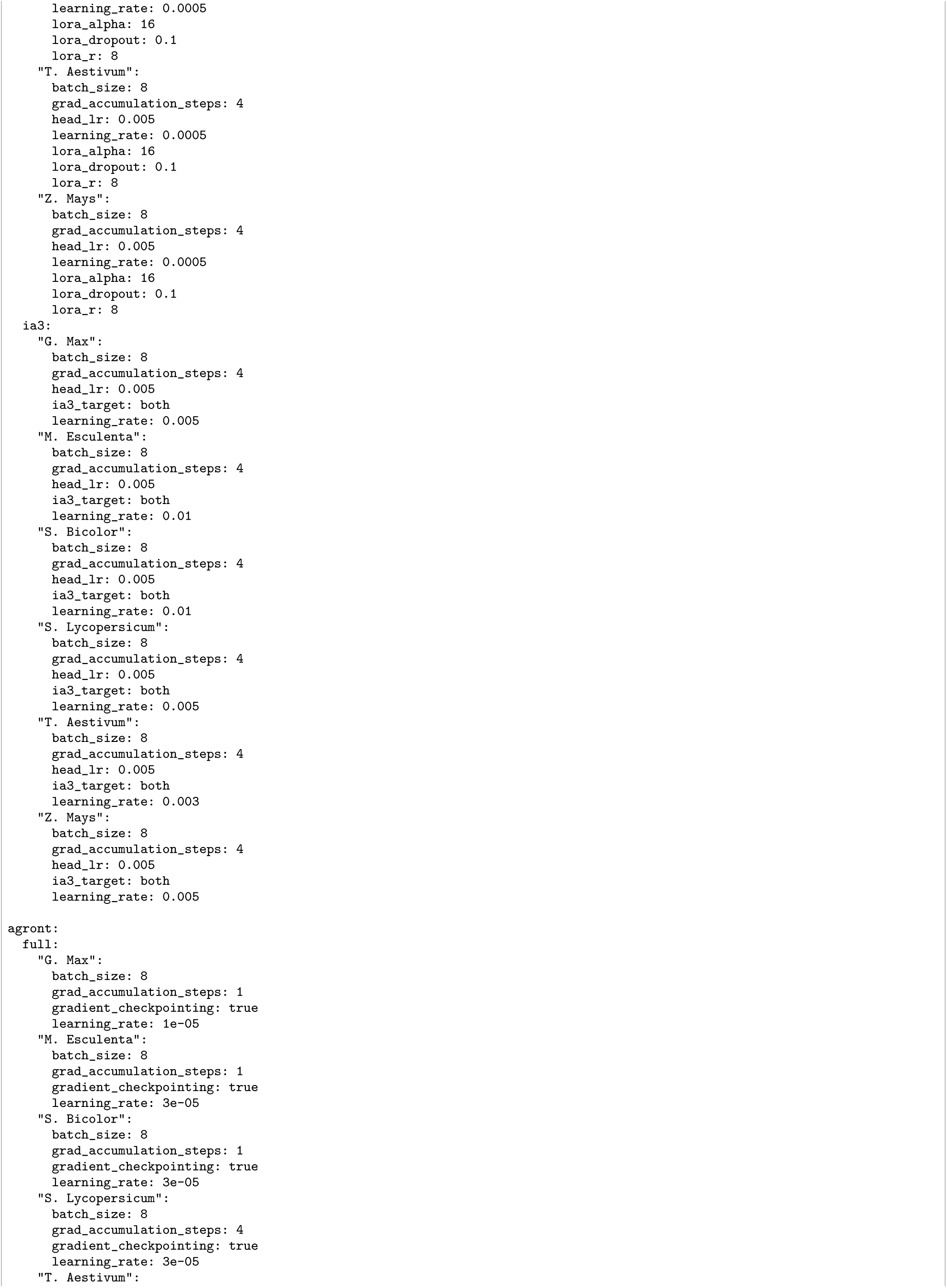
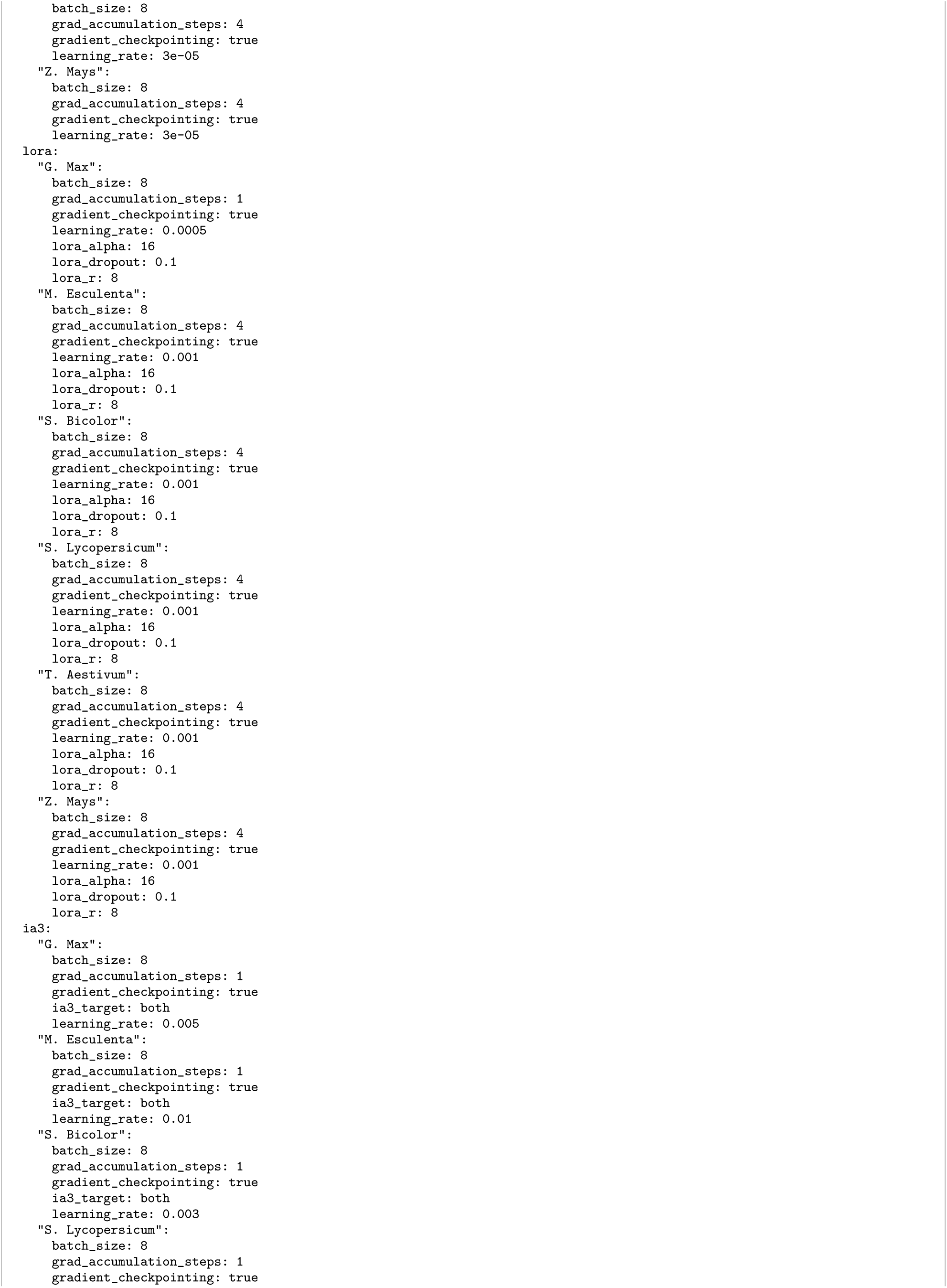
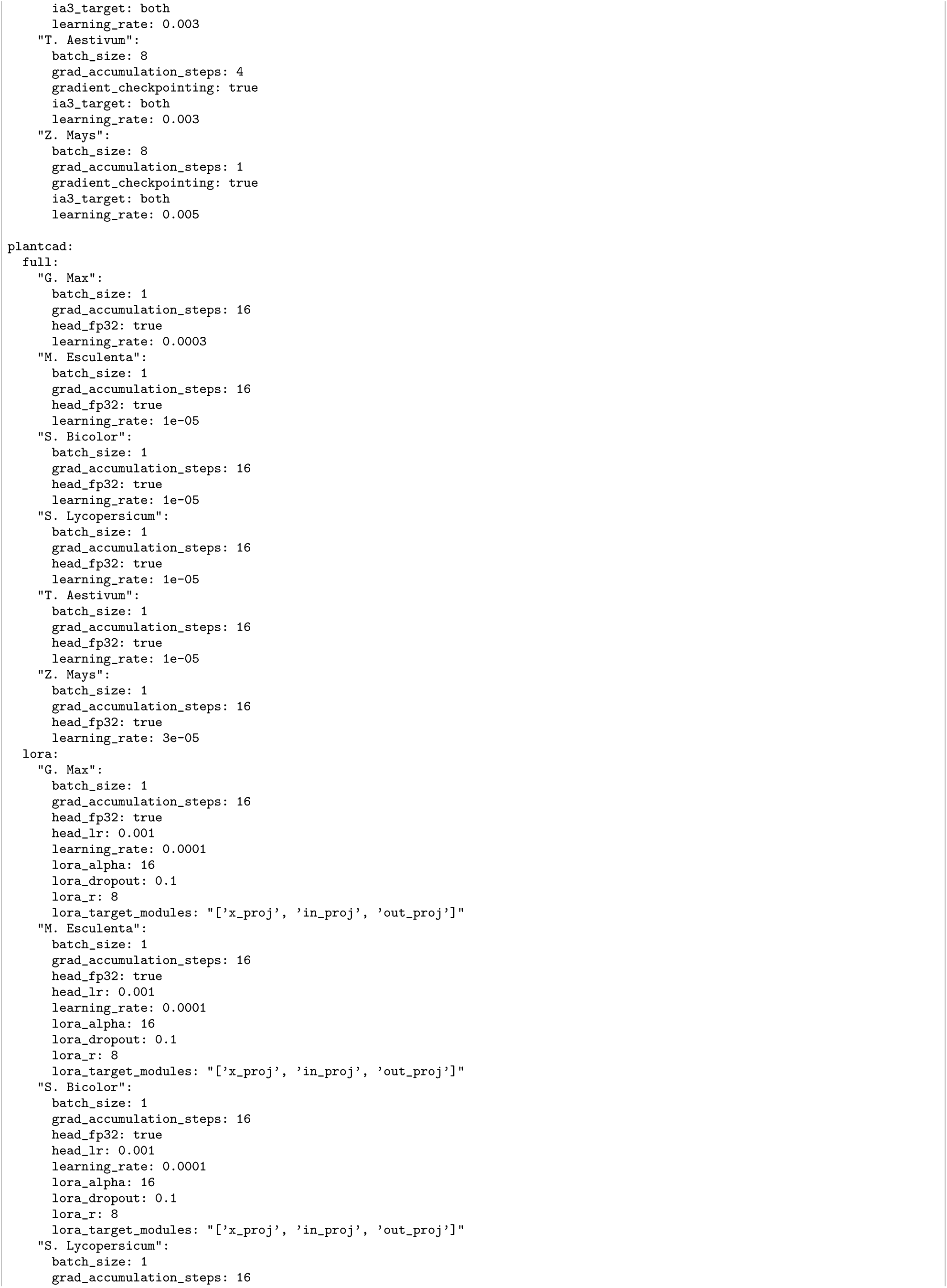
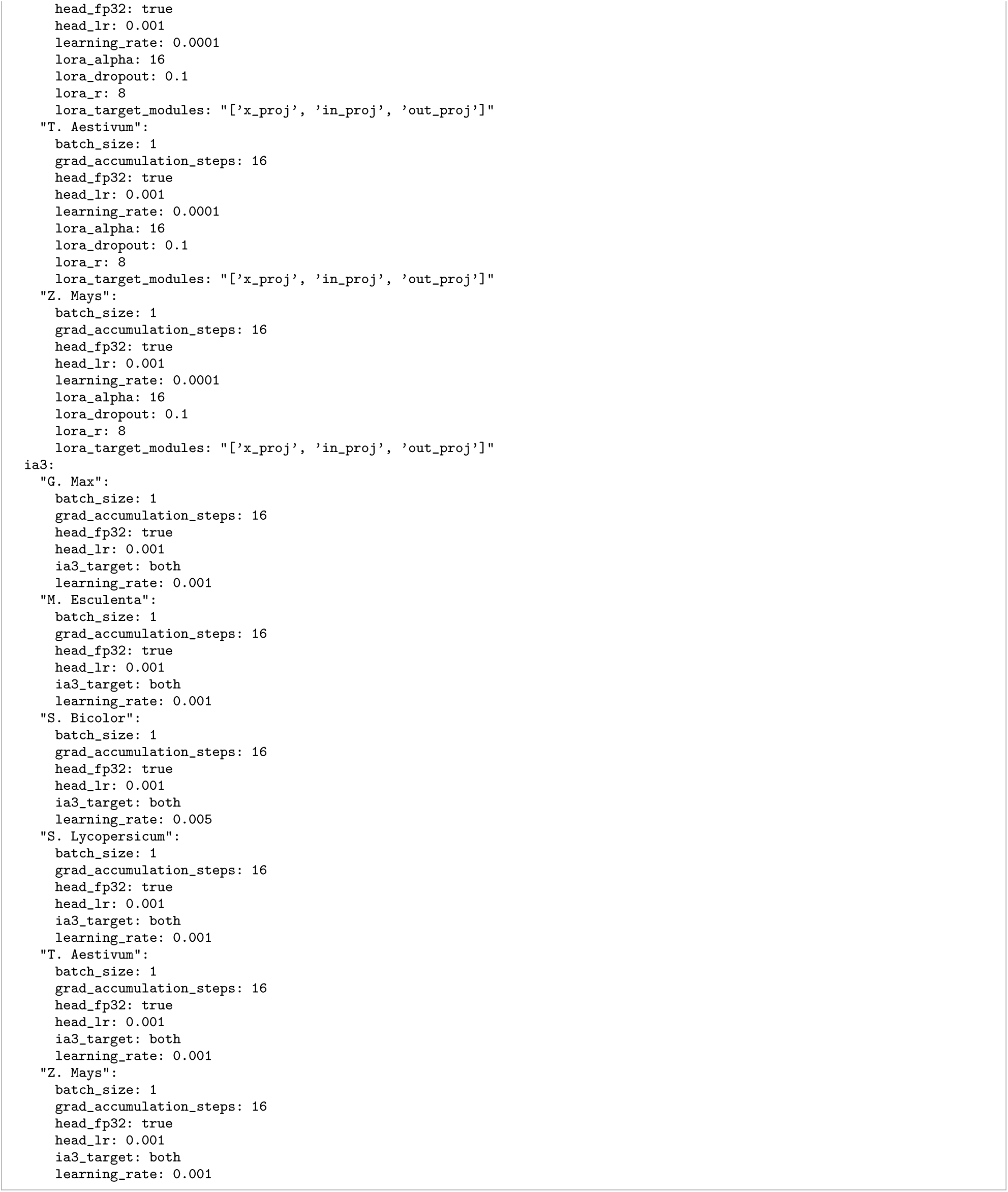
Selected fine-tuning hyperparameters for the lncRNA task.

**Supplementary Figure S27.**
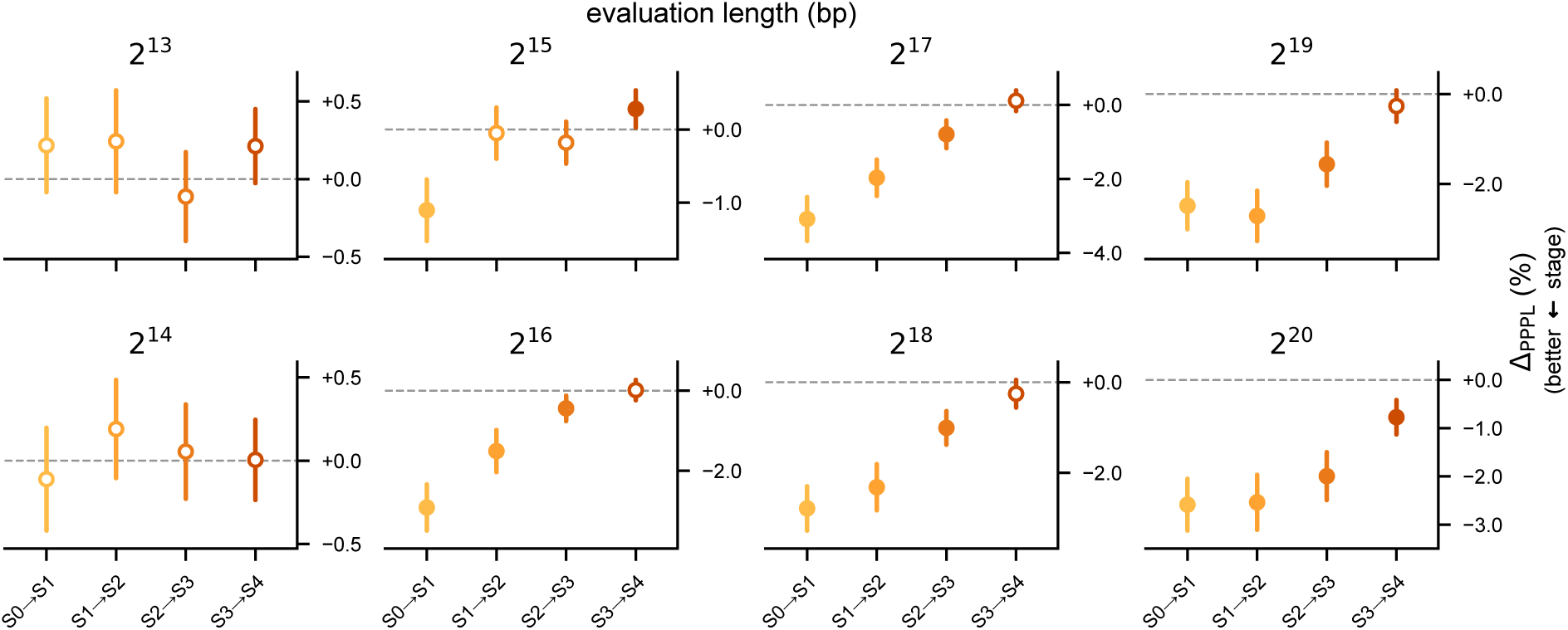
Marginal change in PPPL from successive context-extension stages on heldout sequences at different lengths. Each facet is titled with its evaluation length and displays marginal PPPL changes from each successive context-extension stage (S0 S1 to S3 S4). The scores are percentages of the Botanic1-S0 PPPL baseline at that evaluation length, so the four percentages add up to the cumulative PPPL change. Facets are not comparable: each length scores sequences of a different composition (Section 4.4.1). Bars are simultaneous 95% intervals over the four contrasts, from the same paired bootstrap as Figure 6a. a marker is filled exactly when its interval excludes zero, so the stage it marks changes PPPL significantly.

**Supplementary Figure S28.**
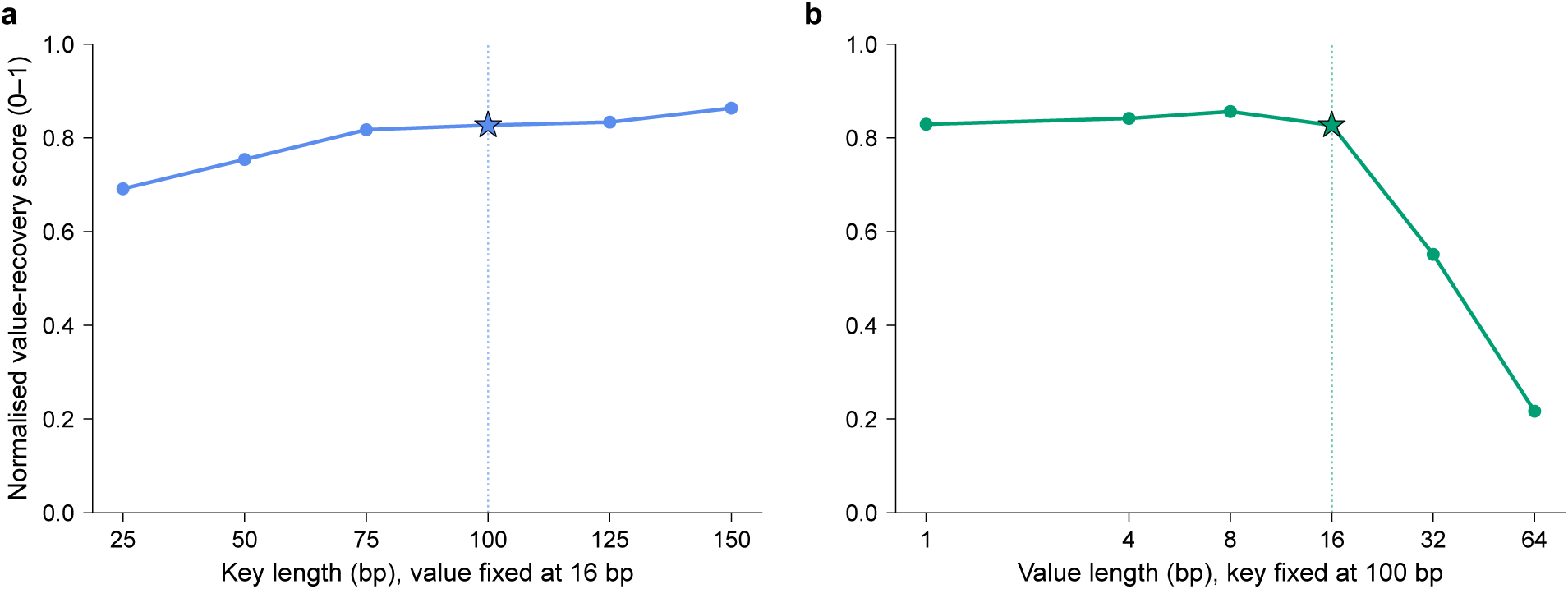
Single-needle key-value retrieval: Key and value lengths sweeps. **a**, Normalised value-recovery score against key length at a 16 bp value. **b**, The same score against value length at a 100 bp key. The settings that are used everywhere else are marked with a star (100 bp key and 16 bp value). The task construction is drawn in Figure 6b.

**Supplementary Figure S29.**
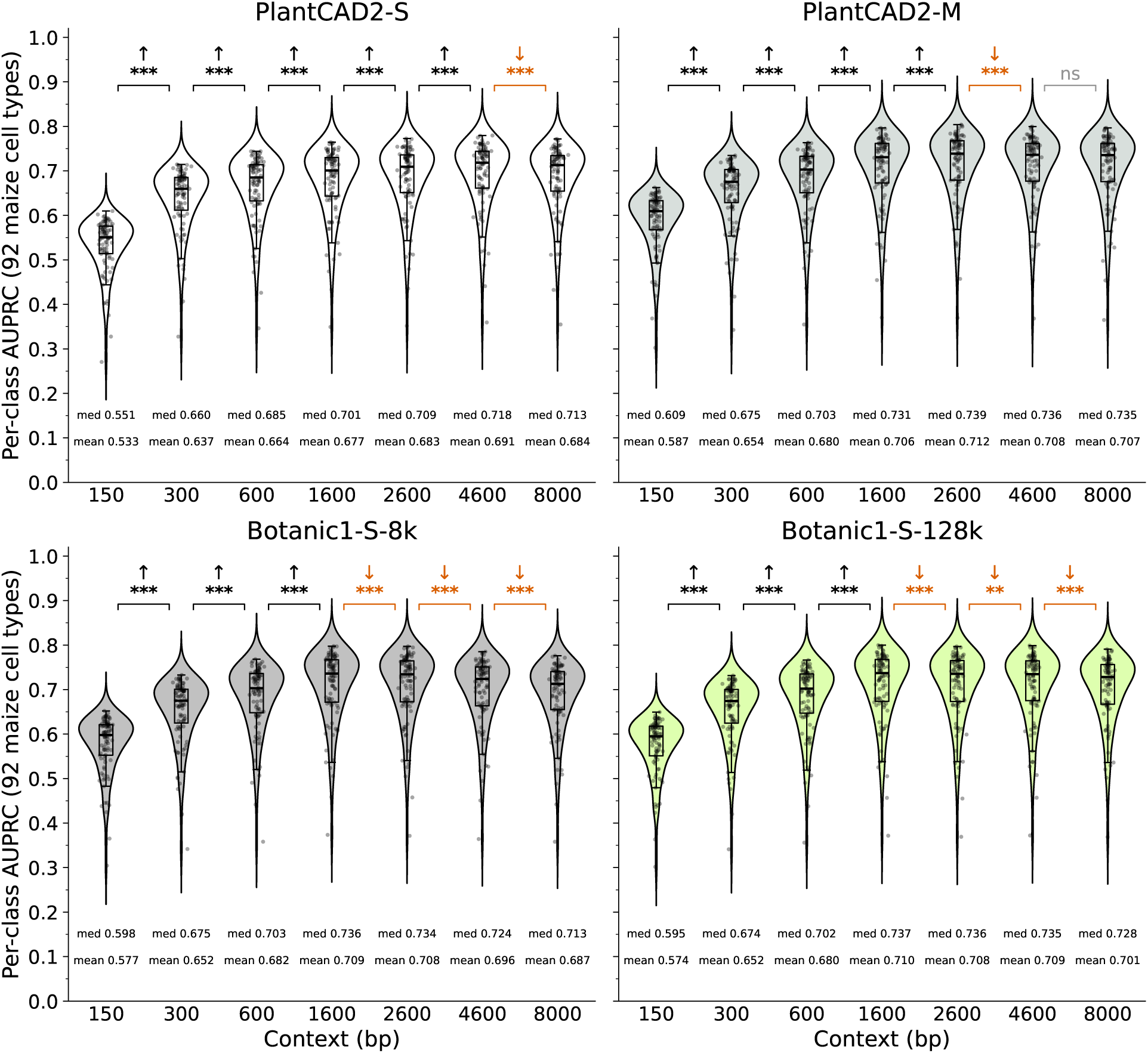
Performance of PlantCAD2-S, PlantCAD2-M, Botanic1-S-8k and Botanic1-S-128k on cell-type-specific accessible chromatin regions prediction, using LoRA fine-tuning on full-window mean-pooled embeddings. For each combination of model and context size, AUPRC is assessed for each of the 92 cell types and the resulting distribution is shown as a violin plot. Brackets between consecutive contexts show the direction of the shift in median AUPRC when it is statistically significant (an upward arrow shows a better performance at the longer context), and the significance marker after Holm-Bonferroni correction within the model (class-paired Wilcoxon signed-rank test; * *p* < 0.05, ** *p* < 0.01, *** *p* < 0.001; ns otherwise); Botanic1-S-8k and Botanic1-S-128k are the two Botanic1-S variants at 8 kbp and 128 kbp native context, respectively.

**Supplementary Table S10:** Statistics on padding used to extend dataset sequences to longer contexts for the maize cell-type-specific chromatin accessibility task. The train/test split corresponds to that reported by Zhai et al. for this dataset [33, 136].

| ctx (bp) | Train |  | Test |  |
| --- | --- | --- | --- | --- |
|  | % padded | mean N (bp) | % padded | mean N (bp) |
| 50 | 0.000 | 0.00 | 0.000 | 0.00 |
| 150 | 0.000 | 0.00 | 0.000 | 0.00 |
| 300 | 0.000 | 0.00 | 0.000 | 0.00 |
| 600 | 0.015 | 0.00 | 0.012 | 0.00 |
| 1,600 | 0.073 | 0.17 | 0.075 | 0.16 |
| 2,600 | 0.124 | 0.58 | 0.118 | 0.55 |
| 4,600 | 0.218 | 2.02 | 0.212 | 2.02 |
| 8,000 | 0.354 | 6.00 | 0.365 | 6.29 |
| 16,000 | 0.602 | 19.9 | 0.641 | 21.7 |
| 32,000 | 0.935 | 58.2 | 1.042 | 67.3 |
| 64,000 | 1.383 | 163.7 | 1.552 | 189.5 |
| 128,000 | 1.897 | 414.6 | 2.112 | 448.9 |

**Supplementary Table S11:** Effective learning rate used in each LoRA fine-tune training run for the cell-typespecific chromatin-accessibility task at long context, strategies 1 and 2 (*Full-window LoRA fine-tuning* and *Central-window LoRA fine-tuning* ).

| context (bp) | PlantCAD2-S | PlantCAD2-M | Botanic1-S-8k | Botanic1-S-128k |
| --- | --- | --- | --- | --- |
| 150 | $2 \times 10^{-4}$ | $4 \times 10^{-4}$ | $1 \times 10^{-4}$ | $4 \times 10^{-4}$ |
| 300 | $4 \times 10^{-4}$ | $4 \times 10^{-4}$ | $1 \times 10^{-4}$ | $4 \times 10^{-4}$ |
| 600 | $2 \times 10^{-4}$ | $4 \times 10^{-4}$ | $2 \times 10^{-4}$ | $2 \times 10^{-4}$ |
| 1,600 | $2 \times 10^{-4}$ | $8 \times 10^{-4}$ | $4 \times 10^{-4}$ | $4 \times 10^{-4}$ |
| 2,600 | $4 \times 10^{-4}$ | $8 \times 10^{-4}$ | $8 \times 10^{-4}$ | $8 \times 10^{-4}$ |
| 4,600 | $8 \times 10^{-4}$ | $1.6 \times 10^{-3}$ | $8 \times 10^{-4}$ | $8 \times 10^{-4}$ |
| 8,000 | $1.6 \times 10^{-3}$ | $3.2 \times 10^{-3}$ | $1.6 \times 10^{-3}$ | $1.6 \times 10^{-3}$ |
| 16,000 | — | — | — | $3.2 \times 10^{-3}$ |
| 32,000 | — | — | — | $6.4 \times 10^{-3}$ |

**Supplementary Table S12:**
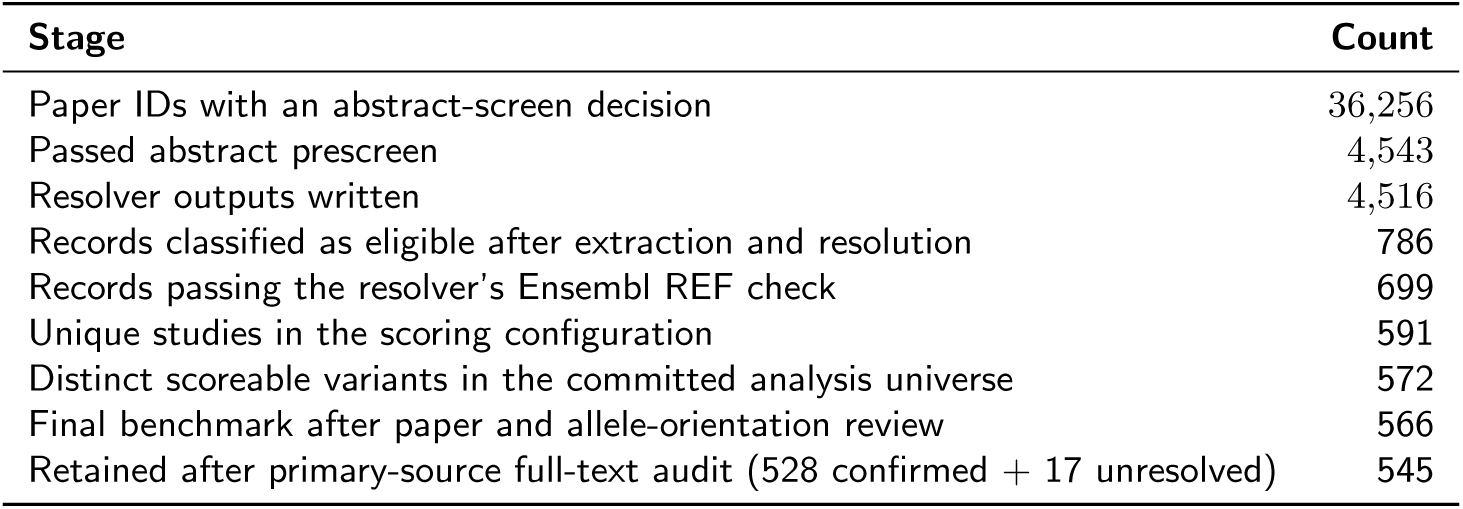
Construction of the causal variant benchmark dataset. The first five rows describe literature search and automated curation. The final three rows describe the construction of the scoring set, including manually configured studies and the removal of duplicate or unscoreable variants.

**Supplementary Figure S30.**
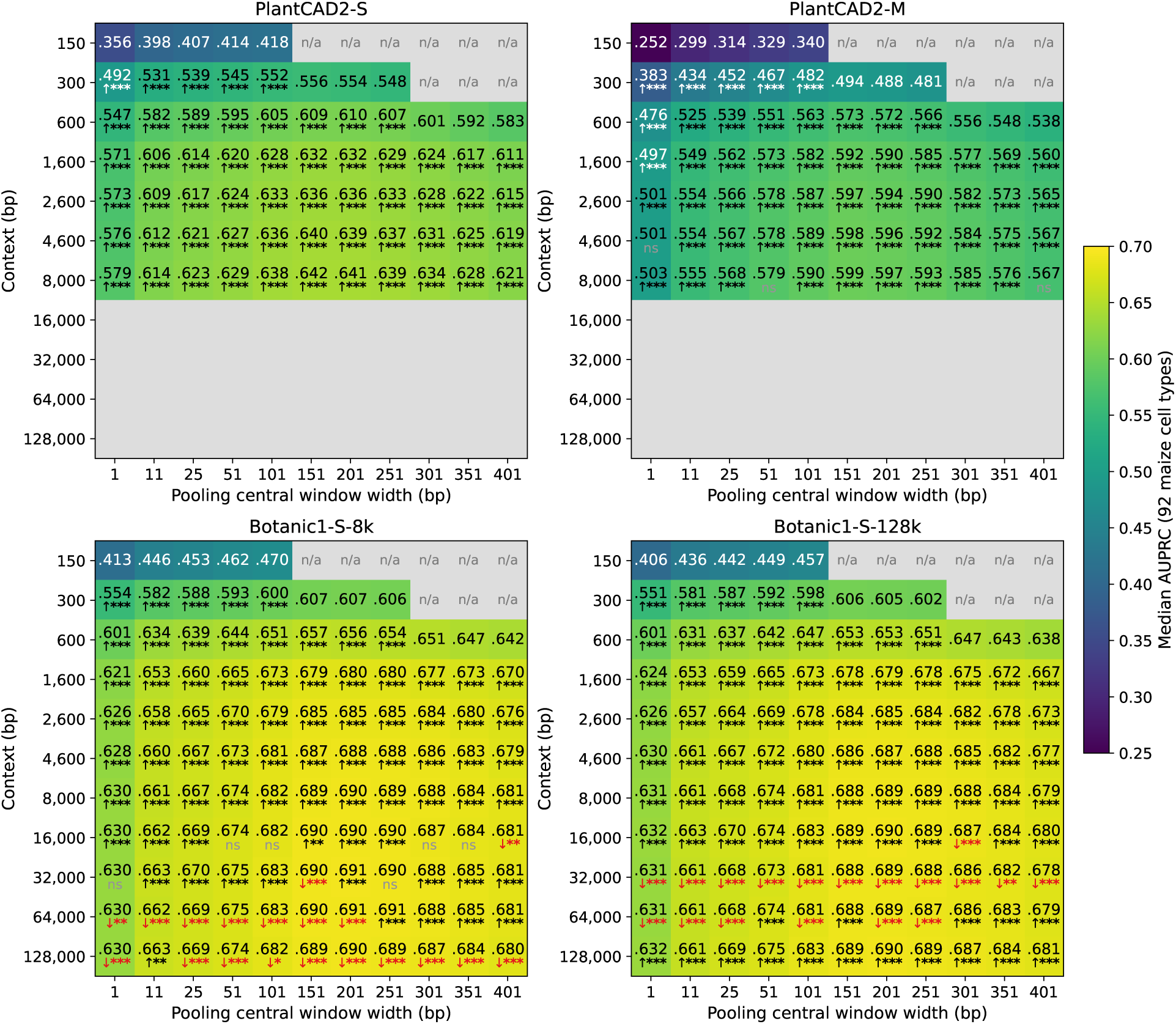
Frozen-embedding probe on the cell-type-specific chromatin accessibility task, with each backbone kept frozen and only a single ESM-like head trained on hidden states mean-pooled over a peak-centred window. Each panel is one backbone; each heatmap sweeps input context length (y-axis) against the pooling window width (x-axis). Cell colour and text report the median per-class AUPRC over the 92 maize cell types. Per-cell arrows and stars report a class-paired Wilcoxon signed-rank test against the same-width cell at the previous context, Holm-corrected within the width column ( increase, decrease in red; * *p* < 0.05, ** *p* < 0.01, *** *p* < 0.001; ns otherwise).

**Supplementary Figure S31.**
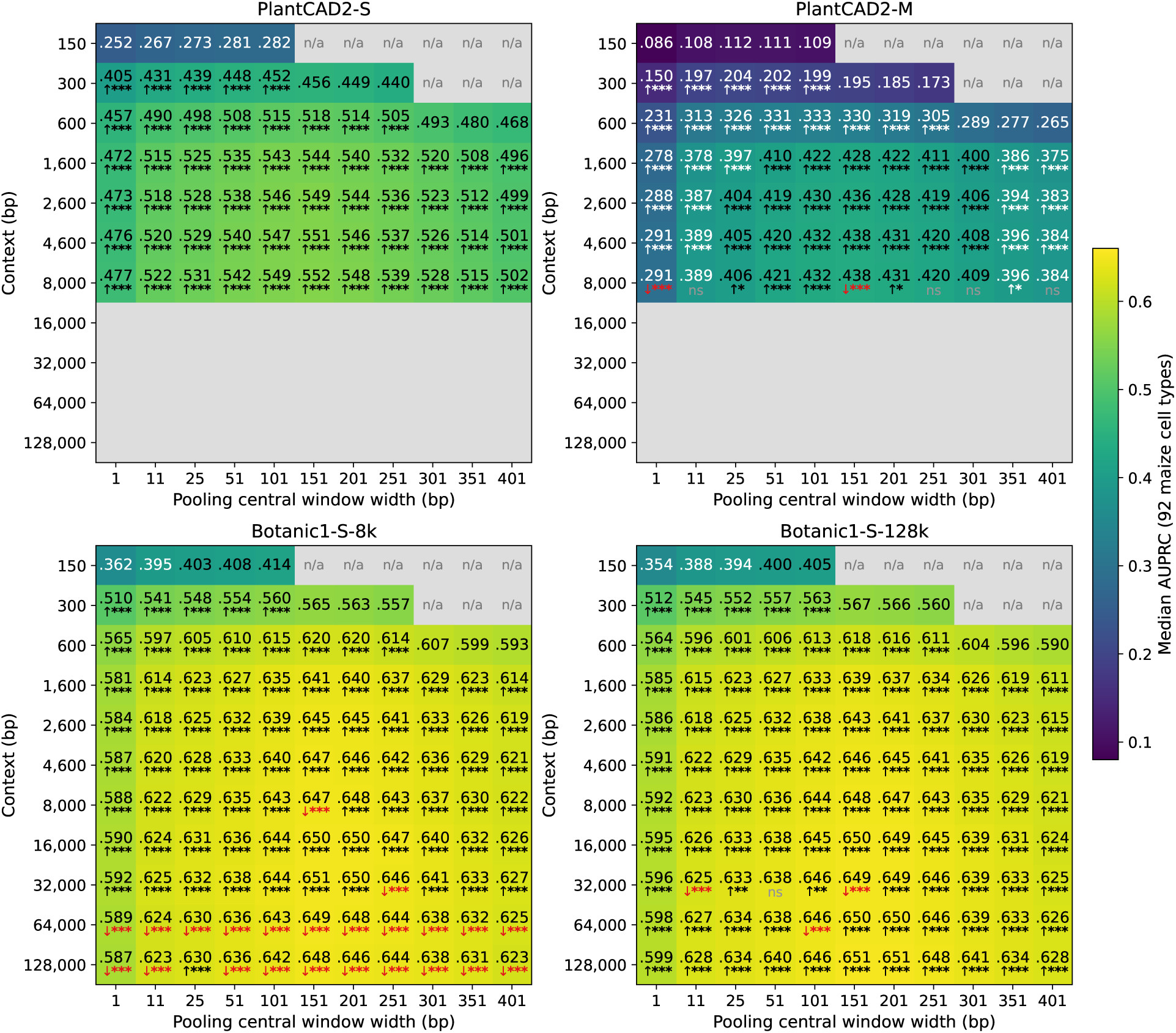
Frozen-embedding probe on the cell-type-specific chromatin accessibility task, with each backbone kept frozen and only a single linear head trained on hidden states mean-pooled over a peak-centred window. Each panel is one backbone; each heatmap sweeps input context (y-axis) against the pooling window width (x-axis). Cell colour and text report the median per-class AUPRC over the 92 maize cell types. Per-cell arrows and stars report a class-paired Wilcoxon signed-rank test against the same-width cell at the previous context, Holm-corrected within the width column ( increase, decrease in red; markers as in Supplementary Figure S30).

**Supplementary Figure S32.**
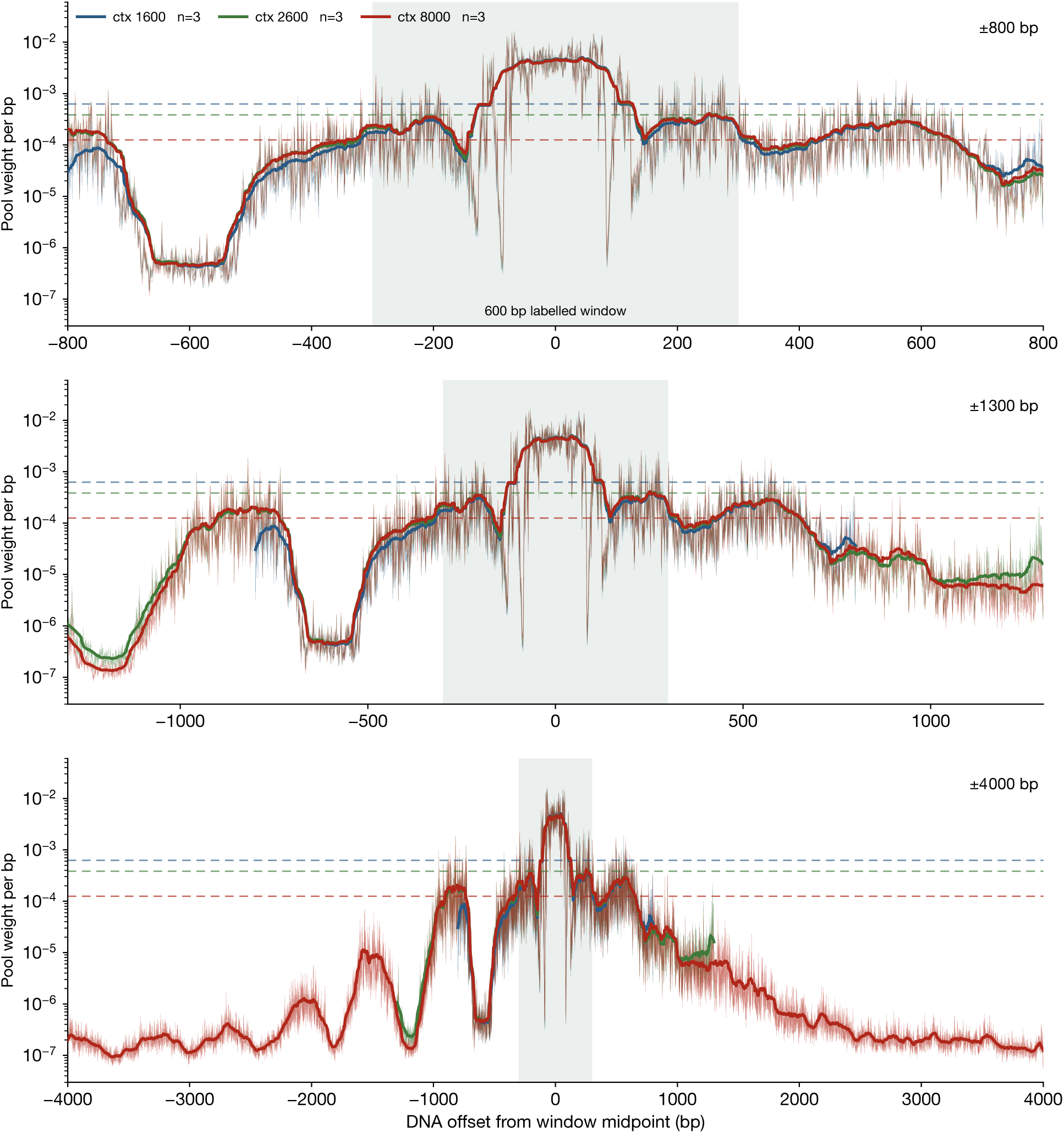
Learned positional pooling weights at 1,600, 2,600 and 8,000 bp of input context on Botanic1-S-8k, three seeds per context, trained at 100 × the classifier head’s learning rate. Thicker lines represent the 51 bp rolling mean of each learned profile, while thinner lines are the raw weights without averaging; the dashed lines are the initialisation value 1/L for each context. The three panels show the same profiles at ±800, ±1,300 and ±4,000 bp; each curve ends where its context does. The recovered window is the same at all three contexts, so 1/L differs between them only because a softmax over more positions puts less weight on each.

**Supplementary Figure S33.**
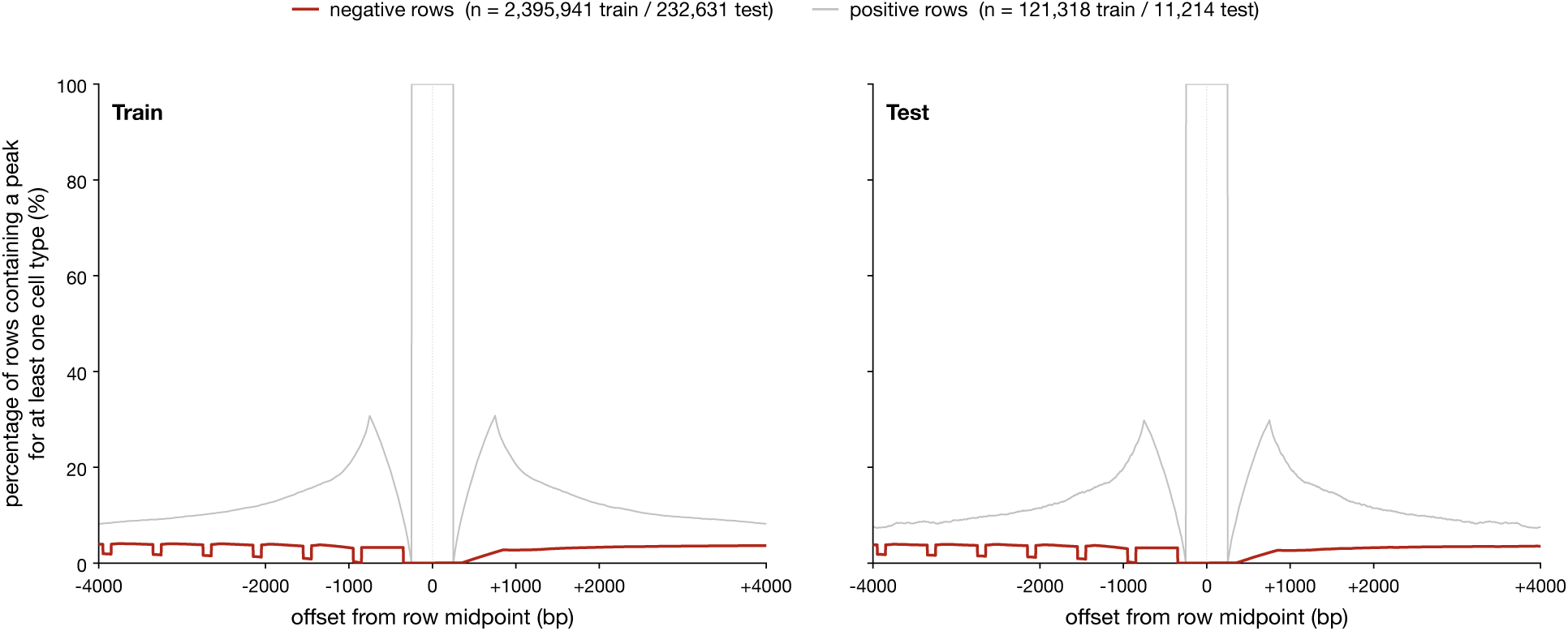
Peak-fraction profile around row midpoints in the maize cell-type-specific chromatin accessibility dataset released by Zhai et al. [33, 136]. For each row and each offset *d* from that midpoint, the curves report the percentage of rows whose 1 bp position at offset *d* overlaps an ACR for at least one of the 92 cell types. Negative rows (dark red) are rows the dataset labels non-accessible in every cell type; positive rows (grey) are rows accessible in at least one cell type.

**Supplementary Table S13:** Causal variant benchmark dataset construction: Prompt roles and required structured outputs.

| Prompt role | Decision | Required output |
| --- | --- | --- |
| Prescreen | Does the abstract plausibly contain a causal plant SNP with functional evidence? | Pass/fail, reason code, and confidence. |
| Extraction | Does the selected paper text meet the benchmark rubric? | Species, variant description and class, causal evidence, alleles/assembly where stated, association-data fields, and confidence. |
| Resolution | Can the structured variant extraction be placed on a usable reference assembly? | Exact coordinate and alleles, reference and VCF resources, liftover steps, evidence metadata, reference-base check, and rejection reason if unresolved. |

**Supplementary Table S14:**
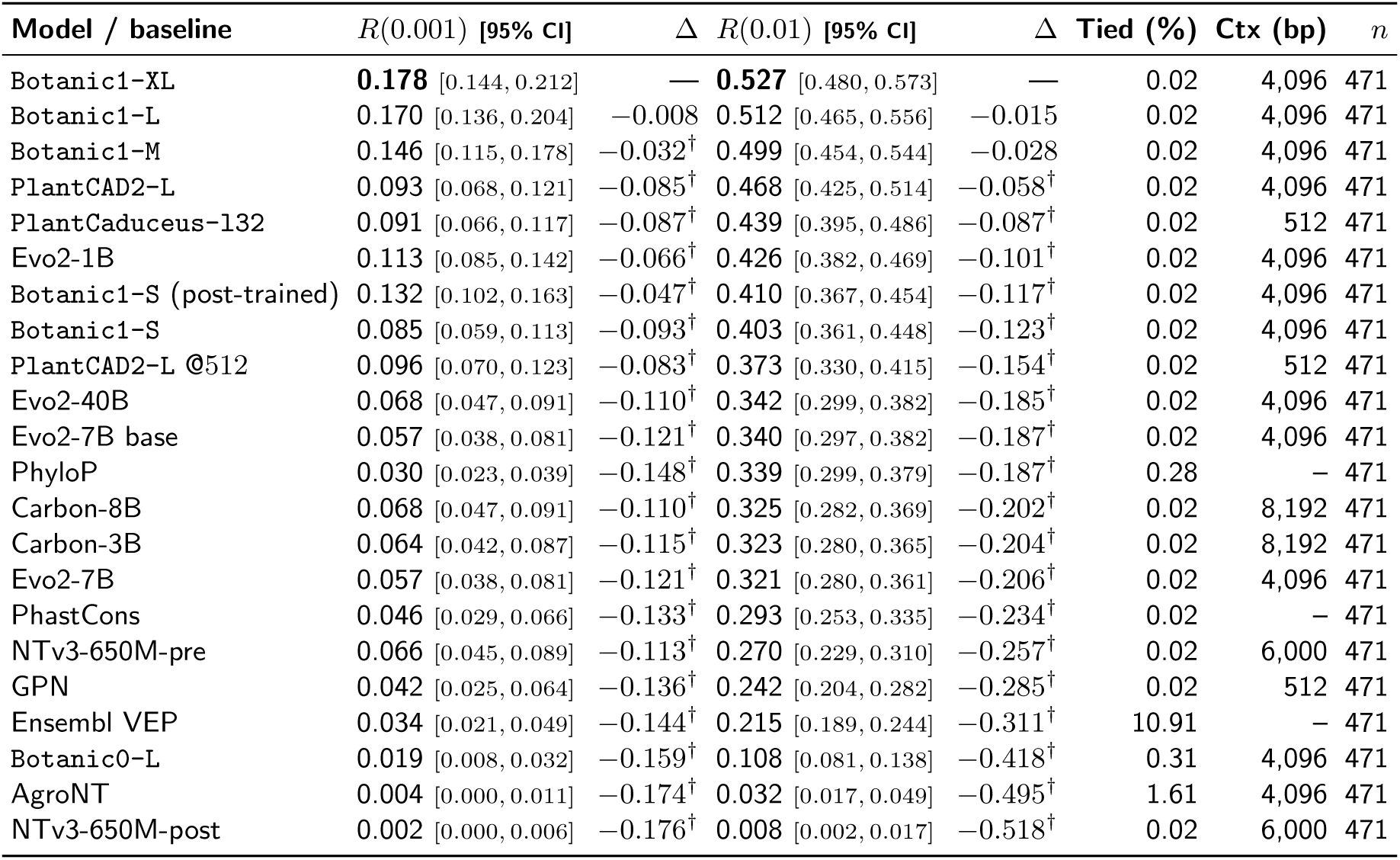
Zero-shot causal-variant prioritisation on the strict intersection set of studies. Same methodology as Table 3, computed on the set of studies that are scored by all methods. The limiting factor are the MSA-derived conservation scores PhastCons and PhyloP which are only supported on 471 studies.

**Supplementary Figure S34.**
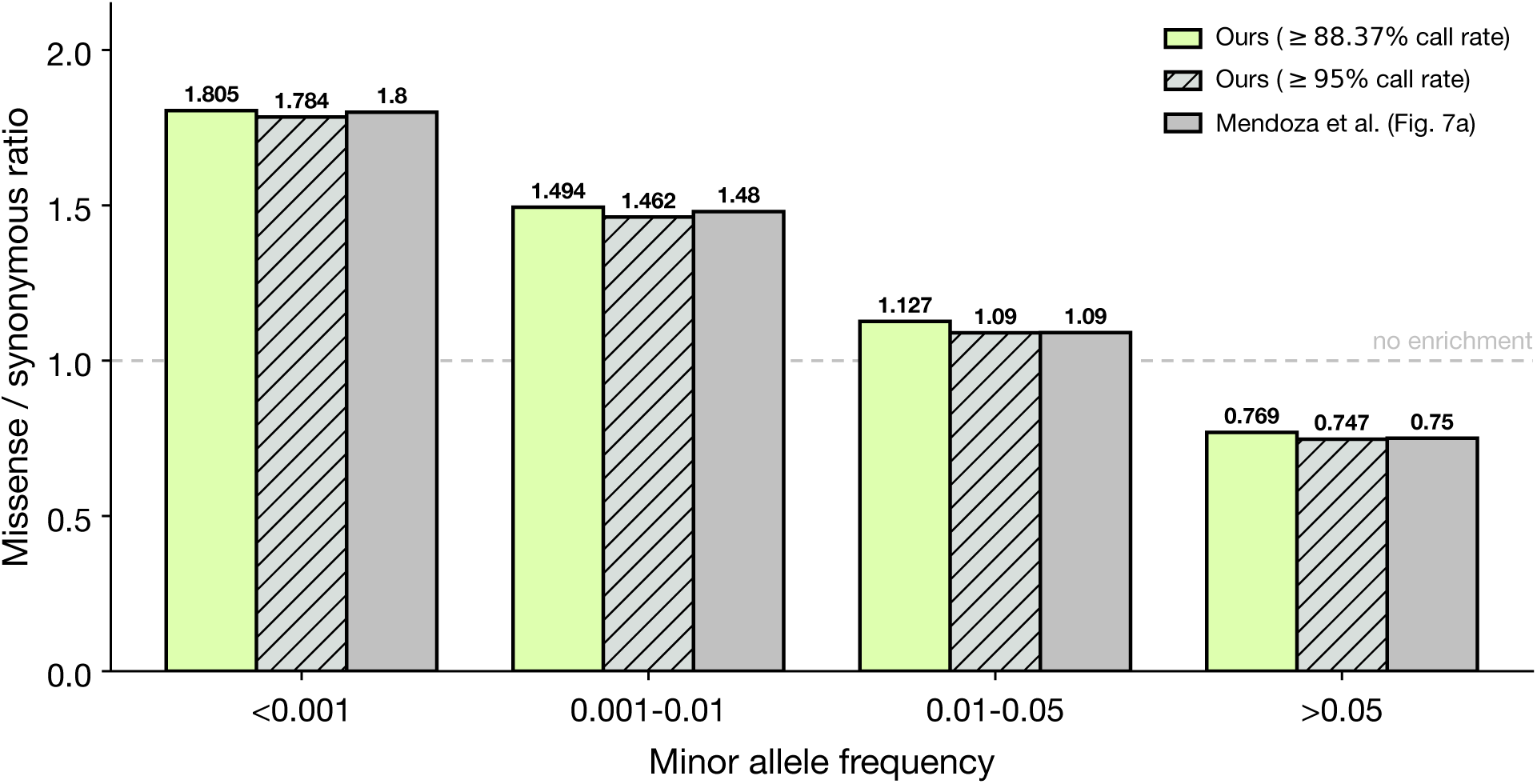
Missense/synonymous ratio of all coding Arabidopsis thaliana SNPs by minor allele frequency. We show the values reported in Figure 7a of Mendoza-Revilla et al. [31], and the values we obtain at 95% and 88.37% call rates on the genomes from the 1001 Genomes Project. 88.37% call rate is the threshold we use for the reported results.

**Supplementary Figure S35.**
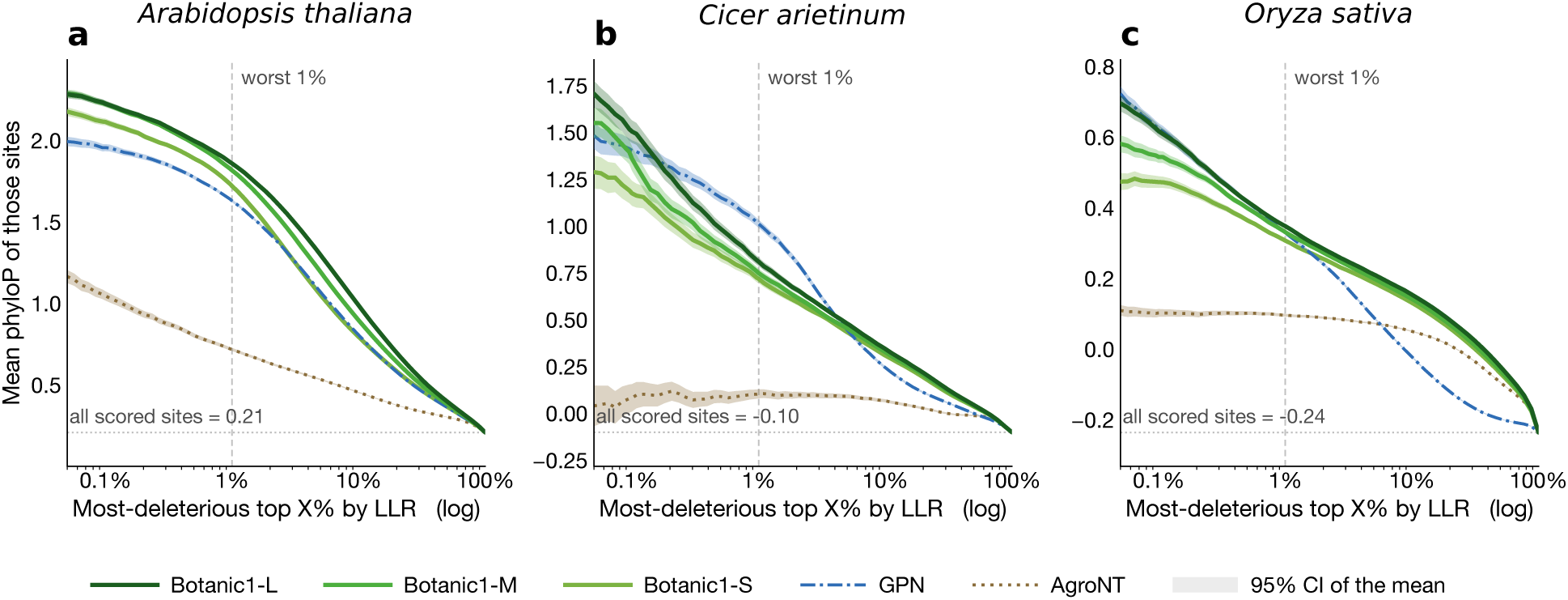
Agreement between LLR and conservation. We rank the variants by their predicted deleteriousness according to each model’s LLR score, and plot the mean phyloP score (indicating their conservation) (y-axis) at variable top percentage (x-axis). The x-axis is in log scale to focus on the lower percentages, and each panel represents the results on one species: **a**, *Arabidopsis thaliana*; **b**, *Cicer arietinum*; **c**, *Oryza sativa*. Figure 9 plots the same values, but by bin (not cumulatively).

**Supplementary Figure S36.**
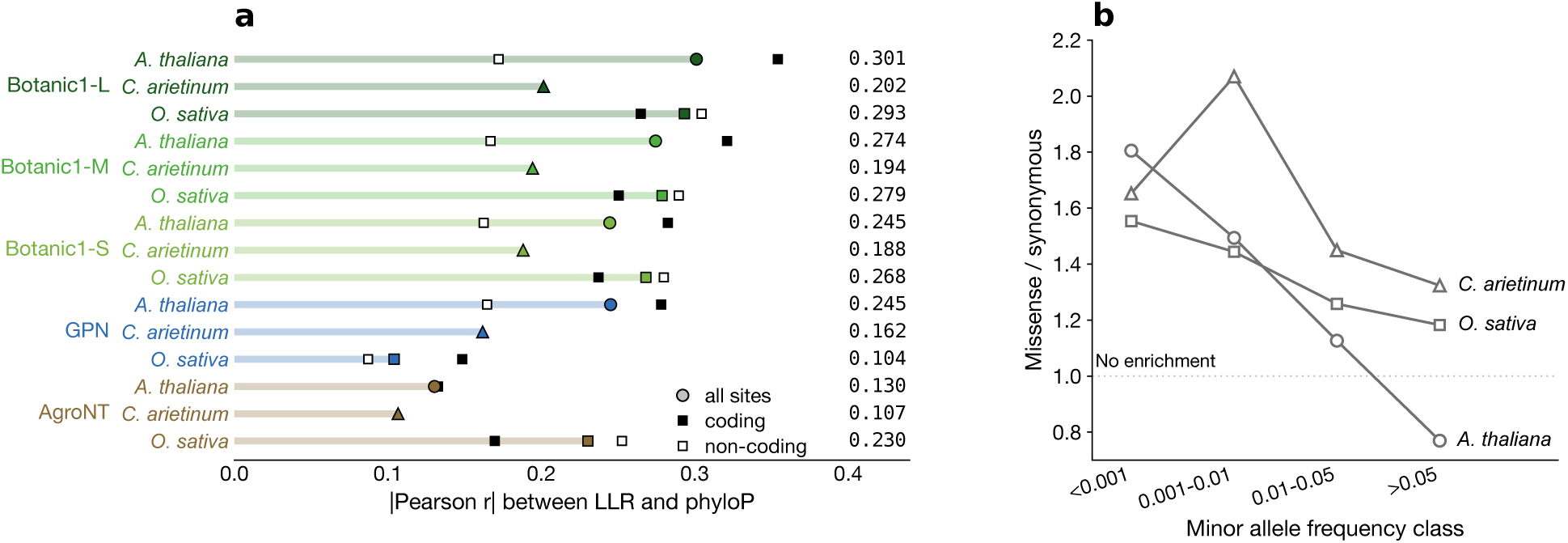
Agreement with conservation by variant class, and the model-independent selection signal. **a**, Pearson correlation coefficient between LLR and phyloP, broken down by variant classification, for each model and species. The *Cicer arietinum* aggregate is computed over all sites and never split by class, so those rows carry the all-sites marker only. **b**, Missense/synonymous ratio across different MAF bins. No model enters this panel: it measures purifying selection in the variant panels themselves. The ratio is not monotonic in *C. arietinum*, rising from 1.65 in the rarest class to 2.07 at 0.1–1% before falling.

**Supplementary Figure S37.**
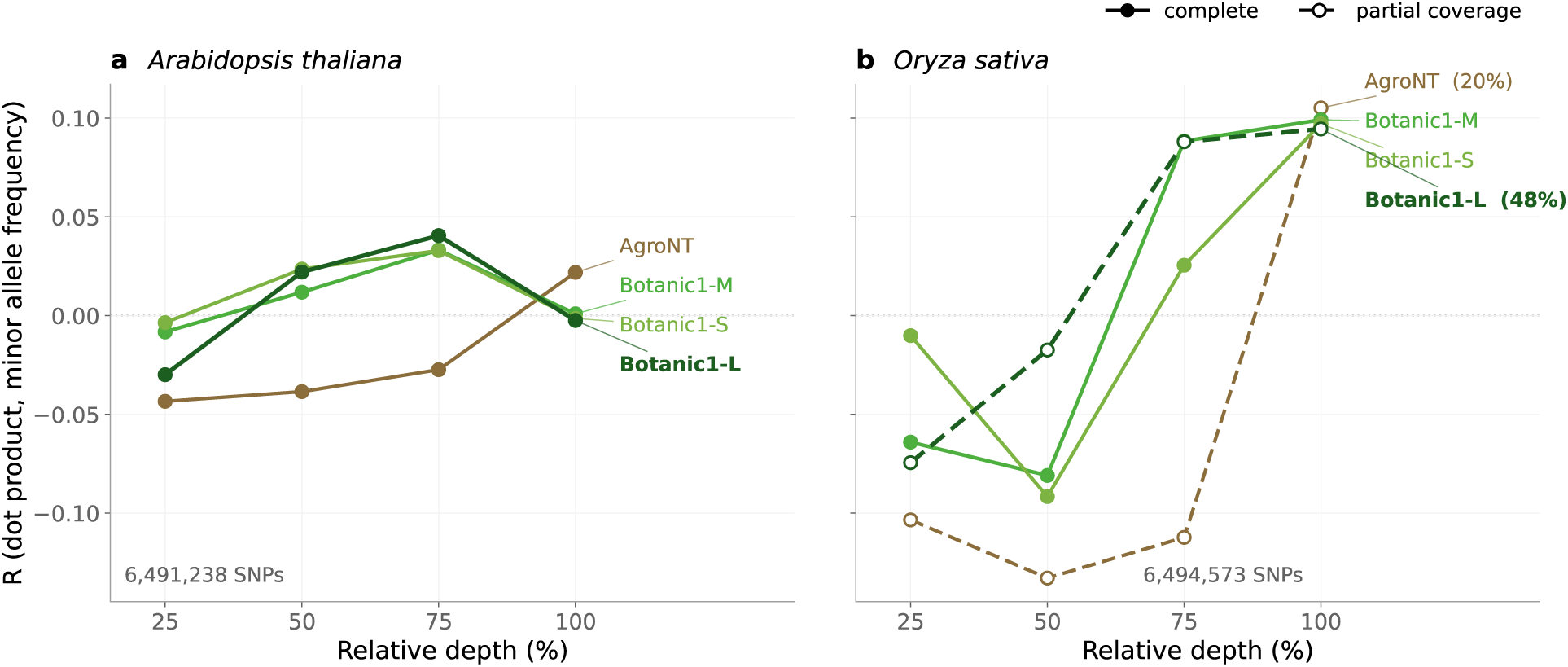
Depth sweep of zero-shot embedding-distance scores on two species. Correlation between the ref-versus-alt embedding dot product and MAF at four layers per model. Layers are matched to be the same relative depth for each model. **a**, *Arabidopsis thaliana* on all 6,491,238 SNPs. **b**, *Oryza sativa* on all 6,494,573 SNPs. Where a model cannot score the full variant set, the fraction of variants it does cover is shown as a percentage next to its label. The peak moves between the two species, from ≈77% relative depth in *Arabidopsis thaliana* to the output layer in *Oryza sativa*.

**Supplementary Table S15:**
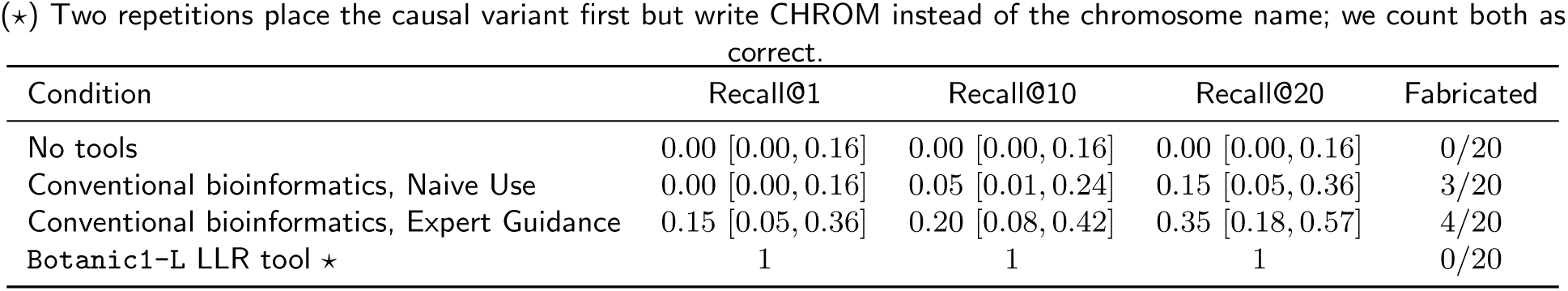
Recovery of the causal melon variant per condition. Recall@k is defined as the fraction of repetitions in which the agent ranks the experimentally validated <u>CmEIN3</u> variant among the top *k* candidates. 95% Wilson intervals are given between brackets. The <u>Fabricated</u> column gives the fraction of repetitions whose output contains variants absent from the provided list of candidates.

**Supplementary Table S16:**
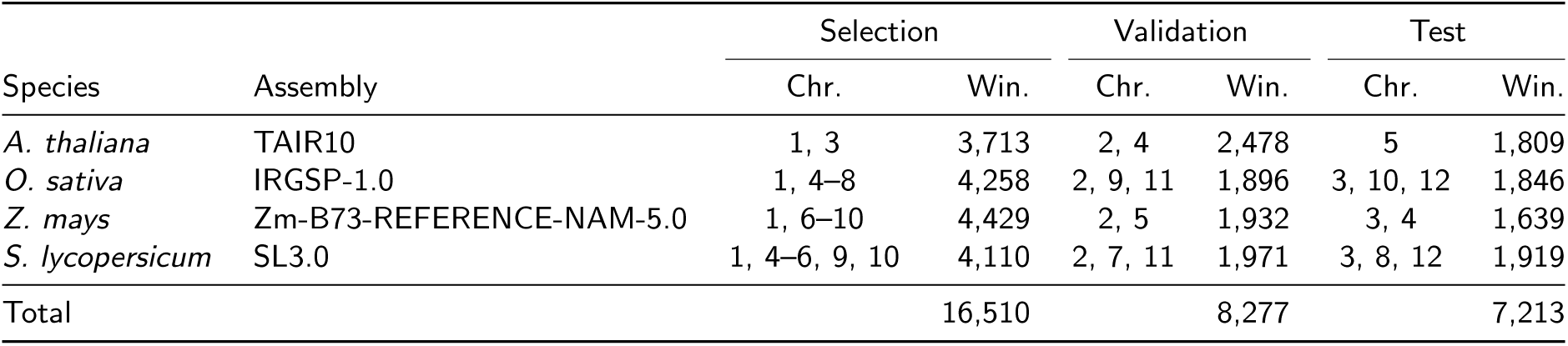
Splits of annotated corpus for interpretability studies. The corpus contains 32,000 coordinate-anchored windows, 8,000 per species, over four annotated reference genomes. Windows are split by chromosome within each species, so no chromosome contributes to more than one split. The reported feature scores are measured on the test split only (Section 4.8). Chromosome numbering follows the Ensembl Plants annotation of each assembly.

**Supplementary Table S17:**
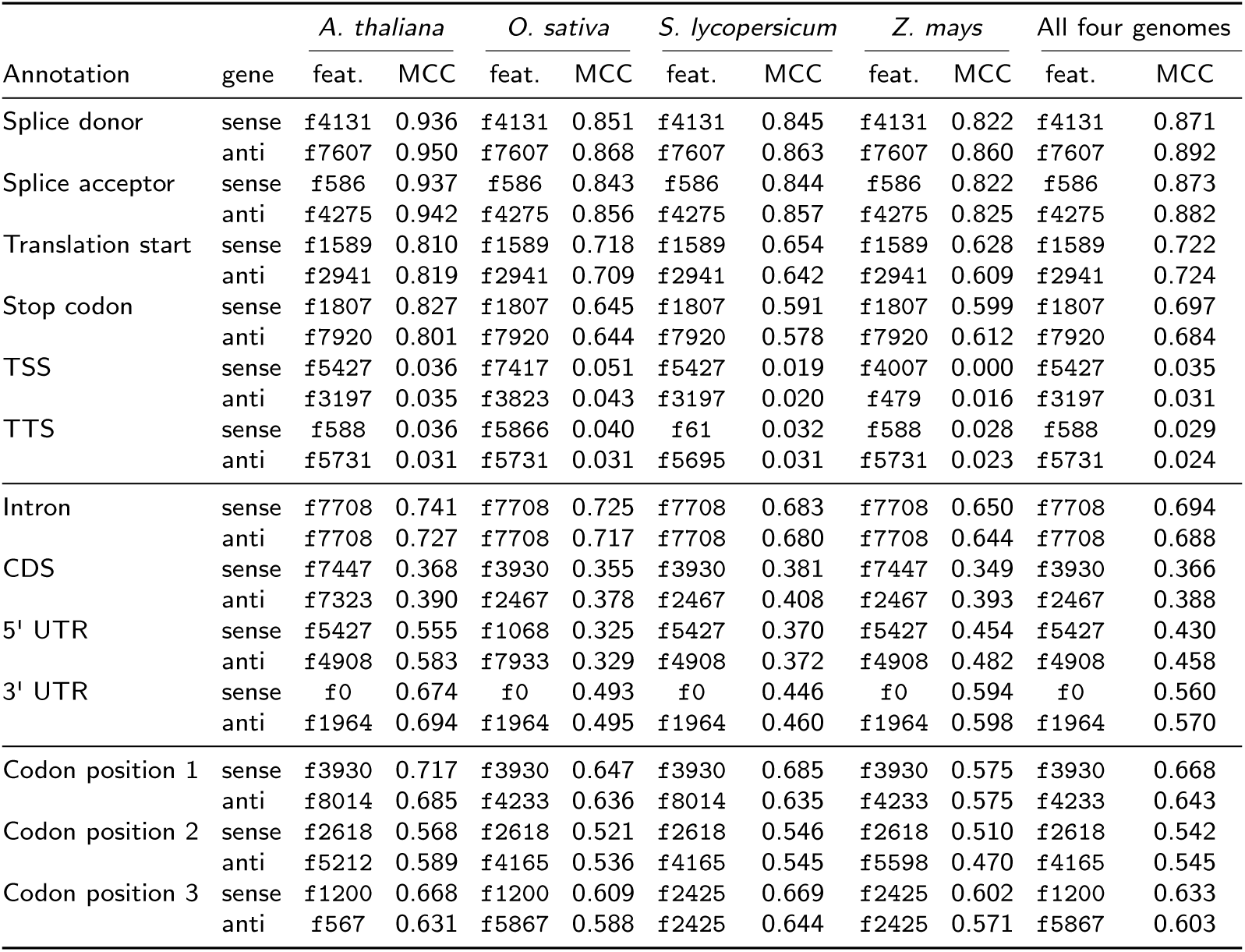
Best SAE feature per gene annotation class and its Matthews correlation coefficient for 4 species. <u>Sense</u> rows score the genes that run in the direction the model reads the window; <u>anti</u> rows score the genes that run against the reading direction.

**Supplementary Table S18:** Botanic1-S attributes contribution to known motif sequence, and not to a motif that is neither enriched nor depleted in accessible chromatin. In silico mutagenesis over 50 held-out Arabidopsis peak windows per motif, each with exactly one motif occurrence, for two functional motifs (HY5, TCP22) and a negative control (HHO3). *z* is the mean per-window contribution at the motif’s bases minus its flanking bases, in flank standard deviations; “> 0” is the fraction of windows above zero; enrichment is the peakversus-background occurrence ratio. *p* is from a two-sided Wilcoxon signed-rank test on the paired motif-to-flank differences (Section 4.5.6).

| Motif | Enrichment | Count head | | Profile head | | $p$ (count) |
| --- | --- | --- | --- | --- | --- | --- |
| | | $z$ | $> 0$ | $z$ | $> 0$ | |
| HY5 (G-box) | 15.4× | +1.87 | 96 % | +0.61 | 64 % | $1.6 \times 10^{-9}$ |
| TCP22 | 9.0× | +2.08 | 78 % | +3.33 | 94 % | $1.9 \times 10^{-6}$ |
| HHO3 (control) | 1.00× | -0.18 | 20 % | -0.07 | 32 % | $2.0 \times 10^{-5}$ |

**Supplementary Figure S38.**
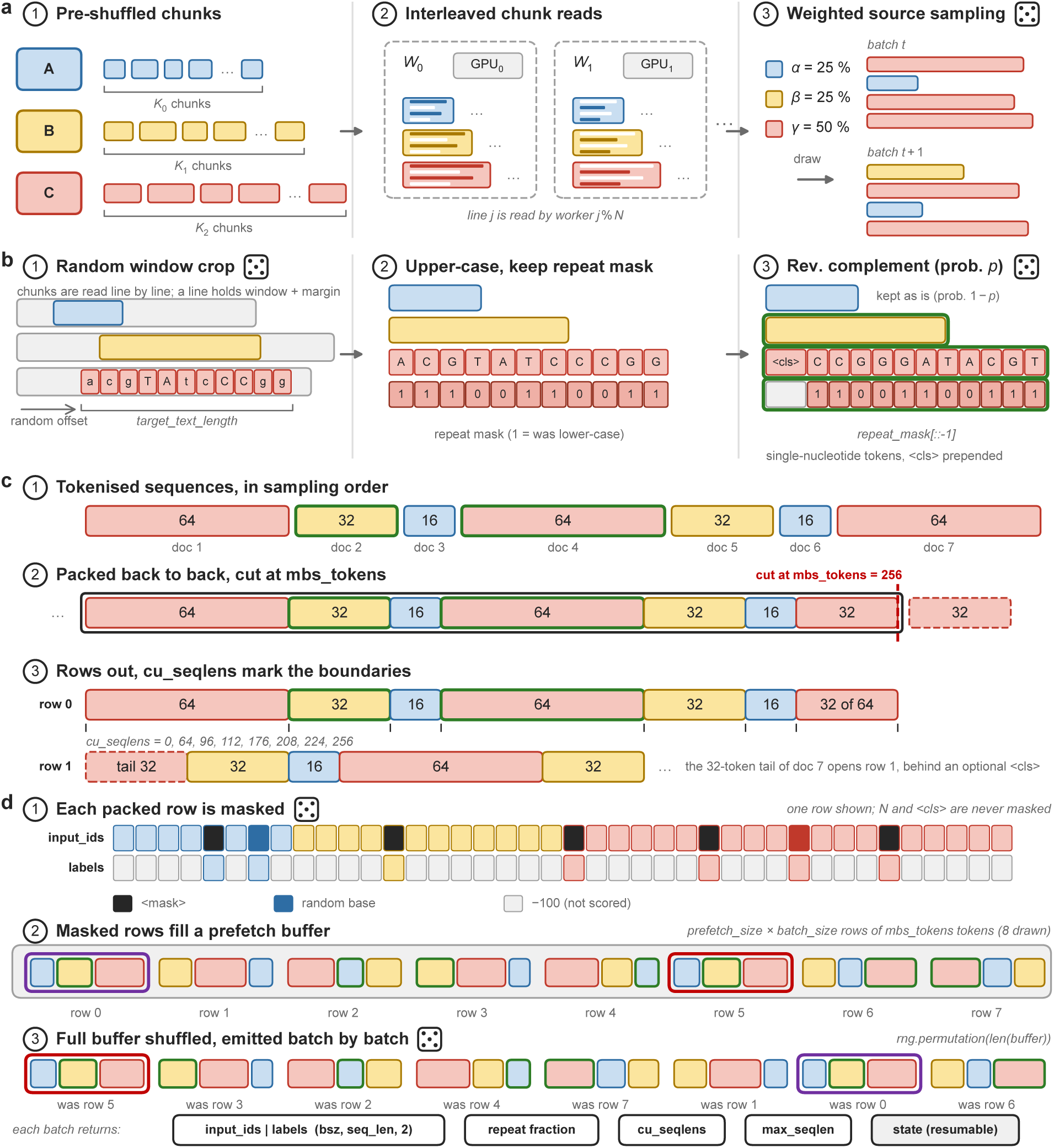
The four stages of batching. Training loader; a die marks a random operation and a green outline marks a reverse-complemented sequence. **a**, Every source is a set of pre-shuffled JSONL chunks (*Ks* per source); chunks are distributed to the data-parallel workers *W_i_*, one per GPU, we colour each line when they are read by the current worker. Each worker draws samples from a source using the mixture weights in this case with three sources (*α*, *β*, *γ*), so a batch mixes sources. **b**, Per-sequence transforms: a start offset is drawn at random. The sequence is upper-cased with its soft-mask pattern stored as a binary mask, and with probability *p* the pair is reverse-complemented before undergoing single-nucleotide tokenisation. **c**, The packed flat strategy writes tokenised sequences back to back into chunks of mbs tokens tokens. As an example here, Document 7 representing a genomic sequence, does not fit, so it is cut and its 32-token tail starts the next chunk, with an optional <cls> token injection at the start of the tail; Cumulative sequence lengths required by the varlen kernels record the boundaries of documents (genomic sequences) within a chunk. **d**, Circled numbers give the order of operations. Rows are masked as they fill a prefetch buffer of here 32 batches. We use black for <mask> and darker shades for a random base, with unscored positions labelled with *−*100 (grey), which is the pytorch convention for values to ignore; the full buffer is then shuffled (two rows tracked in purple and red) and emitted one batch at a time. The loader returns input identifiers and labels, the per-position repeat fraction, the cumulative sequence lengths, as well as the longest sequence in the batch (required by varlen kernels), and its own resumable state. Numbers inside the panels are illustrative.

**Supplementary Figure S39.**
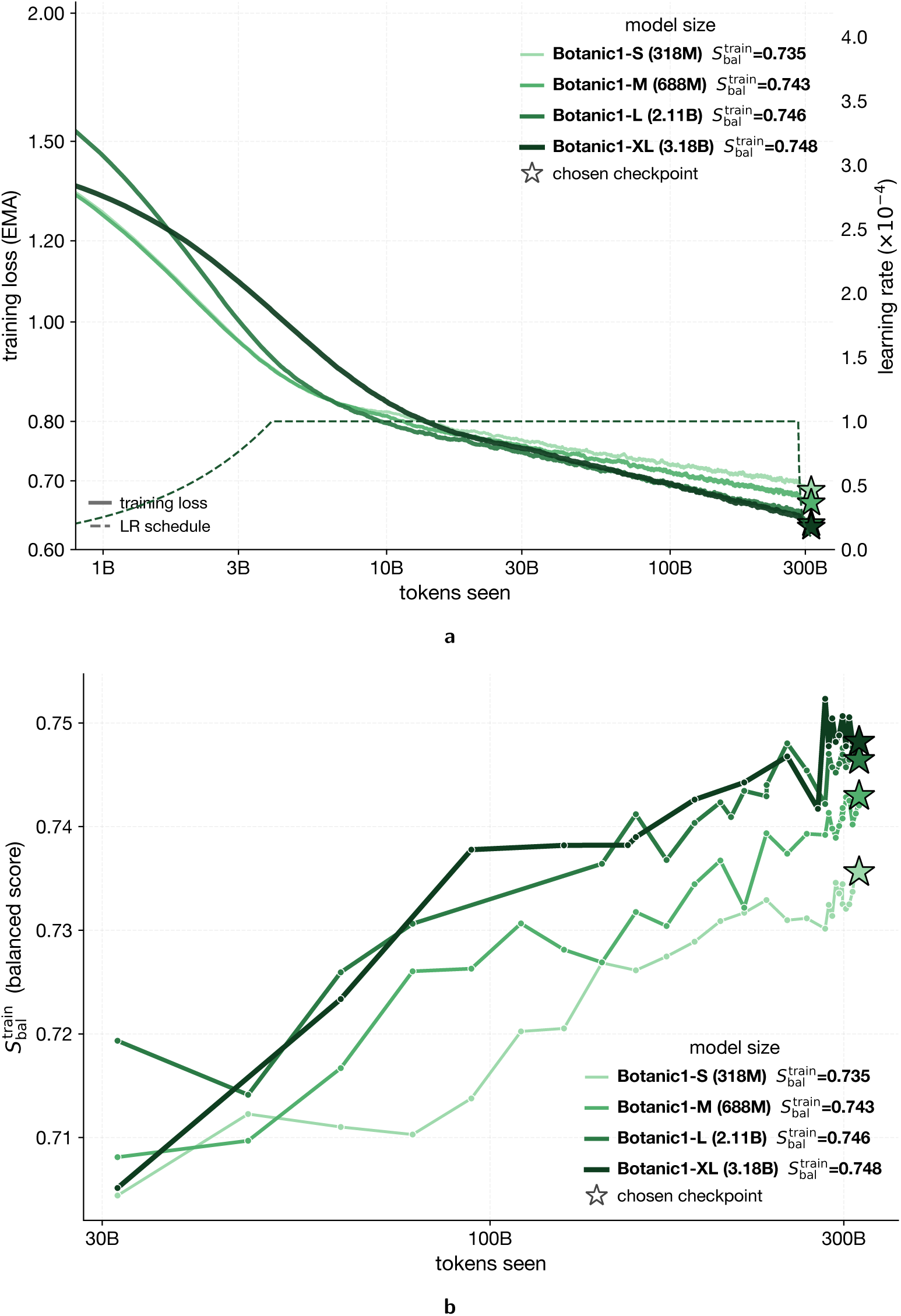
Scaling of the Botanic bidirectional-Mamba2 family. **a**, Training loss and **b**, the species-balanced score S^train^, both against tokens seen, for Botanic1-S (318M), Botanic1-M (688M), Botanic1-L (2.1B) and Botanic1-XL (3.2B). The dashed curve in **a** is the shared learning-rate schedule (right axis); its decay starts near 283B tokens. Stars (*) mark each model’s checkpoint at the target 314.6-billion-token budget: at that matched budget 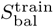 increases monotonically with size, 0.735 (Botanic1-S) < 0.743 (Botanic1-M) < 0.746 (Botanic1-L) < 0.748 (Botanic1-XL). The trajectories and the legend report 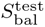.

**Supplementary Figure S40.**
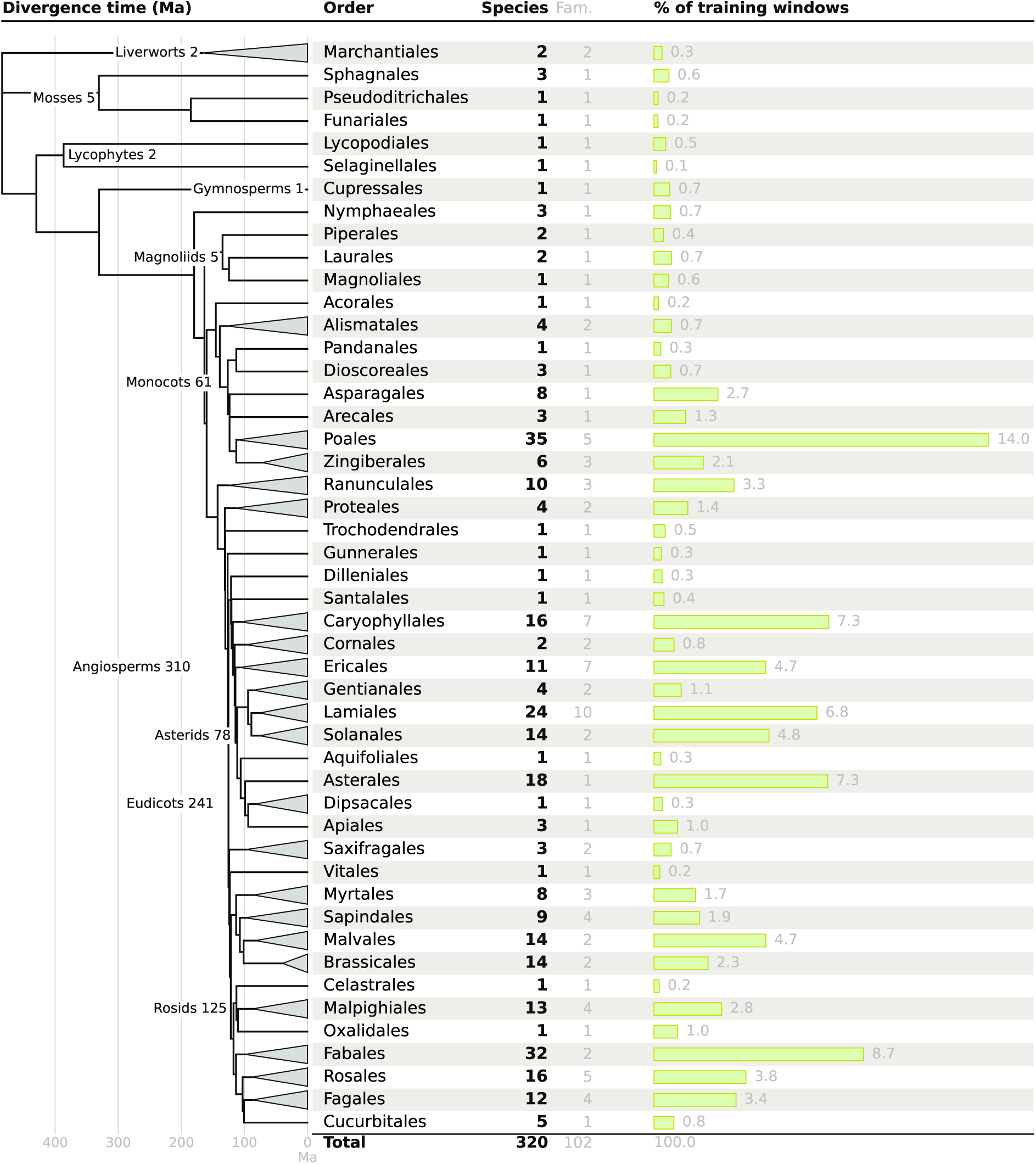
Phylogenetic composition of the 8 kbp pre-training corpus. Each row represents an order contributing at least one species, with its number of species and families and its share of the 7,207,506 8 kbp training windows. Clade labels indicate the total number of sampled species within each clade. The divergence time is retrieved from the TimeTree 5 family-level tree [117], pruned to the 102 families represented in the corpus and collapsed by order. Wedges span the sampled families within each order. Counts include the 320 training species that contribute at least one window; held-out species and two training species left without windows after filtering are excluded.

**Supplementary Figure S41.**
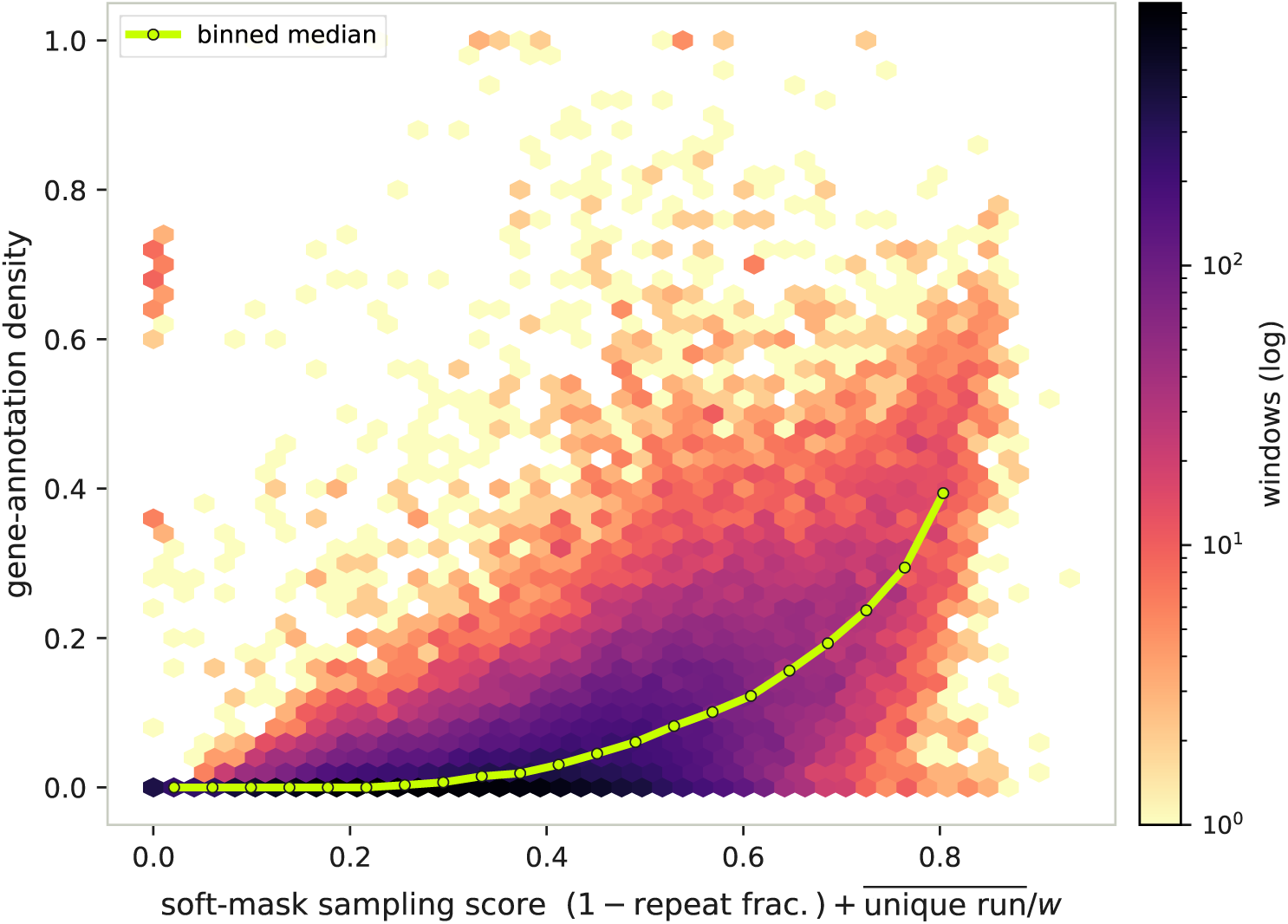
Gene-annotation density against soft-mask derived sampling score. Geneannotation density versus the long-context sampler’s soft-mask score (one minus the window’s repeat fraction plus its mean unique-run length, normalised by window size) for 45,780 non-overlapping 128 kbp windows pooled across 12 soft-masked plant genomes (mean repeat fraction 0.41–0.82). Gene density is correlated with the score (Spearman ρ = +0.63; lime line: binned median).

**Supplementary Table S19:** Pre-training configuration of the Botanic family. Sequences per step is micro batch accumulation replicate shard, and tokens per step is sequences per step × the L = 8,192-token row length; it overestimates what the loss sees, as only 15% of positions are masked and included in the loss. Everything else, objective and schedule included, is shared across the four sizes: all four run on the same 314.6B-token horizon and every reported checkpoint is fully decayed. Throughput is the median training throughput logged over the whole horizon; training time is the wall-clock time spent training, excluding queue time and the segments discarded after pre-emptions, and GPU-hours is time multiplied by the number of GPUs. Model FLOPs utilisation (MFU) is the achieved training FLOP rate per GPU, F FLOP per token × tokens per second per GPU with F from Section 4.1.4, divided by the 2.25 PFLOP/s dense BF16 peak of the B200.

|  | Botanic1-S | Botanic1-M | Botanic1-L | Botanic1-XL |
| --- | --- | --- | --- | --- |
| Model dimension | 1,024 | 1,024 | 1,536 | 1,792 |
| Layers | 24 | 52 | 72 | 80 |
| Parameters | 318M | 688M | 2.1B | 3.2B |
| Micro batch × accumulation | $4 \times 6$ | $4 \times 3$ | $2 \times 6$ | $2 \times 6$ |
| Data parallel (replicate × shard) | $1 \times 8$ | $2 \times 8$ | $2 \times 8$ | $4 \times 8$ |
| Sequences per step | 192 | 192 | 192 | 384 |
| Tokens per step | 1.573M | 1.573M | 1.573M | 3.146M |
| Steps to horizon | 200,000 | 200,000 | 200,000 | 100,000 |
| Horizon (tokens) | 314.6B | 314.6B | 314.6B | 314.6B |
| Reported checkpoint (tokens) | 314.6B | 314.6B | 314.6B | 314.6B |
| NVIDIA B200 GPUs | 8 | 16 | 16 | 32 |
| Throughput (tokens/s) | 1.21M | 1.13M | 0.45M | 0.70M |
| Training time (hours) | 77 | 83 | 202 | 128 |
| GPU-hours | 616 | 1,334 | 3,228 | 4,103 |
| Model FLOPs utilisation | 15.3% | 15.5% | 17.9% | 20.7% |

**Supplementary Table S20:** Pre-training data artefacts by context length.

| Window (bp) | Species (train + held out) | Training windows | Effective training base pairs |
| --- | --- | --- | --- |
| 8,192 | 326 (322 + 4) | 7,207,506 | 59,043,889,152 |
| 16,384 | 285 (282 + 3) | 4,793,670 | 78,539,489,280 |
| 32,768 | 273 (270 + 3) | 3,542,847 | 116,092,010,496 |
| 65,536 | 261 (258 + 3) | 1,677,787 | 109,955,448,832 |
| 131,072 | 238 (235 + 3) | 783,700 | 102,721,126,400 |

**Supplementary Figure S42.**
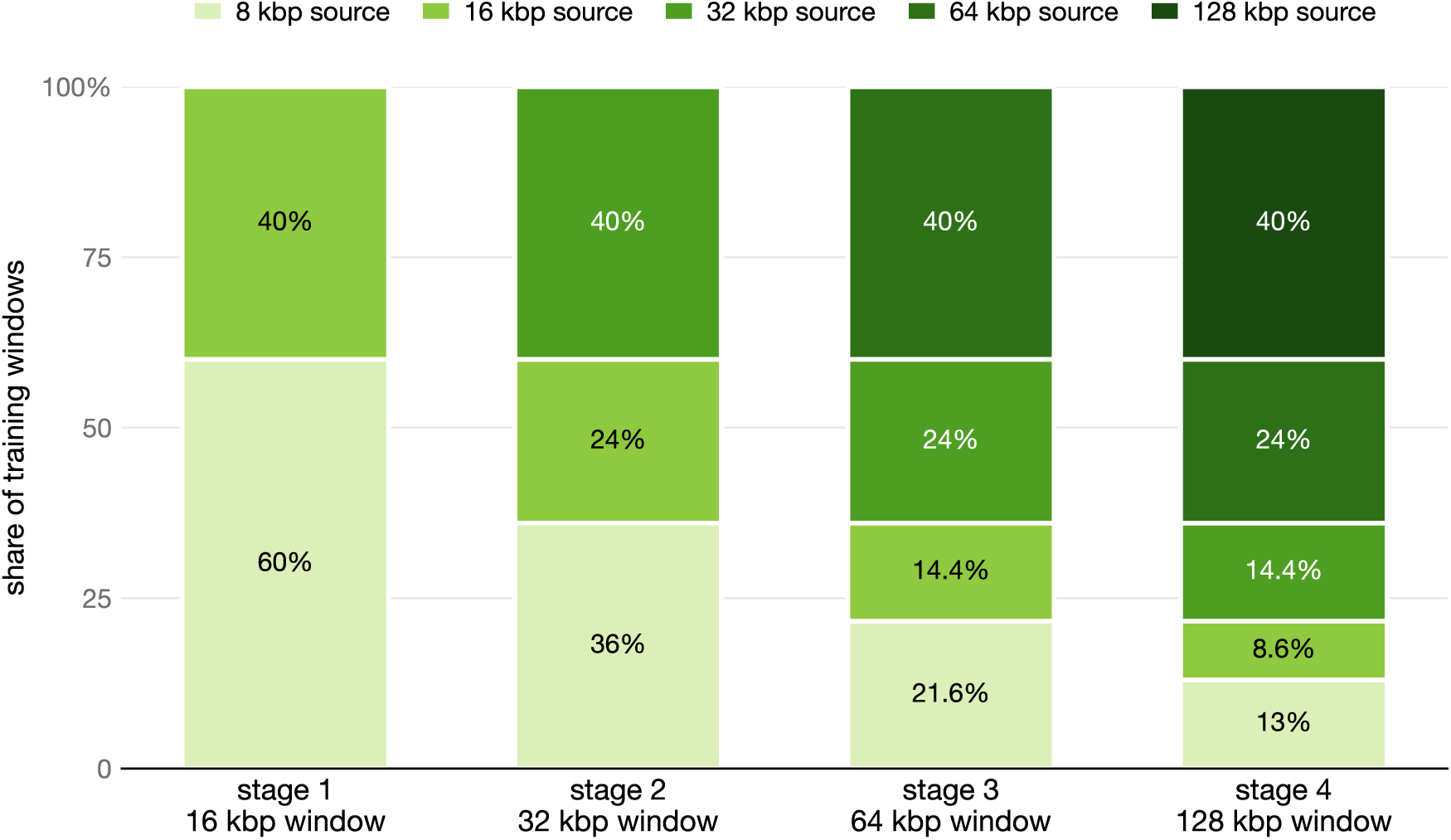
Window-length composition of each context-extension stage. Every stage trains on a static mixture over all window lengths built so far, shaded light to dark with increasing source window length. 40% of windows are sampled from the length newly introduced at that stage and the remaining ones from the already introduced sources, whose weights are rescaled to sum to 60%. The contribution of any source to the mixture then decays geometrically after it has been introduced. The token mixture is consequently more concentrated on the newest length, because a source’s share of tokens is its window share weighted by window length: at the 128 kbp stage the 40% of windows that are 128 kbp long represent 69.6% of total tokens, while the 8 kbp window share of 13% makes up only 1.4% of total tokens.

**Supplementary Figure S43.**
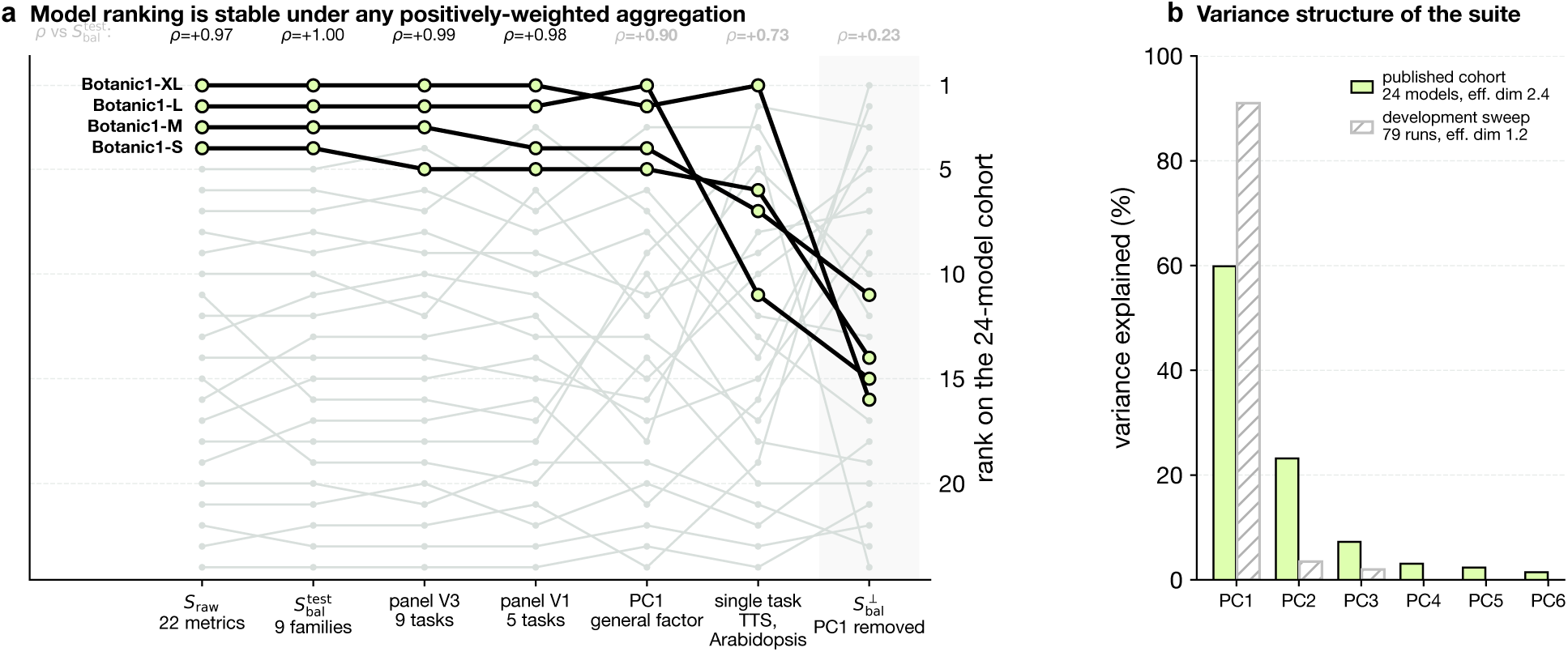
The leaderboard does not depend on how metrics are aggregated. **a**, Model ranking under seven different ways of aggregating the metrics into one scalar: the unweighted mean of all 22 metrics (S_raw_), the capability-balanced score of Equation (1) (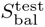), a family-balanced nine-task panel, a greedy five-task panel, the first principal component, the single most discriminative metric, and the nine-task panel with the first principal component projected out (S_bal_*^⊥^* , shaded). Botanic models are highlighted; grey lines are competitors. **b**, Variance explained per principal component between the panel of competitor models vs within different development runs using the BiMamba2 architecture.

**Supplementary Figure S44.**
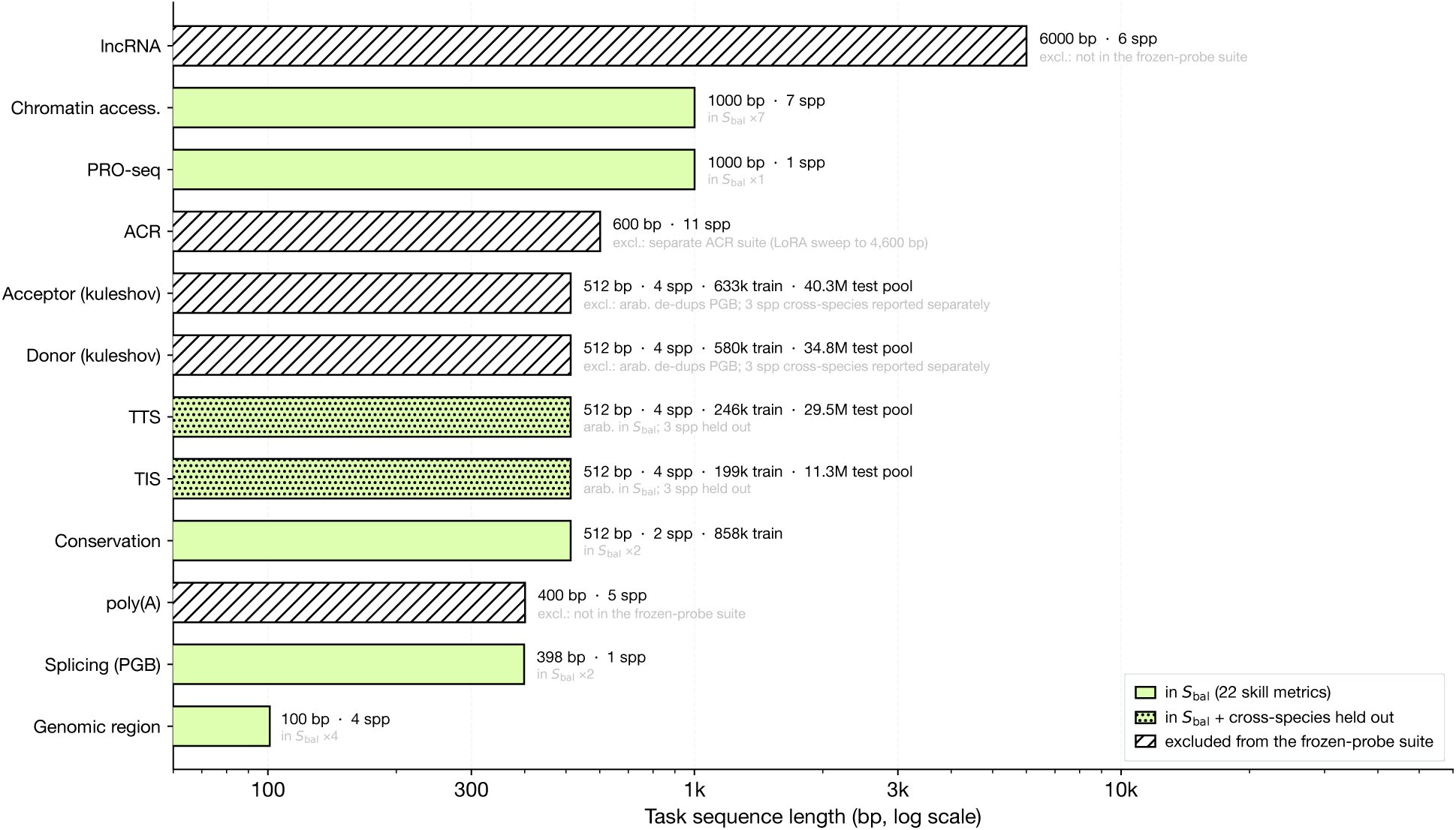
Frozen-probe tasks, ranked by sequence length. Every classification task suitable for frozen model evaluation, ordered by its sequence length (log x). Green bars are the 22 task metrics that enter *S*_bal_, dotted for TIS and TTS; hatched bars are set aside, each labelled with the reason for not including it: absence from the frozen-probe suites (poly(A), lncRNA), the splice overlap with PGB, and ACR’s separate suite. The included lengths span 100 to 1,000 bp.

## Footnotes

‡ At same amount of trainable parameters

* Supported also by PastDB [79] junction data (https://pastdb.crg.eu/gene/AT1G65170@araTha10). From the donor side, in the six samples where this locus is quantified, 24 reads cover the TAIR10 donor position at chr1:24,210,073 and none splice there. From the acceptor side, while the donor at chr1:24,210,127 is absent from the vast-tools event library and cannot be scored directly, some of the reads reaching the shared acceptor are flagged as splicing from an unannotated donor.

